# Identification of an Endothelial Senotype Predicted by Ion Channel Expression in Aged Murine Perigonadal Adipose Tissue

**DOI:** 10.64898/2026.10.26.761925

**Authors:** Ann V. Hertzel, Syed Ali, Alessandro Bartolomucci, Jinhua Wang, Laura J. Niedernhofer, David A. Bernlohr, Christina D. Camell, Paul D. Robbins

## Abstract

Adipose tissue plays a key role in metabolic health, contributing to systemic dysfunction upon obesity or aging. A major driving factor of adipose-driven dysfunction is the accumulation of senescent cells (SnCs), but identification and characterization of specific SnCs is limited. To identify age-dependent SnCs, we performed snRNA-Seq on young and old adipose using two independent analytical strategies. The first, a SenMayo-based approach and the second, dependent on SEAC guidelines and Pearson correlation identified senescent nuclei in endothelial clusters. Subsequent subclustering and cell-to-cell communication analyses identified endothelial capillary nuclei that overexpress macrophage migration inhibitory factor (Mif), which exemplifies the specific characteristics and microenvironment of SnCs. Finally, rank analysis identified ion channel transport genes as the most predictive of the SnCs. Overall, these results highlight new defining features of senescence in endothelial adipose tissue and provide novel age-dependent biomarkers.

## Introduction

Aging leads to a decline in organ function due to a lack of adaptation and repair of molecular and organelle damage with white adipose being one of first tissues to show age-dependent molecular changes^1–4^. Upon aging, adipose redistributes from the subcutaneous depots to the visceral depots, with a changing repertoire of cellular composition, contributing to the increase in metabolic pathologies^4–12^. Most cell types within adipose tissue show effects of aging including increased inflammation, fibrosis that leads to mechanical stress, and metabolic dysfunction^4, 13–16^. Overall, the dysfunction of these cells affects the tissue environment resulting in a chronic, sterile low-grade inflammatory state, that remains unresolved^1, 16, 17^. These conditions are linked to many pathologies, including diabetes, cardiovascular dysfunction, some cancers and dementia^18–20^.

Senescence is a cell fate that is induced by cellular stress, resulting in replication arrest with a repression of apoptosis that allows SnC persistence^21–26^. These cells have altered morphology, are highly metabolic, have increased lysosomal activity and cause dysfunction in neighboring cells due to an elevated inflammatory secretome (senescence-associated-secretory-phenotype; SASP). Although pathogenic in aging, senescence is an important transient component for proper tissue remodeling in wound healing^27, 28^ and is an important biological process associated with cancer prevention^29^. Acutely senescence is beneficial, but SnCs must be cleared to prevent chronic inflammatory cascades and enhanced cellular dysfunction. The immune system efficiently clears SnCs in young organisms, however aging impairs the immune system leading to SnC accumulation^30–32^. Counterintuitively, SnCs can sometimes facilitate tumor growth due to the secretion of inflammatory cytokines^27–29^.

SnCs are highly heterogenous and thus defining biomarkers has been challenging^33–37^. In this work, we utilized single nuclei (sn) sequencing of murine young and old female perigonadal adipose tissue to identify SnCs and have identified a subset of endothelial cells that differentially expressed senescent markers including the cell cycle regulators *Cdkn1a/b*, anti-apoptotic signals, ECM components and modifying enzymes and inflammation. Predicted cell-cell interactions including increased macrophage migration inhibitory factor (Mif) secretion from this endothelial senotype that signals inflammation and NF-κB activation. The most predictive genes of this senotype revealed novel senescent biomarkers including an increased expression of a variety of ion channel genes.

## Results

### Single nuclei sequencing of female perigonadal adipose tissue

To identify age-dependent SnCs in female visceral murine adipose, we collected perigonadal adipose from young (6-month) and old (24-month) C57BL/6J mice (n=3). The average body weights were 23 +/-0.42 g and 29.1 +/-0.26 g from the young and old mice, whereas the average perigonadal weights were 259 +/-34 mg and 386 +/-42 mg from the young and old mice, respectively. From frozen perigonadal adipose tissue, nuclei were isolated and subjected to single nuclei sequencing using the 10X Genomics Chromium 5’ capture platform (**Fig. 1A).** Following quality control, we analyzed a total of 42,327 nuclei from old adipose and 39,265 nuclei from young adipose (**Ext Fig. 1A-D)**. As an overall metric of this dataset, we used UCell to create an inflammatory score, based on the genes of the KEGG inflammation pathway^38^. The resulting curve from the old samples was shifted to the right as compared to the young (**Ext Fig. 1E**), consistent with increased inflammation upon aging^39, 40^.

**Figure 1.**
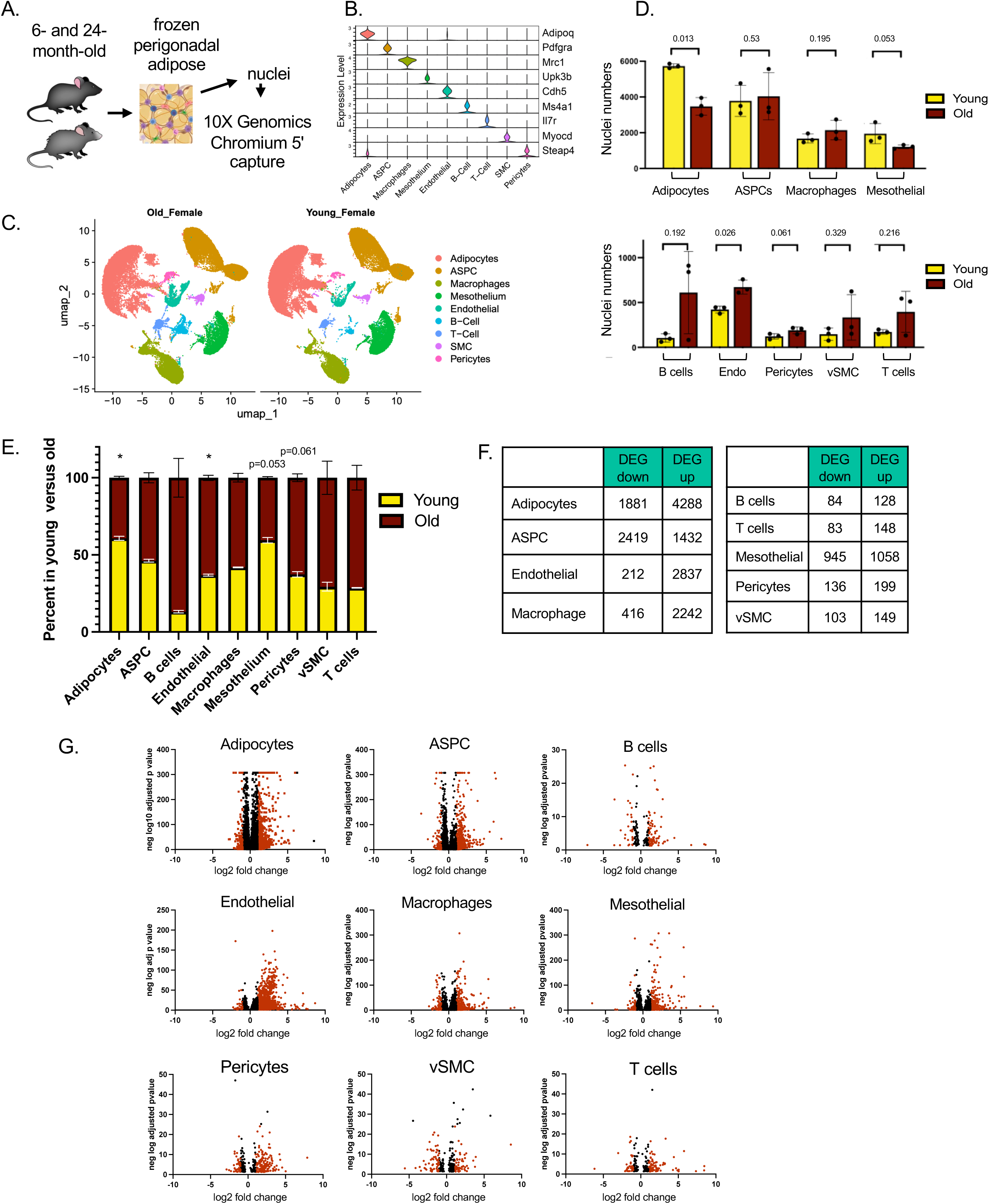
Single nuclei sequencing of female perigonadal adipose tissue from old and young mice. **A**. Schematic of snSEQ experiment on female perigonadal adipose tissue from old (24-month) and young (6-month) C57Bl/6J mice; n=3/group. **B.** Single marker expression for identification of the nine main clusters. **C.** UMAP of old and young single nuclei data. **D.** Nuclei numbers per cluster from old and young adipose; p-values calculated using Student’s t-test. **E.** Percent nuclei from old or young within each cluster. **F.** Number of differentially expressed genes (DEG), separated by directionality in the old compared to the young mice. **G.** Volcano plots of significant DEG: greater than 2-fold (red) and less than 2-fold (black).

Unsupervised clustering identified of nine major clusters representing the major cell types of murine adipose tissue as confirmed by expression of cell-specific markers (**Fig. 1B,C**) and of top genes in each cluster (**Ext Fig. 1F**). Comparison of the nuclei numbers for each cluster indicated a typical adipose tissue cellular composition with the expected age-dependent decrease in adipocyte numbers, stable numbers of ASPCs and an increase in endothelial cells. Vessel support cells and immune cells also trended towards an increase (**Fig. 1D,E**).

Differentially expressed gene (DEG) analyses showed significantly more genes upregulated than downregulated, excluding the ASPCs, although volcano plots show striking changes in gene expression in all cell types (**Fig. 1F,G**). Transcription factor (TF) analyses indicated significant age-dependent alterations including ∼10-20 unique TFs. Although these changes are complex, several themes were revealed. Many cell types have predicted changes in redox and inflammatory TF, such as HIF1α in adipocytes and HIF3α in mesothelial cells, as well as several NF-κB subunits increasing in endothelial cells, macrophages, B cells and T cells. Furthermore, these same cell types all have increases in interferon-regulated TF. Epigenetic modifiers are also in the top changes, with specific demethylase or deacetylase present in young, but not old samples (**Ext Fig. 1G**).

KEGG analyses on specific cellular cluster DEG (separated by increased or decreased gene expression) from old and young nuclei revealed the top pathways changed in nuclei from old mice as compared to young (**Ext Fig. 1H**). In most cell clusters from the old mice, cellular signaling pathways are downregulated, consistent with a reduction in normal cellular processes that each cluster/cell type performs. Alternatively, pathways that are generally elevated in the aged samples are inflammatory pathways, as demonstrated in adipocytes, ASPCs, macrophages and mesothelial cells. B cells from old mice had pathways suggesting increased phagocytosis along with inflammation, whereas macrophage and mesothelial cells had increased lipid metabolism and atherosclerosis pathways. T cells showed pathways elevated in infection, carcinogenesis and diabetes, whereas pericytes and vSMC had the lowest number of pathways altered (**Ext Fig. 1H)**.

### Identification of senescent nuclei: SenMayo panel and SEAC with similarity

To identify nuclei with senescent characteristics, we utilized two orthogonal approaches. The first one employed the SenMayo gene list which identifies genes in SnCs across tissue and species independent of p16^Ink4a^ and p21^Cip1^ expression^41^. For this approach, we identified nuclei that co-expressed the largest number of genes from this list. We first mapped the distribution of the number of SenMayo genes per nuclei, colored by cluster, and focused on those in the upper quartile (25+ SenMayo genes/nuclei; **Fig. 2A** boxed). This resulted in 451 nuclei from the old (range 25-49 SenMayo genes) and 236 nuclei (range 25-41 SenMayo genes) from the young adipose. When comparing the major cell types, the greatest number of nuclei in the upper quartile of SenMayo genes expressed per nuclei were adipocytes. However, the number of young and old nuclei showed a modest age-dependent increase, with limited DEG (n=190 and n=272, respectively; **Fig. 2B**). Endothelial, ASPC and mesothelial populations were the next most abundant, while the remaining cell types had few nuclei in this upper quartile (≤10) **(Fig. 2B)**. Overall, the fold difference in old vs young nuclei in the upper quartile of SenMayo gene expression was greatest in the endothelial cells (**Fig. 2C**). These SenMayo positive nuclei make up 0.39% of the young and 2.4% of the old adipose endothelial nuclei.

**Figure 2.**
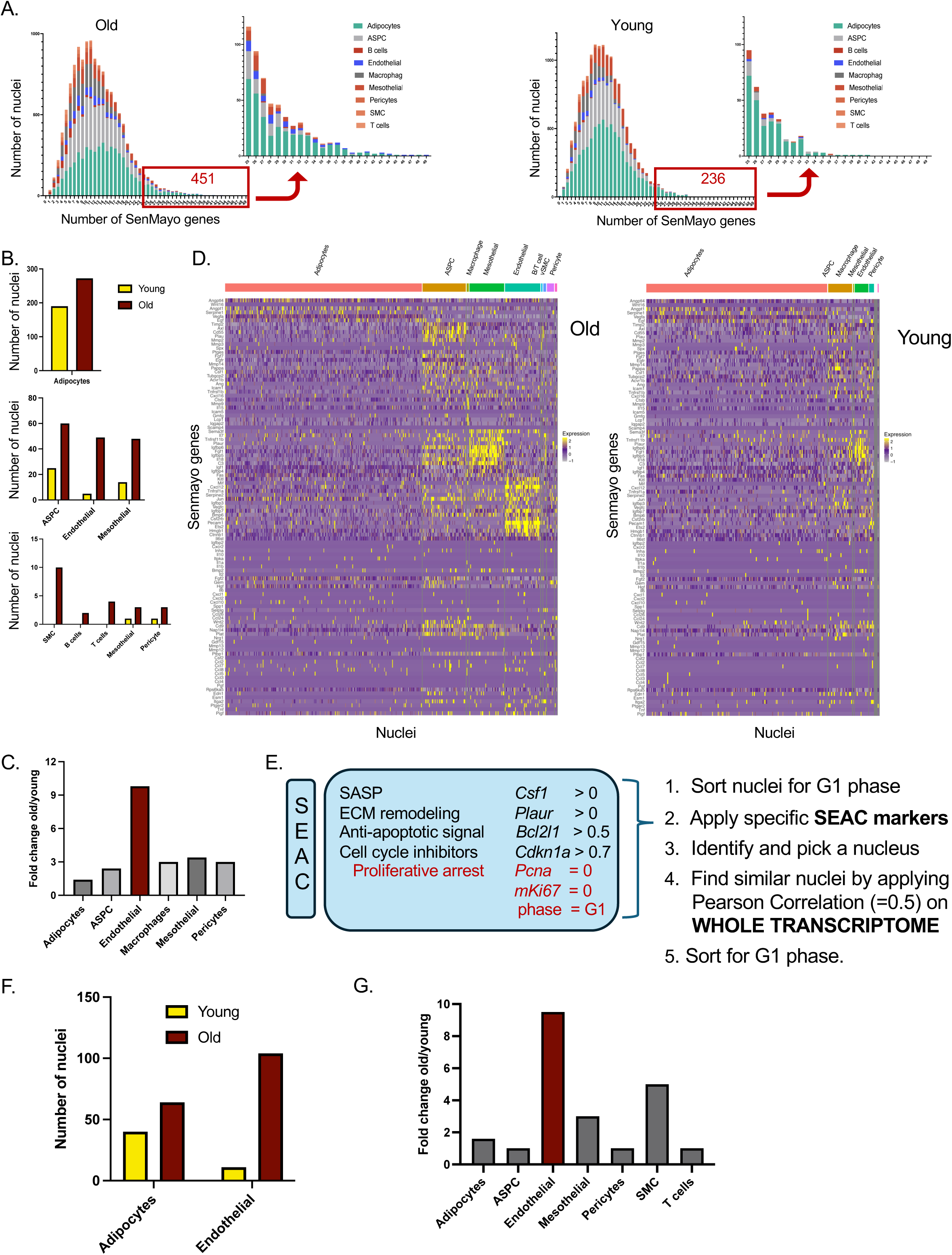
Identification of senescent nuclei using SenMayo or SEAC strategies. **A.** Distribution plot of the number of nuclei that express the specific number of SenMayo genes from snRNA-Seq of old and young adipose. Red box indicates the upper quantile of nuclei (≥25 of the 125 gene panel) and inset highlights these. **B.** By cell type, the number of nuclei identified as senescent from the SenMayo strategy. **C.** Fold change (old to young) in the number of nuclei identified as senescent via the SenMayo strategy. **D.** Heat map showing expression of each SenMayo gene (rows) within each nuclei (columns) from the upper quantile, grouped by cell type (top); Left panel is from old samples and right panel is from young samples. **E.** SEAC strategy consisting of four canonical senescent cell features and requiring proliferative arrest features (left). Specific gene expression utilized in SEAC strategy (middle). SEAC strategy to identify senescent nuclei (right). **F.** Number of nuclei identified as senescent via SEAC, by cell type. **G.** Fold change in the number of nuclei identified by SEAC strategy, by cell type.

Next, we examined the abundance of each SenMayo gene expressed in all nuclei in this upper quartile. Heatmaps of SenMayo gene expression per nuclei (SenMayo 25+) highlighted that individual cell types upregulate a unique subset of the SenMayo genes that is highly visible in the mesothelial and endothelial populations (**Fig. 2D**), an indication of the heterogeneity of SnCs in differing cell types.

Our second approach to define SnCs utilized the characteristics of senescence, including proliferative arrest, structural changes, along with pathological secretion of inflammatory molecules^21–26^. For this we used a knowledge-based binning strategy separating senescence features into four major categories: **S**ASP, **e**xtracellular matrix (ECM), **a**nti-apoptotic, and **c**ell cycle arrest (**SEAC**) represented by *Csf1*, *Bcl2l1*, *Plaur*; and *Cdkn1a* (**Fig. 2E**)^22, 42, 43^. Nuclei were queried for expression of all four genes at a minimum threshold of 0.5 and 0.7 of the log normalized expression for *Bcl2l1* and *Cdkn1a*, to capture the higher expressing populations, as density plots of their expression exhibited a multimodal distribution. Finally, we filtered for nuclei in G1 phase, as indicating non-replicating. As these thresholds were intentionally set high, we expected limited positive hits and found two endothelial nuclei. We hypothesized that these are senescent nuclei and proceeded to identify other nuclei that are most similar in their overall transcriptome, independent of the SEAC markers, by applying a Pearson correlation analysis (set to 0.5) of the identified nucleus to the entire transcriptome of all nuclei in the dataset (**Fig. 2E**). This analysis identified 11 endothelial and 40 adipocyte nuclei in the young and 104 endothelial and 64 adipocyte nuclei in the old adipose (**Fig. 2F**). We performed correlation analysis with other SEAC-positive nuclei, finding comparable results. The identified adipocyte nuclei showed a modest age-dependent increase (similar to **Fig. 2B**), whereas the endothelial nuclei were increased 9-fold in the old as compared to the young adipose (**Fig. 2G**). When considering the entire endothelial cell populations, the SEAC strategy identified 0.87% of the young endothelial nuclei and 5.2% of the old endothelial nuclei as being senescent. The two independent computational approaches identified similar numbers of senescent endothelial nuclei using stringent criteria.

### Senescent endothelial cells

To more specifically interrogate senescent endothelial cells, we subclustered all endothelial nuclei into arterial (A), venule (V), capillary (S) and lymphatic (LEC) endothelial subpopulations (**Fig. 3 A,B**). There was a significant increase in the numbers of EndoV (∼2X) and EndoS2 (3X) subpopulations upon aging (**Fig. 3C,D**).

**Figure 3.**
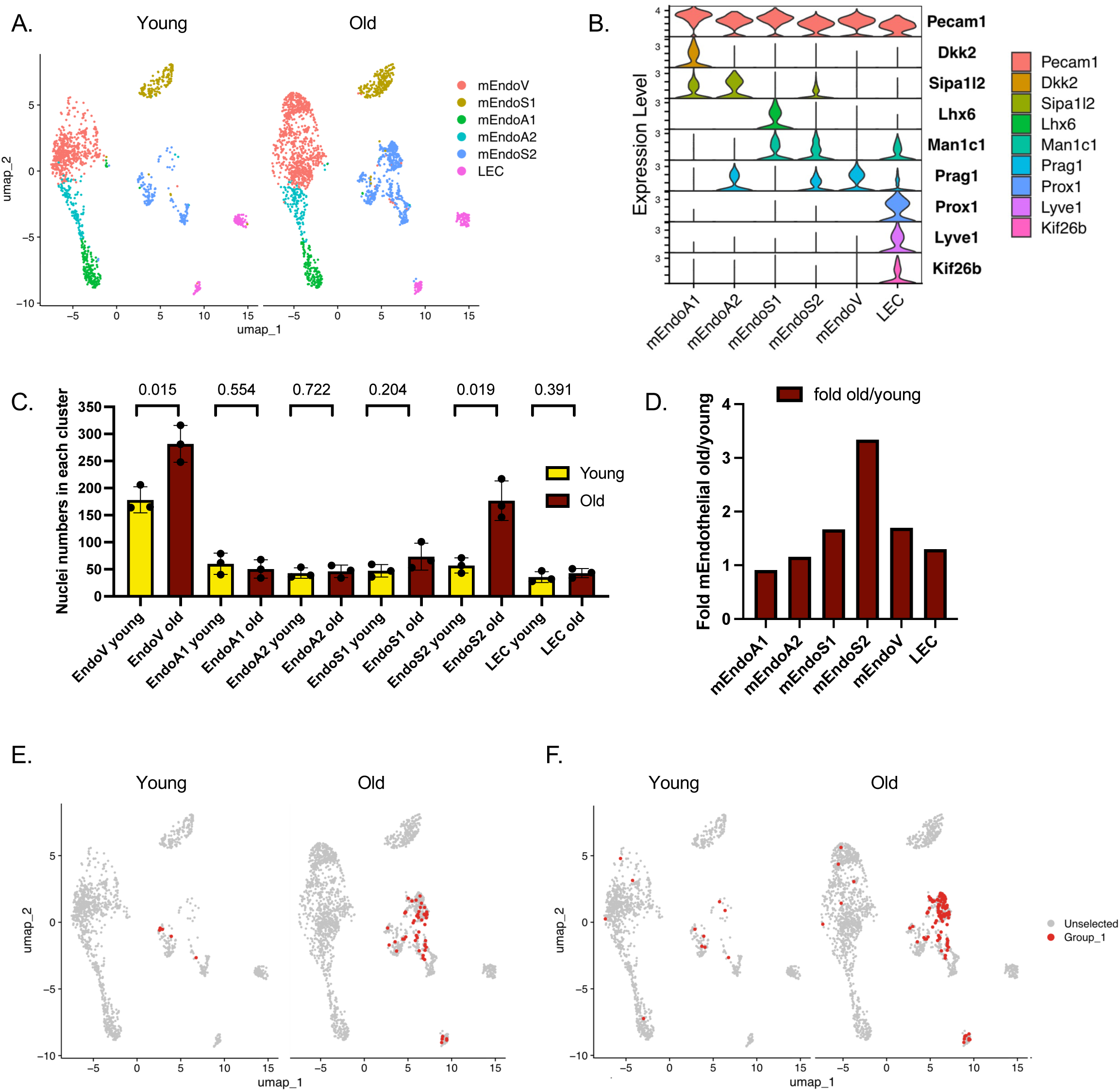
Endothelial cell subclustering and senotype mapping. **A.** UMAP of endothelial subclusters from young and old perigonadal adipose. mEndoV, venule; mEndoS1/2, capillary; mEndo A1/2, arterial; LEC, lymphatic endothelial nuclei. **B.** Violin plots of marker expression for identification of the endothelial cell subclusters. **C.** Nuclei numbers in each endothelial subcluster from young and old adipose tissue. **D.** Fold change (old to young) in nuclei numbers in each endothelial subcluster. **E.** Senescent nuclei (red) identified by the SenMayo strategy mapped onto the endothelial cell subcluster UMAP from old and young adipose tissue. **F.** Senescent nuclei (red) identified by the SEAC strategy mapped onto the endothelial cell subcluster UMAP from old and young adipose tissue.

We next identified the location(s) of the SenMayo-positive and SEAC-identified nuclei within the endothelial subclusters. Most of the senescent nuclei from the SenMayo25+ (**Fig. 3E**) and the SEAC (**Fig. 3F**) strategies resided in the endothelial capillary S2 subcluster, with an 85% overlap of the two strategies. This implicates the capillary population as a reservoir of adipose senescence and suggests the specific microenvironment of these endothelial cells may make them especially vulnerable to the senescent cell fate.

To characterize these senescent endothelial cells, we first determined the DEG (**Fig. 4**). Because of the low number of senescent nuclei identified, we utilized a repeated random down sampling method to compare equal numbers of nuclei in comparison groups, repeating this 100X and summarized the output results. When comparing the old senescent endothelial nuclei to all the young endothelial nuclei, we found 854 and 1,767 genes were differentially expressed in the senescent population identified by the SenMayo and SEAC strategies, respectively, whereas the comparison of the old senescent nuclei to the old non-senescent endothelial nuclei identified 168 and 685 DEG, respectively (**Extended Tables 1-4**). Within the DEG, 16 and 19 of the SenMayo panel genes were significant in the SenMayo and SEAC results, respectively (**Fig. 4A**). Next, we considered each of the four categories from our SEAC strategy. Using proteINside^44^ to predict secreted proteins and matrisomeDB^45^ (a searchable database of ECM) we identified 181 (SenMayo) and 135 (SEAC) DEG that encode secreted proteins (**Fig. 4B, Extended Tables 5,6**) and many alterations in the genes encoding ECM proteins, both within the core matrisome and the matrisome-associated categories (**Fig. 4C**). The anti-apoptotic *Bcl2* and *Mcl1* were significant in the SEAC senotype, whereas *Bcl2* was trending up in the SenMayo strategy **(Fig. 4D)**. And finally, the cell cycle inhibitor, *Cdkn1a*, was significant in both, whereas the SEAC senotype also identified *Cdkn1b* as significantly upregulated (**Fig. 4E)**.

**Figure 4.**
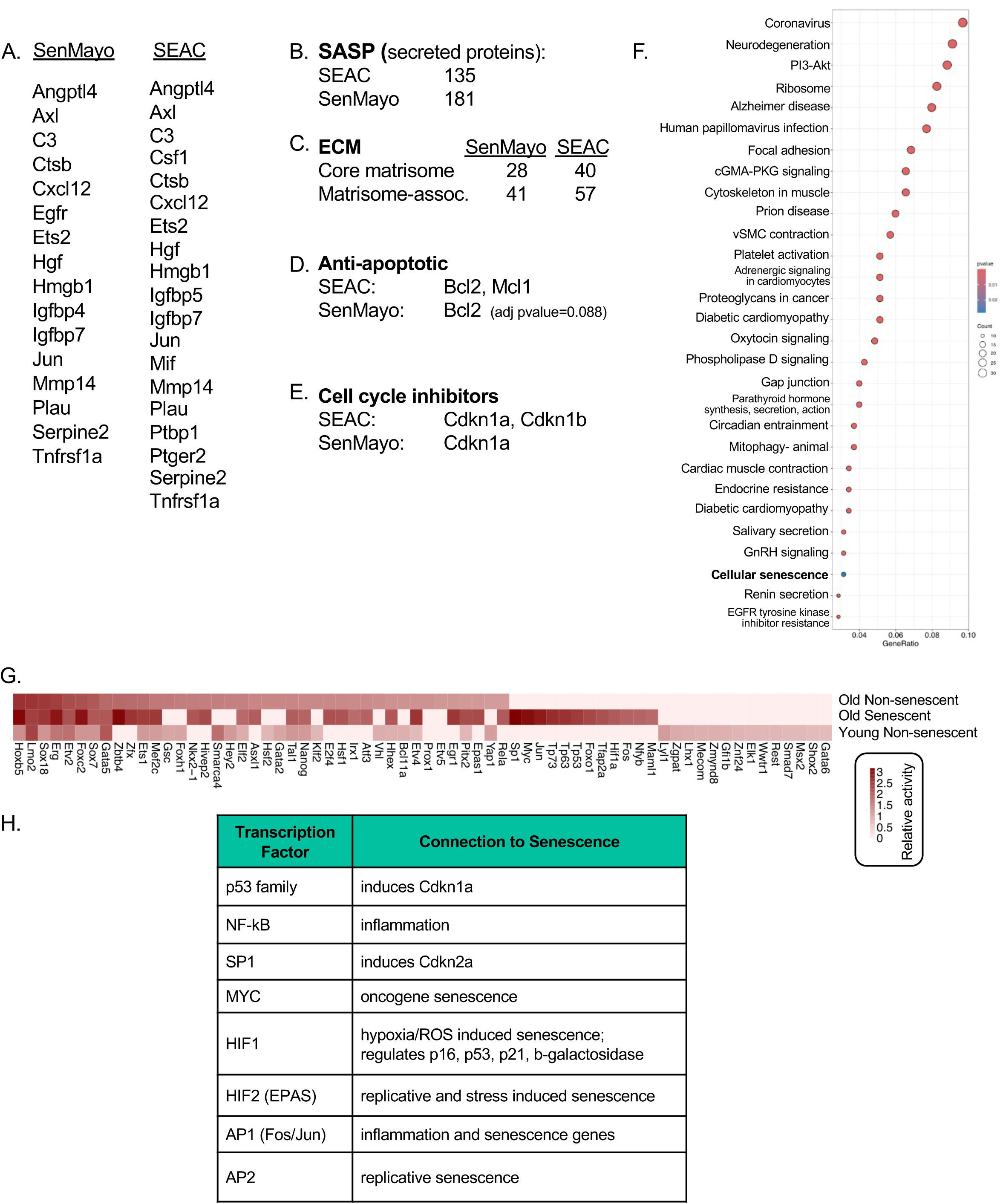
Characteristics of the endothelial senotype. **A.** Significantly upregulated DEG from the SenMayo and SEAC strategies that are part of the SenMayo gene panel. **B.** The number of DEG predicted to be secreted. **C.** The number of DEG altered in the core matrisome and the matrisome-associated genes. **D.** Anti-apoptotic genes upregulated in DEG list. **E.** Cell cycle inhibitor genes upregulated in DEG list. **F.** Kegg pathway of DEG from SEAC strategy. **G.** Top 40 transcription factors whose activity predicts the transcriptome of the old non-senescent, old senescent (SEAC strategy) and young non-senescent nuclei. **H.** The identified TF’s connection to senescence.

KEGG analyses of the DEG from the SEAC-identified senescent endothelial nuclei resulted in pathways consistent with senescence **(Fig. 4F)**. For example, several neurodegeneration and infection pathways were identified consistent with elevated inflammatory genes. Additionally, we identified pathways involved in inducing senescence or increasing with senescence such as p53-or Cdkn1b-dependent hyperactivation of PI3K-AKT^46^, increased ribosome activity, NF-Y regulating cell cycle as seen in replicative senescence^47^, local elevated renin-angiotensin-aldosterone system (RAAS) promoting senescence^48^ and oxidative stress and inflammation. The KEGG GO term cellular senescence was also found, although the overlap of this list with the SenMayo panel is limited.

Next, we predicted the top 40 TF responsible for the transcriptome in the old senescent nuclei, as compared to the young nuclei and old non-senescent nuclei **(Fig. 4G)**. From this, we identified 13 TF whose activity was dramatically increased in the old senescent nuclei compared to both the young endothelial and the old non-senescent endothelial nuclei. Importantly these TF have been shown to have a strong connection to senescence, inflammation, reactive oxygen species, or stress (**Fig. 4H**). Furthermore, this analysis identified 14 TF only in the young endothelial nuclei, largely involved in angiogenesis and survival.

### Interactions of the senescent endothelial cells with other adipose cell types

To interrogate potential interactions of the senescent endothelial cells with other cells, we used analyses of ligand and receptor expression via Cell Chat^49^. When considering the senescent endothelial cells and the main cell type clusters, we identified several incoming ligands and outgoing signals specifically for the senescent endothelial nuclei (**Fig. 5A**). Of particular interest, we identified *Mif*, one of the SenMayo genes, as having a high relative interaction strength and violin plots demonstrated the overexpression of *Mif* in these senescent nuclei. Interestingly, MIF signals to ASPCs via the ACKR3 receptor and to B cells and macrophages via CD74/CD44 receptors^50, 51^ (**Fig. 5B**). These interactions are predicted to inhibit adipogenesis and promote inflammation in ASPCs and B cells/macrophages, respectively^50, 51^. We also found Ceacam interactions with Cd1d1 in adipocytes and Chemerin signaling to adipocytes and macrophages. Also overexpressed in these senescent nuclei, *Lgals9*, *Ntf3*, *Vtn*, *Agrn*, and *Cspg4*, all signal in a fairly broad manner to multiple cell types (**Ext Fig. 2**). Overall, these interactions lead to immune cell regulation (*Ceacam*, *Chemerin*, *Lgals9*, *Itga4* and *Nectin*), migration (*Cspg4*, *Icam2*, *Itga4*, *Kitl*) and cellular/ECM adhesion (*Vtn*, *Agrn*, *Cspg4*, *Icam2*, *Itga4* and *Nectin*). Additionally, we identified three specific receptors in these senescent endothelial nuclei that receive signals via *Vcam* and *Kit*, predominantly from macrophages and *Nectin* from ASPCs (**Ext Fig. 2**). These data are further evidence that the senescent endothelial cells can affect cell-cell communication, cell proliferation, adhesion and inflammation.

**Figure 5.**
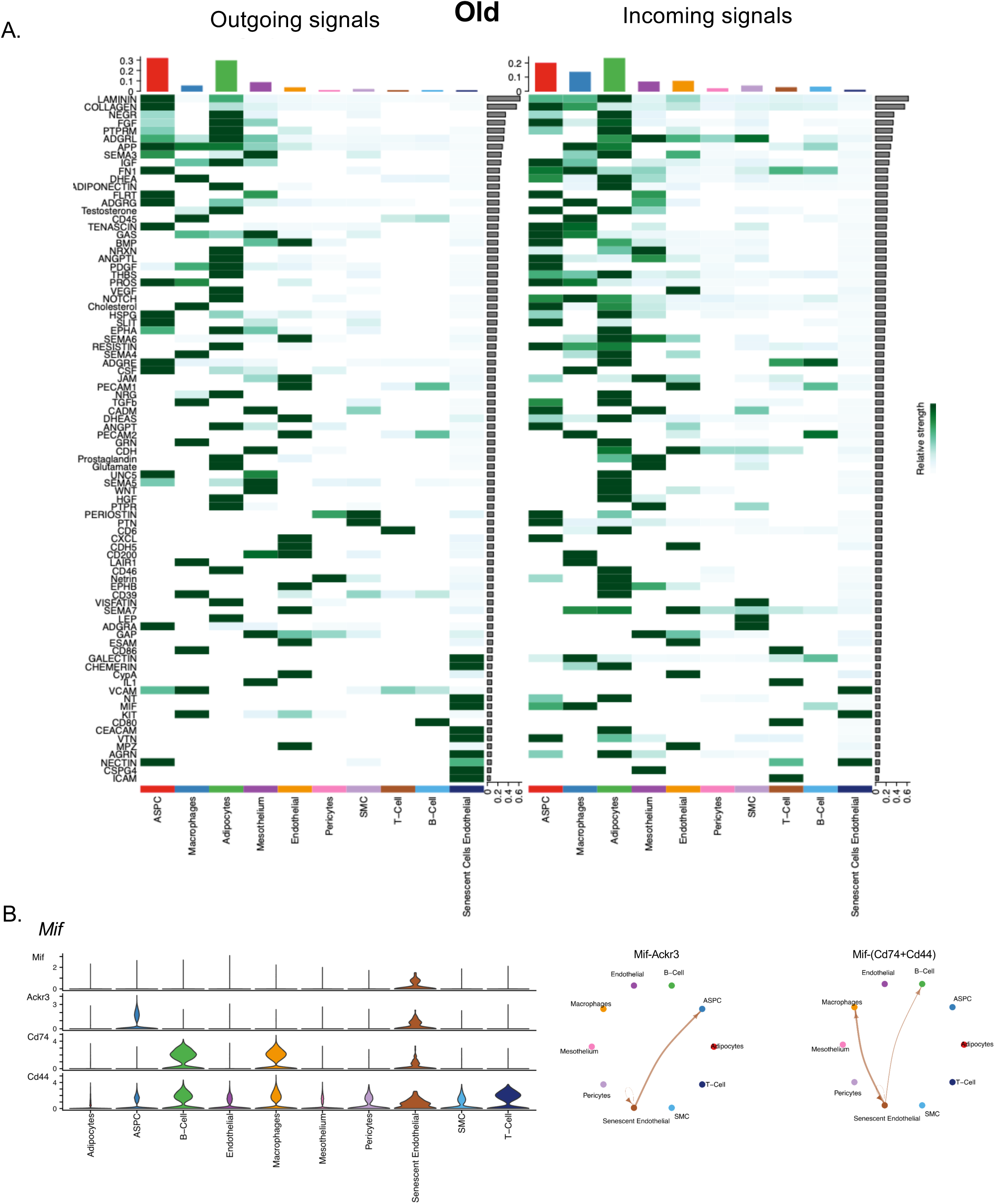
Predicted cell-cell interactions by the endothelial senotype. **A.** Cell Chat predicted outgoing and incoming signals from main nuclei clusters and the old adipose endothelial cell senotype (SEAC strategy). The histogram on top indicates the relative amount of outgoing or incoming signals by cell type. **B.** Violin plots of *Mifa* expression and its receptors, *Ackr3* an *Cd74/Cd44*, by cell type (left). Circle plots indicating predicted cell targets of senescent endothelial cell MIF secretion (right).

### Identification of genes predictive of the senescent endothelial nuclei

We next took a non-parametric bioinformatic approach to rank genes based on their specificity or probability of identifying the senescent endothelial nuclei using genesortR^52, 53^ (**Fig. 6 A,B**). We focused on genes above the 98th quantile ranking comparing senescent to old non-senescent endothelial cells. This identified 14 genes, five of which (*Postn*, *Stac*, *Lin7a*, *Agtr1a*, *Notch3*) are previously linked to senescence^54–60^. Interestingly, eight genes of these 14 encode proteins that are part of or affect ion channels or transporters, including *Kcnq5*, *Plcl1*, *Cacna1c*, *Abcc9*, *Ano1*, *Stac*, *Lin7a*, and *Cdh2*.

**Figure 6.**
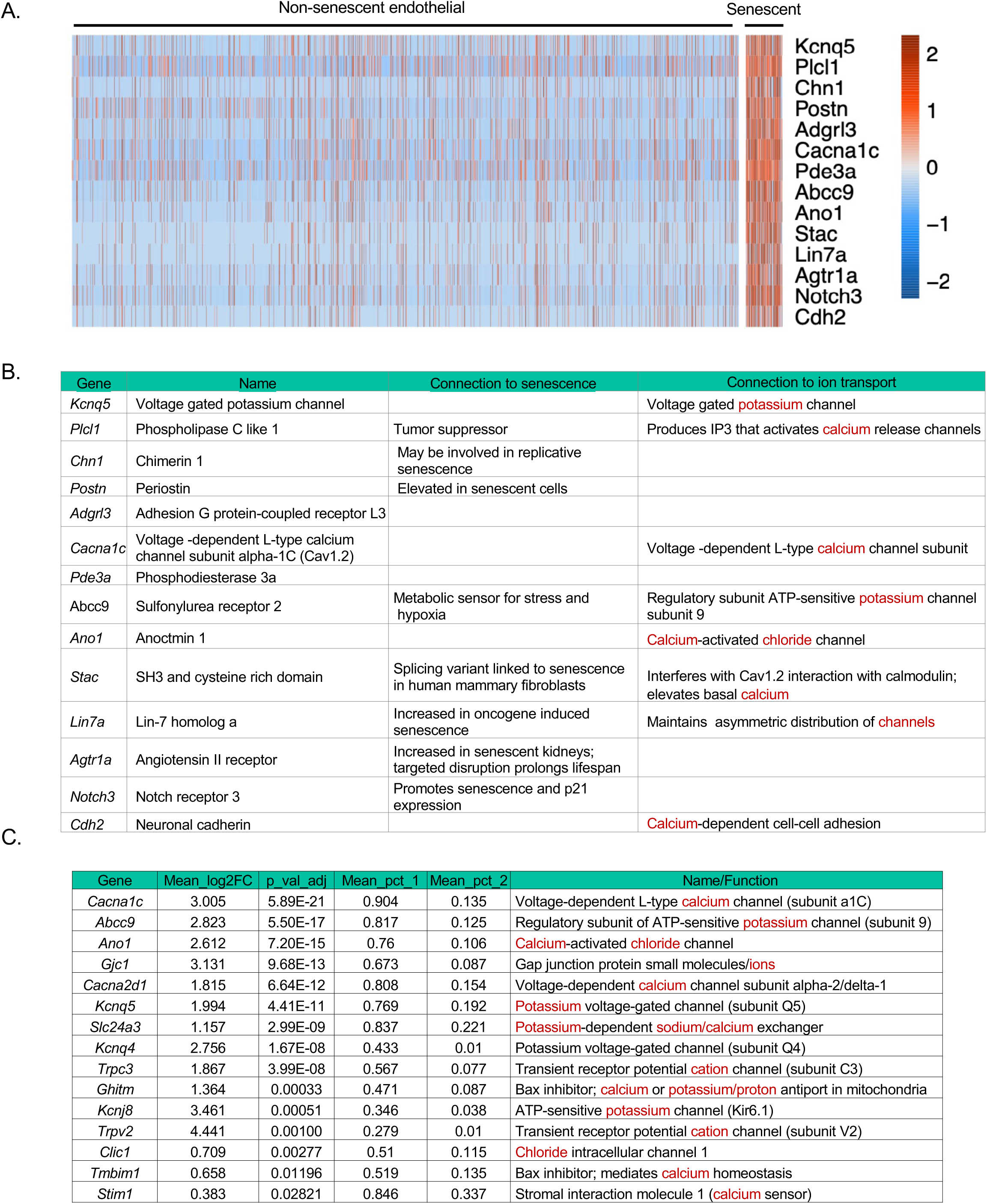
GenesortR prediction of the endothelial senotype and the dysregulation of ion channel expression. **A.** Heat map of the upper 98^th^ quantile of genes most predictive of the senescent endothelial cell nuclei, as compared to non-senescent endothelial nuclei isolated from old murine perigonadal adipose tissue. **B.** Table of the top 98^th^ quantile of predictive genes, their connection to senescence and their connection to ion transport. **C.** Top 98^th^ quantile of DEG in the endothelial senotype compared to old non-senescent endothelial nuclei that are part of the monoatomic ion channel activity gene ontology term GO:0005216.

To determine whether ion channels could be detected on a broader scale, we interrogated the overlap of DEG from senescent endothelial nuclei compared to the old non-senescent nuclei with the GO term monoatomic ion channels, which yielded 15 genes **(Fig. 6C)**. Thirteen of these are ion channel subunits, sensors, or inhibitors of BAX involved in calcium regulation of organelles and blocking of apoptosis. A similar comparison of the senescent endothelial DEG to the young endothelial cells resulted in 27 ion channel genes **(Ext. Table 7)**. The predominance of ion dysregulation in senescence was unexpected but may provide a unique strategy for identification of senescence as well as novel targets for senolytics. Taken as a whole, these results highlight a method to identify and characterize SnCs, which yielded unexpected features of SnCs, including novel senotype and predictions of functional impact.

## Discussion

Senescence is a cell fate that is established to accumulate with chronological age and plays a major role in the risk of chronic disease(s)^21–26^. SnCs have heterogenous features, influenced by means of senescence induction, cell type, and tissue microenvironment^34, 61^. To identify SnCs in murine adipose tissue, we generated snRNAseq data from perigonadal adipose tissue from young and old female mice. We identified a novel age-dependent adipose endothelial SnC population that expresses unique biomarkers, including a dysregulation of ion channel gene expression. Along with the KEGG term cellular senescence, these SnCs significantly upregulated *Cdkn1a* and *Cdkn1b* (representing cell cycle arrest) and *Bcl2* (anti-apoptotic protein). Additionally, numerous ECM and secreted proteins were identified, including a subset of the SenMayo gene panel. Genes upregulated in these senescent endothelial cells included *Jun*, *Tnfrsf1a*, *Igfbp7*, *Serpine2* and *Ets*, a set of genes previously found in *Cdkn1a/p21^Cip1^* expressing cells^62^. Consistent with our recent demonstration that *Cdkn1a* and *Igfbp7* are a marker pair for senescent fibroblasts across tissues^63^, both are increased in these senescent endothelial cells. Our additional analyses focused on TF, cell-cell communications and top gene-correlations to fully characterize and define the senescent endothelial nuclei.

We used TF analyses to identify upstream regulators that drive the dysfunction of endothelial cells upon aging and senescence. Notably, included in the top TF responsible for the transcriptome in the senescent endothelial nuclei were p53, NF-kB, Sp1, Myc, Hif1/2 and AP1/2, all known major TF activities found in senescence. Additionally, loss of TF activity in old-senescent and non-senescent cells included many TF involved in angiogenesis (*Gata6, Rest, Elk1, Znf24, Zmynd8 and Lyll*) although not *Vegfa*, potentially a sign of loss of cellular identity. Similar to Vegfa, other angiogenic factors such as, *Angpt1* and *Pdgfa* were not significantly upregulated in the SnCs, whereas *Angpt2* was increased, a gene known to be upregulated by cellular stress and chronic angiotensin II signaling^64^ and previously identified as a SASP factor. MECOM, a critical zinc-finger TF that plays an important role in endothelial lineage and cellular identity was only found in the young TF. This provides further support to the hypothesis that the senescent endothelial cells have lost their identity. Loss of WWTR1 leads to ER stress, mitochondrial structural and functional abnormalities and impaired Na+/K+ ATPase localization^65^, resulting in organelle dysfunction, consistent with detrimental consequences of senescence. Overall, the identification of these TF provide evidence that we have identified a senescent population as part of the old capillary endothelial cells.

DEG and Cell Chat analyses demonstrated that senescent endothelial nuclei overexpress genes encoding multiple proteins involved in cell adhesion, immune regulation, and inflammation. This is highlighted by the increase expression of *Mif*, which inhibits adipogenesis, a process that decreases with age^4, 51^. MIF increases inflammation in macrophages and B cells and can promote survival of B cells^50, 66^. This is consistent with the fact that although B cell numbers increase with age in adipose tissue, they exhibit diminished function. Furthermore, these results imply that these Mif-expressing SnCs contribute to the reduction of lifespan as Mif knockout mice have been reported to be long-lived^67^. Other ligand-receptor interactions included overexpression of *Icam2* from senescent endothelial cells, known to regulate endothelial barrier function and permeability which may be responsible for leaky vessels that occur upon aging^68^. Other ligands overexpressed from the senescent nuclei include ligands that affect immune cell distribution and/or function, including *Ceacam*, *Chemerin*, *Lgals9*, *Itga4* and *Nectin*. This may contribute to age-dependent SnC accumulation due to impaired immune surveillance and clearance, and therefore reflects a potentially important therapeutic target.

We further characterized the SnC nuclei by ranking genes most predictive of the senescent endothelial senotype. From the top 98^th^ quantile genes, we found five that were previously linked to senescence, including the following. *Postn* encodes a matricellular protein involved in fibrosis that promotes inflammation and is elevated in SnCs^60^. A novel splice variant of *Stac* is implicated in senescence^55^. *Lin7a* is involved in cellular polarity and is increased in oncogene-induced senescence^56^. *Atgr1a* encodes the angiotensin II receptor important for vasoconstriction and ATGR1a causes senescence by promoting NADPH oxidase activity, increasing ROS, mitochondrial dysfunction, and low-grade inflammation (inflammaging)^69^. *Atgr1a* has been found increased in vascular and kidney cell senescence and induces senescence in hepatocellular carcinoma^48, 58, 69^. Disruption of *Atgr1a* delays senescence, reduces oxidative stress and promotes longevity and consistent with this, knockout of *Agtr1a* increases average lifespan in mice by 26%^48^. The AGTR1a ligand, angiotensin II, accelerates the onset of senescence by increasing oxidative stress potentially through telomerase inactivation and blocking proliferative activity^57^. *Notch3* is a tumor suppressor that leads to p21 induction and senescence, dependent on canonical Notch signaling^59^. Other top predictive genes are novel biomarkers of senescence. *Adgrl3* is a member of the latrophilin subfamily of adhesion G protein-coupled receptors that functions in cell-cell adhesion and signal transduction and *Pde3a* is a phosphodiesterase involved in regulating endothelial barrier function.

Surprisingly, eight of the 14 most predictive genes are involved in ion transport. Calcium is an important second messenger in signal transduction pathways and is involved in many cellular processes, including gene expression, cell growth, metabolism, neurotransmitter release, membrane potential and apoptosis^70^, thus requiring exquisite cellular regulation. Aging disrupts calcium homeostasis, with mitochondrial Ca^++^ levels increasing upon ER stress^71^. Mitochondrial calcium levels affect multiple physiological processes, including energy supply, cellular senescence or cell fate^71, 72^, and there are examples of senolytics that alter cellular ion levels^73–75^.

Although many of the ion channel genes predictive of these SnCs were not previously connected to senescence, the role of ions and ion channels has been implicated in senescence^72, 76–78^. This includes *Atgr1a*, a gene correlated with aging that activates phospholipase C and increases intracellular calcium. CAV1.2 (*Cacna1c*) is a pore forming subunit of an L-type voltage-gated calcium channel regulated by calmodulin that allows calcium influx upon strong membrane depolarization^79^. STIM1 is localized in the ER, and when ER Ca^++^ is depleted, STIM1 multimerizes and translocates to the plasma membrane to activate a calcium release-activated Ca^++^ channel which mediates store-operated Ca^++^ entry, causing Ca^++^ influx^80^. *Stim1* expression is controlled by NF-kB and Hif1, two TF identified as upregulated in senescent endothelial cells^80^.

Potassium levels also play important roles in endothelial cells by decreasing stiffness and promoting proper vessel relaxation. Potassium channels were found to be dysregulated in the senescent endothelial cells. KCNQ5 (Kv7.5) is a voltage-gated potassium channel that functions to transport potassium out of the cell and that is involved in regulating blood vessel diameter and vascular tone. Upregulation of *Kcnq5* in senescent endothelial cells is consistent with vascular dysfunction and increased vascular resistance in aged organisms. ABCC9 (sulfonylurea receptor 2) is a regulatory subunit of the ATP-sensitive potassium channel that binds to KCNJ8. ABCC9 is located on plasma and mitochondrial membranes and acts as a metabolic sensor controlling potassium transport. Further indication of altered ion channels in the top predictive genes is increased *Lin7a*, which maintains asymmetric distribution of ion channels. Also, the predictive gene *Cdh2* encodes a protein that in a calcium-dependent manner is involved in cell-cell adhesion.

Endothelial cell senescence was previously demonstrated, both in *in vitro* and in vivo, including in adipose tissue^81, 82,83–86^ and in human microvascular endothelial cells^87^. Our study extends these findings by characterizing a unique endothelial population with newly identified DEG in aged murine visceral adipose tissue. Our data also enables prediction of the functional impact of these age-related senescent endothelial cells on their tissue of origin and potential mechanisms behind that. This reveals a novel endothelial senotype and identifies endothelial cells as important targets for senotherapeutics that could impact age-related chronic inflammation and metabolic diseases.

## Methods

Frozen perigonadal adipose tissue from young and old (6-and 24-months-old), C57Bl/6J mice was used. Nuclei were prepared as protocols.io: The isolation of nuclei from frozen mouse white adipose tissues for single-nucleus genomics--University of Minnesota TMCs; adapted from Van Hauwaert et. al^88^. Nuclei were counted and submitted to the University of Minnesota Genomics Center for 10X Genomics Chromium 5’ nuclei capture and sequencing.

### Processing of FASTQ files

FASTQ files were processed by University of Minnesota Genomic center (UMGC) with cell ranger version 7.1.0, chemistry set to Single Cell 5’ R2-only and mapped to the mouse reference genome version mm10. Quality control included removal of ambient RNA and low-quality nuclei.

### Processing of SnRNA-seq 10x Matrix

10x Genomics Cellranger (V7.0) was used to align demultiplexed fastq files, and the filtered counts matrix retrieved for each sample was processed by Seurat (V5.0)^89^. Matrix files were read into R (V4.4.0) with Seurat’s Read10x function using default settings on raw counts, filtering out cells with fewer than 100 feature_RNA counts. The Vlnplot function was used to visualize feature_RNA counts distribution, and thresholds for feature_RNA counts were set based on the first quantile and upper quantile regions. Cells with above 200 feature_RNA count and less than 7500 feature_RNA count, and less than 5% mitochondrial content were considered for downstream analyses. The sample counts were LogNormalized by Seurat’s NormalizeData function, and 2000 variable features were found for each sample by the FindVariableFeatures function. 2000 integration features were selected by SelectIntegrationFeatures, and integration anchors were calculated using FindIntegrationAnchors with one of the old female mouse samples as a reference. The samples were batch corrected and integrated by the IntegrateData function. The corrected counts were scaled, and 30 principal components (PCs) were calculated by the RunPCA function as per the standard Seurat pipeline. UMAP coordinates were calculated, as well as neighbors, by the RunUMAP and FindNeighbors functions. The clusters were calculated at a resolution of 0.5 by the FindClusters function. The main cell types were annotated based on Emont et. al^90^.

### Subclusters Analysis

Subclusters of endothelial nuclei were created by extracting the cell type, setting it as an integrated assay, following the standard Seurat steps, which include scaling the data, recalculating 30 principal components, finding neighbors, clusters, and calculating the UMAP coordinates. The resolution at which the endothelial clusters were created was 0.5. and the subclusters were annotated based on previously published markers^90^.

### DEG Analyses

Seurat’s Findmarkers function with default settings was used to calculate the DEG between the young and old age groups. The DEG were calculated for each cell type as well as between old versus young cell types. The Wilcoxon rank-sum test, Seurat’s default method, was used to compute differential expression.

### Identification of Senescent nuclei

The SenMayo gene list^41^ was utilized by calculating gene counts through a binary transformation of the data. The rule of transformation was set based on expression – if the gene has an expression value above 0, it was set to 1; otherwise, it was set to 0. Given the binary matrix, the gene counts were summed up for each nucleus and plotted as a distribution and boxplot. The upper quartile boxplot region was used, resulting in selection of counts ≥ 25. In addition, nuclei that were assigned to the G1 phase by Seurat were included. A second strategy (SEAC) used Pearson correlation. Potential senescent genes (*Cdkn1a, Bcl2l1, Pcna, Mki67, Plaur, and Csf1*) were used to extract the nuclei using the following function subset (data, subset = *Cdkn1a* > 0.7, *Bcl2l1* > 0.5, *Pcna* = 0, *Mki67* = 0, *Plaur* > 0, *Csf1* > 0). Nuclei expressing these genes simultaneously were selected – with nuclei having Cdkn1a expression value above 0.7 based on a 3rd quartile value, Bcl2l1 above 0.5 also based on 3rd quantile value. *Mki67* and *Pcna* were required to be 0 as an indication of proliferative arrest and *Plaur,* and *Csf1* were above 0. Finally, cells were required to be in G1 phase. A random nuclei was selected to be a reference. Against this, a Pearson correlation score (threshold of 0.5) was calculated for all nuclei. In addition, only G1 cells were included. To calculate DEG for senotypes, a custom function was created, which took advantage of the bootstrapping technique. Non-senotypes of an equal number to senotypes were compared, and this was done randomly 100 times. The average of log fold change and Bonferroni corrected p-values were calculated (mean_logfc > 0 & mean_adjusted_pvalue < 0.05).

### Cell – Cell Communication Analysis

Cellchat version 2.2.0 was used on the old Seurat objects^49^. Standard steps laid out by the cellchat vignette were followed. The overexpressed genes were identified by celltypes identifyOverExpressedGenes(), upregulated interactions were calculated using identifyOverExpressedInteractions(), communication probability of pathways and ligand receptor pairs in use were calculated using computeCommunProb(), where the population size parameter was set to “True” to avoid bias due to differences in population. Finally, pathways that were expressed in a minimum of ten cells were included. The pathways that had a communication probability above 0.01 were included. We set a low threshold to capture as many pathways as possible. Up-regulated pathways determined by CellChat were extracted, and visualization functions within the CellChat package were used to create cell-cell communication plots. The cell-cell communication was assessed between SnC endothelial and all major cell types.

### Matrisome Analysis

The changes in the ECM were identified using Matrisome AnalyzeR, a tool that annotates and quantifies ECM molecules in big datasets and is available in the following website: https://sites.google.com/uic.edu/matrisome/home^45^.

### KEGG Pathway Analysis

Kegg pathways were retrieved based on significantly (P < 0.05) up-regulated and down-regulated genes in older populations on all the cell types (ASPC, Adipocytes, Endothelial, Mesothelium, Macrophages, T-Cell, B-Cell, Pericytes). Clusterprofiler package version 4.8.3 (Yu et al. 2012) was used to find the KEGG pathways^91^.

### Transcription factors

Transcription involved in the old and young cell types and senotype endothelial cells were calculated using a package called decoupleR^92^. DecoupleR leverages Omnipath, a resource comprising over 100 databases, and calculates a t-value, which was annotated as a mean activity score. The TF above the 90% quantile were considered. Quantile (prob = 0.90) function in R was used to calculate the threshold.

### Ion Channel Gene Analysis

GenesortR version 0.4.3^53^ was acquired from https://github.com/mahmoudibrahim/genesorteR and used to rank genes, and markers above 98th quantile were acquired for old potential endothelial senescent cells and young and old non-senescent cells. The markers specific to potential endothelial senescent cells were annotated, and the frequency of ion channel genes was noted. The markers were also noticed to be significant from the bootstrapping gene differential method. Given the frequency of ion channel genes, we acquired a set of ion channel genes from the GO term monoatomic ion channel activity. The genes present in the GO term list were crossed with DEG to understand how many significant genes were up-regulated and down-regulated.

### Inflammation Score

Gene sets in GO terms acute inflammatory response were acquired and UCell^38^ was used to score the cells for inflammation and stem cell characteristics. Scores were plotted using ggplot2 R package geom_density function.

## Acknowledgements

We would like to acknowledge the Niedernhofer mouse core for the adipose samples, the University of Minnesota Genomics Center for performing the single nuclei capture and sequencing, and the Minnesota Supercomputing Institute. We also would like to acknowledge the Minnesota SenNet team and the larger SenNet community for helpful discussions.

## Author contributions

AVH, DAB, JW, CDC, AB, PDR, LJN Conceptualization AVH Performed experiments

AVH, SA, JW Analyses

AVH, SA, DAB, JW, CDC, AB, PDR, LJN Discussion, Review and Editing

