## Extended figures for "Identification of an Endothelial Senotype Predicted by Ion Channel Expression in Aged Murine Perigonadal Adipose Tissue"

Extended Figure 1. Single nuclei sequencing of female perigonadal adipose tissue from old and young mice.

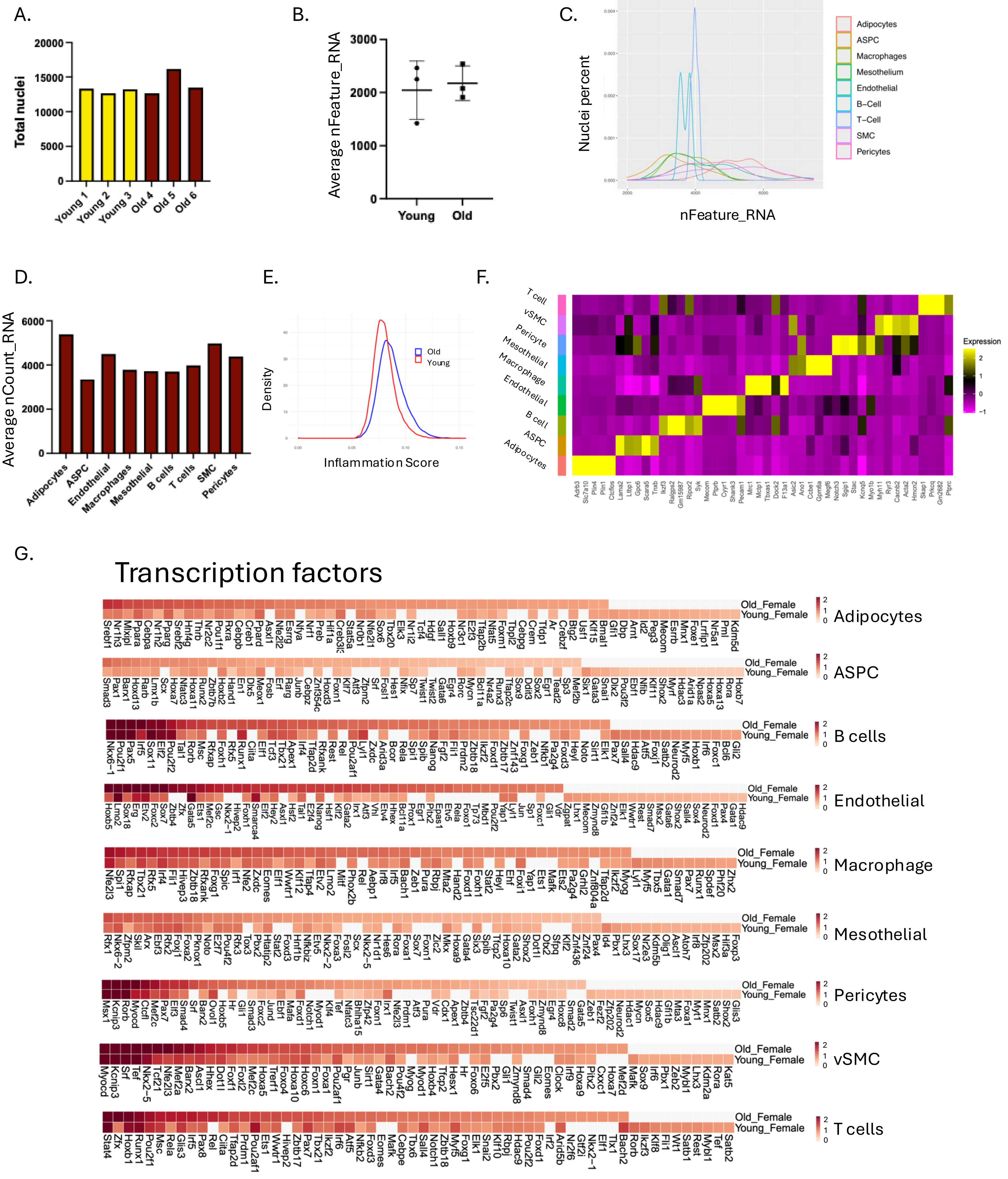

Extended Figure 1 continued. Changes in gene expression in young versus old perigonadal adipose tissue.

H. Kegg Pathways

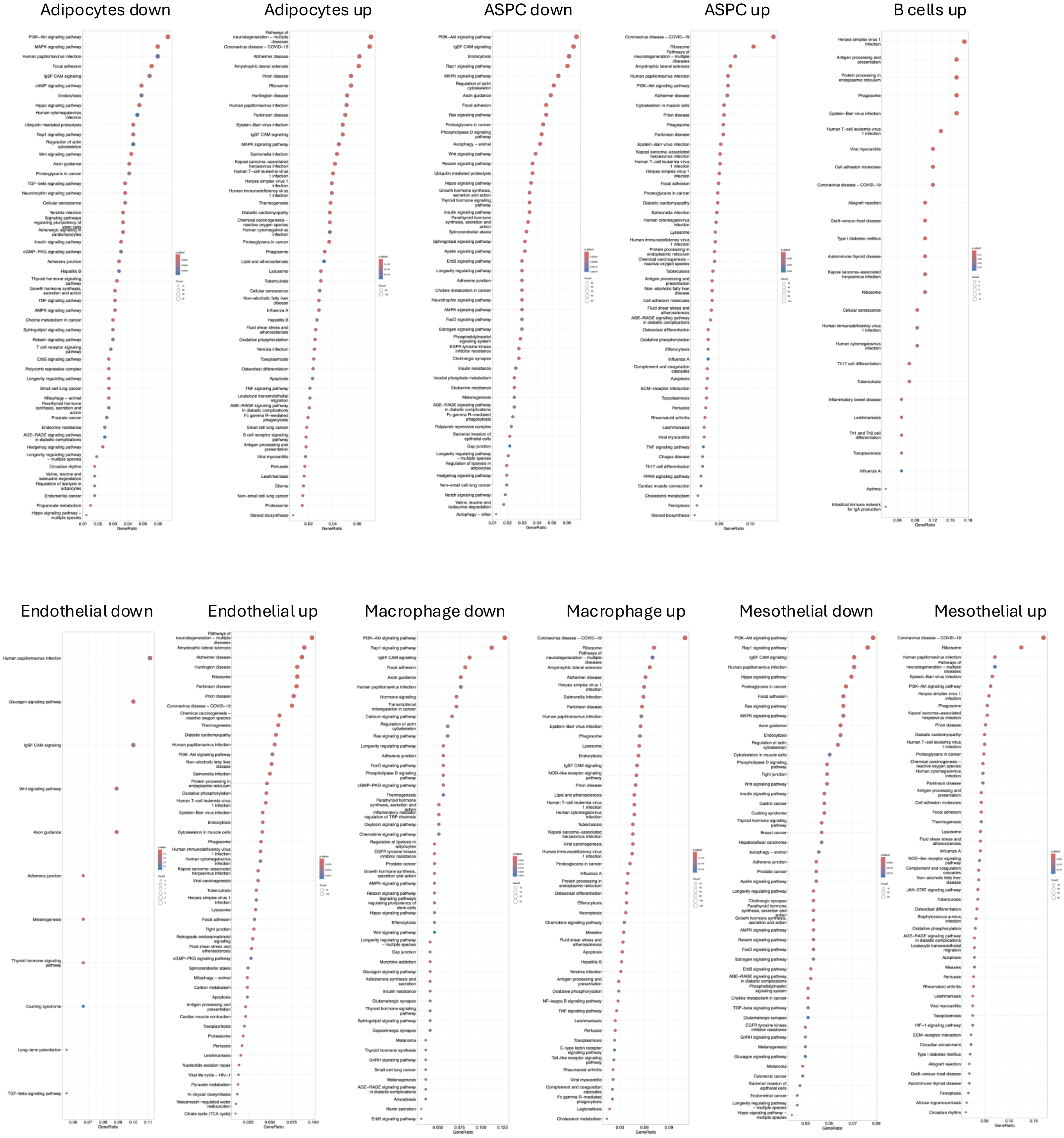

Extended Figure 1 continued. KEGG pathways old to young by cell type

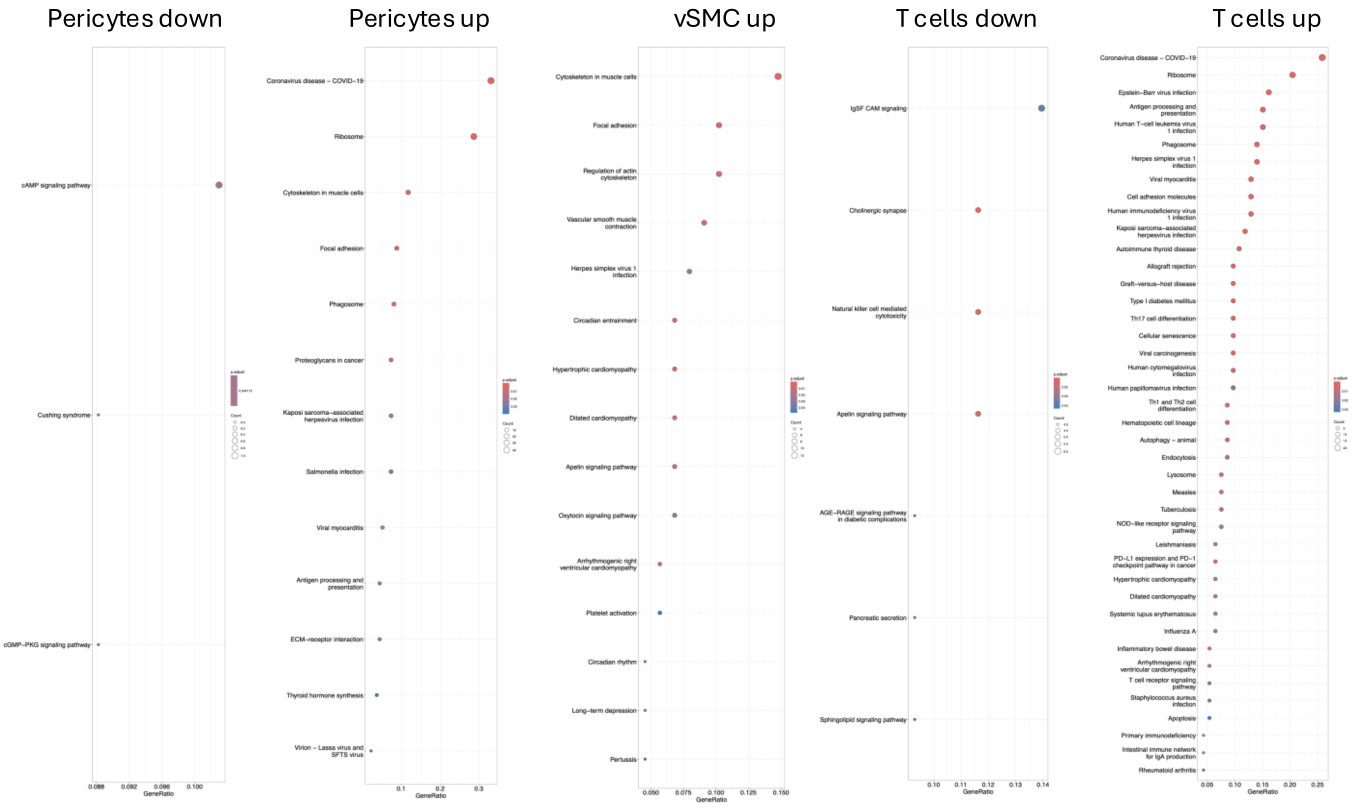


**Extended Figure 1. Single nuclei sequencing of female perigonadal adipose tissue from old and young mice. A.** Total nuclei analyzed in each sample. **B.** Average nFeature_RNA in young and old. **C.** Density of nuclei by nFeatures_RNA. **D.** Average nCount_RNA by cell type. **E.** Inflammation score calculated using the genes from the GO term, “acute inflammatory response”. **F.** Heat map of top gene expression from each cell type. **G.** Top 50 TF whose activity predicts the transcriptomic profile of each cell type in perigonadal murine nuclei from young and old mice. **H.** KEGG pathways of significant DEG, separated by directionality of expression change.

Extended Figure 2. Cell chat SEAC DEG

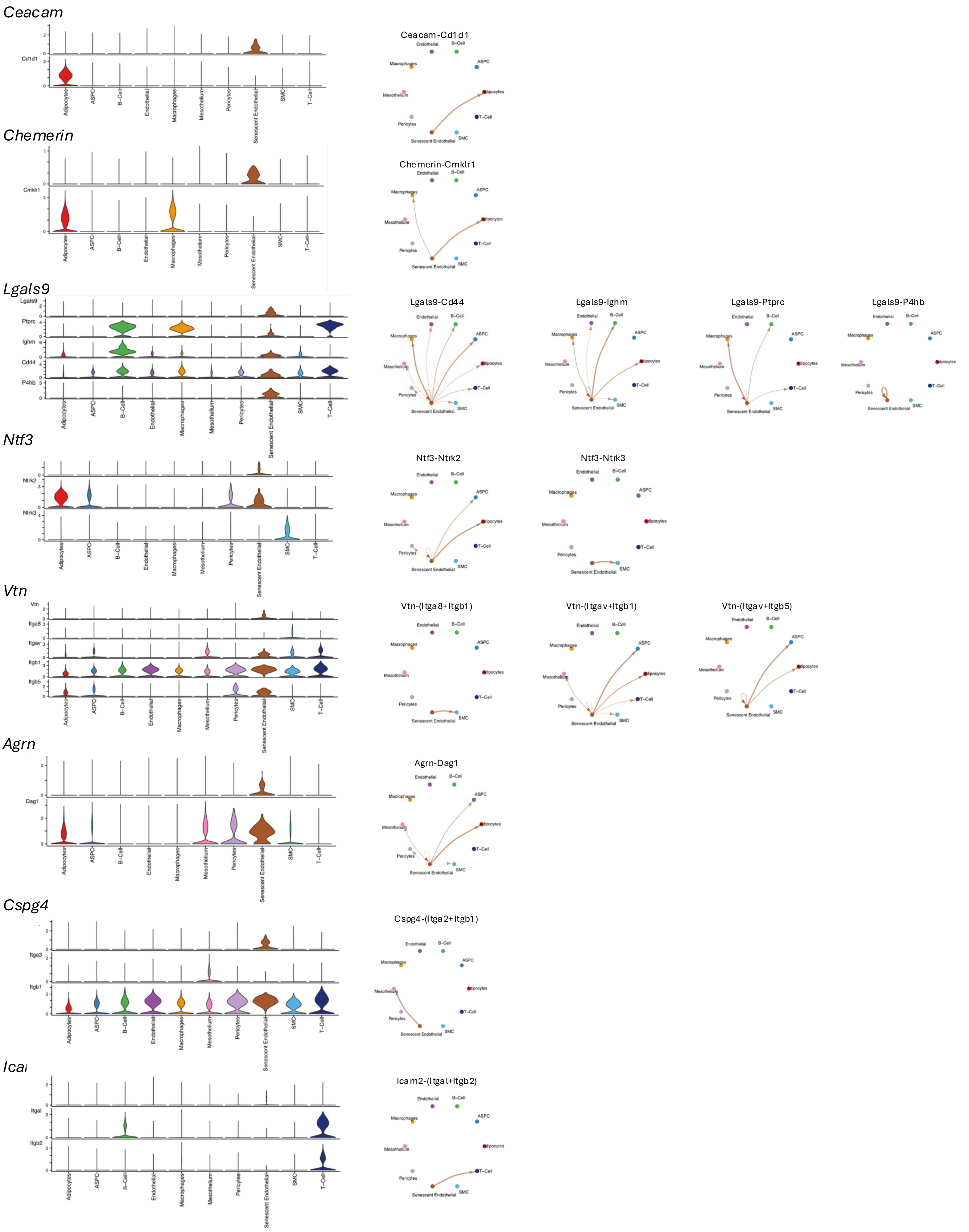

Extended Figure 2. Cell chat SEAC DEG

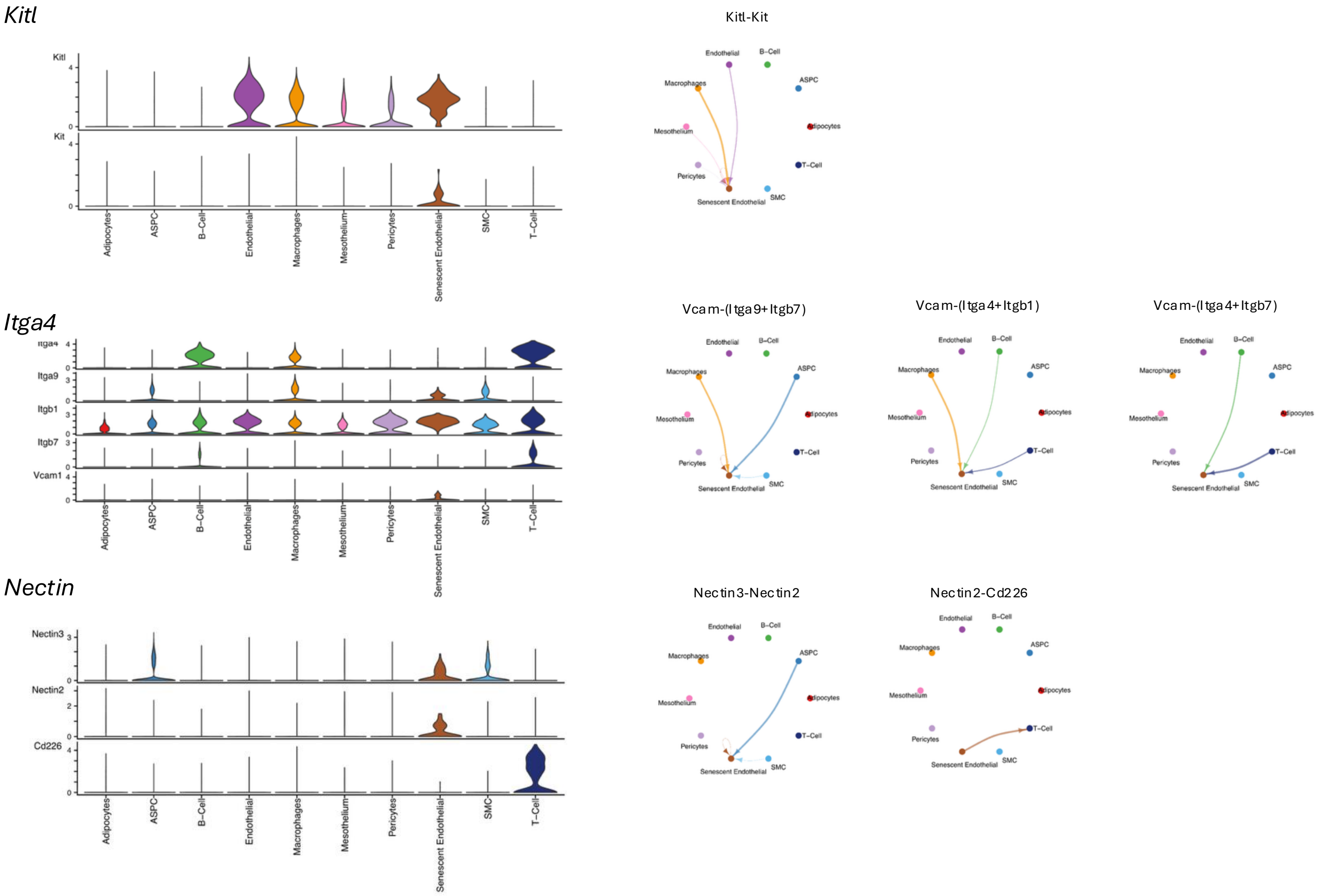


**Extended Figure 2.**

Violin plots of *Ceacam, Chemerin, Lgals9, Ntf3, Vtn, Agrn, Cspg4, Icam2, Kitl, Vcam,* and *Nectin3* expression in the endothelial senotype and their receptors for each of the main cell types in murine adipose tissue (left). Circle plots indicating predicted cell targets of these senescent endothelial upregulated genes (right).

**Extended Table 1**

DEG from endothelial SnC (SenMayo strategy) versus young

| | gene | mean_pval | mean_log2FC | p_val_adj | mean_pct_1 | mean_pct_2 |
|---|---|---|---|---|---|---|
| **1** | Rps24 | 1.10E-16 | 4.428431867 | 2.74E-12 | 0.959 | 0.082 |
| **2** | Rps16 | 3.29E-16 | 5.623591473 | 8.20E-12 | 0.898 | 0.02 |
| **3** | Rpl38 | 4.11E-16 | 4.005168583 | 1.02E-11 | 0.918 | 0.02 |
| **4** | S100a6 | 1.34E-15 | 4.14898877 | 3.34E-11 | 0.939 | 0.061 |
| **5** | Rps21 | 1.92E-15 | 4.856355341 | 4.78E-11 | 0.898 | 0.041 |
| **6** | Cst3 | 2.17E-15 | 3.935157478 | 5.39E-11 | 0.959 | 0.102 |
| **7** | Tmsb4x | 2.44E-15 | 4.093403183 | 6.06E-11 | 0.98 | 0.163 |
| **8** | Ftl1 | 5.16E-15 | 5.292790821 | 1.28E-10 | 0.857 | 0.02 |
| **9** | Rps13 | 5.43E-15 | 6.545160454 | 1.35E-10 | 0.816 | 0 |
| **10** | Rgs5 | 8.00E-15 | 6.127501059 | 1.99E-10 | 0.837 | 0.02 |
| **11** | Ptma | 9.95E-15 | 3.790007381 | 2.47E-10 | 0.918 | 0.082 |
| **12** | Hsp90b1 | 1.26E-14 | 2.901957938 | 3.14E-10 | 0.98 | 0.163 |
| **13** | Gucy1a2 | 1.67E-14 | 7.119042073 | 4.15E-10 | 0.796 | 0 |
| **14** | Npc2 | 1.67E-14 | 5.797202799 | 4.15E-10 | 0.796 | 0 |
| **15** | Col6a2 | 1.88E-14 | 2.816242214 | 4.67E-10 | 0.878 | 0.041 |
| **16** | Rps20 | 2.41E-14 | 4.442640657 | 6.00E-10 | 0.857 | 0.041 |
| **17** | Rps28 | 2.54E-14 | 3.813608688 | 6.32E-10 | 0.898 | 0.061 |
| **18** | Inpp4b | 3.71E-14 | 3.95174399 | 9.24E-10 | 0.837 | 0.02 |
| **19** | Rpl27a | 4.19E-14 | 5.107725644 | 1.04E-09 | 0.816 | 0.02 |
| **20** | Rpl18 | 5.01E-14 | 6.459754223 | 1.25E-09 | 0.776 | 0 |
| **21** | Actb | 7.44E-14 | 3.763731253 | 1.85E-09 | 0.898 | 0.143 |
| **22** | Rpl37a | 8.82E-14 | 3.315702998 | 2.20E-09 | 0.918 | 0.102 |
| **23** | Rps29 | 9.81E-14 | 3.989182545 | 2.44E-09 | 0.878 | 0.082 |
| **24** | Rpl41 | 9.90E-14 | 3.513251029 | 2.46E-09 | 0.918 | 0.122 |
| **25** | Rps2 | 1.47E-13 | 4.760031551 | 3.66E-09 | 0.796 | 0.02 |
| **26** | Notch3 | 1.47E-13 | 7.599283042 | 3.66E-09 | 0.755 | 0 |
| **27** | Fth1 | 1.52E-13 | 3.32107616 | 3.79E-09 | 0.857 | 0.041 |
| **28** | Col6a3 | 1.56E-13 | 2.989574388 | 3.88E-09 | 0.939 | 0.102 |
| **29** | Rpl13a | 1.58E-13 | 3.257674334 | 3.93E-09 | 0.878 | 0.061 |
| **30** | Pla2r1 | 1.72E-13 | 3.601459936 | 4.28E-09 | 0.837 | 0.041 |
| **31** | Rpl9 | 1.81E-13 | 4.433905704 | 4.51E-09 | 0.816 | 0.041 |
| **32** | Rps8 | 2.19E-13 | 3.292060153 | 5.44E-09 | 0.918 | 0.122 |
| **33** | Rpl37 | 2.19E-13 | 5.530434502 | 5.45E-09 | 0.776 | 0.02 |
| **34** | Rps23 | 2.20E-13 | 3.587317408 | 5.47E-09 | 0.878 | 0.082 |
| **35** | Rps3a1 | 2.58E-13 | 4.111979156 | 6.42E-09 | 0.816 | 0.041 |
| **36** | Arpc3 | 3.20E-13 | 3.777457393 | 7.95E-09 | 0.796 | 0.02 |
| **37** | Postn | 3.54E-13 | 5.557672112 | 8.80E-09 | 0.776 | 0.02 |
| **38** | Rpl28 | 3.54E-13 | 5.037123742 | 8.80E-09 | 0.776 | 0.02 |

| | | | | | | |
|---|---|---|---|---|---|---|
| **39** | B230219D22R | 3.60E-13 | 2.891766581 | 8.95E-09 | 0.796 | 0.02 |
| **40** | Rpl35a | 3.74E-13 | 3.705583262 | 9.32E-09 | 0.837 | 0.061 |
| **41** | Cox8a | 4.24E-13 | 6.155833268 | 1.06E-08 | 0.735 | 0 |
| **42** | Rpl23a | 4.24E-13 | 5.979254155 | 1.06E-08 | 0.735 | 0 |
| **43** | Rpl39 | 4.24E-13 | 6.334555099 | 1.06E-08 | 0.735 | 0 |
| **44** | Rpl7 | 4.24E-13 | 5.876970391 | 1.06E-08 | 0.735 | 0 |
| **45** | Rps18 | 4.24E-13 | 5.946700818 | 1.06E-08 | 0.735 | 0 |
| **46** | Cox7c | 4.48E-13 | 4.804789626 | 1.12E-08 | 0.776 | 0.02 |
| **47** | Cd63 | 5.05E-13 | 4.652623617 | 1.26E-08 | 0.776 | 0.02 |
| **48** | Rpl21 | 5.30E-13 | 4.14225853 | 1.32E-08 | 0.837 | 0.082 |
| **49** | Tbc1d1 | 6.64E-13 | 3.177753436 | 1.65E-08 | 0.837 | 0.061 |
| **50** | B2m | 6.94E-13 | 3.004212858 | 1.73E-08 | 0.939 | 0.184 |
| **51** | Agtr1a | 7.42E-13 | 5.236291317 | 1.85E-08 | 0.755 | 0.02 |
| **52** | Rpl30 | 7.78E-13 | 3.64488151 | 1.94E-08 | 0.816 | 0.041 |
| **53** | Eef1a1 | 9.05E-13 | 2.765590346 | 2.25E-08 | 0.959 | 0.204 |
| **54** | Steap4 | 9.69E-13 | 4.148504697 | 2.41E-08 | 0.796 | 0.041 |
| **55** | Tpt1 | 1.15E-12 | 3.194405177 | 2.87E-08 | 0.878 | 0.102 |
| **56** | Rpsa | 1.22E-12 | 3.889372264 | 3.03E-08 | 0.796 | 0.041 |
| **57** | Rps15 | 1.23E-12 | 3.219120386 | 3.06E-08 | 0.816 | 0.041 |
| **58** | Anxa5 | 1.24E-12 | 3.05713295 | 3.08E-08 | 0.857 | 0.061 |
| **59** | Cacna1c | 1.30E-12 | 3.520321285 | 3.23E-08 | 0.878 | 0.122 |
| **60** | Abcc9 | 1.34E-12 | 4.923069012 | 3.32E-08 | 0.755 | 0.02 |
| **61** | Sparcl1 | 1.34E-12 | 2.723761646 | 3.32E-08 | 0.959 | 0.245 |
| **62** | Gpihbp1 | 1.47E-12 | 3.580146914 | 3.65E-08 | 0.837 | 0.061 |
| **63** | Gsn | 1.48E-12 | 2.904166895 | 3.68E-08 | 1 | 0.408 |
| **64** | Calm2 | 1.72E-12 | 3.258272741 | 4.28E-08 | 0.796 | 0.041 |
| **65** | Fau | 1.83E-12 | 3.589874599 | 4.55E-08 | 0.837 | 0.082 |
| **66** | Cfl1 | 2.01E-12 | 4.153385876 | 5.00E-08 | 0.755 | 0.02 |
| **67** | Kcnq5 | 2.05E-12 | 5.935166484 | 5.11E-08 | 0.735 | 0.02 |
| **68** | Eif1 | 2.11E-12 | 4.127926615 | 5.25E-08 | 0.796 | 0.061 |
| **69** | Eef2 | 2.28E-12 | 3.239773679 | 5.66E-08 | 0.816 | 0.061 |
| **70** | Rplp1 | 2.89E-12 | 2.517041927 | 7.20E-08 | 0.918 | 0.122 |
| **71** | Ghr | 3.16E-12 | -1.55323134 | 7.87E-08 | 1 | 1 |
| **72** | Cox6a1 | 3.32E-12 | 6.029811688 | 8.25E-08 | 0.694 | 0 |
| **73** | Mfge8 | 3.32E-12 | 6.692054213 | 8.25E-08 | 0.694 | 0 |
| **74** | Tmed9 | 3.32E-12 | 5.388091361 | 8.25E-08 | 0.694 | 0 |
| **75** | Calm1 | 3.35E-12 | 2.717776014 | 8.35E-08 | 0.898 | 0.122 |
| **76** | Ubc | 3.50E-12 | 3.76643473 | 8.70E-08 | 0.796 | 0.061 |
| **77** | Pdgfrb | 3.63E-12 | 4.359588658 | 9.03E-08 | 0.755 | 0.041 |
| **78** | Rps12 | 3.78E-12 | 3.001940557 | 9.40E-08 | 0.857 | 0.102 |
| **79** | Cldn5 | 3.79E-12 | 2.48762507 | 9.44E-08 | 0.939 | 0.122 |

| | | | | | | |
|---|---|---|---|---|---|---|
| **80** | Pltp | 3.79E-12 | 3.06094152 | 9.44E-08 | 0.796 | 0.041 |
| **81** | Ctsl | 4.13E-12 | 4.561752091 | 1.03E-07 | 0.735 | 0.02 |
| **82** | Rpl17 | 4.75E-12 | 2.753687766 | 1.18E-07 | 0.796 | 0.041 |
| **83** | Actg1 | 5.11E-12 | 3.457449802 | 1.27E-07 | 0.816 | 0.082 |
| **84** | Rpl13 | 5.21E-12 | 3.384619778 | 1.30E-07 | 0.837 | 0.082 |
| **85** | Cd93 | 6.79E-12 | 2.213642663 | 1.69E-07 | 0.939 | 0.122 |
| **86** | Rpl35 | 7.19E-12 | 3.478934727 | 1.79E-07 | 0.796 | 0.061 |
| **87** | Rplp2 | 8.49E-12 | 2.95889726 | 2.11E-07 | 0.837 | 0.082 |
| **88** | Srpr | 9.00E-12 | 5.302111788 | 2.24E-07 | 0.673 | 0 |
| **89** | Stac | 9.00E-12 | 6.939941065 | 2.24E-07 | 0.673 | 0 |
| **90** | Ly6e | 9.05E-12 | 2.515243383 | 2.25E-07 | 0.918 | 0.143 |
| **91** | Angptl2 | 9.16E-12 | 2.923193624 | 2.28E-07 | 0.776 | 0.041 |
| **92** | Dbi | 9.57E-12 | 2.410196327 | 2.38E-07 | 0.837 | 0.061 |
| **93** | Rpl34 | 1.22E-11 | 3.663700067 | 3.03E-07 | 0.776 | 0.061 |
| **94** | C1s1 | 1.33E-11 | 2.549221593 | 3.30E-07 | 0.816 | 0.061 |
| **95** | Rps27a | 1.39E-11 | 3.183410203 | 3.45E-07 | 0.796 | 0.061 |
| **96** | Angptl4 | 1.44E-11 | 2.036780426 | 3.59E-07 | 0.878 | 0.082 |
| **97** | Hmgb1 | 1.45E-11 | 2.598438028 | 3.62E-07 | 0.878 | 0.102 |
| **98** | Rpl27 | 1.65E-11 | 4.016669309 | 4.12E-07 | 0.714 | 0.02 |
| **99** | Id3 | 1.67E-11 | 5.04057075 | 4.15E-07 | 0.694 | 0.02 |
| **100** | Hspa9 | 1.85E-11 | 3.414385941 | 4.61E-07 | 0.714 | 0.02 |
| **101** | Laptm4a | 1.96E-11 | 2.119118775 | 4.88E-07 | 0.98 | 0.184 |
| **102** | Rpl19 | 2.30E-11 | 3.085312275 | 5.72E-07 | 0.755 | 0.041 |
| **103** | Ly6c1 | 2.32E-11 | 2.706800361 | 5.77E-07 | 0.816 | 0.082 |
| **104** | Plin2 | 2.51E-11 | 2.964578491 | 6.25E-07 | 0.796 | 0.061 |
| **105** | Dusp1 | 2.63E-11 | 4.190028484 | 6.53E-07 | 0.694 | 0.02 |
| **106** | Ifitm2 | 2.63E-11 | 4.540397614 | 6.53E-07 | 0.694 | 0.02 |
| **107** | Bgn | 2.64E-11 | 3.732997384 | 6.56E-07 | 0.735 | 0.041 |
| **108** | Cacna2d1 | 2.71E-11 | 2.218784045 | 6.74E-07 | 0.857 | 0.102 |
| **109** | Hnrnpk | 2.89E-11 | 2.38036419 | 7.18E-07 | 0.857 | 0.082 |
| **110** | Marcks | 2.94E-11 | 2.320100083 | 7.33E-07 | 0.939 | 0.163 |
| **111** | Kctd12 | 2.95E-11 | 2.722096551 | 7.34E-07 | 0.796 | 0.061 |
| **112** | Ubb | 3.07E-11 | 3.507438243 | 7.63E-07 | 0.776 | 0.061 |
| **113** | Ywhah | 3.66E-11 | 3.317816577 | 9.12E-07 | 0.735 | 0.041 |
| **114** | Gng11 | 3.68E-11 | 5.764810312 | 9.15E-07 | 0.673 | 0.02 |
| **115** | Rpl6 | 3.68E-11 | 4.429332523 | 9.16E-07 | 0.694 | 0.02 |
| **116** | Rps3 | 3.89E-11 | 3.990001487 | 9.68E-07 | 0.694 | 0.02 |
| **117** | Serf2 | 3.96E-11 | 2.230359895 | 9.86E-07 | 0.837 | 0.082 |
| **118** | Rps11 | 4.12E-11 | 4.064020258 | 1.02E-06 | 0.694 | 0.02 |
| **119** | Adgrl3 | 4.17E-11 | 3.449284615 | 1.04E-06 | 0.776 | 0.082 |
| **120** | H3f3b | 4.55E-11 | 2.516096492 | 1.13E-06 | 0.857 | 0.102 |

| | | | | | | |
|---|---|---|---|---|---|---|
| **121** | Gucy1a1 | 4.71E-11 | 3.011905368 | 1.17E-06 | 0.776 | 0.061 |
| **122** | Ifitm3 | 4.84E-11 | 3.776664 | 1.21E-06 | 0.755 | 0.061 |
| **123** | Crip1 | 5.11E-11 | 3.487714952 | 1.27E-06 | 0.755 | 0.061 |
| **124** | Ly6a | 5.14E-11 | 2.973133817 | 1.28E-06 | 0.816 | 0.102 |
| **125** | Rpl18a | 5.16E-11 | 4.963859461 | 1.28E-06 | 0.673 | 0.02 |
| **126** | Slc38a2 | 5.44E-11 | 2.550405305 | 1.35E-06 | 0.694 | 0.02 |
| **127** | Fabp4 | 5.54E-11 | 3.022736209 | 1.38E-06 | 1 | 0.694 |
| **128** | Myadm | 5.73E-11 | 1.921896596 | 1.43E-06 | 0.918 | 0.143 |
| **129** | Rpl36 | 5.77E-11 | 3.913469107 | 1.44E-06 | 0.714 | 0.041 |
| **130** | Hnrnpa3 | 5.91E-11 | 1.970029985 | 1.47E-06 | 0.898 | 0.143 |
| **131** | Rpl10 | 6.26E-11 | 6.061781928 | 1.56E-06 | 0.633 | 0 |
| **132** | Selenok | 6.26E-11 | 5.392778419 | 1.56E-06 | 0.633 | 0 |
| **133** | Ucp2 | 6.26E-11 | 5.499140877 | 1.56E-06 | 0.633 | 0 |
| **134** | Rpl5 | 6.54E-11 | 2.260838415 | 1.63E-06 | 0.796 | 0.061 |
| **135** | Gas5 | 7.21E-11 | 4.266703837 | 1.79E-06 | 0.673 | 0.02 |
| **136** | Hs2st1 | 7.36E-11 | 2.065004674 | 1.83E-06 | 0.816 | 0.082 |
| **137** | Cfd | 7.54E-11 | 2.065137064 | 1.88E-06 | 0.959 | 0.204 |
| **138** | Sgip1 | 7.54E-11 | 2.487011858 | 1.88E-06 | 0.898 | 0.163 |
| **139** | Rps9 | 8.00E-11 | 2.504739902 | 1.99E-06 | 0.857 | 0.102 |
| **140** | Col1a1 | 8.16E-11 | 2.280055579 | 2.03E-06 | 0.816 | 0.082 |
| **141** | Tmsb10 | 9.01E-11 | 4.33820491 | 2.24E-06 | 0.673 | 0.02 |
| **142** | H2-K1 | 9.52E-11 | 1.818647538 | 2.37E-06 | 1 | 0.306 |
| **143** | Rps7 | 1.02E-10 | 2.911729339 | 2.54E-06 | 0.776 | 0.082 |
| **144** | Ppp2r3a | 1.03E-10 | 1.880837424 | 2.55E-06 | 0.898 | 0.143 |
| **145** | Rpl4 | 1.06E-10 | 3.980525874 | 2.64E-06 | 0.694 | 0.041 |
| **146** | Rps5 | 1.08E-10 | 3.138017987 | 2.68E-06 | 0.755 | 0.061 |
| **147** | Atp6v0b | 1.19E-10 | 3.700545318 | 2.96E-06 | 0.673 | 0.02 |
| **148** | Col5a3 | 1.23E-10 | 2.394566916 | 3.06E-06 | 0.857 | 0.122 |
| **149** | Pde3a | 1.26E-10 | 1.045602758 | 3.13E-06 | 0.98 | 0.245 |
| **150** | Rflnb | 1.30E-10 | 1.755142845 | 3.23E-06 | 0.816 | 0.082 |
| **151** | Rps26 | 1.30E-10 | 2.993340874 | 3.24E-06 | 0.714 | 0.041 |
| **152** | Slc9a3r2 | 1.33E-10 | 3.333089519 | 3.30E-06 | 0.673 | 0.02 |
| **153** | Rnpc3 | 1.40E-10 | 1.881948161 | 3.49E-06 | 0.673 | 0.02 |
| **154** | Col6a1 | 1.50E-10 | 2.501075558 | 3.73E-06 | 0.796 | 0.082 |
| **155** | Arhgef17 | 1.60E-10 | 5.829422689 | 3.99E-06 | 0.612 | 0 |
| **156** | Aspn | 1.60E-10 | 6.414292791 | 3.99E-06 | 0.612 | 0 |
| **157** | Mxra7 | 1.60E-10 | 5.53367516 | 3.99E-06 | 0.612 | 0 |
| **158** | Rpl36a | 1.60E-10 | 6.054657672 | 3.99E-06 | 0.612 | 0 |
| **159** | Rps25 | 1.60E-10 | 5.735114888 | 3.99E-06 | 0.612 | 0 |
| **160** | S100a11 | 1.60E-10 | 5.627650432 | 3.99E-06 | 0.612 | 0 |
| **161** | Dlc1 | 1.64E-10 | 1.457296129 | 4.07E-06 | 1 | 0.776 |

| | | | | | | |
|---|---|---|---|---|---|---|
| **162** | Ddr2 | 1.68E-10 | 1.745122849 | 4.19E-06 | 0.898 | 0.143 |
| **163** | Rps4x | 1.77E-10 | 1.939384857 | 4.40E-06 | 0.878 | 0.122 |
| **164** | Hnrnpul2 | 1.84E-10 | 2.142568615 | 4.57E-06 | 0.796 | 0.082 |
| **165** | Rpl22 | 1.84E-10 | 4.355560254 | 4.57E-06 | 0.653 | 0.02 |
| **166** | Hspa5 | 1.91E-10 | 2.808842526 | 4.76E-06 | 0.755 | 0.061 |
| **167** | Gpc6 | 2.05E-10 | 2.036456822 | 5.10E-06 | 0.878 | 0.163 |
| **168** | Ighm | 2.05E-10 | 4.409174085 | 5.11E-06 | 0.653 | 0.02 |
| **169** | Anxa1 | 2.24E-10 | 2.390611783 | 5.56E-06 | 0.755 | 0.061 |
| **170** | Rps6 | 2.60E-10 | 2.261647021 | 6.46E-06 | 0.714 | 0.041 |
| **171** | Ryk | 2.79E-10 | 2.942848807 | 6.93E-06 | 0.694 | 0.041 |
| **172** | Fip1l1 | 2.80E-10 | 1.878242789 | 6.97E-06 | 0.837 | 0.102 |
| **173** | Arhgap42 | 2.86E-10 | 2.148942573 | 7.13E-06 | 0.857 | 0.122 |
| **174** | Rpl11 | 3.01E-10 | 2.406073737 | 7.48E-06 | 0.776 | 0.082 |
| **175** | Ncl | 3.27E-10 | 2.7079004 | 8.13E-06 | 0.694 | 0.041 |
| **176** | Nmt1 | 3.36E-10 | 3.078342304 | 8.35E-06 | 0.653 | 0.02 |
| **177** | Hsp90ab1 | 3.53E-10 | 1.996262412 | 8.78E-06 | 0.939 | 0.204 |
| **178** | Hprt | 3.54E-10 | 2.963626498 | 8.82E-06 | 0.653 | 0.02 |
| **179** | Npm1 | 3.54E-10 | 2.751007024 | 8.82E-06 | 0.653 | 0.02 |
| **180** | Ctsb | 3.84E-10 | 1.789107243 | 9.57E-06 | 0.857 | 0.122 |
| **181** | Atxn7l3b | 4.04E-10 | 5.128830973 | 1.01E-05 | 0.592 | 0 |
| **182** | Psip1 | 4.04E-10 | 5.156845058 | 1.01E-05 | 0.592 | 0 |
| **183** | Rtn4rl1 | 4.04E-10 | 4.8518221 | 1.01E-05 | 0.592 | 0 |
| **184** | Rps14 | 4.47E-10 | 2.45992323 | 1.11E-05 | 0.796 | 0.102 |
| **185** | Rpl24 | 4.48E-10 | 2.652834097 | 1.12E-05 | 0.694 | 0.041 |
| **186** | Itpripl2 | 4.51E-10 | 1.860026809 | 1.12E-05 | 0.776 | 0.082 |
| **187** | Mrvi1 | 4.55E-10 | 1.845520222 | 1.13E-05 | 0.816 | 0.102 |
| **188** | Chd4 | 4.60E-10 | 1.956441164 | 1.14E-05 | 0.857 | 0.102 |
| **189** | Eif3a | 4.60E-10 | 2.323303928 | 1.14E-05 | 0.837 | 0.122 |
| **190** | Ssb | 4.72E-10 | 3.197191718 | 1.18E-05 | 0.694 | 0.041 |
| **191** | Col3a1 | 5.08E-10 | 1.798246006 | 1.26E-05 | 1 | 0.265 |
| **192** | Zbtb7a | 5.24E-10 | 2.747044686 | 1.30E-05 | 0.694 | 0.041 |
| **193** | Sumo2 | 5.25E-10 | 2.732557536 | 1.31E-05 | 0.735 | 0.061 |
| **194** | Chn1 | 5.42E-10 | 4.284449408 | 1.35E-05 | 0.633 | 0.02 |
| **195** | Dtx3 | 5.53E-10 | 2.968850493 | 1.38E-05 | 0.694 | 0.041 |
| **196** | Dynll1 | 5.72E-10 | 4.168026816 | 1.42E-05 | 0.633 | 0.02 |
| **197** | Klf2 | 5.82E-10 | 2.226209679 | 1.45E-05 | 0.694 | 0.041 |
| **198** | Ppia | 6.16E-10 | 2.686843809 | 1.53E-05 | 0.776 | 0.102 |
| **199** | Rpl32 | 6.20E-10 | 3.339158549 | 1.54E-05 | 0.673 | 0.041 |
| **200** | Sparc | 6.30E-10 | 2.033637206 | 1.57E-05 | 1 | 0.469 |
| **201** | Rpl26 | 6.53E-10 | 3.321604515 | 1.63E-05 | 0.673 | 0.041 |
| **202** | Rabac1 | 6.73E-10 | 3.838641858 | 1.67E-05 | 0.633 | 0.02 |

| | | | | | | |
|---|---|---|---|---|---|---|
| **203** | Enox1 | 6.75E-10 | 1.945766589 | 1.68E-05 | 0.816 | 0.102 |
| **204** | Rpl23 | 6.79E-10 | 2.299791402 | 1.69E-05 | 0.837 | 0.122 |
| **205** | Rplp0 | 7.17E-10 | 2.641698753 | 1.78E-05 | 0.694 | 0.041 |
| **206** | H2-D1 | 7.62E-10 | 1.958200272 | 1.90E-05 | 0.98 | 0.306 |
| **207** | Gstm1 | 8.06E-10 | 3.258743467 | 2.00E-05 | 0.673 | 0.041 |
| **208** | Dse | 8.49E-10 | 3.110332396 | 2.11E-05 | 0.673 | 0.041 |
| **209** | Aldoa | 8.80E-10 | 2.676055244 | 2.19E-05 | 0.633 | 0.02 |
| **210** | Bnip2 | 8.80E-10 | 2.411048308 | 2.19E-05 | 0.633 | 0.02 |
| **211** | Btg1 | 8.80E-10 | 2.949019837 | 2.19E-05 | 0.633 | 0.02 |
| **212** | P4hb | 8.80E-10 | 2.938546213 | 2.19E-05 | 0.633 | 0.02 |
| **213** | Rack1 | 8.80E-10 | 3.375885537 | 2.19E-05 | 0.633 | 0.02 |
| **214** | Sod1 | 8.80E-10 | 2.099364528 | 2.19E-05 | 0.633 | 0.02 |
| **215** | Txn1 | 8.80E-10 | 2.289939323 | 2.19E-05 | 0.633 | 0.02 |
| **216** | Hspa8 | 9.07E-10 | 2.019523293 | 2.26E-05 | 0.837 | 0.122 |
| **217** | Oaz1 | 9.40E-10 | 2.556440459 | 2.34E-05 | 0.714 | 0.061 |
| **218** | H2-T23 | 9.42E-10 | 3.070538467 | 2.34E-05 | 0.673 | 0.041 |
| **219** | Rps10 | 9.55E-10 | 2.996281191 | 2.38E-05 | 0.735 | 0.082 |
| **220** | Calr | 9.89E-10 | 2.831533695 | 2.46E-05 | 0.714 | 0.061 |
| **221** | Atp5e | 9.92E-10 | 2.871043197 | 2.47E-05 | 0.673 | 0.041 |
| **222** | Tenm4 | 9.93E-10 | -2.29013808 | 2.47E-05 | 0.816 | 0.918 |
| **223** | Cdh2 | 1.00E-09 | 6.269814752 | 2.49E-05 | 0.571 | 0 |
| **224** | Lin7a | 1.00E-09 | 6.396091153 | 2.49E-05 | 0.571 | 0 |
| **225** | Naalad2 | 1.00E-09 | 5.053120557 | 2.49E-05 | 0.571 | 0 |
| **226** | Pds5b | 1.00E-09 | 1.923651783 | 2.49E-05 | 0.918 | 0.184 |
| **227** | Zfp36l1 | 1.06E-09 | 1.910903611 | 2.64E-05 | 0.959 | 0.245 |
| **228** | Selenop | 1.11E-09 | 1.719928041 | 2.76E-05 | 0.918 | 0.143 |
| **229** | Nop53 | 1.13E-09 | 3.977394721 | 2.82E-05 | 0.612 | 0.02 |
| **230** | Tomm7 | 1.13E-09 | 3.99460821 | 2.82E-05 | 0.612 | 0.02 |
| **231** | Rhob | 1.15E-09 | 2.057794512 | 2.87E-05 | 0.837 | 0.122 |
| **232** | Arhgap6 | 1.16E-09 | 1.501213209 | 2.89E-05 | 1 | 0.265 |
| **233** | Eif4g2 | 1.20E-09 | 2.113972635 | 2.97E-05 | 0.837 | 0.143 |
| **234** | Sox5 | 1.24E-09 | 0.97859047 | 3.08E-05 | 1 | 0.306 |
| **235** | Tnxb | 1.25E-09 | 1.940927842 | 3.12E-05 | 0.878 | 0.143 |
| **236** | Uqcr11 | 1.33E-09 | 4.084522884 | 3.31E-05 | 0.612 | 0.02 |
| **237** | Cyb5a | 1.34E-09 | 1.290280289 | 3.34E-05 | 0.776 | 0.082 |
| **238** | Ebf2 | 1.37E-09 | 1.676737012 | 3.40E-05 | 0.918 | 0.204 |
| **239** | Ulk2 | 1.41E-09 | 0.429644056 | 3.51E-05 | 0.857 | 0.122 |
| **240** | Hspb1 | 1.48E-09 | 2.447469489 | 3.68E-05 | 0.714 | 0.061 |
| **241** | Lamp1 | 1.58E-09 | 2.307421257 | 3.93E-05 | 0.816 | 0.122 |
| **242** | Eif5b | 1.72E-09 | 2.198715426 | 4.28E-05 | 0.714 | 0.061 |
| **243** | Zfp36l2 | 1.72E-09 | 1.742676964 | 4.28E-05 | 0.714 | 0.061 |

| | | | | | | |
|---|---|---|---|---|---|---|
| **244** | Aoc3 | 1.75E-09 | 2.476931452 | 4.35E-05 | 0.673 | 0.041 |
| **245** | Jund | 1.80E-09 | 1.484252314 | 4.48E-05 | 0.776 | 0.082 |
| **246** | Rbbp7 | 1.83E-09 | 3.779081442 | 4.56E-05 | 0.612 | 0.02 |
| **247** | Ltbp1 | 1.84E-09 | 1.77918755 | 4.59E-05 | 0.796 | 0.102 |
| **248** | Arhgap32 | 1.91E-09 | 1.45422885 | 4.75E-05 | 0.878 | 0.143 |
| **249** | Podxl | 1.98E-09 | 1.526558115 | 4.94E-05 | 0.776 | 0.082 |
| **250** | Jun | 2.06E-09 | 3.139107406 | 5.11E-05 | 0.653 | 0.041 |
| **251** | Ets2 | 2.06E-09 | 2.311127573 | 5.12E-05 | 0.755 | 0.082 |
| **252** | Egflam | 2.15E-09 | 2.144728427 | 5.34E-05 | 0.612 | 0.02 |
| **253** | Serping1 | 2.15E-09 | 2.65514874 | 5.34E-05 | 0.612 | 0.02 |
| **254** | Unc45b | 2.15E-09 | 2.11598272 | 5.34E-05 | 0.612 | 0.02 |
| **255** | Tmf1 | 2.38E-09 | 1.776077329 | 5.92E-05 | 0.755 | 0.082 |
| **256** | Calm3 | 2.43E-09 | 4.811611234 | 6.06E-05 | 0.551 | 0 |
| **257** | Cox7a2l | 2.43E-09 | 5.17952763 | 6.06E-05 | 0.551 | 0 |
| **258** | Crip2 | 2.43E-09 | 5.153712901 | 6.06E-05 | 0.551 | 0 |
| **259** | Daam2 | 2.43E-09 | 5.801533981 | 6.06E-05 | 0.551 | 0 |
| **260** | Ednra | 2.43E-09 | 5.318565054 | 6.06E-05 | 0.551 | 0 |
| **261** | H2afz | 2.43E-09 | 4.910958774 | 6.06E-05 | 0.551 | 0 |
| **262** | Rpl8 | 2.43E-09 | 5.791562039 | 6.06E-05 | 0.551 | 0 |
| **263** | Cdc5l | 2.44E-09 | 0.906388325 | 6.07E-05 | 0.714 | 0.061 |
| **264** | Pttg1ip | 2.48E-09 | 1.004670096 | 6.18E-05 | 0.857 | 0.122 |
| **265** | Cygb | 2.65E-09 | 3.991431149 | 6.59E-05 | 0.633 | 0.041 |
| **266** | Nfkbia | 2.80E-09 | 2.273814617 | 6.96E-05 | 0.653 | 0.041 |
| **267** | Lamb1 | 2.99E-09 | 1.494769901 | 7.44E-05 | 0.857 | 0.122 |
| **268** | Axl | 3.04E-09 | 4.175715461 | 7.57E-05 | 0.592 | 0.02 |
| **269** | Tm4sf1 | 3.06E-09 | 2.135705318 | 7.62E-05 | 0.837 | 0.143 |
| **270** | Ccdc85b | 3.50E-09 | 2.029191589 | 8.72E-05 | 0.755 | 0.082 |
| **271** | Canx | 3.56E-09 | 1.790605791 | 8.86E-05 | 0.918 | 0.204 |
| **272** | Acat1 | 3.61E-09 | 2.030130703 | 8.97E-05 | 0.653 | 0.041 |
| **273** | Vcp | 3.61E-09 | 2.15200183 | 8.97E-05 | 0.735 | 0.082 |
| **274** | Apoe | 3.73E-09 | 2.323611729 | 9.29E-05 | 0.857 | 0.163 |
| **275** | Pdia4 | 3.76E-09 | 3.963697033 | 9.35E-05 | 0.592 | 0.02 |
| **276** | Plcl1 | 3.87E-09 | 1.443688858 | 9.63E-05 | 0.939 | 0.224 |
| **277** | Antxr1 | 3.97E-09 | 2.046200403 | 9.88E-05 | 0.735 | 0.082 |
| **278** | Ror1 | 4.03E-09 | 1.472984475 | 0.000100241 | 0.878 | 0.163 |
| **279** | Itgb1 | 4.04E-09 | 1.665760036 | 0.000100403 | 1 | 0.327 |
| **280** | Gpx4 | 4.24E-09 | 1.564482587 | 0.000105595 | 0.755 | 0.082 |
| **281** | Glyr1 | 4.29E-09 | 1.792188283 | 0.000106799 | 0.694 | 0.061 |
| **282** | S100a16 | 4.64E-09 | 3.110622197 | 0.000115441 | 0.592 | 0.02 |
| **283** | Myl12a | 4.74E-09 | 2.970627902 | 0.000117822 | 0.694 | 0.082 |
| **284** | Akr1a1 | 4.74E-09 | 2.326664154 | 0.000117831 | 0.694 | 0.061 |

| | | | | | | |
|---|---|---|---|---|---|---|
| **285** | Cfl2 | 4.89E-09 | 3.004293914 | 0.00012165 | 0.592 | 0.02 |
| **286** | Rpl12 | 4.89E-09 | 3.007138356 | 0.00012165 | 0.592 | 0.02 |
| **287** | Atp5j2 | 5.15E-09 | 2.13378515 | 0.000128184 | 0.592 | 0.02 |
| **288** | Gnb2 | 5.15E-09 | 3.109024345 | 0.000128184 | 0.592 | 0.02 |
| **289** | Ndrg2 | 5.15E-09 | 2.982736552 | 0.000128184 | 0.592 | 0.02 |
| **290** | Rpl15 | 5.15E-09 | 2.772553912 | 0.000128184 | 0.592 | 0.02 |
| **291** | S100a13 | 5.15E-09 | 2.994034096 | 0.000128184 | 0.592 | 0.02 |
| **292** | Tfcp2 | 5.15E-09 | 2.934450979 | 0.000128184 | 0.592 | 0.02 |
| **293** | Ccnt1 | 5.56E-09 | 1.980807115 | 0.000138308 | 0.735 | 0.082 |
| **294** | Snx18 | 5.76E-09 | 1.691371234 | 0.00014332 | 0.694 | 0.061 |
| **295** | Atp5k | 5.83E-09 | 5.206916277 | 0.000144946 | 0.531 | 0 |
| **296** | Sdha | 5.92E-09 | 0.716461215 | 0.000147388 | 0.755 | 0.082 |
| **297** | Irf2bp2 | 6.10E-09 | 1.19082504 | 0.000151858 | 0.755 | 0.102 |
| **298** | Myh11 | 6.30E-09 | 2.630760633 | 0.000156753 | 0.633 | 0.041 |
| **299** | Tgoln1 | 6.41E-09 | 1.992096411 | 0.000159598 | 0.735 | 0.082 |
| **300** | Tuba1b | 6.45E-09 | 2.510704617 | 0.000160515 | 0.673 | 0.061 |
| **301** | Zfand5 | 6.51E-09 | 1.384844004 | 0.000162093 | 0.837 | 0.122 |
| **302** | Adipoq | 6.60E-09 | 1.368597144 | 0.000164326 | 0.796 | 0.102 |
| **303** | Rpl7a | 6.91E-09 | 3.486842362 | 0.000171999 | 0.612 | 0.041 |
| **304** | Khdrbs1 | 6.96E-09 | 1.449255737 | 0.000173068 | 0.939 | 0.184 |
| **305** | Scp2 | 6.97E-09 | 1.61709388 | 0.00017336 | 0.633 | 0.041 |
| **306** | Lrp1 | 7.40E-09 | 1.355972339 | 0.000184143 | 0.898 | 0.163 |
| **307** | Cavin1 | 7.72E-09 | 1.719722997 | 0.000192037 | 0.98 | 0.306 |
| **308** | Nus1 | 7.95E-09 | 0.343020942 | 0.000197775 | 0.796 | 0.102 |
| **309** | Mrfap1 | 8.02E-09 | 3.881256472 | 0.000199561 | 0.571 | 0.02 |
| **310** | Pim3 | 8.10E-09 | 2.404173865 | 0.000201521 | 0.633 | 0.041 |
| **311** | 1810037I17Rik | 8.51E-09 | 2.109245285 | 0.000211861 | 0.633 | 0.041 |
| **312** | Kirrel | 8.56E-09 | 1.928292517 | 0.000213057 | 0.816 | 0.143 |
| **313** | Klf6 | 8.56E-09 | 1.976658647 | 0.000213057 | 0.816 | 0.143 |
| **314** | Eps15l1 | 8.63E-09 | 1.529166361 | 0.000214764 | 0.776 | 0.102 |
| **315** | Ednrb | 8.90E-09 | 3.78008535 | 0.000221506 | 0.571 | 0.02 |
| **316** | Tmed2 | 8.94E-09 | 1.906717914 | 0.00022242 | 0.735 | 0.082 |
| **317** | Ano1 | 9.36E-09 | 2.922622757 | 0.000232782 | 0.673 | 0.082 |
| **318** | Abi2 | 9.37E-09 | 1.454707986 | 0.000233162 | 0.735 | 0.082 |
| **319** | Bzw1 | 9.37E-09 | 1.473561563 | 0.000233162 | 0.735 | 0.082 |
| **320** | M6pr | 9.38E-09 | 3.546691089 | 0.000233341 | 0.571 | 0.02 |
| **321** | Gucy1b1 | 9.88E-09 | 3.6503143 | 0.00024579 | 0.571 | 0.02 |
| **322** | Enpep | 9.89E-09 | 2.219200661 | 0.000246072 | 0.633 | 0.041 |
| **323** | Sp1 | 1.00E-08 | 0.815987855 | 0.000248986 | 0.796 | 0.102 |
| **324** | Itm2b | 1.00E-08 | 1.601953704 | 0.000249896 | 1 | 0.429 |
| **325** | Znfx1 | 1.03E-08 | 0.949744694 | 0.000256181 | 0.735 | 0.082 |

| | | | | | | |
|---|---|---|---|---|---|---|
| **326** | Ptms | 1.04E-08 | 2.295227414 | 0.000258398 | 0.755 | 0.122 |
| **327** | Chchd2 | 1.04E-08 | 3.370101974 | 0.000258883 | 0.571 | 0.02 |
| **328** | Cnot6 | 1.05E-08 | 0.907585253 | 0.000260675 | 0.796 | 0.102 |
| **329** | Safb | 1.08E-08 | 1.734407429 | 0.000268504 | 0.735 | 0.082 |
| **330** | Psap | 1.09E-08 | 1.705987525 | 0.000271179 | 0.98 | 0.347 |
| **331** | Rps19 | 1.09E-08 | 2.309366634 | 0.000271613 | 0.735 | 0.102 |
| **332** | Ndufa2 | 1.10E-08 | 3.200723461 | 0.000272653 | 0.571 | 0.02 |
| **333** | Rps17 | 1.10E-08 | 3.15329616 | 0.000272653 | 0.571 | 0.02 |
| **334** | Akap12 | 1.10E-08 | 1.0800538 | 0.00027347 | 0.837 | 0.143 |
| **335** | Arl6ip1 | 1.15E-08 | 3.295849137 | 0.000287134 | 0.571 | 0.02 |
| **336** | Os9 | 1.15E-08 | 3.204200578 | 0.000287134 | 0.571 | 0.02 |
| **337** | C1qtnf9 | 1.16E-08 | 2.20532366 | 0.000287854 | 0.673 | 0.061 |
| **338** | Sf3b2 | 1.16E-08 | 1.237880783 | 0.000287854 | 0.673 | 0.061 |
| **339** | Vim | 1.16E-08 | 1.694308454 | 0.000289345 | 0.939 | 0.224 |
| **340** | Brd3 | 1.22E-08 | 2.14172542 | 0.000302362 | 0.571 | 0.02 |
| **341** | Sgk1 | 1.22E-08 | 2.211654247 | 0.000302362 | 0.571 | 0.02 |
| **342** | Ypel3 | 1.22E-08 | 2.360217262 | 0.000302362 | 0.571 | 0.02 |
| **343** | Dmd | 1.23E-08 | 2.192602931 | 0.000305245 | 0.918 | 0.286 |
| **344** | Cox4i1 | 1.23E-08 | 1.6413518 | 0.000305648 | 0.755 | 0.102 |
| **345** | Eef1b2 | 1.33E-08 | 2.959094827 | 0.000330855 | 0.612 | 0.041 |
| **346** | Grn | 1.33E-08 | 3.071230957 | 0.000330855 | 0.612 | 0.041 |
| **347** | Pank3 | 1.34E-08 | 1.847201366 | 0.000332615 | 0.673 | 0.061 |
| **348** | Ndufb9 | 1.37E-08 | 4.791662116 | 0.000341129 | 0.51 | 0 |
| **349** | Pfdn5 | 1.37E-08 | 5.020467344 | 0.000341129 | 0.51 | 0 |
| **350** | Rpl29 | 1.37E-08 | 5.443178549 | 0.000341129 | 0.51 | 0 |
| **351** | Tram1 | 1.40E-08 | 3.002404603 | 0.000347758 | 0.612 | 0.041 |
| **352** | AC149090.1 | 1.41E-08 | 1.86848887 | 0.00035097 | 0.755 | 0.102 |
| **353** | Bicc1 | 1.51E-08 | 1.531242448 | 0.000374982 | 0.796 | 0.122 |
| **354** | Atp5h | 1.54E-08 | 2.892556998 | 0.000384118 | 0.612 | 0.041 |
| **355** | Pcdh19 | 1.55E-08 | 1.856276579 | 0.000384743 | 0.755 | 0.102 |
| **356** | Sdf4 | 1.62E-08 | 1.725901241 | 0.000402936 | 0.673 | 0.061 |
| **357** | Skp1a | 1.62E-08 | 2.91292145 | 0.000403657 | 0.612 | 0.041 |
| **358** | Clic1 | 1.70E-08 | 1.180295552 | 0.000422657 | 0.673 | 0.061 |
| **359** | Klf9 | 1.71E-08 | 1.798615213 | 0.000426427 | 0.918 | 0.245 |
| **360** | Prrx1 | 1.77E-08 | 1.812506147 | 0.000441393 | 0.755 | 0.102 |
| **361** | Ppp1r9b | 1.78E-08 | 1.118800988 | 0.000443315 | 0.673 | 0.061 |
| **362** | Ikbkb | 1.79E-08 | 2.142443126 | 0.000445671 | 0.612 | 0.041 |
| **363** | Pea15a | 1.88E-08 | 2.083926564 | 0.000468242 | 0.612 | 0.041 |
| **364** | Atp5l | 1.96E-08 | 3.359591626 | 0.000486648 | 0.592 | 0.041 |
| **365** | Gns | 1.97E-08 | 3.796296592 | 0.000490014 | 0.551 | 0.02 |
| **366** | Meis1 | 1.97E-08 | 1.737240917 | 0.000491078 | 0.776 | 0.143 |

| | | | | | | |
|---|---|---|---|---|---|---|
| **367** | Abcb7 | 1.98E-08 | 2.148857091 | 0.000493657 | 0.694 | 0.082 |
| **368** | Retn | 1.99E-08 | 1.904727438 | 0.00049539 | 0.735 | 0.102 |
| **369** | Rpl14 | 2.07E-08 | 3.712467489 | 0.000515911 | 0.551 | 0.02 |
| **370** | Dag1 | 2.14E-08 | 0.752364872 | 0.000532612 | 0.837 | 0.143 |
| **371** | Ecm2 | 2.19E-08 | 2.477343007 | 0.000543915 | 0.653 | 0.061 |
| **372** | Txnip | 2.26E-08 | 1.906995092 | 0.000562427 | 0.939 | 0.449 |
| **373** | Pakap | 2.29E-08 | 1.437849377 | 0.000570143 | 0.612 | 0.041 |
| **374** | Rbms3 | 2.33E-08 | 1.480193232 | 0.000580098 | 0.98 | 0.306 |
| **375** | Rnase4 | 2.51E-08 | 3.241957338 | 0.000623743 | 0.592 | 0.041 |
| **376** | Mlf2 | 2.55E-08 | 3.357471642 | 0.000633438 | 0.551 | 0.02 |
| **377** | Abca8a | 2.65E-08 | 2.111907407 | 0.000658172 | 0.653 | 0.061 |
| **378** | Cyp4b1 | 2.72E-08 | 1.819020587 | 0.000677953 | 0.776 | 0.163 |
| **379** | Cox6c | 2.75E-08 | 1.888623265 | 0.000683939 | 0.694 | 0.082 |
| **380** | Capns1 | 2.77E-08 | 1.996588 | 0.000688498 | 0.592 | 0.041 |
| **381** | Prpf8 | 2.77E-08 | 2.994517277 | 0.000688498 | 0.592 | 0.041 |
| **382** | Ywhaz | 2.79E-08 | 1.52204956 | 0.00069323 | 0.898 | 0.224 |
| **383** | Btf3 | 2.82E-08 | 1.449190108 | 0.000701563 | 0.551 | 0.02 |
| **384** | Ndufs5 | 2.82E-08 | 1.941928726 | 0.000701563 | 0.551 | 0.02 |
| **385** | Plxna1 | 2.82E-08 | 1.573082108 | 0.000701563 | 0.551 | 0.02 |
| **386** | Purb | 2.95E-08 | 1.980734061 | 0.000734723 | 0.755 | 0.122 |
| **387** | Ahnak | 3.01E-08 | 1.30209208 | 0.00074863 | 0.98 | 0.633 |
| **388** | Bnip3l | 3.05E-08 | 2.466607164 | 0.000759756 | 0.592 | 0.041 |
| **389** | Sema5a | 3.07E-08 | 1.498039907 | 0.000764676 | 0.837 | 0.163 |
| **390** | C3 | 3.10E-08 | 2.070425711 | 0.000770159 | 0.755 | 0.143 |
| **391** | Ubr2 | 3.13E-08 | 1.180834585 | 0.000779051 | 0.898 | 0.163 |
| **392** | Mamdc2 | 3.18E-08 | 5.567560031 | 0.000790056 | 0.49 | 0 |
| **393** | Per1 | 3.18E-08 | 4.843020755 | 0.000790056 | 0.49 | 0 |
| **394** | Psme1 | 3.18E-08 | 5.134448988 | 0.000790056 | 0.49 | 0 |
| **395** | Ralgds | 3.18E-08 | 4.524488806 | 0.000790056 | 0.49 | 0 |
| **396** | Rgs4 | 3.18E-08 | 5.231927369 | 0.000790056 | 0.49 | 0 |
| **397** | Rnasek | 3.18E-08 | 4.811492785 | 0.000790056 | 0.49 | 0 |
| **398** | Extl3 | 3.22E-08 | 1.018455825 | 0.000800829 | 0.796 | 0.122 |
| **399** | Pros1 | 3.22E-08 | 0.985169248 | 0.000800829 | 0.796 | 0.122 |
| **400** | Srsf2 | 3.35E-08 | 1.722744513 | 0.000834057 | 0.653 | 0.061 |
| **401** | Efcab14 | 3.38E-08 | 2.634681397 | 0.000839876 | 0.633 | 0.061 |
| **402** | Cxcl12 | 3.47E-08 | 2.00491756 | 0.000862389 | 0.898 | 0.286 |
| **403** | Clec3b | 3.51E-08 | 2.362797245 | 0.000874553 | 0.673 | 0.082 |
| **404** | Tnfrsf1a | 3.54E-08 | 1.660263682 | 0.000880799 | 0.633 | 0.061 |
| **405** | Trf | 3.83E-08 | 2.22373679 | 0.000953894 | 0.714 | 0.102 |
| **406** | Uqcrh | 3.90E-08 | 2.411870804 | 0.000970633 | 0.592 | 0.041 |
| **407** | 6720427I07Rik | 4.05E-08 | 1.46038551 | 0.001006836 | 0.653 | 0.061 |

| | | | | | | |
|---|---|---|---|---|---|---|
| **408** | Ssr1 | 4.05E-08 | 1.456260769 | 0.001006836 | 0.653 | 0.061 |
| **409** | Ngf | 4.10E-08 | 2.229631275 | 0.001019148 | 0.592 | 0.041 |
| **410** | Eif4a1 | 4.11E-08 | 1.518134481 | 0.001023322 | 0.735 | 0.102 |
| **411** | Mrc2 | 4.23E-08 | 3.318764281 | 0.001053104 | 0.571 | 0.041 |
| **412** | AW112010 | 4.30E-08 | 2.163223226 | 0.001070011 | 0.592 | 0.041 |
| **413** | Med23 | 4.30E-08 | 2.835347398 | 0.001070011 | 0.592 | 0.041 |
| **414** | Pdia6 | 4.30E-08 | 1.398363479 | 0.001070011 | 0.592 | 0.041 |
| **415** | Erap1 | 4.37E-08 | 1.248164455 | 0.001087652 | 0.755 | 0.102 |
| **416** | Pard3 | 4.49E-08 | 0.614052243 | 0.001117429 | 0.918 | 0.224 |
| **417** | Hsp90aa1 | 4.51E-08 | 2.324890933 | 0.001123331 | 0.592 | 0.041 |
| **418** | Set | 4.52E-08 | 3.786276502 | 0.001123728 | 0.531 | 0.02 |
| **419** | Adgra3 | 4.64E-08 | 1.184891588 | 0.001153715 | 0.673 | 0.082 |
| **420** | Rdx | 4.73E-08 | 0.928399179 | 0.001176843 | 0.878 | 0.163 |
| **421** | Atp5g3 | 4.74E-08 | 2.256064044 | 0.001179223 | 0.592 | 0.041 |
| **422** | Cd81 | 4.79E-08 | 1.799117811 | 0.001191989 | 0.857 | 0.204 |
| **423** | Ptpn13 | 4.93E-08 | 2.202923004 | 0.001226535 | 0.633 | 0.061 |
| **424** | Nfe2l1 | 4.93E-08 | 1.659455841 | 0.001227825 | 0.857 | 0.184 |
| **425** | C1qtnf1 | 5.22E-08 | 0.664708484 | 0.001299207 | 0.592 | 0.041 |
| **426** | Dusp6 | 5.22E-08 | 2.264709613 | 0.001299207 | 0.592 | 0.041 |
| **427** | Gbp7 | 5.22E-08 | 1.542101854 | 0.001299207 | 0.592 | 0.041 |
| **428** | Tagln2 | 5.22E-08 | 1.401876057 | 0.001299207 | 0.592 | 0.041 |
| **429** | Myl6 | 5.36E-08 | 1.849310421 | 0.001332751 | 0.918 | 0.286 |
| **430** | Yy1 | 5.37E-08 | 0.531931191 | 0.00133629 | 0.878 | 0.163 |
| **431** | Zfp91 | 5.38E-08 | 1.501151708 | 0.001337859 | 0.735 | 0.102 |
| **432** | Pfn1 | 5.57E-08 | 1.948338038 | 0.001385943 | 0.673 | 0.082 |
| **433** | Cdkn1b | 5.68E-08 | 1.972265061 | 0.001412105 | 0.633 | 0.061 |
| **434** | Ostf1 | 5.68E-08 | 1.206989196 | 0.001414472 | 0.796 | 0.122 |
| **435** | Arl6ip5 | 5.72E-08 | 1.134122575 | 0.001424184 | 0.694 | 0.082 |
| **436** | Tpm4 | 6.11E-08 | 1.520124464 | 0.001519165 | 0.837 | 0.163 |
| **437** | Arf5 | 6.12E-08 | 3.108077856 | 0.001523079 | 0.531 | 0.02 |
| **438** | Adap2 | 6.14E-08 | 1.24494568 | 0.00152845 | 0.735 | 0.102 |
| **439** | Atp5mpl | 6.44E-08 | 1.795366048 | 0.001601815 | 0.531 | 0.02 |
| **440** | Gabarapl1 | 6.44E-08 | 2.319943571 | 0.001601815 | 0.531 | 0.02 |
| **441** | Ndufb1-ps | 6.44E-08 | 2.271162867 | 0.001601815 | 0.531 | 0.02 |
| **442** | Ost4 | 6.44E-08 | 2.221047337 | 0.001601815 | 0.531 | 0.02 |
| **443** | Lasp1 | 6.50E-08 | 1.249542498 | 0.001618277 | 0.816 | 0.143 |
| **444** | Naca | 6.68E-08 | 1.939050659 | 0.001663195 | 0.673 | 0.082 |
| **445** | Igfbp7 | 7.08E-08 | 1.514720819 | 0.001762365 | 0.959 | 0.327 |
| **446** | Anp32e | 7.24E-08 | 4.466953558 | 0.001801317 | 0.469 | 0 |
| **447** | Camk2n1 | 7.24E-08 | 4.772188421 | 0.001801317 | 0.469 | 0 |
| **448** | Chmp2a | 7.24E-08 | 4.709957476 | 0.001801317 | 0.469 | 0 |

| | | | | | | |
|---|---|---|---|---|---|---|
| **449** | Creg1 | 7.24E-08 | 4.526310786 | 0.001801317 | 0.469 | 0 |
| **450** | Lgals1 | 7.24E-08 | 5.312817238 | 0.001801317 | 0.469 | 0 |
| **451** | Nol7 | 7.24E-08 | 4.772605505 | 0.001801317 | 0.469 | 0 |
| **452** | Plod3 | 7.24E-08 | 4.436178963 | 0.001801317 | 0.469 | 0 |
| **453** | Ppfia2 | 7.24E-08 | 5.442997997 | 0.001801317 | 0.469 | 0 |
| **454** | Ubxn1 | 7.24E-08 | 4.345783081 | 0.001801317 | 0.469 | 0 |
| **455** | Zbtb11 | 7.24E-08 | 4.951593246 | 0.001801317 | 0.469 | 0 |
| **456** | Gpx3 | 7.32E-08 | 1.827546032 | 0.001821267 | 0.673 | 0.082 |
| **457** | Tmed10 | 7.49E-08 | 1.124317012 | 0.001863909 | 0.837 | 0.143 |
| **458** | Cdk11b | 7.51E-08 | 1.45465541 | 0.001868288 | 0.633 | 0.061 |
| **459** | Naa50 | 7.51E-08 | 1.571711514 | 0.001868288 | 0.633 | 0.061 |
| **460** | Csnk1g3 | 7.54E-08 | 0.901449553 | 0.001877118 | 0.837 | 0.163 |
| **461** | Atp1a1 | 7.60E-08 | 2.941823183 | 0.001890887 | 0.571 | 0.041 |
| **462** | Fgfr1op2 | 7.69E-08 | 0.46560699 | 0.001913605 | 0.796 | 0.122 |
| **463** | Atp5a1 | 7.79E-08 | 1.509072874 | 0.001938229 | 0.755 | 0.122 |
| **464** | Fap | 7.85E-08 | 1.508321511 | 0.001951917 | 0.714 | 0.102 |
| **465** | Ybx1 | 7.87E-08 | 1.499124652 | 0.001957924 | 0.837 | 0.163 |
| **466** | Rarres2 | 7.98E-08 | 2.499492862 | 0.001984451 | 0.571 | 0.041 |
| **467** | Cmpk1 | 8.00E-08 | 1.234407512 | 0.001991669 | 0.735 | 0.102 |
| **468** | Sem1 | 8.14E-08 | 1.549476878 | 0.002024215 | 0.755 | 0.122 |
| **469** | Camk2g | 8.57E-08 | 1.574675587 | 0.002132298 | 0.714 | 0.102 |
| **470** | Trim56 | 8.67E-08 | 1.864604833 | 0.002156173 | 0.612 | 0.061 |
| **471** | Cd44 | 9.13E-08 | 0.457883953 | 0.002271655 | 0.735 | 0.102 |
| **472** | Ramp2 | 9.54E-08 | 2.079907227 | 0.002373183 | 0.714 | 0.122 |
| **473** | Adam23 | 9.67E-08 | 2.272717217 | 0.002405585 | 0.571 | 0.041 |
| **474** | N4bp2l2 | 9.74E-08 | 1.116928269 | 0.002422207 | 0.918 | 0.204 |
| **475** | Gnai2 | 9.95E-08 | 1.229241786 | 0.002474754 | 0.816 | 0.143 |
| **476** | Ephb4 | 9.96E-08 | 1.0154595 | 0.002479097 | 0.735 | 0.102 |
| **477** | Immt | 1.05E-07 | 1.455704452 | 0.002612002 | 0.673 | 0.082 |
| **478** | Arpc5 | 1.06E-07 | 2.353556877 | 0.002647398 | 0.571 | 0.041 |
| **479** | Synm | 1.07E-07 | 3.617793237 | 0.002667419 | 0.51 | 0.02 |
| **480** | Gjc1 | 1.09E-07 | 1.717593593 | 0.002704227 | 0.694 | 0.102 |
| **481** | Gpx1 | 1.09E-07 | 2.328334202 | 0.002720564 | 0.612 | 0.061 |
| **482** | Zc3h13 | 1.11E-07 | 0.791732492 | 0.002759382 | 0.796 | 0.143 |
| **483** | Dusp3 | 1.12E-07 | 1.272183866 | 0.002793513 | 0.857 | 0.163 |
| **484** | Fabp5 | 1.13E-07 | 3.493205015 | 0.002804687 | 0.51 | 0.02 |
| **485** | Ilk | 1.13E-07 | 3.340858805 | 0.002804687 | 0.51 | 0.02 |
| **486** | Sp3 | 1.15E-07 | 0.94473849 | 0.00285391 | 0.837 | 0.163 |
| **487** | Selenof | 1.17E-07 | 1.545158741 | 0.002899714 | 0.714 | 0.102 |
| **488** | Ddx23 | 1.17E-07 | 1.201578514 | 0.00291266 | 0.571 | 0.041 |
| **489** | Socs4 | 1.17E-07 | 1.819037732 | 0.00291266 | 0.571 | 0.041 |

| | | | | | | |
|---|---|---|---|---|---|---|
| **490** | Tmx3 | 1.18E-07 | 0.6237588 | 0.002928398 | 0.816 | 0.143 |
| **491** | Tcf15 | 1.20E-07 | 2.162630024 | 0.002984274 | 0.612 | 0.061 |
| **492** | Pabpn1 | 1.20E-07 | 1.173712447 | 0.002987004 | 0.673 | 0.082 |
| **493** | Cox6b1 | 1.25E-07 | 3.096418386 | 0.003100023 | 0.51 | 0.02 |
| **494** | Gng5 | 1.25E-07 | 3.427533089 | 0.003100023 | 0.51 | 0.02 |
| **495** | Tomm6 | 1.25E-07 | 3.30925823 | 0.003100023 | 0.51 | 0.02 |
| **496** | mt-Co1 | 1.25E-07 | 1.057406744 | 0.003100208 | 1 | 0.898 |
| **497** | Phf12 | 1.26E-07 | 0.895482824 | 0.003131489 | 0.796 | 0.143 |
| **498** | Rps27 | 1.36E-07 | 1.837282122 | 0.003371763 | 0.755 | 0.143 |
| **499** | Col1a2 | 1.38E-07 | 1.15181363 | 0.003425206 | 0.816 | 0.163 |
| **500** | Il34 | 1.38E-07 | 2.922034941 | 0.003425346 | 0.51 | 0.02 |
| **501** | Rsrc2 | 1.38E-07 | 0.626094136 | 0.003444031 | 0.776 | 0.122 |
| **502** | Smc6 | 1.42E-07 | 1.009096311 | 0.003539774 | 0.755 | 0.122 |
| **503** | St13 | 1.42E-07 | 1.20149995 | 0.003539774 | 0.755 | 0.122 |
| **504** | Gabarapl2 | 1.44E-07 | 2.352215178 | 0.003587881 | 0.612 | 0.061 |
| **505** | Rnf152 | 1.44E-07 | 1.554155996 | 0.003587881 | 0.612 | 0.061 |
| **506** | Impad1 | 1.45E-07 | 2.756911188 | 0.003600156 | 0.51 | 0.02 |
| **507** | Kctd12b | 1.45E-07 | 1.880327933 | 0.003600156 | 0.51 | 0.02 |
| **508** | Lbh | 1.45E-07 | 2.512704136 | 0.003600156 | 0.51 | 0.02 |
| **509** | Mlec | 1.45E-07 | 1.9694049 | 0.003600156 | 0.51 | 0.02 |
| **510** | Ppp1r15b | 1.45E-07 | 1.825290575 | 0.003600156 | 0.51 | 0.02 |
| **511** | Uqcrq | 1.45E-07 | 2.177361815 | 0.003600156 | 0.51 | 0.02 |
| **512** | G3bp1 | 1.45E-07 | 1.245731736 | 0.003605059 | 0.714 | 0.102 |
| **513** | Btg3 | 1.48E-07 | 1.199505022 | 0.003693731 | 0.755 | 0.122 |
| **514** | Rbx1 | 1.51E-07 | 1.189045841 | 0.003756321 | 0.612 | 0.061 |
| **515** | Pnpla8 | 1.54E-07 | 1.90510407 | 0.003823484 | 0.714 | 0.122 |
| **516** | Robo1 | 1.54E-07 | 1.996868732 | 0.003840618 | 0.694 | 0.102 |
| **517** | Bsg | 1.55E-07 | 1.110123252 | 0.003854156 | 0.755 | 0.122 |
| **518** | Slfn5 | 1.57E-07 | 1.226132535 | 0.003893967 | 0.98 | 0.408 |
| **519** | Prkd3 | 1.57E-07 | 0.449232179 | 0.00391015 | 0.776 | 0.122 |
| **520** | Gcnt2 | 1.58E-07 | 1.598810909 | 0.00392282 | 0.796 | 0.163 |
| **521** | Map3k20 | 1.60E-07 | 1.186294593 | 0.003977804 | 0.878 | 0.204 |
| **522** | Ifi27l2a | 1.61E-07 | 1.546283289 | 0.004011654 | 0.694 | 0.102 |
| **523** | Atp6ap2 | 1.63E-07 | 4.562276445 | 0.00404467 | 0.449 | 0 |
| **524** | Eif3k | 1.63E-07 | 4.593388554 | 0.00404467 | 0.449 | 0 |
| **525** | Hgf | 1.63E-07 | 4.601166591 | 0.00404467 | 0.449 | 0 |
| **526** | Itm2c | 1.63E-07 | 4.719497521 | 0.00404467 | 0.449 | 0 |
| **527** | Mesd | 1.63E-07 | 4.925369082 | 0.00404467 | 0.449 | 0 |
| **528** | Rem1 | 1.63E-07 | 4.415845513 | 0.00404467 | 0.449 | 0 |
| **529** | Romo1 | 1.63E-07 | 5.111202784 | 0.00404467 | 0.449 | 0 |
| **530** | S100a1 | 1.63E-07 | 4.920712596 | 0.00404467 | 0.449 | 0 |

| | | | | | | |
|---|---|---|---|---|---|---|
| **531** | Smdt1 | 1.63E-07 | 4.855138776 | 0.00404467 | 0.449 | 0 |
| **532** | Suds3 | 1.63E-07 | 4.227337835 | 0.00404467 | 0.449 | 0 |
| **533** | Tmbim4 | 1.63E-07 | 4.765608613 | 0.00404467 | 0.449 | 0 |
| **534** | Ddx3x | 1.63E-07 | 1.490131724 | 0.004056596 | 0.898 | 0.245 |
| **535** | Scd1 | 1.66E-07 | 1.699753559 | 0.004129106 | 1 | 0.429 |
| **536** | Cpd | 1.66E-07 | 1.15951072 | 0.00413871 | 0.918 | 0.224 |
| **537** | Ndufa4 | 1.67E-07 | 1.359642735 | 0.004145183 | 0.653 | 0.082 |
| **538** | Rassf1 | 1.73E-07 | 1.776126665 | 0.004308805 | 0.612 | 0.061 |
| **539** | Tln1 | 1.73E-07 | 1.100925699 | 0.004314772 | 0.837 | 0.163 |
| **540** | Ppp1r10 | 1.90E-07 | 1.727906056 | 0.004719936 | 0.612 | 0.061 |
| **541** | Elob | 1.95E-07 | 2.633317084 | 0.00484163 | 0.551 | 0.041 |
| **542** | Gbp9 | 1.95E-07 | 1.14907367 | 0.00484163 | 0.551 | 0.041 |
| **543** | Eif2s2 | 1.99E-07 | 0.997532308 | 0.004939474 | 0.612 | 0.061 |
| **544** | Tmem184b | 2.00E-07 | 1.059374361 | 0.004967267 | 0.816 | 0.163 |
| **545** | Arcn1 | 2.08E-07 | 1.006539386 | 0.005167221 | 0.796 | 0.143 |
| **546** | Ccdc47 | 2.08E-07 | 1.783666308 | 0.005173894 | 0.653 | 0.082 |
| **547** | Fcgrt | 2.08E-07 | 1.103347455 | 0.005181482 | 0.755 | 0.122 |
| **548** | Gspt1 | 2.12E-07 | 2.053104176 | 0.005276492 | 0.673 | 0.102 |
| **549** | Brip1os | 2.14E-07 | 2.10221966 | 0.005322451 | 0.551 | 0.041 |
| **550** | Hint1 | 2.14E-07 | 2.392665571 | 0.005322451 | 0.551 | 0.041 |
| **551** | Rpl3 | 2.17E-07 | 1.73176586 | 0.00540738 | 0.653 | 0.082 |
| **552** | Ddx42 | 2.23E-07 | 0.492446019 | 0.005546452 | 0.714 | 0.102 |
| **553** | Uqcrb | 2.24E-07 | 1.93821489 | 0.005579853 | 0.551 | 0.041 |
| **554** | Pds5a | 2.27E-07 | 0.755100595 | 0.005650613 | 0.898 | 0.224 |
| **555** | Cyp20a1 | 2.28E-07 | 1.98649871 | 0.005672643 | 0.592 | 0.061 |
| **556** | Cd151 | 2.35E-07 | 1.68293766 | 0.005849263 | 0.551 | 0.041 |
| **557** | Tmx1 | 2.35E-07 | 1.706464939 | 0.005849263 | 0.551 | 0.041 |
| **558** | Atp5f1 | 2.46E-07 | 0.906729802 | 0.006128141 | 0.755 | 0.122 |
| **559** | Ppib | 2.46E-07 | 1.976797606 | 0.006131221 | 0.551 | 0.041 |
| **560** | Vat1 | 2.46E-07 | 1.63251816 | 0.006131221 | 0.551 | 0.041 |
| **561** | Dpysl2 | 2.47E-07 | 0.82208267 | 0.006148609 | 0.857 | 0.184 |
| **562** | Tmem176b | 2.50E-07 | 3.454315733 | 0.006226756 | 0.49 | 0.02 |
| **563** | Tm9sf2 | 2.50E-07 | 1.171806563 | 0.006229401 | 0.816 | 0.184 |
| **564** | Erp29 | 2.58E-07 | 1.549701577 | 0.006426287 | 0.551 | 0.041 |
| **565** | Pip5k1c | 2.58E-07 | 1.305122742 | 0.006426287 | 0.551 | 0.041 |
| **566** | Sgpl1 | 2.58E-07 | 1.66881295 | 0.006426287 | 0.551 | 0.041 |
| **567** | Lbr | 2.59E-07 | 1.409677833 | 0.00644733 | 0.653 | 0.082 |
| **568** | Brd2 | 2.59E-07 | 1.582690181 | 0.00645171 | 0.694 | 0.102 |
| **569** | Irf1 | 2.61E-07 | 2.482478238 | 0.006503378 | 0.592 | 0.061 |
| **570** | Cox5b | 2.63E-07 | 3.340529308 | 0.006542627 | 0.49 | 0.02 |
| **571** | Srsf10 | 2.63E-07 | 1.084853448 | 0.006543079 | 0.735 | 0.122 |

| | | | | | | |
|---|---|---|---|---|---|---|
| **572** | Col5a1 | 2.65E-07 | 1.344635841 | 0.00659802 | 0.755 | 0.143 |
| **573** | Mmrn2 | 2.69E-07 | 1.280050876 | 0.006705302 | 0.878 | 0.204 |
| **574** | Angpt2 | 2.76E-07 | 3.25680986 | 0.006873948 | 0.49 | 0.02 |
| **575** | Tpr | 2.77E-07 | 1.173196405 | 0.006894044 | 0.796 | 0.143 |
| **576** | mt-Nd1 | 2.80E-07 | 1.2548522 | 0.006974085 | 1 | 0.653 |
| **577** | Plod2 | 2.82E-07 | 1.847850465 | 0.007010986 | 0.694 | 0.122 |
| **578** | Glud1 | 2.86E-07 | 0.427332725 | 0.007115433 | 0.735 | 0.122 |
| **579** | Glul | 2.86E-07 | 1.406542761 | 0.007115433 | 0.735 | 0.122 |
| **580** | Rnf10 | 2.86E-07 | 1.80643161 | 0.007122192 | 0.633 | 0.082 |
| **581** | Zfp592 | 2.89E-07 | 1.043664426 | 0.007182279 | 0.796 | 0.143 |
| **582** | B3glct | 2.90E-07 | 3.212338708 | 0.007221446 | 0.49 | 0.02 |
| **583** | Cavin3 | 2.90E-07 | 3.213456675 | 0.007221446 | 0.49 | 0.02 |
| **584** | Ftx | 2.90E-07 | 3.242962035 | 0.007221446 | 0.49 | 0.02 |
| **585** | Ikbip | 2.90E-07 | 3.267772553 | 0.007221446 | 0.49 | 0.02 |
| **586** | Sec62 | 2.91E-07 | 1.267542707 | 0.007240111 | 0.714 | 0.122 |
| **587** | Setbp1 | 2.93E-07 | 1.106155714 | 0.007289612 | 0.837 | 0.184 |
| **588** | Ifngr1 | 2.94E-07 | 1.171010555 | 0.007323164 | 0.837 | 0.163 |
| **589** | Zscan26 | 3.03E-07 | 0.360019677 | 0.007548169 | 0.755 | 0.122 |
| **590** | Grcc10 | 3.05E-07 | 2.689084785 | 0.007585878 | 0.49 | 0.02 |
| **591** | Psme2 | 3.05E-07 | 2.730506209 | 0.007585878 | 0.49 | 0.02 |
| **592** | Trpc3 | 3.05E-07 | 3.141628691 | 0.007600425 | 0.531 | 0.041 |
| **593** | Atpif1 | 3.20E-07 | 0.468138297 | 0.007968037 | 0.49 | 0.02 |
| **594** | Ids | 3.20E-07 | 2.451535156 | 0.007968037 | 0.49 | 0.02 |
| **595** | Sox18 | 3.26E-07 | 2.50622605 | 0.008114913 | 0.571 | 0.061 |
| **596** | Btg2 | 3.28E-07 | 1.519993814 | 0.008155488 | 0.592 | 0.061 |
| **597** | Arf1 | 3.34E-07 | 1.054695178 | 0.008309374 | 0.776 | 0.143 |
| **598** | Ppp4r2 | 3.51E-07 | 0.519986994 | 0.008728774 | 0.857 | 0.163 |
| **599** | Atox1 | 3.52E-07 | 2.77239941 | 0.008755075 | 0.531 | 0.041 |
| **600** | Sclt1 | 3.52E-07 | 2.973764218 | 0.008755075 | 0.531 | 0.041 |
| **601** | Ep300 | 3.53E-07 | 0.901981898 | 0.008775808 | 0.816 | 0.163 |
| **602** | Egfr | 3.56E-07 | 2.387353804 | 0.008848742 | 0.837 | 0.245 |
| **603** | Pdia3 | 3.56E-07 | 1.266043084 | 0.008848742 | 0.878 | 0.204 |
| **604** | Arf6 | 3.60E-07 | 4.436942691 | 0.008947572 | 0.429 | 0 |
| **605** | Eif5a | 3.60E-07 | 4.413957491 | 0.008947572 | 0.429 | 0 |
| **606** | Hrct1 | 3.60E-07 | 4.633927275 | 0.008947572 | 0.429 | 0 |
| **607** | Myl9 | 3.60E-07 | 4.442680111 | 0.008947572 | 0.429 | 0 |
| **608** | Ndufb7 | 3.60E-07 | 4.647455438 | 0.008947572 | 0.429 | 0 |
| **609** | Pde8b | 3.60E-07 | 4.495883644 | 0.008947572 | 0.429 | 0 |
| **610** | Psmb10 | 3.60E-07 | 4.514364329 | 0.008947572 | 0.429 | 0 |
| **611** | Sec24c | 3.60E-07 | 4.332722411 | 0.008947572 | 0.429 | 0 |
| **612** | Selenos | 3.60E-07 | 4.115522042 | 0.008947572 | 0.429 | 0 |

| | | | | | | |
|---|---|---|---|---|---|---|
| **613** | 4632427E13Ri | 3.65E-07 | 0.599159104 | 0.009071591 | 0.694 | 0.102 |
| **614** | 4930523C07Ri | 3.76E-07 | 1.485012819 | 0.009360035 | 0.776 | 0.163 |
| **615** | Calu | 3.82E-07 | 0.92430755 | 0.009510541 | 0.816 | 0.163 |
| **616** | Gsr | 3.87E-07 | 2.022634352 | 0.009617027 | 0.531 | 0.041 |
| **617** | Rpl10a | 3.87E-07 | 1.432476365 | 0.009635472 | 0.673 | 0.102 |
| **618** | Bcam | 3.91E-07 | 2.113505842 | 0.009731913 | 0.571 | 0.061 |
| **619** | Nr1h3 | 3.92E-07 | 1.668508334 | 0.009762216 | 0.592 | 0.061 |
| **620** | Dmxl1 | 4.00E-07 | 0.991655066 | 0.00995763 | 0.878 | 0.204 |
| **621** | Tmbim6 | 4.00E-07 | 1.195253358 | 0.00995763 | 0.878 | 0.204 |
| **622** | Atp6v1a | 4.04E-07 | 1.460685058 | 0.010054185 | 0.673 | 0.102 |
| **623** | Phldb1 | 4.24E-07 | 1.632047903 | 0.010560259 | 0.633 | 0.082 |
| **624** | Wipi1 | 4.24E-07 | 2.362656487 | 0.010560592 | 0.531 | 0.041 |
| **625** | Ube2v2 | 4.28E-07 | 2.259197494 | 0.010652909 | 0.571 | 0.061 |
| **626** | Mff | 4.29E-07 | 1.124327333 | 0.010676141 | 0.592 | 0.061 |
| **627** | Bzw2 | 4.43E-07 | 0.495582118 | 0.011029017 | 0.633 | 0.082 |
| **628** | Ogn | 4.50E-07 | 3.850948994 | 0.011190259 | 0.469 | 0.02 |
| **629** | Srgn | 4.60E-07 | 1.417125129 | 0.011456005 | 0.776 | 0.163 |
| **630** | Ensa | 4.66E-07 | 1.906773802 | 0.011593167 | 0.531 | 0.041 |
| **631** | Fam43a | 4.66E-07 | 2.341862957 | 0.011593167 | 0.531 | 0.041 |
| **632** | Ddx21 | 4.88E-07 | 1.499230139 | 0.01214532 | 0.531 | 0.041 |
| **633** | Spcs2 | 4.88E-07 | 2.314192452 | 0.01214532 | 0.531 | 0.041 |
| **634** | Tmem109 | 4.88E-07 | 2.079360086 | 0.01214532 | 0.531 | 0.041 |
| **635** | Uxs1 | 4.88E-07 | 2.446328694 | 0.01214532 | 0.531 | 0.041 |
| **636** | Wars | 4.88E-07 | 1.553235284 | 0.01214532 | 0.531 | 0.041 |
| **637** | Atp2a2 | 4.96E-07 | 0.882729727 | 0.012331378 | 0.959 | 0.245 |
| **638** | Amotl2 | 4.97E-07 | 1.090390891 | 0.012372056 | 0.735 | 0.143 |
| **639** | Serpinh1 | 4.97E-07 | 1.497805982 | 0.012372056 | 0.735 | 0.143 |
| **640** | Slf2 | 5.00E-07 | 1.206653674 | 0.012452244 | 0.714 | 0.122 |
| **641** | Msl1 | 5.05E-07 | 1.065486081 | 0.012558858 | 0.633 | 0.082 |
| **642** | Mtr | 5.11E-07 | 2.545856945 | 0.012722792 | 0.531 | 0.041 |
| **643** | Dpt | 5.13E-07 | 2.00628677 | 0.012753637 | 0.571 | 0.061 |
| **644** | Atf7 | 5.20E-07 | 1.346067282 | 0.012935628 | 0.918 | 0.224 |
| **645** | Brd8 | 5.21E-07 | 1.033698413 | 0.012960203 | 0.673 | 0.102 |
| **646** | Rev1 | 5.32E-07 | 0.722886166 | 0.013243938 | 0.735 | 0.122 |
| **647** | Arhgef10l | 5.36E-07 | 2.346184681 | 0.013326697 | 0.531 | 0.041 |
| **648** | Gas1 | 5.36E-07 | 1.700088618 | 0.013326697 | 0.531 | 0.041 |
| **649** | Tmem260 | 5.43E-07 | 1.415897535 | 0.013517393 | 0.673 | 0.102 |
| **650** | Rad23b | 5.47E-07 | 0.883109604 | 0.013597918 | 0.857 | 0.184 |
| **651** | Rps15a | 5.50E-07 | 1.842943198 | 0.01368789 | 0.653 | 0.102 |
| **652** | Mon2 | 5.53E-07 | 0.923013609 | 0.013762894 | 0.837 | 0.184 |
| **653** | Mmp14 | 5.55E-07 | 1.972259184 | 0.013804406 | 0.612 | 0.082 |

| | | | | | | |
|---|---|---|---|---|---|---|
| **654** | mt-Nd5 | 5.58E-07 | 1.205402343 | 0.013871245 | 1 | 0.571 |
| **655** | Cggbp1 | 5.61E-07 | 1.516504233 | 0.013958195 | 0.531 | 0.041 |
| **656** | Trappc6b | 5.74E-07 | 1.56593405 | 0.014292346 | 0.633 | 0.082 |
| **657** | Zup1 | 5.74E-07 | 0.923526105 | 0.014292346 | 0.633 | 0.082 |
| **658** | Ntrk2 | 5.86E-07 | 2.122350547 | 0.014585829 | 0.571 | 0.061 |
| **659** | Ubtf | 5.86E-07 | 1.422958697 | 0.014585829 | 0.571 | 0.061 |
| **660** | Jade1 | 6.00E-07 | 1.254505585 | 0.014919819 | 0.633 | 0.082 |
| **661** | Sox7 | 6.00E-07 | 1.035834252 | 0.014919819 | 0.633 | 0.082 |
| **662** | Ap1ar | 6.03E-07 | 3.068807396 | 0.015014873 | 0.469 | 0.02 |
| **663** | D8Ertd738e | 6.03E-07 | 3.102987634 | 0.015014873 | 0.469 | 0.02 |
| **664** | Tspan3 | 6.03E-07 | 3.340884343 | 0.015014873 | 0.469 | 0.02 |
| **665** | Ywhae | 6.04E-07 | 1.152394675 | 0.015018763 | 0.98 | 0.327 |
| **666** | Pten | 6.06E-07 | 1.199049293 | 0.015083703 | 1 | 0.367 |
| **667** | Capn7 | 6.13E-07 | 1.38398377 | 0.015251098 | 0.571 | 0.061 |
| **668** | Piezo2 | 6.13E-07 | 1.179087439 | 0.015251098 | 0.571 | 0.061 |
| **669** | Cp | 6.16E-07 | 0.803524227 | 0.015334113 | 0.816 | 0.163 |
| **670** | Il10rb | 6.26E-07 | 0.712887507 | 0.015573804 | 0.633 | 0.082 |
| **671** | Rab31 | 6.26E-07 | 0.781826864 | 0.015573804 | 0.633 | 0.082 |
| **672** | Spry4 | 6.29E-07 | 2.900767289 | 0.015657778 | 0.51 | 0.041 |
| **673** | Nr1d1 | 6.59E-07 | 2.961886181 | 0.01640492 | 0.51 | 0.041 |
| **674** | Lgals8 | 6.60E-07 | 1.640407282 | 0.016414851 | 0.612 | 0.082 |
| **675** | Rnf7 | 6.65E-07 | 2.677778524 | 0.016549476 | 0.469 | 0.02 |
| **676** | Flna | 6.66E-07 | 1.945237733 | 0.016575978 | 0.714 | 0.143 |
| **677** | Atp5b | 6.70E-07 | 1.827460133 | 0.016670463 | 0.571 | 0.061 |
| **678** | Nucks1 | 6.88E-07 | 1.141739378 | 0.017117762 | 0.735 | 0.143 |
| **679** | Aamp | 6.98E-07 | 2.460060566 | 0.017372395 | 0.469 | 0.02 |
| **680** | Ankle2 | 6.98E-07 | 2.547155703 | 0.017372395 | 0.469 | 0.02 |
| **681** | Arl5a | 6.98E-07 | 2.614951834 | 0.017372395 | 0.469 | 0.02 |
| **682** | Arpc4 | 6.98E-07 | 2.396539754 | 0.017372395 | 0.469 | 0.02 |
| **683** | Golga5 | 6.98E-07 | 2.721340123 | 0.017372395 | 0.469 | 0.02 |
| **684** | Khsrp | 6.98E-07 | 1.6379286 | 0.017372395 | 0.469 | 0.02 |
| **685** | Ndufa5 | 6.98E-07 | 2.116724805 | 0.017372395 | 0.469 | 0.02 |
| **686** | Ppp3r1 | 6.98E-07 | 2.530386183 | 0.017372395 | 0.469 | 0.02 |
| **687** | Slc3a2 | 6.98E-07 | 2.550670753 | 0.017372395 | 0.469 | 0.02 |
| **688** | Vamp8 | 6.98E-07 | 2.438323322 | 0.017372395 | 0.469 | 0.02 |
| **689** | Tpm1 | 7.15E-07 | 1.449066262 | 0.01777875 | 0.776 | 0.163 |
| **690** | Dek | 7.32E-07 | 1.733389249 | 0.018216695 | 0.571 | 0.061 |
| **691** | Ube2z | 7.32E-07 | 0.655994233 | 0.018216695 | 0.571 | 0.061 |
| **692** | Prrc2a | 7.34E-07 | 0.841464351 | 0.018268819 | 0.796 | 0.163 |
| **693** | Ctla2a | 7.58E-07 | 2.438494997 | 0.018858229 | 0.51 | 0.041 |
| **694** | Csnk1e | 7.65E-07 | 1.188287933 | 0.01904074 | 0.571 | 0.061 |

| | | | | | | |
|---|---|---|---|---|---|---|
| **695** | Fscn1 | 7.65E-07 | 1.65625436 | 0.01904074 | 0.571 | 0.061 |
| **696** | Thap3 | 7.65E-07 | 0.858650435 | 0.01904074 | 0.571 | 0.061 |
| **697** | Reep3 | 7.82E-07 | 0.476279122 | 0.019445538 | 0.959 | 0.245 |
| **698** | Ctbp1 | 7.84E-07 | 1.313498966 | 0.019497818 | 0.612 | 0.082 |
| **699** | Alms1 | 7.84E-07 | 4.565489884 | 0.019508642 | 0.408 | 0 |
| **700** | Brox | 7.84E-07 | 4.435167315 | 0.019508642 | 0.408 | 0 |
| **701** | Cops9 | 7.84E-07 | 4.660448508 | 0.019508642 | 0.408 | 0 |
| **702** | Dusp10 | 7.84E-07 | 4.706290771 | 0.019508642 | 0.408 | 0 |
| **703** | Eif3f | 7.84E-07 | 4.430249716 | 0.019508642 | 0.408 | 0 |
| **704** | Gipr | 7.84E-07 | 4.482793175 | 0.019508642 | 0.408 | 0 |
| **705** | Gstm2 | 7.84E-07 | 4.549972835 | 0.019508642 | 0.408 | 0 |
| **706** | Med28 | 7.84E-07 | 4.361266592 | 0.019508642 | 0.408 | 0 |
| **707** | Ndufb8 | 7.84E-07 | 4.39043009 | 0.019508642 | 0.408 | 0 |
| **708** | Pde5a | 7.84E-07 | 5.061166653 | 0.019508642 | 0.408 | 0 |
| **709** | Ptpdc1 | 7.84E-07 | 4.30249573 | 0.019508642 | 0.408 | 0 |
| **710** | Ran | 7.84E-07 | 4.314332435 | 0.019508642 | 0.408 | 0 |
| **711** | Rps27l | 7.84E-07 | 4.771805279 | 0.019508642 | 0.408 | 0 |
| **712** | Selenom | 7.84E-07 | 4.412643717 | 0.019508642 | 0.408 | 0 |
| **713** | Serpine2 | 7.84E-07 | 4.619259131 | 0.019508642 | 0.408 | 0 |
| **714** | Sh3bgrl3 | 7.84E-07 | 4.57350126 | 0.019508642 | 0.408 | 0 |
| **715** | Slc25a5 | 7.84E-07 | 4.267482159 | 0.019508642 | 0.408 | 0 |
| **716** | Sspn | 7.84E-07 | 4.278985773 | 0.019508642 | 0.408 | 0 |
| **717** | Tmem219 | 7.84E-07 | 4.295416327 | 0.019508642 | 0.408 | 0 |
| **718** | Tmem43 | 7.84E-07 | 4.241885829 | 0.019508642 | 0.408 | 0 |
| **719** | Tprgl | 7.84E-07 | 4.53124809 | 0.019508642 | 0.408 | 0 |
| **720** | Vstm4 | 7.84E-07 | 5.078661157 | 0.019508642 | 0.408 | 0 |
| **721** | Zfp672 | 7.84E-07 | 4.275349767 | 0.019508642 | 0.408 | 0 |
| **722** | Eif4a2 | 7.93E-07 | 1.17983767 | 0.019723488 | 0.837 | 0.163 |
| **723** | C4b | 7.94E-07 | 2.130676363 | 0.019751865 | 0.51 | 0.041 |
| **724** | Vezf1 | 8.00E-07 | 1.972052514 | 0.019900634 | 0.571 | 0.061 |
| **725** | Slc25a3 | 8.08E-07 | 1.354816525 | 0.020106267 | 0.735 | 0.143 |
| **726** | Zeb2 | 8.18E-07 | 0.997180905 | 0.020348948 | 0.98 | 0.694 |
| **727** | Tmem167 | 8.18E-07 | 1.531729464 | 0.02035166 | 0.612 | 0.082 |
| **728** | Nars | 8.31E-07 | 1.567587767 | 0.020686218 | 0.51 | 0.041 |
| **729** | Ranbp2 | 8.36E-07 | 0.922054263 | 0.020796265 | 0.878 | 0.204 |
| **730** | Gatad1 | 8.36E-07 | 1.392659165 | 0.02079787 | 0.571 | 0.061 |
| **731** | Glg1 | 8.45E-07 | 1.101201706 | 0.021034223 | 0.878 | 0.224 |
| **732** | Lama2 | 8.49E-07 | 1.694938758 | 0.021122378 | 0.673 | 0.143 |
| **733** | Klhl9 | 8.71E-07 | 2.433114087 | 0.021663065 | 0.51 | 0.041 |
| **734** | Mtch2 | 8.71E-07 | 2.273143385 | 0.021663065 | 0.51 | 0.041 |
| **735** | Rab14 | 8.71E-07 | 1.11102535 | 0.021682791 | 0.857 | 0.204 |

| | | | | | | |
|---|---|---|---|---|---|---|
| **736** | Larp1 | 8.79E-07 | 1.153329624 | 0.021870485 | 0.735 | 0.163 |
| **737** | Xbp1 | 8.91E-07 | 1.073603556 | 0.022168657 | 0.612 | 0.082 |
| **738** | Dock10 | 9.05E-07 | 1.136787229 | 0.022529356 | 0.857 | 0.204 |
| **739** | Kpna6 | 9.12E-07 | 2.234165879 | 0.022684255 | 0.51 | 0.041 |
| **740** | Ptar1 | 9.13E-07 | 1.162941691 | 0.022710638 | 0.571 | 0.061 |
| **741** | Tmed5 | 9.33E-07 | 0.71233411 | 0.023212913 | 0.755 | 0.143 |
| **742** | Ccar1 | 9.48E-07 | 0.830506405 | 0.023590174 | 0.816 | 0.163 |
| **743** | Dync1li2 | 9.49E-07 | 1.12672565 | 0.023613615 | 0.653 | 0.102 |
| **744** | Yme1l1 | 9.89E-07 | 1.211902665 | 0.024614124 | 0.653 | 0.102 |
| **745** | Fbln1 | 9.99E-07 | 2.511792278 | 0.024867447 | 0.51 | 0.041 |
| **746** | Slc24a3 | 1.00E-06 | 1.448007777 | 0.024973751 | 0.918 | 0.306 |
| **747** | Ttc37 | 1.01E-06 | 0.941564075 | 0.025126708 | 0.755 | 0.143 |
| **748** | Mapk1ip1l | 1.01E-06 | 1.435064298 | 0.025190018 | 0.612 | 0.082 |
| **749** | Sptssa | 1.02E-06 | 3.516254401 | 0.025379228 | 0.449 | 0.02 |
| **750** | Vti1b | 1.04E-06 | 2.170613445 | 0.025805477 | 0.551 | 0.061 |
| **751** | Zfp68 | 1.05E-06 | 2.282391825 | 0.026033543 | 0.51 | 0.041 |
| **752** | Anp32b | 1.06E-06 | 1.269052089 | 0.026282466 | 0.612 | 0.082 |
| **753** | Hnrnpa0 | 1.06E-06 | 2.033077144 | 0.026409793 | 0.592 | 0.082 |
| **754** | Slc29a1 | 1.07E-06 | 3.496474174 | 0.026641175 | 0.449 | 0.02 |
| **755** | Lix1l | 1.10E-06 | 1.278505227 | 0.027252175 | 0.51 | 0.041 |
| **756** | Anxa4 | 1.13E-06 | 2.107721321 | 0.028185564 | 0.551 | 0.061 |
| **757** | Hnrnpa2b1 | 1.14E-06 | 1.220154325 | 0.028298164 | 1 | 0.388 |
| **758** | Ccdc82 | 1.15E-06 | 0.664281318 | 0.028489319 | 0.776 | 0.163 |
| **759** | Eif5 | 1.15E-06 | 1.087073957 | 0.028525607 | 0.51 | 0.041 |
| **760** | Cnbp | 1.18E-06 | 0.828520455 | 0.029276998 | 0.878 | 0.204 |
| **761** | Sfr1 | 1.18E-06 | 3.25428152 | 0.029348652 | 0.449 | 0.02 |
| **762** | Tmem50a | 1.20E-06 | 1.34307817 | 0.029840284 | 0.612 | 0.082 |
| **763** | Cox17 | 1.20E-06 | 1.077636126 | 0.029856195 | 0.51 | 0.041 |
| **764** | Pgrmc1 | 1.20E-06 | 2.033122204 | 0.029856195 | 0.51 | 0.041 |
| **765** | Ifnar2 | 1.20E-06 | 1.377959947 | 0.029939261 | 0.735 | 0.143 |
| **766** | Zc3hav1 | 1.22E-06 | 0.990125832 | 0.030345221 | 0.796 | 0.163 |
| **767** | Igfbp4 | 1.23E-06 | 1.843792336 | 0.030581766 | 0.633 | 0.102 |
| **768** | Arfgef2 | 1.27E-06 | 0.619201663 | 0.031540606 | 0.796 | 0.163 |
| **769** | Ceacam1 | 1.29E-06 | 1.87618945 | 0.032155888 | 0.551 | 0.061 |
| **770** | Sri | 1.29E-06 | 1.944497009 | 0.032155888 | 0.551 | 0.061 |
| **771** | Emc7 | 1.30E-06 | 1.311590461 | 0.032464903 | 0.612 | 0.082 |
| **772** | Vdac2 | 1.30E-06 | 0.730315107 | 0.032464903 | 0.612 | 0.082 |
| **773** | Eda | 1.32E-06 | 0.731452578 | 0.032964188 | 0.673 | 0.143 |
| **774** | Rpl31 | 1.36E-06 | 2.927603301 | 0.033911934 | 0.449 | 0.02 |
| **775** | Mndal | 1.38E-06 | 0.948675675 | 0.034239043 | 0.857 | 0.184 |
| **776** | Wtap | 1.38E-06 | 0.270941786 | 0.034323915 | 0.714 | 0.122 |

| | | | | | | |
|---|---|---|---|---|---|---|
| **777** | Cdk6 | 1.39E-06 | 0.330421736 | 0.034692218 | 0.816 | 0.163 |
| **778** | Fbh1 | 1.42E-06 | 0.947752396 | 0.035310829 | 0.612 | 0.082 |
| **779** | Irf2bpl | 1.43E-06 | 2.623222306 | 0.035579291 | 0.449 | 0.02 |
| **780** | Plau | 1.43E-06 | 2.772357638 | 0.035579291 | 0.449 | 0.02 |
| **781** | Eif4ebp2 | 1.44E-06 | 0.83553218 | 0.035785209 | 0.755 | 0.143 |
| **782** | Hnrnph3 | 1.50E-06 | 0.932325757 | 0.037208277 | 0.755 | 0.143 |
| **783** | C1ra | 1.50E-06 | 2.549939694 | 0.037325332 | 0.449 | 0.02 |
| **784** | Cdkn1a | 1.50E-06 | 2.145387687 | 0.037325332 | 0.449 | 0.02 |
| **785** | Cenpb | 1.50E-06 | 1.344722246 | 0.037325332 | 0.449 | 0.02 |
| **786** | Colgalt1 | 1.50E-06 | 2.686794796 | 0.037325332 | 0.449 | 0.02 |
| **787** | Eif3b | 1.50E-06 | 1.466128794 | 0.037325332 | 0.449 | 0.02 |
| **788** | Prrc1 | 1.50E-06 | 2.205718026 | 0.037325332 | 0.449 | 0.02 |
| **789** | Rnps1 | 1.50E-06 | 1.170847853 | 0.037325332 | 0.449 | 0.02 |
| **790** | Sema4c | 1.50E-06 | 2.755450286 | 0.037325332 | 0.449 | 0.02 |
| **791** | Snhg20 | 1.50E-06 | 2.842119888 | 0.037325332 | 0.449 | 0.02 |
| **792** | Snrpg | 1.50E-06 | 2.462416526 | 0.037325332 | 0.449 | 0.02 |
| **793** | Syf2 | 1.50E-06 | 1.441411237 | 0.037325332 | 0.449 | 0.02 |
| **794** | Ube2j1 | 1.50E-06 | 2.301631921 | 0.037325332 | 0.449 | 0.02 |
| **795** | Zfp260 | 1.50E-06 | 2.660831134 | 0.037325332 | 0.449 | 0.02 |
| **796** | Ndufc2 | 1.55E-06 | 0.927541705 | 0.03868866 | 0.653 | 0.102 |
| **797** | Aatf | 1.56E-06 | 1.150993535 | 0.038714683 | 0.592 | 0.082 |
| **798** | Fzd4 | 1.57E-06 | 1.415392464 | 0.03915221 | 0.857 | 0.245 |
| **799** | Synpo2 | 1.59E-06 | 1.061570534 | 0.039462815 | 0.694 | 0.122 |
| **800** | Trim2 | 1.61E-06 | 0.60189573 | 0.040016195 | 0.673 | 0.122 |
| **801** | Ralgapb | 1.63E-06 | 0.670378487 | 0.040635532 | 0.898 | 0.204 |
| **802** | Ddx46 | 1.64E-06 | 1.088923442 | 0.040813336 | 0.633 | 0.102 |
| **803** | mt-Co2 | 1.66E-06 | 0.988498049 | 0.041343102 | 1 | 0.816 |
| **804** | Ndufa3 | 1.66E-06 | 1.836658129 | 0.041384782 | 0.531 | 0.061 |
| **805** | Acsl5 | 1.68E-06 | 1.189578497 | 0.041770524 | 0.551 | 0.061 |
| **806** | Stx5a | 1.68E-06 | 1.750425064 | 0.041770524 | 0.551 | 0.061 |
| **807** | Pcnt | 1.68E-06 | 2.186875752 | 0.041882136 | 0.49 | 0.041 |
| **808** | Atp5o | 1.69E-06 | 4.26723282 | 0.041939368 | 0.388 | 0 |
| **809** | Ccl11 | 1.69E-06 | 4.616159101 | 0.041939368 | 0.388 | 0 |
| **810** | Cdk16 | 1.69E-06 | 4.204081732 | 0.041939368 | 0.388 | 0 |
| **811** | Cspg4 | 1.69E-06 | 4.504225814 | 0.041939368 | 0.388 | 0 |
| **812** | Cycs | 1.69E-06 | 4.60677675 | 0.041939368 | 0.388 | 0 |
| **813** | Dpm3 | 1.69E-06 | 4.69136233 | 0.041939368 | 0.388 | 0 |
| **814** | Dusp7 | 1.69E-06 | 4.587540839 | 0.041939368 | 0.388 | 0 |
| **815** | Emg1 | 1.69E-06 | 3.980564744 | 0.041939368 | 0.388 | 0 |
| **816** | Gpx8 | 1.69E-06 | 4.463014113 | 0.041939368 | 0.388 | 0 |
| **817** | Irs2 | 1.69E-06 | 4.541466264 | 0.041939368 | 0.388 | 0 |

| | | | | | | |
|---|---|---|---|---|---|---|
| **818** | Kcnq4 | 1.69E-06 | 4.465297118 | 0.041939368 | 0.388 | 0 |
| **819** | Lyz2 | 1.69E-06 | 5.19795694 | 0.041939368 | 0.388 | 0 |
| **820** | Mybbp1a | 1.69E-06 | 4.147982706 | 0.041939368 | 0.388 | 0 |
| **821** | Myo7a | 1.69E-06 | 4.49658464 | 0.041939368 | 0.388 | 0 |
| **822** | Ndufaf8 | 1.69E-06 | 4.060511399 | 0.041939368 | 0.388 | 0 |
| **823** | Pdcd6 | 1.69E-06 | 4.13444293 | 0.041939368 | 0.388 | 0 |
| **824** | Pi4k2b | 1.69E-06 | 4.144577462 | 0.041939368 | 0.388 | 0 |
| **825** | Rex1bd | 1.69E-06 | 4.006236548 | 0.041939368 | 0.388 | 0 |
| **826** | Rpl22l1 | 1.69E-06 | 4.319216936 | 0.041939368 | 0.388 | 0 |
| **827** | Serpinf1 | 1.69E-06 | 4.487024787 | 0.041939368 | 0.388 | 0 |
| **828** | Slc50a1 | 1.69E-06 | 4.128347056 | 0.041939368 | 0.388 | 0 |
| **829** | Snrpe | 1.69E-06 | 4.500198931 | 0.041939368 | 0.388 | 0 |
| **830** | Tm2d1 | 1.69E-06 | 4.166727004 | 0.041939368 | 0.388 | 0 |
| **831** | Tmem134 | 1.69E-06 | 4.37818828 | 0.041939368 | 0.388 | 0 |
| **832** | Tppp3 | 1.69E-06 | 4.590320121 | 0.041939368 | 0.388 | 0 |
| **833** | Trappc3 | 1.69E-06 | 4.103433096 | 0.041939368 | 0.388 | 0 |
| **834** | Tubb4b | 1.69E-06 | 4.102893449 | 0.041939368 | 0.388 | 0 |
| **835** | Yipf6 | 1.69E-06 | 4.228618059 | 0.041939368 | 0.388 | 0 |
| **836** | Rab18 | 1.69E-06 | 1.114871176 | 0.04196914 | 0.653 | 0.102 |
| **837** | Timp3 | 1.69E-06 | 1.335262062 | 0.042000517 | 0.98 | 0.49 |
| **838** | Gimap6 | 1.69E-06 | 1.298523471 | 0.042115975 | 0.571 | 0.082 |
| **839** | 1700025G04R | 1.72E-06 | 0.826266594 | 0.042876656 | 0.776 | 0.184 |
| **840** | Sec24d | 1.74E-06 | 1.428990436 | 0.04320628 | 0.714 | 0.143 |
| **841** | Arhgdia | 1.75E-06 | 1.16776845 | 0.043620799 | 0.551 | 0.061 |
| **842** | Bach1 | 1.77E-06 | 1.381810884 | 0.043924627 | 0.592 | 0.082 |
| **843** | Aff3 | 1.79E-06 | 1.1019253 | 0.044544774 | 0.612 | 0.102 |
| **844** | Oxct1 | 1.79E-06 | 1.636635277 | 0.044544774 | 0.612 | 0.102 |
| **845** | Nr2f2 | 1.83E-06 | 1.007778261 | 0.04551609 | 0.653 | 0.102 |
| **846** | Mtch1 | 1.85E-06 | 1.328096504 | 0.04614175 | 0.633 | 0.102 |
| **847** | Tap2 | 1.89E-06 | 1.214368058 | 0.046955402 | 0.673 | 0.122 |
| **848** | Itih5 | 1.89E-06 | 1.194190269 | 0.047030938 | 0.837 | 0.224 |
| **849** | Nes | 1.91E-06 | 0.62862995 | 0.047560426 | 0.551 | 0.061 |
| **850** | Pon2 | 1.91E-06 | 1.195423081 | 0.047560426 | 0.551 | 0.061 |
| **851** | Mgst1 | 1.91E-06 | 1.128588232 | 0.047582599 | 0.755 | 0.163 |
| **852** | Mapk3 | 1.92E-06 | 0.918384842 | 0.047765427 | 0.592 | 0.082 |
| **853** | Arap2 | 1.93E-06 | 0.632035108 | 0.047938993 | 0.796 | 0.184 |
| **854** | Smarca5 | 1.93E-06 | 0.755837781 | 0.048126325 | 0.694 | 0.122 |

**Extended Table 2**

DEG from endothelial SnC (SEAC strategy) versus young

| | gene | mean_pval | mean_log2FC | p_val_adj | mean_pct_1 | mean_pct_2 |
|---|---|---|---|---|---|---|
| **1** | Rgs5 | 3.28E-33 | 7.508444755 | 8.16E-29 | 0.885 | 0.01 |
| **2** | Tmsb4x | 6.90E-30 | 4.241165294 | 1.72E-25 | 0.942 | 0.154 |
| **3** | Abcc9 | 2.15E-29 | 6.397679653 | 5.35E-25 | 0.817 | 0.01 |
| **4** | Pdgfrb | 1.14E-28 | 4.844779275 | 2.85E-24 | 0.837 | 0.029 |
| **5** | Inpp4b | 1.19E-28 | 3.507236129 | 2.97E-24 | 0.894 | 0.048 |
| **6** | Notch3 | 1.25E-28 | 5.266863029 | 3.10E-24 | 0.827 | 0.029 |
| **7** | Cst3 | 1.35E-28 | 4.146962775 | 3.35E-24 | 0.894 | 0.087 |
| **8** | Actb | 2.74E-28 | 3.316818923 | 6.81E-24 | 0.962 | 0.163 |
| **9** | Sgip1 | 1.57E-27 | 3.622448899 | 3.92E-23 | 0.913 | 0.144 |
| **10** | Adgrl3 | 1.59E-27 | 5.001825885 | 3.96E-23 | 0.817 | 0.038 |
| **11** | Rpl41 | 5.17E-27 | 3.716529146 | 1.29E-22 | 0.894 | 0.115 |
| **12** | Kcnq5 | 6.13E-27 | 7.287645432 | 1.52E-22 | 0.769 | 0.019 |
| **13** | Rps28 | 6.40E-27 | 3.793226529 | 1.59E-22 | 0.846 | 0.048 |
| **14** | Ptma | 6.44E-27 | 3.370931163 | 1.60E-22 | 0.894 | 0.087 |
| **15** | Rps16 | 1.01E-26 | 5.291823113 | 2.51E-22 | 0.788 | 0.029 |
| **16** | Fabp4 | 1.54E-26 | 3.051974459 | 3.82E-22 | 1 | 0.654 |
| **17** | Cacna1c | 1.55E-26 | 3.34970755 | 3.86E-22 | 0.904 | 0.135 |
| **18** | Rps24 | 2.51E-26 | 3.252641142 | 6.23E-22 | 0.875 | 0.106 |
| **19** | Rps20 | 3.11E-26 | 4.213811039 | 7.73E-22 | 0.817 | 0.048 |
| **20** | Stac | 3.96E-26 | 6.480125287 | 9.86E-22 | 0.75 | 0.01 |
| **21** | Rps3a1 | 4.43E-26 | 3.90218425 | 1.10E-21 | 0.817 | 0.048 |
| **22** | Postn | 1.21E-25 | 4.2429109 | 3.02E-21 | 0.827 | 0.077 |
| **23** | Rps21 | 1.89E-25 | 3.843154063 | 4.70E-21 | 0.837 | 0.077 |
| **24** | Rpl38 | 3.32E-25 | 3.355356353 | 8.26E-21 | 0.846 | 0.077 |
| **25** | Rpl37 | 6.88E-25 | 6.056401638 | 1.71E-20 | 0.731 | 0.019 |
| **26** | Rpl35a | 7.10E-25 | 3.187502608 | 1.77E-20 | 0.856 | 0.077 |
| **27** | Gpihbp1 | 1.12E-24 | 4.05238462 | 2.79E-20 | 0.798 | 0.048 |
| **28** | Eef1a1 | 1.16E-24 | 2.701404846 | 2.87E-20 | 0.952 | 0.202 |
| **29** | Rps29 | 1.37E-24 | 3.851248155 | 3.41E-20 | 0.817 | 0.067 |
| **30** | Gucy1a1 | 1.68E-24 | 3.700754863 | 4.17E-20 | 0.808 | 0.058 |
| **31** | Rps23 | 2.04E-24 | 3.554203994 | 5.08E-20 | 0.808 | 0.058 |
| **32** | Rps8 | 2.59E-24 | 3.441578332 | 6.43E-20 | 0.837 | 0.096 |
| **33** | Pla2r1 | 2.69E-24 | 2.831057816 | 6.69E-20 | 0.808 | 0.048 |
| **34** | Mfge8 | 3.27E-24 | 5.90484765 | 8.13E-20 | 0.712 | 0.01 |
| **35** | Col6a3 | 4.24E-24 | 2.867193003 | 1.06E-19 | 0.885 | 0.106 |
| **36** | B2m | 6.67E-24 | 2.499689748 | 1.66E-19 | 0.981 | 0.24 |
| **37** | Rpl13 | 7.84E-24 | 3.667455317 | 1.95E-19 | 0.798 | 0.058 |
| **38** | Steap4 | 1.42E-23 | 3.774327595 | 3.52E-19 | 0.779 | 0.058 |

| | | | | | | |
|---|---|---|---|---|---|---|
| **39** | Rpl27a | 1.85E-23 | 5.298218413 | 4.61E-19 | 0.712 | 0.019 |
| **40** | Rplp1 | 1.96E-23 | 2.794948896 | 4.86E-19 | 0.875 | 0.115 |
| **41** | Rpl18 | 4.18E-23 | 5.045344554 | 1.04E-18 | 0.692 | 0.01 |
| **42** | Sparc | 4.54E-23 | 2.277696736 | 1.13E-18 | 0.99 | 0.375 |
| **43** | Rpl37a | 5.01E-23 | 3.171925938 | 1.25E-18 | 0.837 | 0.106 |
| **44** | Gucy1a2 | 5.67E-23 | 3.865000674 | 1.41E-18 | 0.75 | 0.038 |
| **45** | Dlc1 | 6.18E-23 | 1.734826392 | 1.54E-18 | 1 | 0.75 |
| **46** | Sparcl1 | 6.45E-23 | 2.441665827 | 1.60E-18 | 0.962 | 0.288 |
| **47** | Ano1 | 8.88E-23 | 3.827470038 | 2.21E-18 | 0.76 | 0.058 |
| **48** | Calm1 | 9.82E-23 | 2.712328452 | 2.44E-18 | 0.885 | 0.125 |
| **49** | Rpl34 | 1.17E-22 | 4.462168499 | 2.92E-18 | 0.712 | 0.029 |
| **50** | Lin7a | 1.51E-22 | 8.190950681 | 3.75E-18 | 0.654 | 0 |
| **51** | Rps12 | 1.52E-22 | 3.229973358 | 3.78E-18 | 0.798 | 0.077 |
| **52** | Tpt1 | 1.91E-22 | 3.158606062 | 4.76E-18 | 0.827 | 0.106 |
| **53** | Actg1 | 2.21E-22 | 3.592585954 | 5.50E-18 | 0.788 | 0.087 |
| **54** | S100a6 | 2.89E-22 | 3.433066992 | 7.19E-18 | 0.779 | 0.067 |
| **55** | Ly6c1 | 3.34E-22 | 3.565200822 | 8.32E-18 | 0.75 | 0.058 |
| **56** | Agtr1a | 3.76E-22 | 4.375145798 | 9.35E-18 | 0.702 | 0.029 |
| **57** | Txnip | 4.07E-22 | 2.372230896 | 1.01E-17 | 0.971 | 0.346 |
| **58** | Aspn | 4.55E-22 | 5.649492007 | 1.13E-17 | 0.663 | 0.01 |
| **59** | Ghr | 7.33E-22 | -1.62708166 | 1.82E-17 | 0.99 | 0.981 |
| **60** | Ly6e | 7.34E-22 | 2.622078426 | 1.83E-17 | 0.885 | 0.144 |
| **61** | Cdh2 | 1.02E-21 | 7.99692797 | 2.54E-17 | 0.635 | 0 |
| **62** | Rps15 | 1.10E-21 | 3.315553386 | 2.73E-17 | 0.721 | 0.038 |
| **63** | Marcks | 1.20E-21 | 2.829542117 | 2.98E-17 | 0.837 | 0.115 |
| **64** | Hnrnpa3 | 2.14E-21 | 2.503058521 | 5.32E-17 | 0.817 | 0.096 |
| **65** | Rpl19 | 2.77E-21 | 3.470488279 | 6.89E-17 | 0.721 | 0.048 |
| **66** | Cyp4b1 | 2.95E-21 | 2.498637532 | 7.35E-17 | 0.846 | 0.144 |
| **67** | Eef2 | 3.20E-21 | 3.105950765 | 7.97E-17 | 0.75 | 0.058 |
| **68** | Ifitm3 | 3.35E-21 | 3.929186513 | 8.34E-17 | 0.731 | 0.058 |
| **69** | Rplp2 | 4.09E-21 | 2.69812263 | 1.02E-16 | 0.817 | 0.096 |
| **70** | Arhgef17 | 5.68E-21 | 4.405523337 | 1.41E-16 | 0.663 | 0.019 |
| **71** | Cfl1 | 6.22E-21 | 3.787201938 | 1.55E-16 | 0.683 | 0.029 |
| **72** | Gng11 | 9.48E-21 | 4.053589143 | 2.36E-16 | 0.683 | 0.038 |
| **73** | Ptms | 1.03E-20 | 3.09440353 | 2.58E-16 | 0.75 | 0.067 |
| **74** | Rhob | 1.34E-20 | 2.452258521 | 3.32E-16 | 0.788 | 0.077 |
| **75** | H3f3b | 1.75E-20 | 2.707724637 | 4.35E-16 | 0.808 | 0.106 |
| **76** | H2-D1 | 1.96E-20 | 2.145313937 | 4.87E-16 | 0.952 | 0.26 |
| **77** | Hsp90ab1 | 2.28E-20 | 2.609482019 | 5.66E-16 | 0.808 | 0.115 |
| **78** | Rpl28 | 2.60E-20 | 4.365224804 | 6.46E-16 | 0.663 | 0.029 |
| **79** | Rpsa | 2.94E-20 | 3.666568421 | 7.32E-16 | 0.683 | 0.038 |

| | | | | | | |
|---|---|---|---|---|---|---|
| **80** | Chn1 | 3.18E-20 | 4.311688079 | 7.92E-16 | 0.673 | 0.038 |
| **81** | Rps2 | 3.97E-20 | 3.265505214 | 9.87E-16 | 0.683 | 0.038 |
| **82** | Pde3a | 5.38E-20 | 1.521339706 | 1.34E-15 | 0.933 | 0.24 |
| **83** | Pcdh19 | 5.52E-20 | 2.541383397 | 1.37E-15 | 0.808 | 0.115 |
| **84** | Id3 | 5.53E-20 | 4.169225079 | 1.38E-15 | 0.654 | 0.029 |
| **85** | Rpl21 | 6.08E-20 | 3.28133633 | 1.51E-15 | 0.712 | 0.067 |
| **86** | Selenop | 6.09E-20 | 1.925443545 | 1.51E-15 | 0.875 | 0.135 |
| **87** | Rps4x | 6.87E-20 | 2.777893198 | 1.71E-15 | 0.721 | 0.058 |
| **88** | Cacna2d1 | 7.11E-20 | 2.300128605 | 1.77E-15 | 0.808 | 0.106 |
| **89** | Enox1 | 7.32E-20 | 2.54845217 | 1.82E-15 | 0.76 | 0.077 |
| **90** | Rps13 | 8.49E-20 | 3.981043688 | 2.11E-15 | 0.654 | 0.029 |
| **91** | Col5a3 | 1.01E-19 | 2.398600687 | 2.50E-15 | 0.817 | 0.115 |
| **92** | Rpl13a | 1.01E-19 | 2.230158761 | 2.51E-15 | 0.808 | 0.115 |
| **93** | Daam2 | 1.04E-19 | 4.020347636 | 2.58E-15 | 0.635 | 0.019 |
| **94** | Col6a2 | 1.14E-19 | 2.481571466 | 2.84E-15 | 0.721 | 0.058 |
| **95** | Myadm | 1.45E-19 | 2.04479304 | 3.61E-15 | 0.837 | 0.125 |
| **96** | Rpl6 | 1.70E-19 | 4.221392916 | 4.24E-15 | 0.644 | 0.029 |
| **97** | Tmem273 | 1.75E-19 | 4.9761405 | 4.35E-15 | 0.606 | 0.01 |
| **98** | Fau | 1.79E-19 | 3.210731254 | 4.46E-15 | 0.74 | 0.096 |
| **99** | Ednra | 1.79E-19 | 4.974010023 | 4.46E-15 | 0.606 | 0.01 |
| **100** | Rpl7 | 1.98E-19 | 4.351062063 | 4.92E-15 | 0.625 | 0.019 |
| **101** | Cd63 | 2.36E-19 | 2.890032262 | 5.88E-15 | 0.654 | 0.029 |
| **102** | Rpl30 | 2.52E-19 | 3.703332788 | 6.27E-15 | 0.673 | 0.048 |
| **103** | Rpl23 | 2.86E-19 | 2.5551323 | 7.11E-15 | 0.769 | 0.096 |
| **104** | Mrvi1 | 3.15E-19 | 2.272074085 | 7.85E-15 | 0.827 | 0.125 |
| **105** | Rps27a | 3.25E-19 | 2.926758613 | 8.09E-15 | 0.74 | 0.077 |
| **106** | Ly6a | 3.30E-19 | 3.114537293 | 8.20E-15 | 0.74 | 0.096 |
| **107** | Angptl4 | 3.60E-19 | 2.526968553 | 8.96E-15 | 0.74 | 0.067 |
| **108** | Hmgb1 | 3.99E-19 | 2.333947145 | 9.92E-15 | 0.798 | 0.115 |
| **109** | Rpl17 | 4.47E-19 | 2.832835645 | 1.11E-14 | 0.673 | 0.038 |
| **110** | Cd93 | 5.17E-19 | 2.146675557 | 1.29E-14 | 0.865 | 0.163 |
| **111** | Chd4 | 5.79E-19 | 2.188164221 | 1.44E-14 | 0.798 | 0.106 |
| **112** | Myh11 | 6.20E-19 | 4.024363216 | 1.54E-14 | 0.615 | 0.019 |
| **113** | Pakap | 8.62E-19 | 2.236782165 | 2.14E-14 | 0.731 | 0.067 |
| **114** | Tmsb10 | 8.69E-19 | 3.072695916 | 2.16E-14 | 0.663 | 0.038 |
| **115** | Rpl22 | 9.36E-19 | 4.585656623 | 2.33E-14 | 0.615 | 0.029 |
| **116** | Myo1b | 9.59E-19 | 1.981970728 | 2.39E-14 | 0.933 | 0.25 |
| **117** | Fth1 | 1.02E-18 | 2.468850709 | 2.53E-14 | 0.74 | 0.077 |
| **118** | Cox8a | 1.03E-18 | 4.563014343 | 2.56E-14 | 0.606 | 0.019 |
| **119** | Anxa1 | 1.07E-18 | 2.656359172 | 2.65E-14 | 0.702 | 0.058 |
| **120** | Eif1 | 1.34E-18 | 3.093932699 | 3.34E-14 | 0.721 | 0.096 |

| | | | | | | |
|---|---|---|---|---|---|---|
| **121** | Tbc1d1 | 1.69E-18 | 2.061652461 | 4.21E-14 | 0.779 | 0.106 |
| **122** | Ftl1 | 1.95E-18 | 2.443998165 | 4.86E-14 | 0.692 | 0.058 |
| **123** | Arhgap42 | 2.00E-18 | 2.301685314 | 4.99E-14 | 0.76 | 0.096 |
| **124** | Rps11 | 2.07E-18 | 4.430187585 | 5.16E-14 | 0.596 | 0.019 |
| **125** | Ctsl | 2.10E-18 | 2.674900504 | 5.22E-14 | 0.692 | 0.058 |
| **126** | Rpl39 | 2.13E-18 | 4.549332176 | 5.29E-14 | 0.596 | 0.019 |
| **127** | Ifitm2 | 2.47E-18 | 4.621015247 | 6.15E-14 | 0.596 | 0.019 |
| **128** | Col6a1 | 2.65E-18 | 2.432834631 | 6.60E-14 | 0.692 | 0.058 |
| **129** | Cxcl12 | 2.65E-18 | 2.322024978 | 6.60E-14 | 0.885 | 0.231 |
| **130** | Rps10 | 3.14E-18 | 3.697109506 | 7.81E-14 | 0.635 | 0.038 |
| **131** | Laptm4a | 3.38E-18 | 1.962631897 | 8.41E-14 | 0.894 | 0.212 |
| **132** | Rpl8 | 3.64E-18 | 6.984568988 | 9.05E-14 | 0.548 | 0 |
| **133** | Rpl18a | 3.94E-18 | 3.263370454 | 9.80E-14 | 0.654 | 0.048 |
| **134** | Gucy1b1 | 3.96E-18 | 3.698177868 | 9.86E-14 | 0.596 | 0.019 |
| **135** | Ubb | 4.26E-18 | 3.170159119 | 1.06E-13 | 0.663 | 0.058 |
| **136** | Cox7c | 4.41E-18 | 2.703976693 | 1.10E-13 | 0.663 | 0.048 |
| **137** | Serf2 | 4.63E-18 | 2.786081729 | 1.15E-13 | 0.663 | 0.048 |
| **138** | Plin2 | 6.32E-18 | 2.162493866 | 1.57E-13 | 0.779 | 0.106 |
| **139** | Kctd12 | 6.69E-18 | 2.810355382 | 1.67E-13 | 0.683 | 0.058 |
| **140** | Dmd | 7.18E-18 | 2.288427855 | 1.79E-13 | 0.904 | 0.308 |
| **141** | Rnf152 | 7.66E-18 | 3.277572987 | 1.90E-13 | 0.644 | 0.048 |
| **142** | Ubc | 9.19E-18 | 2.330032592 | 2.29E-13 | 0.731 | 0.096 |
| **143** | Jund | 9.30E-18 | 1.863305503 | 2.31E-13 | 0.692 | 0.058 |
| **144** | Rpl23a | 1.38E-17 | 3.199416337 | 3.43E-13 | 0.625 | 0.038 |
| **145** | Rpl32 | 1.39E-17 | 4.370964769 | 3.46E-13 | 0.577 | 0.019 |
| **146** | Ppia | 1.49E-17 | 2.782939203 | 3.71E-13 | 0.692 | 0.077 |
| **147** | Rpl27 | 1.52E-17 | 3.336054355 | 3.77E-13 | 0.625 | 0.038 |
| **148** | Cldn5 | 1.69E-17 | 1.929584959 | 4.20E-13 | 0.846 | 0.163 |
| **149** | Rflnb | 1.72E-17 | 2.054833441 | 4.27E-13 | 0.76 | 0.106 |
| **150** | Arpc3 | 1.84E-17 | 2.3714205 | 4.57E-13 | 0.635 | 0.038 |
| **151** | Bgn | 2.25E-17 | 3.388209278 | 5.61E-13 | 0.615 | 0.038 |
| **152** | Dbi | 2.46E-17 | 1.668237142 | 6.11E-13 | 0.75 | 0.087 |
| **153** | Igfbp7 | 2.52E-17 | 1.373113556 | 6.28E-13 | 0.981 | 0.317 |
| **154** | Srpr | 2.79E-17 | 5.084816822 | 6.95E-13 | 0.548 | 0.01 |
| **155** | Rpl11 | 2.96E-17 | 2.779972974 | 7.36E-13 | 0.654 | 0.058 |
| **156** | Cox6a1 | 3.03E-17 | 4.198814092 | 7.53E-13 | 0.567 | 0.019 |
| **157** | Itih5 | 3.11E-17 | 2.367420564 | 7.75E-13 | 0.779 | 0.144 |
| **158** | Car4 | 3.52E-17 | 1.694399438 | 8.76E-13 | 0.808 | 0.125 |
| **159** | Npc2 | 3.88E-17 | 2.987485163 | 9.66E-13 | 0.615 | 0.038 |
| **160** | Rpl4 | 3.94E-17 | 2.888814845 | 9.80E-13 | 0.625 | 0.048 |
| **161** | Cavin1 | 3.94E-17 | 1.736456931 | 9.81E-13 | 0.971 | 0.317 |

| | | | | | | |
|---|---|---|---|---|---|---|
| **162** | Ahnak | 4.14E-17 | 1.410055191 | 1.03E-12 | 1 | 0.587 |
| **163** | Angptl2 | 4.63E-17 | 2.368900445 | 1.15E-12 | 0.644 | 0.048 |
| **164** | Dock10 | 4.83E-17 | 1.894062065 | 1.20E-12 | 0.837 | 0.154 |
| **165** | Plcl1 | 5.31E-17 | 1.584036157 | 1.32E-12 | 0.933 | 0.269 |
| **166** | Rpl9 | 6.05E-17 | 2.628986277 | 1.51E-12 | 0.692 | 0.087 |
| **167** | Slfn5 | 7.50E-17 | 1.586432793 | 1.87E-12 | 0.981 | 0.365 |
| **168** | Pten | 7.67E-17 | 1.629617607 | 1.91E-12 | 0.99 | 0.385 |
| **169** | C1s1 | 8.10E-17 | 2.735332769 | 2.01E-12 | 0.635 | 0.048 |
| **170** | Ywhah | 9.96E-17 | 3.258761944 | 2.48E-12 | 0.596 | 0.038 |
| **171** | Rpl35 | 1.08E-16 | 2.345939018 | 2.68E-12 | 0.673 | 0.077 |
| **172** | Ddr2 | 1.23E-16 | 1.486548768 | 3.05E-12 | 0.817 | 0.144 |
| **173** | Rpl24 | 1.32E-16 | 3.402001669 | 3.29E-12 | 0.558 | 0.019 |
| **174** | Rpl10 | 1.37E-16 | 5.333343485 | 3.40E-12 | 0.529 | 0.01 |
| **175** | Atxn7l3b | 1.73E-16 | 3.53155884 | 4.29E-12 | 0.558 | 0.019 |
| **176** | Calm2 | 1.77E-16 | 2.457502705 | 4.41E-12 | 0.683 | 0.077 |
| **177** | Ywhaz | 1.92E-16 | 1.589861618 | 4.77E-12 | 0.894 | 0.212 |
| **178** | Psap | 1.94E-16 | 1.695526208 | 4.82E-12 | 0.962 | 0.356 |
| **179** | Ror1 | 1.97E-16 | 1.528711084 | 4.91E-12 | 0.808 | 0.144 |
| **180** | Fzd4 | 2.01E-16 | 2.025416364 | 5.00E-12 | 0.846 | 0.202 |
| **181** | Hspb1 | 2.07E-16 | 2.701032194 | 5.16E-12 | 0.635 | 0.058 |
| **182** | Synpo2 | 2.14E-16 | 2.357101222 | 5.32E-12 | 0.663 | 0.067 |
| **183** | Ighm | 2.27E-16 | 3.019370054 | 5.65E-12 | 0.577 | 0.029 |
| **184** | Cygb | 2.28E-16 | 3.713741949 | 5.66E-12 | 0.587 | 0.038 |
| **185** | S1pr1 | 2.33E-16 | 1.567641902 | 5.81E-12 | 0.875 | 0.202 |
| **186** | Lamp1 | 2.47E-16 | 2.442533434 | 6.15E-12 | 0.712 | 0.106 |
| **187** | Rpl7a | 2.48E-16 | 4.199885699 | 6.17E-12 | 0.558 | 0.029 |
| **188** | B230219D22R | 2.55E-16 | 2.491804698 | 6.34E-12 | 0.615 | 0.048 |
| **189** | Crip1 | 2.62E-16 | 3.980577368 | 6.53E-12 | 0.577 | 0.038 |
| **190** | Trpc3 | 3.09E-16 | 2.775907976 | 7.68E-12 | 0.567 | 0.029 |
| **191** | Hsp90b1 | 3.27E-16 | 1.824247917 | 8.13E-12 | 0.837 | 0.231 |
| **192** | Gsn | 3.86E-16 | 2.119185945 | 9.59E-12 | 0.952 | 0.394 |
| **193** | Rps9 | 3.95E-16 | 2.226116247 | 9.83E-12 | 0.721 | 0.106 |
| **194** | Akap12 | 3.96E-16 | 1.519474028 | 9.86E-12 | 0.769 | 0.125 |
| **195** | H2-T23 | 4.40E-16 | 2.774374847 | 1.09E-11 | 0.615 | 0.048 |
| **196** | Il34 | 4.44E-16 | 4.569951294 | 1.10E-11 | 0.519 | 0.01 |
| **197** | Pltp | 4.75E-16 | 1.647957855 | 1.18E-11 | 0.683 | 0.077 |
| **198** | Rpl26 | 4.85E-16 | 4.119266907 | 1.21E-11 | 0.538 | 0.019 |
| **199** | Crip2 | 5.04E-16 | 3.147958965 | 1.25E-11 | 0.577 | 0.038 |
| **200** | Ncl | 5.20E-16 | 2.130622403 | 1.29E-11 | 0.663 | 0.077 |
| **201** | Zfp36l1 | 5.32E-16 | 1.838308831 | 1.32E-11 | 0.875 | 0.231 |
| **202** | Rpl5 | 5.33E-16 | 2.21249327 | 1.33E-11 | 0.635 | 0.058 |

| | | | | | | |
|---|---|---|---|---|---|---|
| **203** | Rplp0 | 5.43E-16 | 2.491380487 | 1.35E-11 | 0.596 | 0.038 |
| **204** | H2-K1 | 5.77E-16 | 1.556270059 | 1.44E-11 | 0.962 | 0.356 |
| **205** | Ppfia2 | 5.87E-16 | 7.174271095 | 1.46E-11 | 0.49 | 0 |
| **206** | Gpx4 | 6.09E-16 | 2.253086349 | 1.51E-11 | 0.635 | 0.058 |
| **207** | Egflam | 6.52E-16 | 1.583245506 | 1.62E-11 | 0.596 | 0.038 |
| **208** | Eif4g2 | 6.70E-16 | 1.835762023 | 1.67E-11 | 0.817 | 0.163 |
| **209** | Rps7 | 6.82E-16 | 2.745582776 | 1.70E-11 | 0.644 | 0.077 |
| **210** | Rps27 | 7.61E-16 | 2.84858874 | 1.89E-11 | 0.644 | 0.077 |
| **211** | Itgb1 | 7.62E-16 | 1.515359035 | 1.90E-11 | 0.99 | 0.394 |
| **212** | Scd1 | 7.94E-16 | 1.805120408 | 1.98E-11 | 0.962 | 0.385 |
| **213** | Timp3 | 7.99E-16 | 1.509388886 | 1.99E-11 | 1 | 0.462 |
| **214** | Pros1 | 8.35E-16 | 1.182716062 | 2.08E-11 | 0.779 | 0.125 |
| **215** | Atp5e | 8.86E-16 | 3.385483254 | 2.21E-11 | 0.558 | 0.029 |
| **216** | C1qtnf9 | 8.88E-16 | 2.101973402 | 2.21E-11 | 0.654 | 0.067 |
| **217** | Tpm4 | 9.03E-16 | 1.694721071 | 2.25E-11 | 0.798 | 0.154 |
| **218** | Tuba1b | 9.57E-16 | 2.545872706 | 2.38E-11 | 0.644 | 0.067 |
| **219** | Ccdc85b | 1.03E-15 | 2.335595057 | 2.57E-11 | 0.654 | 0.067 |
| **220** | Ddx3x | 1.03E-15 | 1.528427709 | 2.57E-11 | 0.885 | 0.212 |
| **221** | Serping1 | 1.23E-15 | 3.588934849 | 3.05E-11 | 0.51 | 0.01 |
| **222** | Anxa5 | 1.24E-15 | 1.790378399 | 3.08E-11 | 0.74 | 0.115 |
| **223** | Rpl36a | 1.33E-15 | 4.112540967 | 3.30E-11 | 0.538 | 0.029 |
| **224** | Purb | 1.43E-15 | 2.323213779 | 3.55E-11 | 0.673 | 0.087 |
| **225** | Itm2b | 1.44E-15 | 1.515260293 | 3.59E-11 | 0.981 | 0.404 |
| **226** | Rps26 | 1.48E-15 | 2.54588502 | 3.67E-11 | 0.596 | 0.048 |
| **227** | Rpl15 | 1.69E-15 | 3.377647918 | 4.20E-11 | 0.529 | 0.019 |
| **228** | Dusp1 | 1.70E-15 | 2.893889329 | 4.24E-11 | 0.577 | 0.038 |
| **229** | Sox18 | 2.04E-15 | 3.568057717 | 5.08E-11 | 0.548 | 0.029 |
| **230** | Rps18 | 2.14E-15 | 3.416762261 | 5.32E-11 | 0.548 | 0.029 |
| **231** | Rpl3 | 2.19E-15 | 2.816989223 | 5.45E-11 | 0.577 | 0.038 |
| **232** | Rps3 | 2.35E-15 | 3.366292662 | 5.84E-11 | 0.548 | 0.029 |
| **233** | Myl6 | 2.51E-15 | 1.897605715 | 6.24E-11 | 0.865 | 0.24 |
| **234** | Tenm4 | 2.52E-15 | -2.15300187 | 6.27E-11 | 0.817 | 0.875 |
| **235** | Itpripl2 | 2.58E-15 | 1.993602402 | 6.42E-11 | 0.625 | 0.067 |
| **236** | Cox4i1 | 2.74E-15 | 2.341629194 | 6.82E-11 | 0.644 | 0.077 |
| **237** | P4hb | 3.17E-15 | 3.08430867 | 7.89E-11 | 0.548 | 0.029 |
| **238** | Sox5 | 4.07E-15 | 1.15487792 | 1.01E-10 | 0.971 | 0.365 |
| **239** | Dynll1 | 4.37E-15 | 2.65129245 | 1.09E-10 | 0.558 | 0.038 |
| **240** | Ifngr1 | 4.52E-15 | 1.740778542 | 1.13E-10 | 0.731 | 0.115 |
| **241** | Morrbid | 4.64E-15 | 1.784034714 | 1.15E-10 | 0.76 | 0.135 |
| **242** | Hspa8 | 4.76E-15 | 1.999591483 | 1.18E-10 | 0.721 | 0.125 |
| **243** | Chchd2 | 5.91E-15 | 4.121483934 | 1.47E-10 | 0.49 | 0.01 |

| | | | | | | |
|---|---|---|---|---|---|---|
| **244** | Rps19 | 6.08E-15 | 2.156095149 | 1.51E-10 | 0.692 | 0.106 |
| **245** | Rasl12 | 6.08E-15 | 4.229560683 | 1.51E-10 | 0.51 | 0.019 |
| **246** | Tnfrsf1a | 6.14E-15 | 2.289112132 | 1.53E-10 | 0.558 | 0.038 |
| **247** | Tcf15 | 6.43E-15 | 3.020438406 | 1.60E-10 | 0.558 | 0.038 |
| **248** | Gpc6 | 6.53E-15 | 1.053788572 | 1.62E-10 | 0.817 | 0.192 |
| **249** | Ctsb | 7.66E-15 | 1.583472839 | 1.91E-10 | 0.731 | 0.125 |
| **250** | Gjc1 | 7.67E-15 | 1.754442501 | 1.91E-10 | 0.673 | 0.096 |
| **251** | Adipoq | 8.34E-15 | 1.327539083 | 2.08E-10 | 0.721 | 0.106 |
| **252** | Ccdc47 | 8.65E-15 | 2.173650127 | 2.15E-10 | 0.625 | 0.067 |
| **253** | Rps25 | 9.27E-15 | 3.542432794 | 2.31E-10 | 0.51 | 0.019 |
| **254** | Spry4 | 9.31E-15 | 3.951532795 | 2.32E-10 | 0.519 | 0.029 |
| **255** | 4930523C07Ri | 9.48E-15 | 1.906304348 | 2.36E-10 | 0.721 | 0.125 |
| **256** | Cfd | 1.03E-14 | 1.581437678 | 2.57E-10 | 0.904 | 0.269 |
| **257** | Rps5 | 1.07E-14 | 2.336191934 | 2.67E-10 | 0.615 | 0.067 |
| **258** | Arhgap32 | 1.09E-14 | 1.660070118 | 2.70E-10 | 0.76 | 0.144 |
| **259** | Nfe2l1 | 1.25E-14 | 1.595594841 | 3.12E-10 | 0.808 | 0.163 |
| **260** | Aldoa | 1.32E-14 | 3.336728602 | 3.27E-10 | 0.5 | 0.019 |
| **261** | Ost4 | 1.38E-14 | 3.4377679 | 3.43E-10 | 0.481 | 0.01 |
| **262** | Arl6ip1 | 1.38E-14 | 2.972805609 | 3.44E-10 | 0.548 | 0.038 |
| **263** | Ybx1 | 1.47E-14 | 1.640994907 | 3.67E-10 | 0.76 | 0.135 |
| **264** | Sema5a | 1.59E-14 | 1.388040241 | 3.95E-10 | 0.827 | 0.183 |
| **265** | Eef1b2 | 1.65E-14 | 3.619794842 | 4.11E-10 | 0.519 | 0.029 |
| **266** | Oaz1 | 1.81E-14 | 2.511121507 | 4.49E-10 | 0.567 | 0.048 |
| **267** | Mxra7 | 2.00E-14 | 3.034105194 | 4.98E-10 | 0.529 | 0.029 |
| **268** | Map3k20 | 2.01E-14 | 1.552167133 | 5.00E-10 | 0.837 | 0.192 |
| **269** | Mamdc2 | 2.15E-14 | 3.320739051 | 5.34E-10 | 0.5 | 0.019 |
| **270** | Mylk | 2.15E-14 | 1.176170363 | 5.35E-10 | 0.769 | 0.144 |
| **271** | Gas5 | 2.22E-14 | 3.2812092 | 5.53E-10 | 0.519 | 0.029 |
| **272** | Tnxb | 2.26E-14 | 1.599283682 | 5.62E-10 | 0.769 | 0.144 |
| **273** | Adap2 | 2.29E-14 | 1.533096225 | 5.70E-10 | 0.692 | 0.106 |
| **274** | Znfx1 | 2.68E-14 | 1.445910299 | 6.67E-10 | 0.615 | 0.067 |
| **275** | Atp6v0b | 2.86E-14 | 2.656912425 | 7.12E-10 | 0.596 | 0.067 |
| **276** | Eif4a1 | 2.87E-14 | 2.223308996 | 7.14E-10 | 0.587 | 0.058 |
| **277** | Rack1 | 2.97E-14 | 4.006095858 | 7.40E-10 | 0.471 | 0.01 |
| **278** | Dag1 | 3.13E-14 | 1.142404506 | 7.79E-10 | 0.76 | 0.144 |
| **279** | Vstm4 | 3.30E-14 | 6.759959352 | 8.20E-10 | 0.442 | 0 |
| **280** | Tpm1 | 3.52E-14 | 1.873062099 | 8.76E-10 | 0.731 | 0.135 |
| **281** | Ssb | 3.77E-14 | 2.49274596 | 9.39E-10 | 0.606 | 0.067 |
| **282** | Carmn | 4.00E-14 | 2.554546125 | 9.95E-10 | 0.606 | 0.077 |
| **283** | Btg2 | 4.31E-14 | 2.154876217 | 1.07E-09 | 0.587 | 0.058 |
| **284** | Atp5h | 4.32E-14 | 2.024537556 | 1.07E-09 | 0.567 | 0.048 |

| | | | | | | |
|---|---|---|---|---|---|---|
| **285** | Npm1 | 4.46E-14 | 2.206835052 | 1.11E-09 | 0.558 | 0.048 |
| **286** | Tomm7 | 5.20E-14 | 3.427017922 | 1.29E-09 | 0.51 | 0.029 |
| **287** | Hspa5 | 5.22E-14 | 2.337120845 | 1.30E-09 | 0.654 | 0.106 |
| **288** | Prpf8 | 5.70E-14 | 2.714630346 | 1.42E-09 | 0.548 | 0.048 |
| **289** | Tpr | 6.12E-14 | 1.513532638 | 1.52E-09 | 0.769 | 0.154 |
| **290** | Cd81 | 6.29E-14 | 1.666459134 | 1.57E-09 | 0.817 | 0.202 |
| **291** | Rarres2 | 6.97E-14 | 3.000481116 | 1.73E-09 | 0.51 | 0.029 |
| **292** | Rpl36 | 7.48E-14 | 2.455463769 | 1.86E-09 | 0.577 | 0.067 |
| **293** | Vim | 7.79E-14 | 1.7330617 | 1.94E-09 | 0.817 | 0.192 |
| **294** | AW112010 | 9.20E-14 | 3.168755531 | 2.29E-09 | 0.5 | 0.029 |
| **295** | Rabac1 | 9.22E-14 | 2.545633737 | 2.29E-09 | 0.538 | 0.048 |
| **296** | Pfn1 | 9.31E-14 | 2.137995943 | 2.32E-09 | 0.596 | 0.067 |
| **297** | Btg1 | 1.04E-13 | 2.155454481 | 2.59E-09 | 0.51 | 0.029 |
| **298** | Uqcrh | 1.12E-13 | 2.132260748 | 2.79E-09 | 0.596 | 0.067 |
| **299** | Ets2 | 1.12E-13 | 1.623236421 | 2.79E-09 | 0.721 | 0.125 |
| **300** | Gcnt2 | 1.14E-13 | 1.56296791 | 2.85E-09 | 0.798 | 0.192 |
| **301** | Rps6 | 1.29E-13 | 2.438653431 | 3.22E-09 | 0.529 | 0.038 |
| **302** | Rps14 | 1.32E-13 | 1.649026086 | 3.29E-09 | 0.731 | 0.144 |
| **303** | 1810037I17Rik | 1.38E-13 | 2.773609016 | 3.43E-09 | 0.5 | 0.029 |
| **304** | Irf2bp2 | 1.53E-13 | 1.240042917 | 3.80E-09 | 0.663 | 0.106 |
| **305** | Rgs4 | 1.57E-13 | 6.506987699 | 3.90E-09 | 0.423 | 0 |
| **306** | Gnb2 | 1.61E-13 | 2.694725674 | 4.00E-09 | 0.519 | 0.038 |
| **307** | Tmem123 | 1.63E-13 | 1.412573177 | 4.06E-09 | 0.577 | 0.058 |
| **308** | Naca | 1.84E-13 | 2.028696791 | 4.57E-09 | 0.615 | 0.077 |
| **309** | Nus1 | 2.34E-13 | 0.928928595 | 5.82E-09 | 0.692 | 0.115 |
| **310** | Hprt | 2.38E-13 | 1.920441093 | 5.92E-09 | 0.538 | 0.048 |
| **311** | Ebf2 | 2.44E-13 | 0.757726628 | 6.08E-09 | 0.885 | 0.25 |
| **312** | Tmem30a | 2.47E-13 | 1.432341397 | 6.14E-09 | 0.76 | 0.154 |
| **313** | Bmpr1a | 2.58E-13 | 1.55815383 | 6.42E-09 | 0.692 | 0.125 |
| **314** | Zfand5 | 2.94E-13 | 1.444014929 | 7.31E-09 | 0.731 | 0.135 |
| **315** | Klf9 | 3.06E-13 | 1.376377284 | 7.60E-09 | 0.894 | 0.24 |
| **316** | Col1a2 | 3.07E-13 | 1.650909018 | 7.64E-09 | 0.654 | 0.106 |
| **317** | Atp5j2 | 3.14E-13 | 2.988627154 | 7.80E-09 | 0.442 | 0.01 |
| **318** | Ednrb | 3.72E-13 | 1.911023401 | 9.26E-09 | 0.577 | 0.067 |
| **319** | Tmem50a | 3.75E-13 | 1.932624458 | 9.33E-09 | 0.567 | 0.058 |
| **320** | Ndufc2 | 3.76E-13 | 1.39890883 | 9.35E-09 | 0.635 | 0.087 |
| **321** | Uqcr11 | 4.08E-13 | 2.47709419 | 1.02E-08 | 0.51 | 0.038 |
| **322** | Rps17 | 4.60E-13 | 2.30510562 | 1.15E-08 | 0.49 | 0.029 |
| **323** | Selenow | 4.76E-13 | 1.459299183 | 1.18E-08 | 0.712 | 0.135 |
| **324** | Atp5g3 | 4.83E-13 | 2.675587928 | 1.20E-08 | 0.519 | 0.048 |
| **325** | Scp2 | 4.86E-13 | 1.661836437 | 1.21E-08 | 0.577 | 0.067 |

| | | | | | | |
|---|---|---|---|---|---|---|
| **326** | Selenok | 5.07E-13 | 2.366573104 | 1.26E-08 | 0.51 | 0.038 |
| **327** | Mnd1 | 5.09E-13 | 4.645674848 | 1.27E-08 | 0.433 | 0.01 |
| **328** | Btf3 | 5.70E-13 | 2.098792523 | 1.42E-08 | 0.462 | 0.019 |
| **329** | Sult1a1 | 5.98E-13 | 1.720832262 | 1.49E-08 | 0.673 | 0.125 |
| **330** | Kank3 | 6.06E-13 | 1.168050628 | 1.51E-08 | 0.788 | 0.192 |
| **331** | Zfp91 | 6.31E-13 | 1.26666792 | 1.57E-08 | 0.654 | 0.106 |
| **332** | Tmbim6 | 7.10E-13 | 0.783094698 | 1.77E-08 | 0.846 | 0.202 |
| **333** | Sod1 | 7.15E-13 | 1.47624783 | 1.78E-08 | 0.51 | 0.038 |
| **334** | Cox6b1 | 7.23E-13 | 3.716514601 | 1.80E-08 | 0.452 | 0.019 |
| **335** | Cd38 | 8.04E-13 | 1.405991595 | 2.00E-08 | 0.644 | 0.106 |
| **336** | Ranbp2 | 8.27E-13 | 1.036490441 | 2.06E-08 | 0.827 | 0.183 |
| **337** | Prkar1a | 8.34E-13 | 1.129774241 | 2.08E-08 | 0.837 | 0.202 |
| **338** | Arpc5 | 8.77E-13 | 2.327500706 | 2.18E-08 | 0.481 | 0.029 |
| **339** | Nr1d1 | 8.79E-13 | 2.79549925 | 2.19E-08 | 0.471 | 0.029 |
| **340** | Sf3b2 | 9.18E-13 | 1.491440895 | 2.28E-08 | 0.587 | 0.077 |
| **341** | S100a11 | 9.19E-13 | 3.529752675 | 2.29E-08 | 0.471 | 0.029 |
| **342** | Hspa9 | 9.36E-13 | 2.59544094 | 2.33E-08 | 0.481 | 0.029 |
| **343** | M6pr | 9.39E-13 | 3.346432492 | 2.34E-08 | 0.471 | 0.029 |
| **344** | Selenof | 1.00E-12 | 1.505373501 | 2.49E-08 | 0.654 | 0.106 |
| **345** | Ccnt1 | 1.06E-12 | 1.965467998 | 2.63E-08 | 0.587 | 0.077 |
| **346** | Rad21 | 1.10E-12 | 1.271273066 | 2.73E-08 | 0.635 | 0.096 |
| **347** | Uqcrq | 1.11E-12 | 2.823448127 | 2.76E-08 | 0.452 | 0.019 |
| **348** | Sumo2 | 1.12E-12 | 1.977963184 | 2.77E-08 | 0.606 | 0.096 |
| **349** | Hnrnpk | 1.12E-12 | 1.505898272 | 2.80E-08 | 0.76 | 0.173 |
| **350** | Hsp90aa1 | 1.21E-12 | 2.93588292 | 3.02E-08 | 0.452 | 0.019 |
| **351** | Myl12a | 1.22E-12 | 1.788580472 | 3.04E-08 | 0.596 | 0.087 |
| **352** | Ddx24 | 1.32E-12 | 1.574689168 | 3.28E-08 | 0.625 | 0.096 |
| **353** | Rpl14 | 1.42E-12 | 3.771825318 | 3.54E-08 | 0.462 | 0.029 |
| **354** | Klf2 | 1.51E-12 | 1.40643393 | 3.76E-08 | 0.673 | 0.115 |
| **355** | Gipr | 1.54E-12 | 6.020222676 | 3.84E-08 | 0.394 | 0 |
| **356** | Pim3 | 1.56E-12 | 1.7576581 | 3.89E-08 | 0.538 | 0.058 |
| **357** | Ctla2a | 1.59E-12 | 2.493821455 | 3.96E-08 | 0.471 | 0.029 |
| **358** | Cnot6 | 1.60E-12 | 1.078453972 | 3.97E-08 | 0.625 | 0.087 |
| **359** | Dtx3 | 1.67E-12 | 1.800718302 | 4.15E-08 | 0.587 | 0.077 |
| **360** | Gbp7 | 1.81E-12 | 1.368200378 | 4.50E-08 | 0.577 | 0.077 |
| **361** | Srgn | 1.81E-12 | 1.891030235 | 4.51E-08 | 0.654 | 0.125 |
| **362** | Dusp3 | 1.92E-12 | 1.28298802 | 4.78E-08 | 0.76 | 0.163 |
| **363** | Arhgdia | 2.11E-12 | 2.134184238 | 5.24E-08 | 0.49 | 0.038 |
| **364** | Lhfp | 2.18E-12 | 1.247085003 | 5.42E-08 | 0.731 | 0.163 |
| **365** | Atp5l | 2.20E-12 | 2.284723929 | 5.48E-08 | 0.548 | 0.067 |
| **366** | Bzw1 | 2.30E-12 | 1.460832534 | 5.73E-08 | 0.567 | 0.067 |

| | | | | | | |
|---|---|---|---|---|---|---|
| **367** | Myh9 | 2.32E-12 | 1.056106836 | 5.76E-08 | 1 | 0.538 |
| **368** | Gstm1 | 2.39E-12 | 2.722451597 | 5.95E-08 | 0.5 | 0.048 |
| **369** | Cd300lg | 2.41E-12 | 0.912540474 | 6.00E-08 | 0.933 | 0.298 |
| **370** | Abi2 | 2.57E-12 | 1.685664859 | 6.39E-08 | 0.596 | 0.087 |
| **371** | Zfp36l2 | 2.64E-12 | 1.64603644 | 6.56E-08 | 0.577 | 0.077 |
| **372** | Eif3a | 2.73E-12 | 1.192138944 | 6.79E-08 | 0.712 | 0.144 |
| **373** | Psme1 | 3.03E-12 | 3.141310268 | 7.54E-08 | 0.413 | 0.01 |
| **374** | Tmed9 | 3.03E-12 | 3.420794858 | 7.54E-08 | 0.413 | 0.01 |
| **375** | Arhgap6 | 3.08E-12 | 0.837574939 | 7.66E-08 | 0.942 | 0.365 |
| **376** | Cspg4 | 3.27E-12 | 5.825305379 | 8.13E-08 | 0.385 | 0 |
| **377** | Myl9 | 3.27E-12 | 5.741073072 | 8.13E-08 | 0.385 | 0 |
| **378** | Rnasek | 3.27E-12 | 5.661702171 | 8.13E-08 | 0.385 | 0 |
| **379** | Tprgl | 3.27E-12 | 5.717028352 | 8.13E-08 | 0.385 | 0 |
| **380** | Tgoln1 | 3.36E-12 | 2.292784191 | 8.36E-08 | 0.529 | 0.058 |
| **381** | Cxcl9 | 3.43E-12 | 5.184507887 | 8.52E-08 | 0.404 | 0.01 |
| **382** | Pawr | 3.56E-12 | 0.579806645 | 8.86E-08 | 0.894 | 0.221 |
| **383** | Pdia3 | 3.61E-12 | 1.089392754 | 8.98E-08 | 0.798 | 0.192 |
| **384** | Slc12a2 | 3.70E-12 | 2.71924195 | 9.21E-08 | 0.481 | 0.038 |
| **385** | Tagln2 | 3.70E-12 | 1.800285362 | 9.21E-08 | 0.51 | 0.048 |
| **386** | Pnrc1 | 3.79E-12 | 1.693064405 | 9.44E-08 | 0.558 | 0.067 |
| **387** | Fam110b | 4.12E-12 | 1.905709396 | 1.02E-07 | 0.529 | 0.058 |
| **388** | Aoc3 | 4.19E-12 | 1.980152433 | 1.04E-07 | 0.51 | 0.048 |
| **389** | Ssr1 | 4.23E-12 | 1.382544523 | 1.05E-07 | 0.538 | 0.058 |
| **390** | Dusp10 | 4.41E-12 | 4.801975172 | 1.10E-07 | 0.404 | 0.01 |
| **391** | Mrfap1 | 4.61E-12 | 3.299584556 | 1.15E-07 | 0.433 | 0.019 |
| **392** | Itm2c | 5.15E-12 | 2.890244878 | 1.28E-07 | 0.433 | 0.019 |
| **393** | Sp1 | 5.16E-12 | 0.909122121 | 1.28E-07 | 0.673 | 0.115 |
| **394** | Rps15a | 5.29E-12 | 1.635282337 | 1.32E-07 | 0.625 | 0.106 |
| **395** | Lgals1 | 5.43E-12 | 4.271191694 | 1.35E-07 | 0.404 | 0.01 |
| **396** | Adamts12 | 5.45E-12 | 1.287959385 | 1.36E-07 | 0.933 | 0.423 |
| **397** | Retn | 5.74E-12 | 1.346884846 | 1.43E-07 | 0.615 | 0.106 |
| **398** | Nmt1 | 5.77E-12 | 2.210161112 | 1.43E-07 | 0.481 | 0.038 |
| **399** | Rpl22l1 | 5.81E-12 | 4.081518632 | 1.45E-07 | 0.404 | 0.01 |
| **400** | Pcbp1 | 6.11E-12 | 2.628019017 | 1.52E-07 | 0.452 | 0.029 |
| **401** | Hnrnph3 | 6.36E-12 | 1.050571332 | 1.58E-07 | 0.692 | 0.135 |
| **402** | Nr2f2 | 6.65E-12 | 1.161053499 | 1.65E-07 | 0.673 | 0.125 |
| **403** | Adamts15 | 6.86E-12 | 5.714805705 | 1.71E-07 | 0.375 | 0 |
| **404** | Atp1b2 | 6.86E-12 | 5.703548842 | 1.71E-07 | 0.375 | 0 |
| **405** | S100a1 | 6.86E-12 | 5.909161482 | 1.71E-07 | 0.375 | 0 |
| **406** | Klf6 | 6.88E-12 | 1.665981243 | 1.71E-07 | 0.702 | 0.163 |
| **407** | Snx18 | 7.78E-12 | 1.113977249 | 1.94E-07 | 0.606 | 0.096 |

| | | | | | | |
|---|---|---|---|---|---|---|
| **408** | Rpl12 | 8.06E-12 | 3.237499094 | 2.01E-07 | 0.423 | 0.019 |
| **409** | Thap3 | 8.29E-12 | 1.897748677 | 2.06E-07 | 0.49 | 0.048 |
| **410** | Hnrnpa0 | 8.95E-12 | 2.140148291 | 2.23E-07 | 0.529 | 0.067 |
| **411** | Sema4c | 9.00E-12 | 2.48279168 | 2.24E-07 | 0.452 | 0.029 |
| **412** | Gimap6 | 9.02E-12 | 1.348080478 | 2.24E-07 | 0.548 | 0.077 |
| **413** | Gnai2 | 9.11E-12 | 1.000304054 | 2.27E-07 | 0.817 | 0.202 |
| **414** | Psip1 | 9.20E-12 | 2.908376099 | 2.29E-07 | 0.423 | 0.019 |
| **415** | Hint1 | 9.41E-12 | 3.262614885 | 2.34E-07 | 0.423 | 0.019 |
| **416** | Apoe | 9.97E-12 | 1.633030908 | 2.48E-07 | 0.74 | 0.183 |
| **417** | Set | 1.06E-11 | 2.478048288 | 2.64E-07 | 0.462 | 0.038 |
| **418** | Ubxn1 | 1.10E-11 | 3.048373112 | 2.73E-07 | 0.423 | 0.019 |
| **419** | Tubb5 | 1.15E-11 | 2.606109379 | 2.85E-07 | 0.423 | 0.019 |
| **420** | Smc6 | 1.19E-11 | 0.988338675 | 2.95E-07 | 0.644 | 0.115 |
| **421** | S100a13 | 1.29E-11 | 2.135226744 | 3.22E-07 | 0.471 | 0.038 |
| **422** | Calm3 | 1.30E-11 | 3.553072559 | 3.23E-07 | 0.394 | 0.01 |
| **423** | Cnbp | 1.33E-11 | 0.975426223 | 3.31E-07 | 0.779 | 0.183 |
| **424** | Lamc3 | 1.43E-11 | 6.050534066 | 3.56E-07 | 0.365 | 0 |
| **425** | P3h2 | 1.49E-11 | 1.465447864 | 3.70E-07 | 0.606 | 0.115 |
| **426** | Ngf | 1.49E-11 | 1.91192595 | 3.72E-07 | 0.529 | 0.067 |
| **427** | Lman1 | 1.54E-11 | 2.447942221 | 3.84E-07 | 0.462 | 0.038 |
| **428** | Calr | 1.67E-11 | 2.412569422 | 4.15E-07 | 0.519 | 0.067 |
| **429** | Tram1 | 1.69E-11 | 1.810500096 | 4.19E-07 | 0.5 | 0.058 |
| **430** | Srsf3 | 1.69E-11 | 1.106238987 | 4.21E-07 | 0.75 | 0.173 |
| **431** | Aplp2 | 1.70E-11 | 1.054090634 | 4.24E-07 | 1 | 0.49 |
| **432** | Elob | 1.75E-11 | 3.457998076 | 4.35E-07 | 0.413 | 0.019 |
| **433** | H2afz | 1.75E-11 | 2.713216123 | 4.35E-07 | 0.413 | 0.019 |
| **434** | Nab1 | 1.84E-11 | 1.232835836 | 4.57E-07 | 0.615 | 0.106 |
| **435** | Hs2st1 | 1.87E-11 | 0.646473425 | 4.66E-07 | 0.702 | 0.144 |
| **436** | Vezf1 | 1.90E-11 | 1.747102808 | 4.72E-07 | 0.538 | 0.067 |
| **437** | Bst2 | 1.95E-11 | 2.317310121 | 4.85E-07 | 0.519 | 0.067 |
| **438** | Cds2 | 1.96E-11 | 0.487673635 | 4.87E-07 | 0.808 | 0.202 |
| **439** | Efnb1 | 1.96E-11 | 1.167632349 | 4.87E-07 | 0.548 | 0.077 |
| **440** | Gnb4 | 1.96E-11 | 1.16079243 | 4.88E-07 | 0.663 | 0.135 |
| **441** | Kdr | 2.00E-11 | 1.0840411 | 4.98E-07 | 0.894 | 0.317 |
| **442** | Nid1 | 2.03E-11 | 0.883099096 | 5.04E-07 | 0.933 | 0.317 |
| **443** | Mlf2 | 2.04E-11 | 3.168280589 | 5.07E-07 | 0.413 | 0.019 |
| **444** | Tmem176b | 2.11E-11 | 2.468606906 | 5.24E-07 | 0.433 | 0.029 |
| **445** | Sqstm1 | 2.13E-11 | 1.427375842 | 5.30E-07 | 0.625 | 0.115 |
| **446** | Ppp2r3a | 2.36E-11 | 1.201621742 | 5.87E-07 | 0.769 | 0.202 |
| **447** | Tmed5 | 2.39E-11 | 1.023227558 | 5.94E-07 | 0.663 | 0.135 |
| **448** | Col1a1 | 2.40E-11 | 1.477444068 | 5.97E-07 | 0.567 | 0.087 |

| | | | | | | |
|---|---|---|---|---|---|---|
| **449** | Vasp | 2.41E-11 | 1.322821915 | 6.01E-07 | 0.558 | 0.077 |
| **450** | Atp2a2 | 2.42E-11 | 0.987377785 | 6.02E-07 | 0.808 | 0.221 |
| **451** | Cox6c | 2.47E-11 | 1.703611846 | 6.16E-07 | 0.596 | 0.106 |
| **452** | Tinagl1 | 2.57E-11 | 1.768704914 | 6.40E-07 | 0.587 | 0.106 |
| **453** | Tm4sf1 | 2.60E-11 | 1.670962312 | 6.46E-07 | 0.692 | 0.173 |
| **454** | Lrp1 | 2.67E-11 | 1.095281402 | 6.64E-07 | 0.731 | 0.173 |
| **455** | Bzw2 | 2.74E-11 | 0.918816773 | 6.82E-07 | 0.596 | 0.096 |
| **456** | Ppp1r10 | 2.94E-11 | 2.14140084 | 7.30E-07 | 0.462 | 0.038 |
| **457** | Akr1a1 | 2.96E-11 | 1.564939886 | 7.36E-07 | 0.606 | 0.106 |
| **458** | Smdt1 | 2.97E-11 | 5.568214055 | 7.38E-07 | 0.356 | 0 |
| **459** | Tmf1 | 2.98E-11 | 1.573813269 | 7.41E-07 | 0.538 | 0.077 |
| **460** | Aff3 | 3.03E-11 | 1.153097514 | 7.54E-07 | 0.596 | 0.106 |
| **461** | Prkg1 | 3.07E-11 | 0.849714716 | 7.63E-07 | 0.952 | 0.365 |
| **462** | Bnip2 | 3.19E-11 | 1.168893263 | 7.95E-07 | 0.529 | 0.067 |
| **463** | Cpd | 3.33E-11 | 0.899142429 | 8.29E-07 | 0.827 | 0.221 |
| **464** | Atp5md | 3.43E-11 | 2.245669043 | 8.54E-07 | 0.433 | 0.029 |
| **465** | Dnaja1 | 3.51E-11 | 1.161874067 | 8.73E-07 | 0.731 | 0.173 |
| **466** | Canx | 3.58E-11 | 1.335376708 | 8.92E-07 | 0.817 | 0.24 |
| **467** | Nop53 | 3.64E-11 | 1.714352232 | 9.07E-07 | 0.452 | 0.038 |
| **468** | Ndufs5 | 3.66E-11 | 2.086884104 | 9.10E-07 | 0.433 | 0.029 |
| **469** | Ndufa2 | 3.73E-11 | 2.318390239 | 9.29E-07 | 0.433 | 0.029 |
| **470** | Cav2 | 3.76E-11 | 1.495607865 | 9.36E-07 | 0.712 | 0.173 |
| **471** | Eps8 | 3.89E-11 | 1.08346283 | 9.68E-07 | 0.894 | 0.308 |
| **472** | Naalad2 | 4.01E-11 | 3.230351942 | 9.99E-07 | 0.404 | 0.019 |
| **473** | Mmp14 | 4.03E-11 | 2.232417517 | 1.00E-06 | 0.471 | 0.048 |
| **474** | Parp14 | 4.11E-11 | 1.275307188 | 1.02E-06 | 0.76 | 0.192 |
| **475** | Eif2s2 | 4.20E-11 | 1.659195774 | 1.05E-06 | 0.481 | 0.048 |
| **476** | Serpinh1 | 4.35E-11 | 1.511019414 | 1.08E-06 | 0.654 | 0.144 |
| **477** | Cox7a2l | 4.40E-11 | 2.737525175 | 1.09E-06 | 0.423 | 0.029 |
| **478** | Mndal | 4.41E-11 | 1.153767313 | 1.10E-06 | 0.827 | 0.212 |
| **479** | Irf1 | 4.46E-11 | 1.209964293 | 1.11E-06 | 0.587 | 0.096 |
| **480** | Bnip3l | 4.59E-11 | 1.356702373 | 1.14E-06 | 0.519 | 0.067 |
| **481** | Uqcrb | 4.66E-11 | 2.193794454 | 1.16E-06 | 0.452 | 0.038 |
| **482** | Khsrp | 4.67E-11 | 2.395250416 | 1.16E-06 | 0.404 | 0.019 |
| **483** | Nes | 4.96E-11 | 0.827947818 | 1.23E-06 | 0.519 | 0.067 |
| **484** | Adnp | 5.00E-11 | 0.797425392 | 1.24E-06 | 0.635 | 0.115 |
| **485** | Rasgrp2 | 5.16E-11 | 0.767969016 | 1.28E-06 | 0.75 | 0.163 |
| **486** | Slc38a2 | 5.24E-11 | 1.368492955 | 1.30E-06 | 0.596 | 0.106 |
| **487** | Atp5b | 5.35E-11 | 1.832132216 | 1.33E-06 | 0.529 | 0.077 |
| **488** | Pde8b | 5.67E-11 | 3.548329982 | 1.41E-06 | 0.375 | 0.01 |
| **489** | Snrpg | 5.67E-11 | 3.448953005 | 1.41E-06 | 0.375 | 0.01 |

| | | | | | | |
|---|---|---|---|---|---|---|
| **490** | Gnas | 5.72E-11 | 0.992192596 | 1.42E-06 | 0.923 | 0.365 |
| **491** | Kcnj8 | 6.11E-11 | 6.45075592 | 1.52E-06 | 0.346 | 0 |
| **492** | Nol7 | 6.11E-11 | 5.591219238 | 1.52E-06 | 0.346 | 0 |
| **493** | Snrpe | 6.11E-11 | 5.619680828 | 1.52E-06 | 0.346 | 0 |
| **494** | Ubr2 | 6.11E-11 | 0.695583 | 1.52E-06 | 0.827 | 0.202 |
| **495** | Frmd4a | 6.27E-11 | 0.819326732 | 1.56E-06 | 0.856 | 0.25 |
| **496** | Sdf4 | 6.38E-11 | 1.126036551 | 1.59E-06 | 0.625 | 0.115 |
| **497** | Ddx42 | 6.45E-11 | 0.798037207 | 1.60E-06 | 0.606 | 0.106 |
| **498** | Sec24d | 6.57E-11 | 1.412940692 | 1.63E-06 | 0.654 | 0.135 |
| **499** | 7-Sep | 6.57E-11 | 1.063809862 | 1.64E-06 | 0.942 | 0.337 |
| **500** | Peli2 | 6.73E-11 | 1.034260377 | 1.67E-06 | 0.885 | 0.26 |
| **501** | Lhfpl3 | 6.75E-11 | 3.475619177 | 1.68E-06 | 0.394 | 0.019 |
| **502** | Atox1 | 6.80E-11 | 2.277924954 | 1.69E-06 | 0.442 | 0.038 |
| **503** | Ndrg2 | 6.85E-11 | 2.121173951 | 1.71E-06 | 0.423 | 0.029 |
| **504** | Tln1 | 6.98E-11 | 0.995334992 | 1.74E-06 | 0.74 | 0.173 |
| **505** | Elk4 | 7.11E-11 | 0.807020792 | 1.77E-06 | 0.846 | 0.231 |
| **506** | AC149090.1 | 7.19E-11 | 1.088531583 | 1.79E-06 | 0.663 | 0.144 |
| **507** | Naa50 | 7.27E-11 | 1.387676199 | 1.81E-06 | 0.519 | 0.067 |
| **508** | Ifi27l2a | 7.38E-11 | 1.140383485 | 1.84E-06 | 0.596 | 0.106 |
| **509** | Axl | 7.49E-11 | 2.342265473 | 1.86E-06 | 0.462 | 0.048 |
| **510** | Col3a1 | 7.57E-11 | 1.462503819 | 1.88E-06 | 0.827 | 0.269 |
| **511** | Kirrel | 7.74E-11 | 1.308366679 | 1.93E-06 | 0.692 | 0.163 |
| **512** | Rpl29 | 7.76E-11 | 4.866002743 | 1.93E-06 | 0.365 | 0.01 |
| **513** | Prrc2a | 7.93E-11 | 1.149935457 | 1.97E-06 | 0.635 | 0.125 |
| **514** | Chmp2a | 8.11E-11 | 4.365422946 | 2.02E-06 | 0.365 | 0.01 |
| **515** | Peg13 | 8.21E-11 | 2.70851016 | 2.04E-06 | 0.394 | 0.019 |
| **516** | Tcea1 | 8.32E-11 | 0.593736703 | 2.07E-06 | 0.673 | 0.135 |
| **517** | Fabp5 | 8.55E-11 | 2.446515519 | 2.13E-06 | 0.452 | 0.048 |
| **518** | Trarg1 | 8.61E-11 | 2.407858923 | 2.14E-06 | 0.462 | 0.048 |
| **519** | Brd2 | 9.15E-11 | 1.432446874 | 2.28E-06 | 0.596 | 0.106 |
| **520** | Ppib | 9.15E-11 | 1.62322245 | 2.28E-06 | 0.471 | 0.048 |
| **521** | Itga1 | 9.27E-11 | 0.861798258 | 2.31E-06 | 1 | 0.548 |
| **522** | Csnk1e | 9.62E-11 | 1.423339882 | 2.39E-06 | 0.49 | 0.058 |
| **523** | Camk2n1 | 9.71E-11 | 4.295779463 | 2.42E-06 | 0.365 | 0.01 |
| **524** | Rpn1 | 1.03E-10 | 2.247175373 | 2.56E-06 | 0.462 | 0.048 |
| **525** | Rsrc2 | 1.03E-10 | 0.485671055 | 2.57E-06 | 0.721 | 0.163 |
| **526** | Sdha | 1.05E-10 | 0.739589878 | 2.61E-06 | 0.615 | 0.106 |
| **527** | Nucks1 | 1.06E-10 | 1.301744096 | 2.63E-06 | 0.625 | 0.125 |
| **528** | Ccnd2 | 1.06E-10 | 1.752662254 | 2.64E-06 | 0.51 | 0.067 |
| **529** | Tbx18 | 1.09E-10 | 3.555035532 | 2.70E-06 | 0.365 | 0.01 |
| **530** | Pip5k1b | 1.16E-10 | 0.834623571 | 2.88E-06 | 0.529 | 0.077 |

| | | | | | | |
|---|---|---|---|---|---|---|
| **531** | Atp6ap2 | 1.16E-10 | 3.02105294 | 2.89E-06 | 0.365 | 0.01 |
| **532** | Gapdh | 1.16E-10 | 3.641820867 | 2.89E-06 | 0.365 | 0.01 |
| **533** | Rem1 | 1.16E-10 | 2.359972036 | 2.89E-06 | 0.365 | 0.01 |
| **534** | Arcn1 | 1.19E-10 | 0.953024573 | 2.95E-06 | 0.635 | 0.125 |
| **535** | Prrx1 | 1.22E-10 | 0.89693806 | 3.02E-06 | 0.644 | 0.135 |
| **536** | Jph1 | 1.26E-10 | 5.215942724 | 3.14E-06 | 0.356 | 0.01 |
| **537** | Tmem182 | 1.29E-10 | 0.900431875 | 3.20E-06 | 0.702 | 0.154 |
| **538** | Meox2 | 1.31E-10 | 0.75395777 | 3.26E-06 | 0.798 | 0.202 |
| **539** | Sfr1 | 1.32E-10 | 2.41770803 | 3.27E-06 | 0.385 | 0.019 |
| **540** | Hnrnpul2 | 1.34E-10 | 1.091489876 | 3.34E-06 | 0.673 | 0.144 |
| **541** | Cycs | 1.35E-10 | 4.827079421 | 3.36E-06 | 0.356 | 0.01 |
| **542** | Atp5f1 | 1.40E-10 | 1.204675842 | 3.49E-06 | 0.596 | 0.106 |
| **543** | Lrrc58 | 1.41E-10 | 0.988827912 | 3.50E-06 | 0.837 | 0.24 |
| **544** | Srsf1 | 1.41E-10 | 0.88266594 | 3.51E-06 | 0.548 | 0.087 |
| **545** | Pear1 | 1.42E-10 | 1.132260142 | 3.53E-06 | 0.644 | 0.135 |
| **546** | Ifi203 | 1.42E-10 | 0.873625372 | 3.54E-06 | 0.808 | 0.221 |
| **547** | Extl3 | 1.47E-10 | 0.464136014 | 3.66E-06 | 0.731 | 0.163 |
| **548** | Atp5a1 | 1.48E-10 | 0.960220019 | 3.67E-06 | 0.567 | 0.096 |
| **549** | 6720427I07Rik | 1.53E-10 | 0.909683337 | 3.82E-06 | 0.596 | 0.106 |
| **550** | Atp5j | 1.58E-10 | 1.920906356 | 3.93E-06 | 0.51 | 0.077 |
| **551** | Papss2 | 1.60E-10 | 0.607270632 | 3.98E-06 | 0.702 | 0.154 |
| **552** | Cd151 | 1.64E-10 | 2.126114114 | 4.09E-06 | 0.433 | 0.038 |
| **553** | Piezo2 | 1.65E-10 | 1.271680017 | 4.10E-06 | 0.519 | 0.077 |
| **554** | Pdia4 | 1.68E-10 | 1.841197389 | 4.19E-06 | 0.452 | 0.048 |
| **555** | Sem1 | 1.69E-10 | 1.121591235 | 4.20E-06 | 0.702 | 0.173 |
| **556** | C1qtnf1 | 1.69E-10 | 0.921326371 | 4.20E-06 | 0.462 | 0.048 |
| **557** | Ucp2 | 1.78E-10 | 2.00437749 | 4.43E-06 | 0.433 | 0.038 |
| **558** | Rtn4rl1 | 1.83E-10 | 1.639692779 | 4.56E-06 | 0.481 | 0.058 |
| **559** | Dok6 | 1.84E-10 | 1.555950725 | 4.58E-06 | 0.548 | 0.096 |
| **560** | Emc7 | 1.86E-10 | 1.534766082 | 4.63E-06 | 0.5 | 0.067 |
| **561** | Cox17 | 1.97E-10 | 1.932188579 | 4.90E-06 | 0.433 | 0.038 |
| **562** | Atp5mpl | 2.02E-10 | 1.989880712 | 5.03E-06 | 0.385 | 0.019 |
| **563** | Zbtb16 | 2.08E-10 | -1.7231881 | 5.18E-06 | 0.933 | 0.846 |
| **564** | Glyr1 | 2.09E-10 | 0.863641541 | 5.19E-06 | 0.625 | 0.125 |
| **565** | Lbh | 2.18E-10 | 1.550459243 | 5.42E-06 | 0.452 | 0.048 |
| **566** | Runx1t1 | 2.25E-10 | 1.206431781 | 5.61E-06 | 0.615 | 0.125 |
| **567** | Pfdn5 | 2.26E-10 | 3.730896788 | 5.62E-06 | 0.356 | 0.01 |
| **568** | Txn1 | 2.27E-10 | 1.301804471 | 5.65E-06 | 0.433 | 0.038 |
| **569** | Rnf10 | 2.27E-10 | 1.716503742 | 5.65E-06 | 0.529 | 0.087 |
| **570** | Srsf6 | 2.29E-10 | 0.79488822 | 5.70E-06 | 0.577 | 0.096 |
| **571** | Eif3f | 2.31E-10 | 3.626085719 | 5.74E-06 | 0.356 | 0.01 |

| | | | | | | |
|---|---|---|---|---|---|---|
| **572** | Psmb10 | 2.36E-10 | 2.407737874 | 5.87E-06 | 0.356 | 0.01 |
| **573** | Atp5g2 | 2.41E-10 | 1.964451212 | 6.00E-06 | 0.471 | 0.058 |
| **574** | Cyp2d22 | 2.59E-10 | 1.090440693 | 6.44E-06 | 0.481 | 0.058 |
| **575** | Depp1 | 2.71E-10 | 1.979018276 | 6.74E-06 | 0.471 | 0.058 |
| **576** | Fmo2 | 2.73E-10 | 1.161193856 | 6.80E-06 | 0.558 | 0.096 |
| **577** | Ndufb1-ps | 2.84E-10 | 2.953720158 | 7.06E-06 | 0.375 | 0.019 |
| **578** | Rgcc | 2.99E-10 | 1.262272696 | 7.43E-06 | 0.683 | 0.183 |
| **579** | H2afv | 3.10E-10 | 2.322831098 | 7.70E-06 | 0.404 | 0.029 |
| **580** | Rock1 | 3.15E-10 | 0.896505161 | 7.84E-06 | 0.981 | 0.404 |
| **581** | Tap2 | 3.16E-10 | 1.135190499 | 7.87E-06 | 0.625 | 0.135 |
| **582** | Trip10 | 3.22E-10 | 2.814413604 | 8.02E-06 | 0.375 | 0.019 |
| **583** | Rnpc3 | 3.33E-10 | 0.387997936 | 8.28E-06 | 0.5 | 0.067 |
| **584** | Atf4 | 3.47E-10 | 1.677513338 | 8.64E-06 | 0.471 | 0.058 |
| **585** | Fcgrt | 3.48E-10 | 1.215956879 | 8.66E-06 | 0.587 | 0.106 |
| **586** | Pgrmc1 | 3.48E-10 | 1.318419522 | 8.67E-06 | 0.452 | 0.048 |
| **587** | Cbx3 | 3.50E-10 | 1.268897918 | 8.71E-06 | 0.587 | 0.115 |
| **588** | B130024G19R | 3.54E-10 | 2.326410114 | 8.81E-06 | 0.442 | 0.048 |
| **589** | Timp4 | 3.56E-10 | 1.186402876 | 8.86E-06 | 0.625 | 0.135 |
| **590** | Ndufb9 | 3.59E-10 | 2.286806767 | 8.93E-06 | 0.375 | 0.019 |
| **591** | Ywhag | 3.64E-10 | 0.892457093 | 9.06E-06 | 0.74 | 0.192 |
| **592** | Daam1 | 3.77E-10 | 1.065632051 | 9.38E-06 | 0.837 | 0.298 |
| **593** | Atp5k | 3.77E-10 | 1.644089011 | 9.39E-06 | 0.423 | 0.038 |
| **594** | Slc24a3 | 3.90E-10 | 1.226678782 | 9.72E-06 | 0.837 | 0.308 |
| **595** | Ik | 3.93E-10 | 2.833939403 | 9.78E-06 | 0.394 | 0.029 |
| **596** | Lpar6 | 3.99E-10 | 0.921314108 | 9.93E-06 | 0.596 | 0.115 |
| **597** | Acat1 | 4.05E-10 | 1.521575253 | 1.01E-05 | 0.5 | 0.077 |
| **598** | Sppl2a | 4.05E-10 | 0.761625852 | 1.01E-05 | 0.798 | 0.231 |
| **599** | H1f0 | 4.08E-10 | 1.757757175 | 1.01E-05 | 0.375 | 0.019 |
| **600** | Rpl31 | 4.13E-10 | 3.413256684 | 1.03E-05 | 0.365 | 0.019 |
| **601** | Col5a1 | 4.28E-10 | 0.882992551 | 1.07E-05 | 0.644 | 0.135 |
| **602** | Tspan15 | 4.30E-10 | 0.647281472 | 1.07E-05 | 0.606 | 0.115 |
| **603** | Ubtf | 4.34E-10 | 1.471380184 | 1.08E-05 | 0.481 | 0.067 |
| **604** | Tmed2 | 4.68E-10 | 1.998112815 | 1.16E-05 | 0.481 | 0.067 |
| **605** | Gabarapl1 | 4.77E-10 | 2.933660446 | 1.19E-05 | 0.346 | 0.01 |
| **606** | Dek | 4.95E-10 | 1.763649044 | 1.23E-05 | 0.481 | 0.067 |
| **607** | Snrnp200 | 4.98E-10 | 1.02310058 | 1.24E-05 | 0.471 | 0.058 |
| **608** | Rab12 | 5.07E-10 | 0.772052771 | 1.26E-05 | 0.577 | 0.106 |
| **609** | Btg3 | 5.10E-10 | 1.027031368 | 1.27E-05 | 0.567 | 0.106 |
| **610** | Robo1 | 5.11E-10 | 1.410815373 | 1.27E-05 | 0.548 | 0.096 |
| **611** | Gpx8 | 5.13E-10 | 5.944199788 | 1.28E-05 | 0.317 | 0 |
| **612** | Ndufa1 | 5.13E-10 | 5.495476914 | 1.28E-05 | 0.317 | 0 |

| | | | | | | |
|---|---|---|---|---|---|---|
| **613** | Aqp1 | 5.13E-10 | 1.720683577 | 1.28E-05 | 0.644 | 0.183 |
| **614** | Arl6ip5 | 5.20E-10 | 0.911897215 | 1.29E-05 | 0.606 | 0.115 |
| **615** | Taf1 | 5.33E-10 | 0.923132447 | 1.33E-05 | 0.625 | 0.125 |
| **616** | Hnrnpa2b1 | 5.52E-10 | 1.112034532 | 1.37E-05 | 0.913 | 0.356 |
| **617** | Rdx | 5.60E-10 | 0.830009014 | 1.39E-05 | 0.76 | 0.202 |
| **618** | Rab18 | 5.75E-10 | 1.22591275 | 1.43E-05 | 0.558 | 0.096 |
| **619** | Arap2 | 5.75E-10 | 0.77842213 | 1.43E-05 | 0.779 | 0.221 |
| **620** | Sox7 | 5.99E-10 | 1.555103338 | 1.49E-05 | 0.442 | 0.048 |
| **621** | Hrct1 | 6.00E-10 | 1.62689996 | 1.49E-05 | 0.413 | 0.038 |
| **622** | Ndufa3 | 6.24E-10 | 2.691713867 | 1.55E-05 | 0.385 | 0.029 |
| **623** | Capns1 | 6.83E-10 | 1.711324356 | 1.70E-05 | 0.433 | 0.048 |
| **624** | Tmed7 | 6.99E-10 | 1.465193478 | 1.74E-05 | 0.442 | 0.048 |
| **625** | Pard3 | 7.14E-10 | 0.550602953 | 1.78E-05 | 0.846 | 0.269 |
| **626** | Tmem109 | 7.24E-10 | 1.974570653 | 1.80E-05 | 0.433 | 0.048 |
| **627** | Klhl29 | 7.38E-10 | 1.982675167 | 1.84E-05 | 0.433 | 0.048 |
| **628** | Plekha2 | 7.48E-10 | 0.660726213 | 1.86E-05 | 0.692 | 0.154 |
| **629** | Slc3a2 | 7.63E-10 | 1.44753596 | 1.90E-05 | 0.452 | 0.058 |
| **630** | Cdk11b | 7.65E-10 | 0.978737706 | 1.90E-05 | 0.558 | 0.106 |
| **631** | Bcl2 | 7.72E-10 | 0.709290176 | 1.92E-05 | 0.692 | 0.192 |
| **632** | Cdh5 | 7.94E-10 | 1.077034632 | 1.98E-05 | 0.962 | 0.481 |
| **633** | Polr2a | 7.94E-10 | 1.072381363 | 1.98E-05 | 0.596 | 0.125 |
| **634** | Pcmtd2 | 8.05E-10 | 0.588570442 | 2.00E-05 | 0.596 | 0.115 |
| **635** | Kcnq4 | 8.14E-10 | 1.993632479 | 2.02E-05 | 0.433 | 0.048 |
| **636** | Msl1 | 8.17E-10 | 1.040727735 | 2.03E-05 | 0.481 | 0.067 |
| **637** | Gpx1 | 8.18E-10 | 1.771402667 | 2.04E-05 | 0.519 | 0.087 |
| **638** | Grn | 8.23E-10 | 1.813687002 | 2.05E-05 | 0.452 | 0.058 |
| **639** | Ryk | 8.36E-10 | 1.152685764 | 2.08E-05 | 0.596 | 0.125 |
| **640** | Os9 | 8.42E-10 | 1.868969045 | 2.09E-05 | 0.413 | 0.038 |
| **641** | Ehd2 | 8.55E-10 | 0.831080182 | 2.13E-05 | 0.913 | 0.327 |
| **642** | Klhl9 | 8.67E-10 | 2.537535329 | 2.16E-05 | 0.385 | 0.029 |
| **643** | Ppp4r2 | 8.82E-10 | 0.477663706 | 2.19E-05 | 0.731 | 0.173 |
| **644** | H3f3a | 9.09E-10 | 1.061143504 | 2.26E-05 | 0.683 | 0.173 |
| **645** | Gng5 | 9.14E-10 | 2.689123835 | 2.27E-05 | 0.404 | 0.038 |
| **646** | Ubl5 | 9.16E-10 | 3.827248854 | 2.28E-05 | 0.337 | 0.01 |
| **647** | Trim2 | 9.20E-10 | 1.020048276 | 2.29E-05 | 0.635 | 0.154 |
| **648** | Ddit4 | 9.22E-10 | 1.589749186 | 2.29E-05 | 0.471 | 0.067 |
| **649** | Rbbp7 | 9.47E-10 | 1.195035294 | 2.36E-05 | 0.481 | 0.067 |
| **650** | Ndufa5 | 9.57E-10 | 3.252202189 | 2.38E-05 | 0.337 | 0.01 |
| **651** | Prr13 | 9.57E-10 | 2.16797744 | 2.38E-05 | 0.337 | 0.01 |
| **652** | Sgk1 | 9.68E-10 | 1.663077614 | 2.41E-05 | 0.433 | 0.048 |
| **653** | Spry1 | 9.71E-10 | 1.161723299 | 2.42E-05 | 0.462 | 0.058 |

| | | | | | | |
|---|---|---|---|---|---|---|
| **654** | Ifi35 | 9.87E-10 | 1.806048633 | 2.46E-05 | 0.433 | 0.048 |
| **655** | Rnf44 | 9.89E-10 | 1.186527694 | 2.46E-05 | 0.462 | 0.058 |
| **656** | Rbms3 | 9.90E-10 | 0.819367503 | 2.46E-05 | 0.933 | 0.404 |
| **657** | Uvssa | 1.01E-09 | 2.107926678 | 2.51E-05 | 0.404 | 0.038 |
| **658** | Trim56 | 1.02E-09 | 1.070451796 | 2.54E-05 | 0.529 | 0.096 |
| **659** | Mcam | 1.03E-09 | 1.560462647 | 2.56E-05 | 0.51 | 0.087 |
| **660** | Ccdc82 | 1.05E-09 | 0.548758445 | 2.61E-05 | 0.692 | 0.173 |
| **661** | Cct6a | 1.09E-09 | 1.420717011 | 2.72E-05 | 0.452 | 0.058 |
| **662** | Tomm6 | 1.11E-09 | 2.206756066 | 2.76E-05 | 0.385 | 0.029 |
| **663** | Bicc1 | 1.12E-09 | 0.727775452 | 2.78E-05 | 0.644 | 0.154 |
| **664** | Enpep | 1.12E-09 | 1.050127787 | 2.79E-05 | 0.615 | 0.135 |
| **665** | Zbtb7a | 1.12E-09 | 1.219257385 | 2.80E-05 | 0.548 | 0.096 |
| **666** | Ecm2 | 1.15E-09 | 1.174863948 | 2.85E-05 | 0.519 | 0.087 |
| **667** | G3bp1 | 1.18E-09 | 0.680358343 | 2.95E-05 | 0.721 | 0.173 |
| **668** | Chmp4b | 1.19E-09 | 0.482333216 | 2.97E-05 | 0.635 | 0.135 |
| **669** | Apool | 1.20E-09 | 1.857546918 | 2.99E-05 | 0.385 | 0.029 |
| **670** | Plod2 | 1.20E-09 | 1.378771566 | 2.99E-05 | 0.548 | 0.106 |
| **671** | Cald1 | 1.22E-09 | 0.792900307 | 3.03E-05 | 0.952 | 0.423 |
| **672** | Frg1 | 1.25E-09 | 1.765728026 | 3.11E-05 | 0.385 | 0.029 |
| **673** | Angpt2 | 1.26E-09 | 2.280308131 | 3.12E-05 | 0.404 | 0.038 |
| **674** | Sf1 | 1.35E-09 | 0.798002628 | 3.37E-05 | 0.673 | 0.163 |
| **675** | Kctd12b | 1.39E-09 | 1.173090752 | 3.47E-05 | 0.433 | 0.048 |
| **676** | Tmod3 | 1.40E-09 | 0.80243881 | 3.48E-05 | 0.913 | 0.308 |
| **677** | Igf2r | 1.42E-09 | 0.764410088 | 3.52E-05 | 0.798 | 0.231 |
| **678** | Srsf2 | 1.47E-09 | 1.239827981 | 3.66E-05 | 0.558 | 0.106 |
| **679** | Pgrmc2 | 1.50E-09 | 1.391505019 | 3.72E-05 | 0.51 | 0.087 |
| **680** | Tmem59 | 1.53E-09 | 1.345378563 | 3.81E-05 | 0.548 | 0.106 |
| **681** | Cavin2 | 1.56E-09 | 1.015560607 | 3.88E-05 | 0.933 | 0.404 |
| **682** | Ttc19 | 1.56E-09 | 1.420177912 | 3.88E-05 | 0.452 | 0.058 |
| **683** | Spcs2 | 1.56E-09 | 2.253317731 | 3.88E-05 | 0.404 | 0.038 |
| **684** | Cox5b | 1.60E-09 | 4.055168998 | 3.99E-05 | 0.327 | 0.01 |
| **685** | Ubr1 | 1.63E-09 | 0.832523333 | 4.04E-05 | 0.817 | 0.25 |
| **686** | Ran | 1.64E-09 | 3.983187298 | 4.08E-05 | 0.327 | 0.01 |
| **687** | Calu | 1.64E-09 | 0.779415243 | 4.08E-05 | 0.644 | 0.144 |
| **688** | Tmed10 | 1.66E-09 | 0.822433144 | 4.13E-05 | 0.692 | 0.173 |
| **689** | Atp9a | 1.68E-09 | 0.451164672 | 4.17E-05 | 0.75 | 0.192 |
| **690** | Jun | 1.69E-09 | 1.265941583 | 4.21E-05 | 0.519 | 0.096 |
| **691** | Serinc3 | 1.73E-09 | 0.868825668 | 4.31E-05 | 0.981 | 0.423 |
| **692** | Impad1 | 1.78E-09 | 1.309014699 | 4.42E-05 | 0.452 | 0.058 |
| **693** | Eif4h | 1.82E-09 | 0.815626773 | 4.52E-05 | 0.548 | 0.106 |
| **694** | Plau | 1.87E-09 | 3.418337323 | 4.65E-05 | 0.327 | 0.01 |

| | | | | | | |
|---|---|---|---|---|---|---|
| **695** | Romo1 | 1.87E-09 | 3.320232857 | 4.65E-05 | 0.327 | 0.01 |
| **696** | Larp4 | 1.87E-09 | 0.919450639 | 4.65E-05 | 0.538 | 0.096 |
| **697** | Arf6 | 1.91E-09 | 2.923955654 | 4.75E-05 | 0.327 | 0.01 |
| **698** | Itpripl1 | 1.91E-09 | 2.495000093 | 4.75E-05 | 0.327 | 0.01 |
| **699** | Mapk3 | 1.92E-09 | 0.883505473 | 4.78E-05 | 0.471 | 0.067 |
| **700** | Nbeal1 | 1.95E-09 | 0.877877196 | 4.86E-05 | 0.981 | 0.462 |
| **701** | Tubgcp5 | 1.96E-09 | 1.522564165 | 4.87E-05 | 0.423 | 0.048 |
| **702** | Dock7 | 1.96E-09 | 0.801460227 | 4.88E-05 | 0.712 | 0.192 |
| **703** | Pttg1ip | 1.97E-09 | 0.57004414 | 4.90E-05 | 0.625 | 0.135 |
| **704** | Rbm8a | 2.02E-09 | 2.408563672 | 5.01E-05 | 0.375 | 0.029 |
| **705** | Rsad2 | 2.05E-09 | 1.483932785 | 5.11E-05 | 0.5 | 0.087 |
| **706** | Dgkb | 2.06E-09 | 7.225936213 | 5.12E-05 | 0.298 | 0 |
| **707** | Sh3bgrl3 | 2.06E-09 | 5.100060391 | 5.12E-05 | 0.298 | 0 |
| **708** | Vtn | 2.06E-09 | 5.606087872 | 5.12E-05 | 0.298 | 0 |
| **709** | Nfkbia | 2.09E-09 | 1.197298999 | 5.20E-05 | 0.51 | 0.087 |
| **710** | Il31ra | 2.11E-09 | 1.321016012 | 5.25E-05 | 0.952 | 0.462 |
| **711** | Hnrnpab | 2.16E-09 | 1.361070607 | 5.38E-05 | 0.462 | 0.067 |
| **712** | Khdrbs1 | 2.16E-09 | 0.81670972 | 5.38E-05 | 0.75 | 0.202 |
| **713** | Sclt1 | 2.22E-09 | 1.983421232 | 5.51E-05 | 0.452 | 0.067 |
| **714** | Aldh2 | 2.22E-09 | 0.856581269 | 5.52E-05 | 0.538 | 0.096 |
| **715** | Eif4e3 | 2.23E-09 | 1.144443279 | 5.55E-05 | 0.375 | 0.029 |
| **716** | Tspo | 2.23E-09 | 2.747532964 | 5.55E-05 | 0.375 | 0.029 |
| **717** | Cul2 | 2.23E-09 | 1.62892866 | 5.55E-05 | 0.442 | 0.058 |
| **718** | Srp14 | 2.28E-09 | 2.115992155 | 5.66E-05 | 0.375 | 0.029 |
| **719** | Nr1h3 | 2.29E-09 | 1.584317278 | 5.69E-05 | 0.462 | 0.067 |
| **720** | Nfic | 2.36E-09 | 1.04503133 | 5.88E-05 | 0.51 | 0.087 |
| **721** | Mcm9 | 2.38E-09 | 0.284970178 | 5.91E-05 | 0.49 | 0.077 |
| **722** | Slc16a2 | 2.39E-09 | 0.76456778 | 5.96E-05 | 0.538 | 0.106 |
| **723** | Cyp20a1 | 2.39E-09 | 1.691209854 | 5.96E-05 | 0.49 | 0.087 |
| **724** | Rbx1 | 2.40E-09 | 1.396017915 | 5.98E-05 | 0.442 | 0.058 |
| **725** | Olfm2 | 2.45E-09 | 1.856217428 | 6.09E-05 | 0.413 | 0.048 |
| **726** | Zhx1 | 2.47E-09 | 1.51966016 | 6.14E-05 | 0.375 | 0.029 |
| **727** | Sbno1 | 2.49E-09 | 0.321609731 | 6.20E-05 | 0.683 | 0.163 |
| **728** | Trf | 2.57E-09 | 1.5387748 | 6.38E-05 | 0.529 | 0.106 |
| **729** | Lamb2 | 2.59E-09 | 1.316320927 | 6.44E-05 | 0.481 | 0.077 |
| **730** | Bsg | 2.61E-09 | 1.059924326 | 6.50E-05 | 0.587 | 0.125 |
| **731** | Ushbp1 | 2.67E-09 | 0.700850529 | 6.64E-05 | 0.856 | 0.279 |
| **732** | Eif5a | 2.78E-09 | 2.801343161 | 6.90E-05 | 0.346 | 0.019 |
| **733** | Tfcp2 | 2.84E-09 | 1.51366514 | 7.06E-05 | 0.442 | 0.058 |
| **734** | Myl12b | 2.84E-09 | 0.7843665 | 7.06E-05 | 0.683 | 0.173 |
| **735** | Chst1 | 2.84E-09 | 1.44128196 | 7.07E-05 | 0.462 | 0.067 |

| | | | | | | |
|---|---|---|---|---|---|---|
| **736** | Ramp2 | 2.86E-09 | 1.829561307 | 7.12E-05 | 0.548 | 0.125 |
| **737** | Fgfr1op2 | 2.88E-09 | 0.55463365 | 7.16E-05 | 0.644 | 0.144 |
| **738** | Mat2a | 2.94E-09 | 0.754904655 | 7.31E-05 | 0.673 | 0.163 |
| **739** | Rhoa | 2.97E-09 | 0.977761781 | 7.39E-05 | 0.865 | 0.308 |
| **740** | Kpnb1 | 3.04E-09 | 0.858108814 | 7.57E-05 | 0.558 | 0.106 |
| **741** | Sub1 | 3.06E-09 | 1.282065629 | 7.60E-05 | 0.442 | 0.058 |
| **742** | Myo7a | 3.08E-09 | 2.129555921 | 7.66E-05 | 0.346 | 0.019 |
| **743** | Gas1 | 3.10E-09 | 1.730243663 | 7.72E-05 | 0.394 | 0.038 |
| **744** | Cdk16 | 3.14E-09 | 1.943730758 | 7.82E-05 | 0.346 | 0.019 |
| **745** | Abcb7 | 3.18E-09 | 1.057483509 | 7.92E-05 | 0.587 | 0.125 |
| **746** | Cdkn1a | 3.21E-09 | 2.006946382 | 7.99E-05 | 0.346 | 0.019 |
| **747** | Tle5 | 3.21E-09 | 2.189117939 | 7.99E-05 | 0.346 | 0.019 |
| **748** | Cdkn1b | 3.26E-09 | 1.563571568 | 8.10E-05 | 0.413 | 0.048 |
| **749** | Fscn1 | 3.26E-09 | 2.05648477 | 8.10E-05 | 0.413 | 0.048 |
| **750** | Ndufa7 | 3.26E-09 | 1.869368322 | 8.12E-05 | 0.442 | 0.067 |
| **751** | Mmrn2 | 3.32E-09 | 0.771319436 | 8.25E-05 | 0.721 | 0.192 |
| **752** | Mxd4 | 3.36E-09 | 0.559484671 | 8.36E-05 | 0.712 | 0.183 |
| **753** | Slf2 | 3.40E-09 | 0.696675071 | 8.45E-05 | 0.587 | 0.125 |
| **754** | Rad23b | 3.42E-09 | 0.8459653 | 8.51E-05 | 0.683 | 0.173 |
| **755** | Ghitm | 3.42E-09 | 1.82293706 | 8.52E-05 | 0.471 | 0.077 |
| **756** | Eif4g1 | 3.43E-09 | 0.898482766 | 8.53E-05 | 0.692 | 0.183 |
| **757** | Capn7 | 3.56E-09 | 1.465398966 | 8.86E-05 | 0.452 | 0.067 |
| **758** | S100a16 | 3.56E-09 | 1.457325939 | 8.87E-05 | 0.481 | 0.077 |
| **759** | Ypel3 | 3.72E-09 | 1.657726516 | 9.25E-05 | 0.413 | 0.048 |
| **760** | Lamb1 | 3.74E-09 | 0.676084971 | 9.31E-05 | 0.712 | 0.183 |
| **761** | Cyb5r3 | 3.78E-09 | 2.057153476 | 9.42E-05 | 0.317 | 0.01 |
| **762** | Kctd17 | 3.78E-09 | 2.975726778 | 9.42E-05 | 0.317 | 0.01 |
| **763** | Med28 | 3.78E-09 | 2.425323352 | 9.42E-05 | 0.317 | 0.01 |
| **764** | Pcyox1 | 3.78E-09 | 2.418692266 | 9.42E-05 | 0.317 | 0.01 |
| **765** | Ptpdc1 | 3.78E-09 | 2.691248914 | 9.42E-05 | 0.317 | 0.01 |
| **766** | Uba1 | 3.84E-09 | 1.518484449 | 9.56E-05 | 0.394 | 0.038 |
| **767** | Anp32e | 3.87E-09 | 2.520695741 | 9.64E-05 | 0.365 | 0.029 |
| **768** | Dhx40 | 3.97E-09 | 0.919338551 | 9.89E-05 | 0.51 | 0.087 |
| **769** | mt-Nd4 | 4.05E-09 | 0.81057963 | 0.000100783 | 1 | 0.75 |
| **770** | Dpm3 | 4.08E-09 | 5.337308588 | 0.000101506 | 0.288 | 0 |
| **771** | Tpm2 | 4.08E-09 | 5.54306909 | 0.000101506 | 0.288 | 0 |
| **772** | Ccnt2 | 4.09E-09 | 0.551180367 | 0.000101754 | 0.769 | 0.221 |
| **773** | Ddrgk1 | 4.20E-09 | 1.071676553 | 0.000104396 | 0.433 | 0.058 |
| **774** | Cox5a | 4.40E-09 | 1.518830957 | 0.000109595 | 0.413 | 0.048 |
| **775** | Arpc4 | 4.49E-09 | 1.466622919 | 0.000111676 | 0.413 | 0.048 |
| **776** | Ube2b | 4.50E-09 | 0.753567583 | 0.00011199 | 0.76 | 0.221 |

| | | | | | | |
|---|---|---|---|---|---|---|
| **777** | Ddx23 | 4.52E-09 | 1.390666845 | 0.000112373 | 0.433 | 0.058 |
| **778** | S100a10 | 4.55E-09 | 1.398729789 | 0.00011312 | 0.471 | 0.077 |
| **779** | Lclat1 | 4.56E-09 | 1.281632547 | 0.000113498 | 0.481 | 0.077 |
| **780** | Rab31 | 4.68E-09 | 0.557164235 | 0.00011654 | 0.615 | 0.135 |
| **781** | Iws1 | 4.80E-09 | 0.775303775 | 0.000119521 | 0.529 | 0.096 |
| **782** | Mprip | 4.93E-09 | 0.791681389 | 0.000122675 | 0.971 | 0.433 |
| **783** | Syde1 | 4.97E-09 | 0.631451461 | 0.000123595 | 0.51 | 0.087 |
| **784** | Ech1 | 5.06E-09 | 2.367114619 | 0.000125837 | 0.385 | 0.038 |
| **785** | Flna | 5.08E-09 | 1.387776512 | 0.000126463 | 0.615 | 0.154 |
| **786** | Cct8 | 5.16E-09 | 1.915500972 | 0.0001283 | 0.385 | 0.038 |
| **787** | Samd4 | 5.24E-09 | 0.374252212 | 0.000130343 | 0.798 | 0.24 |
| **788** | Ube2z | 5.33E-09 | 0.984287242 | 0.000132551 | 0.433 | 0.058 |
| **789** | Grcc10 | 5.35E-09 | 2.855211658 | 0.00013301 | 0.337 | 0.019 |
| **790** | Cfl2 | 5.41E-09 | 1.260673873 | 0.000134727 | 0.413 | 0.048 |
| **791** | Eif3k | 5.46E-09 | 3.027870071 | 0.000135795 | 0.337 | 0.019 |
| **792** | Per1 | 5.52E-09 | 0.709567079 | 0.000137272 | 0.413 | 0.048 |
| **793** | Rnf7 | 5.57E-09 | 2.774299034 | 0.000138638 | 0.337 | 0.019 |
| **794** | mt-Co1 | 5.69E-09 | 0.712956707 | 0.000141461 | 1 | 0.942 |
| **795** | Ecm1 | 5.82E-09 | 2.807483157 | 0.000144821 | 0.356 | 0.029 |
| **796** | Pdia6 | 5.84E-09 | 0.892713245 | 0.000145209 | 0.538 | 0.106 |
| **797** | Scoc | 5.93E-09 | 1.949360255 | 0.000147517 | 0.337 | 0.019 |
| **798** | Stimate | 6.02E-09 | 0.466531299 | 0.000149703 | 0.567 | 0.115 |
| **799** | Psma1 | 6.05E-09 | 2.570926521 | 0.000150597 | 0.337 | 0.019 |
| **800** | Arpc1a | 6.17E-09 | 1.357018998 | 0.000153415 | 0.433 | 0.058 |
| **801** | Trim35 | 6.22E-09 | 1.265019157 | 0.000154749 | 0.452 | 0.067 |
| **802** | Rhoj | 6.23E-09 | 0.8415403 | 0.000154963 | 0.971 | 0.5 |
| **803** | Nhs | 6.25E-09 | 1.143252799 | 0.000155549 | 0.51 | 0.096 |
| **804** | Atpif1 | 6.31E-09 | 0.930352572 | 0.000156948 | 0.337 | 0.019 |
| **805** | Lsm12 | 6.31E-09 | 2.101954649 | 0.000156948 | 0.337 | 0.019 |
| **806** | 2-Mar | 6.37E-09 | 0.981394042 | 0.000158489 | 0.596 | 0.135 |
| **807** | Ndufb10 | 6.55E-09 | 3.941617327 | 0.000163012 | 0.308 | 0.01 |
| **808** | Syf2 | 6.56E-09 | 1.747505101 | 0.000163299 | 0.356 | 0.029 |
| **809** | Sp3 | 6.63E-09 | 0.703832618 | 0.000164949 | 0.683 | 0.183 |
| **810** | Atp6v1b2 | 6.68E-09 | 0.777952537 | 0.000166195 | 0.452 | 0.067 |
| **811** | Sec62 | 6.75E-09 | 0.964989532 | 0.000167889 | 0.615 | 0.154 |
| **812** | Stk38l | 6.78E-09 | 0.698969937 | 0.000168581 | 0.538 | 0.106 |
| **813** | Zfp664 | 6.80E-09 | 1.047773846 | 0.000169268 | 0.51 | 0.096 |
| **814** | Tspan13 | 6.91E-09 | 0.698611247 | 0.000171984 | 0.827 | 0.269 |
| **815** | Nvl | 7.05E-09 | 1.094001981 | 0.000175315 | 0.452 | 0.067 |
| **816** | BC031181 | 7.14E-09 | 3.640386236 | 0.000177713 | 0.308 | 0.01 |
| **817** | Cbx1 | 7.22E-09 | 0.920347799 | 0.000179731 | 0.577 | 0.125 |

| | | | | | | |
|---|---|---|---|---|---|---|
| **818** | Fap | 7.23E-09 | 0.755160019 | 0.000180007 | 0.596 | 0.135 |
| **819** | Slc30a1 | 7.30E-09 | 3.549716537 | 0.000181585 | 0.308 | 0.01 |
| **820** | Tspyl1 | 7.30E-09 | 3.367472734 | 0.000181585 | 0.308 | 0.01 |
| **821** | Thbs2 | 7.44E-09 | 1.116087903 | 0.000185056 | 0.385 | 0.038 |
| **822** | Tuba1a | 7.46E-09 | 3.305770562 | 0.00018554 | 0.308 | 0.01 |
| **823** | Ndufa4 | 7.53E-09 | 1.527579415 | 0.000187331 | 0.462 | 0.077 |
| **824** | Eda | 7.66E-09 | 0.837400611 | 0.000190472 | 0.596 | 0.154 |
| **825** | Nek9 | 7.69E-09 | 0.895005911 | 0.000191279 | 0.548 | 0.115 |
| **826** | Mlec | 7.85E-09 | 2.04926782 | 0.000195385 | 0.356 | 0.029 |
| **827** | Arl4a | 8.05E-09 | 5.280818326 | 0.000200197 | 0.279 | 0 |
| **828** | Des | 8.05E-09 | 5.402162283 | 0.000200197 | 0.279 | 0 |
| **829** | Gstm2 | 8.05E-09 | 5.2927768 | 0.000200197 | 0.279 | 0 |
| **830** | Hes1 | 8.05E-09 | 5.170271096 | 0.000200197 | 0.279 | 0 |
| **831** | Hic1 | 8.05E-09 | 5.433261165 | 0.000200197 | 0.279 | 0 |
| **832** | Imp3 | 8.05E-09 | 5.307843515 | 0.000200197 | 0.279 | 0 |
| **833** | Pcdh18 | 8.05E-09 | 5.545761527 | 0.000200197 | 0.279 | 0 |
| **834** | Psmb6 | 8.05E-09 | 5.155762 | 0.000200197 | 0.279 | 0 |
| **835** | Runx2 | 8.05E-09 | 5.303436932 | 0.000200197 | 0.279 | 0 |
| **836** | Selenom | 8.05E-09 | 5.30756172 | 0.000200197 | 0.279 | 0 |
| **837** | Igfbp5 | 8.08E-09 | 1.38683077 | 0.00020099 | 0.548 | 0.125 |
| **838** | Arl5a | 8.12E-09 | 1.271029215 | 0.000202081 | 0.452 | 0.067 |
| **839** | Pik3r1 | 8.38E-09 | 0.997148642 | 0.000208614 | 0.952 | 0.462 |
| **840** | Rnase4 | 8.50E-09 | 3.457566326 | 0.000211516 | 0.327 | 0.019 |
| **841** | Tmem256 | 8.50E-09 | 1.589408397 | 0.000211541 | 0.356 | 0.029 |
| **842** | Cggbp1 | 8.59E-09 | 1.451934677 | 0.000213731 | 0.423 | 0.058 |
| **843** | Elmo1 | 8.65E-09 | -1.54442626 | 0.000215228 | 0.942 | 0.808 |
| **844** | Psma7 | 8.85E-09 | 1.667339456 | 0.000220098 | 0.356 | 0.029 |
| **845** | Fam76a | 8.93E-09 | 1.180979188 | 0.000222243 | 0.404 | 0.048 |
| **846** | Arf4 | 8.94E-09 | 0.719608019 | 0.0002224 | 0.615 | 0.154 |
| **847** | Pdap1 | 9.02E-09 | 2.354524089 | 0.000224502 | 0.356 | 0.029 |
| **848** | Cobll1 | 9.31E-09 | 0.452670166 | 0.000231518 | 0.846 | 0.288 |
| **849** | Ssr3 | 9.52E-09 | 1.06978911 | 0.000236895 | 0.51 | 0.096 |
| **850** | Eif5b | 9.55E-09 | 0.91566397 | 0.000237581 | 0.587 | 0.135 |
| **851** | Gabarap | 9.57E-09 | 2.205191742 | 0.000238153 | 0.375 | 0.038 |
| **852** | Dusp6 | 9.62E-09 | 1.47721584 | 0.000239398 | 0.404 | 0.048 |
| **853** | Limd1 | 9.69E-09 | 0.892814892 | 0.000241079 | 0.625 | 0.154 |
| **854** | Eif5 | 9.69E-09 | 0.740043424 | 0.000241168 | 0.452 | 0.067 |
| **855** | Eif4a2 | 9.69E-09 | 0.852791506 | 0.000241171 | 0.74 | 0.212 |
| **856** | Pola2 | 9.99E-09 | 0.941863704 | 0.000248452 | 0.404 | 0.048 |
| **857** | Hspa4 | 1.01E-08 | 0.439718175 | 0.0002525 | 0.615 | 0.144 |
| **858** | Dstn | 1.02E-08 | 1.048074597 | 0.000253683 | 0.538 | 0.115 |

| | | | | | | |
|---|---|---|---|---|---|---|
| **859** | Stam | 1.05E-08 | 0.897894491 | 0.000260479 | 0.615 | 0.144 |
| **860** | 4-Sep | 1.05E-08 | 0.87542119 | 0.000262467 | 0.625 | 0.154 |
| **861** | Antxr1 | 1.06E-08 | 0.617189531 | 0.000264941 | 0.529 | 0.106 |
| **862** | Llph | 1.09E-08 | 2.834509723 | 0.000270928 | 0.327 | 0.019 |
| **863** | Elp4 | 1.11E-08 | 0.481865209 | 0.000275525 | 0.683 | 0.183 |
| **864** | Cavin3 | 1.11E-08 | 2.420434381 | 0.000276553 | 0.327 | 0.019 |
| **865** | Ube2q1 | 1.12E-08 | 1.677592189 | 0.000277643 | 0.375 | 0.038 |
| **866** | Sdc3 | 1.14E-08 | 0.350580092 | 0.000282831 | 0.625 | 0.144 |
| **867** | Arhgap10 | 1.14E-08 | 0.693874463 | 0.000283089 | 0.894 | 0.327 |
| **868** | Hnrnpd | 1.14E-08 | 0.305716364 | 0.000284834 | 0.779 | 0.212 |
| **869** | Fam162a | 1.15E-08 | 0.491003628 | 0.000286705 | 0.394 | 0.048 |
| **870** | Mier1 | 1.18E-08 | 0.86801881 | 0.000292457 | 0.817 | 0.269 |
| **871** | Skp1a | 1.19E-08 | 1.081968916 | 0.000295387 | 0.5 | 0.096 |
| **872** | Agpat5 | 1.19E-08 | 0.863592208 | 0.000296151 | 0.423 | 0.058 |
| **873** | Jcad | 1.19E-08 | 0.433639221 | 0.000296618 | 0.817 | 0.25 |
| **874** | Bcam | 1.20E-08 | 1.27970864 | 0.000299479 | 0.51 | 0.106 |
| **875** | Afg3l1 | 1.23E-08 | 1.80746485 | 0.000306429 | 0.327 | 0.019 |
| **876** | Tril | 1.23E-08 | 1.616349488 | 0.000306429 | 0.327 | 0.019 |
| **877** | Copz1 | 1.23E-08 | 1.124850936 | 0.000307024 | 0.423 | 0.058 |
| **878** | Anxa7 | 1.23E-08 | 0.885440184 | 0.000307047 | 0.529 | 0.106 |
| **879** | Ddx21 | 1.25E-08 | 1.202655696 | 0.000311366 | 0.375 | 0.038 |
| **880** | Drap1 | 1.26E-08 | 4.023259865 | 0.000312788 | 0.298 | 0.01 |
| **881** | Uqcr10 | 1.26E-08 | 3.848801434 | 0.000312788 | 0.298 | 0.01 |
| **882** | Thbd | 1.26E-08 | 0.988066677 | 0.000313299 | 0.538 | 0.115 |
| **883** | Synm | 1.26E-08 | 1.25897632 | 0.000313427 | 0.413 | 0.058 |
| **884** | Btbd1 | 1.27E-08 | 0.5444526 | 0.000315673 | 0.683 | 0.173 |
| **885** | Smtn | 1.29E-08 | 1.157612031 | 0.000321103 | 0.5 | 0.096 |
| **886** | Arf5 | 1.30E-08 | 1.857171192 | 0.000323466 | 0.375 | 0.038 |
| **887** | Klf4 | 1.31E-08 | 1.847244813 | 0.000326154 | 0.433 | 0.067 |
| **888** | Ifnar2 | 1.35E-08 | 0.911504308 | 0.000335405 | 0.654 | 0.173 |
| **889** | Cdc5l | 1.37E-08 | 0.19791149 | 0.000341067 | 0.538 | 0.106 |
| **890** | Fnbp4 | 1.37E-08 | 0.704245741 | 0.000342062 | 0.606 | 0.144 |
| **891** | Slc25a3 | 1.37E-08 | 0.614416088 | 0.000342062 | 0.606 | 0.144 |
| **892** | Rac1 | 1.39E-08 | 0.863860063 | 0.000346192 | 0.769 | 0.24 |
| **893** | Dctn4 | 1.39E-08 | 1.225913788 | 0.000346359 | 0.519 | 0.106 |
| **894** | Iscu | 1.40E-08 | 3.082132778 | 0.000348225 | 0.298 | 0.01 |
| **895** | Ugdh | 1.40E-08 | 3.598862949 | 0.000348225 | 0.298 | 0.01 |
| **896** | Ikbip | 1.40E-08 | 2.750963466 | 0.000349408 | 0.346 | 0.029 |
| **897** | Prnp | 1.42E-08 | 1.538737026 | 0.000352143 | 0.471 | 0.087 |
| **898** | mt-Nd4l | 1.43E-08 | 0.827723127 | 0.000355169 | 0.99 | 0.798 |
| **899** | Psme2 | 1.43E-08 | 3.170885657 | 0.000355766 | 0.298 | 0.01 |

| | | | | | | |
|---|---|---|---|---|---|---|
| **900** | Ndufa8 | 1.43E-08 | 2.414427144 | 0.000356379 | 0.346 | 0.029 |
| **901** | Mesd | 1.44E-08 | 1.490066701 | 0.00035827 | 0.394 | 0.048 |
| **902** | Adam23 | 1.44E-08 | 0.833416938 | 0.000359484 | 0.442 | 0.067 |
| **903** | Slc9a3r2 | 1.45E-08 | 0.972034451 | 0.000359704 | 0.548 | 0.115 |
| **904** | Gstp1 | 1.46E-08 | 2.208423065 | 0.000363466 | 0.298 | 0.01 |
| **905** | Leprotl1 | 1.46E-08 | 2.733452738 | 0.000363466 | 0.298 | 0.01 |
| **906** | Inf2 | 1.48E-08 | 0.934396238 | 0.000368802 | 0.462 | 0.077 |
| **907** | Rcn2 | 1.49E-08 | 2.007715582 | 0.000370728 | 0.346 | 0.029 |
| **908** | Mylip | 1.50E-08 | 0.664704485 | 0.000372269 | 0.712 | 0.212 |
| **909** | Dhx36 | 1.52E-08 | 0.938199529 | 0.000377694 | 0.567 | 0.125 |
| **910** | Tsnax | 1.53E-08 | 0.547642595 | 0.000380662 | 0.558 | 0.125 |
| **911** | Btbd3 | 1.53E-08 | 0.6633976 | 0.000381028 | 0.712 | 0.192 |
| **912** | Pabpc1 | 1.55E-08 | 1.260455886 | 0.000385734 | 0.567 | 0.135 |
| **913** | Atp5o | 1.58E-08 | 4.966089765 | 0.000392562 | 0.269 | 0 |
| **914** | Rex1bd | 1.58E-08 | 4.863084991 | 0.000392562 | 0.269 | 0 |
| **915** | Slc25a5 | 1.58E-08 | 4.934264129 | 0.000392562 | 0.269 | 0 |
| **916** | Slu7 | 1.58E-08 | 5.11595024 | 0.000392562 | 0.269 | 0 |
| **917** | Tagln | 1.58E-08 | 5.479553765 | 0.000392562 | 0.269 | 0 |
| **918** | Sox4 | 1.59E-08 | 3.453035834 | 0.00039498 | 0.317 | 0.019 |
| **919** | Zc3hav1 | 1.60E-08 | 0.652644864 | 0.000397884 | 0.673 | 0.173 |
| **920** | Cpne3 | 1.60E-08 | 0.96608456 | 0.000398564 | 0.5 | 0.096 |
| **921** | Arid5b | 1.62E-08 | 0.603568605 | 0.000402025 | 0.865 | 0.308 |
| **922** | Serinc1 | 1.63E-08 | 1.221677166 | 0.000404759 | 0.644 | 0.183 |
| **923** | Vps35 | 1.64E-08 | 0.299578694 | 0.000408738 | 0.567 | 0.125 |
| **924** | Pla1a | 1.64E-08 | 2.115213501 | 0.0004091 | 0.346 | 0.029 |
| **925** | Magt1 | 1.66E-08 | 0.956849411 | 0.000413582 | 0.442 | 0.067 |
| **926** | Zdhhc14 | 1.68E-08 | 0.404441077 | 0.000417025 | 0.769 | 0.24 |
| **927** | Ids | 1.68E-08 | 1.872983476 | 0.000417225 | 0.346 | 0.029 |
| **928** | Tm9sf2 | 1.68E-08 | 0.692610075 | 0.000417448 | 0.779 | 0.24 |
| **929** | Eps15l1 | 1.71E-08 | 0.53926366 | 0.000426331 | 0.702 | 0.183 |
| **930** | Cdc42ep2 | 1.73E-08 | 1.583665209 | 0.00043115 | 0.365 | 0.038 |
| **931** | Ap1ar | 1.74E-08 | 1.699034823 | 0.000433947 | 0.346 | 0.029 |
| **932** | Timm17a | 1.78E-08 | 2.00603214 | 0.00044255 | 0.346 | 0.029 |
| **933** | Srsf10 | 1.80E-08 | 0.721090356 | 0.000448104 | 0.596 | 0.144 |
| **934** | Ankrd50 | 1.80E-08 | 1.747804263 | 0.000448119 | 0.433 | 0.067 |
| **935** | Nrp1 | 1.80E-08 | 0.672078191 | 0.00044841 | 1 | 0.625 |
| **936** | Sec24c | 1.85E-08 | 1.840253401 | 0.000460256 | 0.346 | 0.029 |
| **937** | Zfp706 | 1.85E-08 | 1.583530038 | 0.000460256 | 0.346 | 0.029 |
| **938** | Hecw2 | 1.86E-08 | 0.463312795 | 0.000462587 | 0.933 | 0.317 |
| **939** | Kdm6b | 1.89E-08 | 1.408816096 | 0.000469365 | 0.346 | 0.029 |
| **940** | Ep300 | 1.94E-08 | 0.750910893 | 0.000482491 | 0.683 | 0.192 |

| | | | | | | |
|---|---|---|---|---|---|---|
| **941** | Rpl36al | 1.94E-08 | 2.258243612 | 0.00048333 | 0.365 | 0.038 |
| **942** | Gabarapl2 | 1.97E-08 | 1.546002065 | 0.00048921 | 0.433 | 0.067 |
| **943** | Slc30a4 | 1.99E-08 | 2.104424559 | 0.000496182 | 0.385 | 0.048 |
| **944** | Cdc42ep1 | 2.01E-08 | 1.463952837 | 0.000500766 | 0.413 | 0.058 |
| **945** | Ppp4r3b | 2.03E-08 | 0.703571467 | 0.000504234 | 0.683 | 0.192 |
| **946** | mt-Co2 | 2.17E-08 | 0.725040382 | 0.000539127 | 1 | 0.885 |
| **947** | D8Ertd738e | 2.18E-08 | 1.443539657 | 0.00054162 | 0.365 | 0.038 |
| **948** | Ginm1 | 2.23E-08 | 0.893637737 | 0.000555964 | 0.462 | 0.077 |
| **949** | Terf2 | 2.26E-08 | 0.920785455 | 0.000563486 | 0.5 | 0.096 |
| **950** | Creg1 | 2.29E-08 | 2.560737121 | 0.000570897 | 0.317 | 0.019 |
| **951** | Irf2bpl | 2.29E-08 | 2.377174966 | 0.000570897 | 0.317 | 0.019 |
| **952** | Notch1 | 2.31E-08 | 0.293322563 | 0.000574869 | 0.817 | 0.269 |
| **953** | Sri | 2.34E-08 | 1.079103668 | 0.000581235 | 0.442 | 0.077 |
| **954** | Cops9 | 2.34E-08 | 1.93603233 | 0.000582634 | 0.317 | 0.019 |
| **955** | Nono | 2.38E-08 | 0.888539353 | 0.000591894 | 0.5 | 0.096 |
| **956** | Spcs1 | 2.39E-08 | 2.099517786 | 0.000594604 | 0.317 | 0.019 |
| **957** | Zfp646 | 2.39E-08 | 2.279425962 | 0.000594604 | 0.317 | 0.019 |
| **958** | Baz2a | 2.44E-08 | 0.350496138 | 0.000606399 | 0.587 | 0.135 |
| **959** | Ppp1r9b | 2.48E-08 | 0.452113335 | 0.000618142 | 0.548 | 0.115 |
| **960** | Dpysl2 | 2.53E-08 | 0.570920176 | 0.000628695 | 0.865 | 0.279 |
| **961** | Rala | 2.53E-08 | 0.532583353 | 0.000630246 | 0.654 | 0.163 |
| **962** | Il10rb | 2.54E-08 | 0.915410041 | 0.00063194 | 0.5 | 0.096 |
| **963** | Rpl10a | 2.59E-08 | 1.275808817 | 0.000645338 | 0.49 | 0.096 |
| **964** | Ube2v2 | 2.60E-08 | 1.40551719 | 0.00064615 | 0.481 | 0.096 |
| **965** | Sh3bgrl | 2.62E-08 | 0.643264878 | 0.000652854 | 0.923 | 0.356 |
| **966** | Neurl1b | 2.68E-08 | 1.069624294 | 0.000666927 | 0.356 | 0.038 |
| **967** | 1700025G04R | 2.69E-08 | 0.418536491 | 0.000668099 | 0.788 | 0.25 |
| **968** | Prxl2c | 2.69E-08 | 0.814844949 | 0.000669667 | 0.433 | 0.067 |
| **969** | Sfmbt2 | 2.73E-08 | 3.128934045 | 0.000678341 | 0.288 | 0.01 |
| **970** | Nudt4 | 2.83E-08 | 1.082995285 | 0.000704773 | 0.452 | 0.077 |
| **971** | Plscr3 | 2.84E-08 | 2.195420812 | 0.000707848 | 0.288 | 0.01 |
| **972** | Prkar1b | 2.84E-08 | 2.542952871 | 0.000707848 | 0.288 | 0.01 |
| **973** | Rars | 2.84E-08 | 2.312341747 | 0.000707848 | 0.288 | 0.01 |
| **974** | Rnpepl1 | 2.84E-08 | 2.921649126 | 0.000707848 | 0.288 | 0.01 |
| **975** | Taldo1 | 2.84E-08 | 2.913513939 | 0.000707848 | 0.288 | 0.01 |
| **976** | Prdx1 | 2.87E-08 | 0.878564651 | 0.000713754 | 0.529 | 0.115 |
| **977** | Tmem140 | 2.89E-08 | 0.640910989 | 0.000718671 | 0.721 | 0.212 |
| **978** | Gcc2 | 2.91E-08 | 0.952019187 | 0.000723054 | 0.51 | 0.106 |
| **979** | Cd46 | 2.92E-08 | 1.477130034 | 0.000726338 | 0.442 | 0.077 |
| **980** | Enpp2 | 2.93E-08 | 0.633347654 | 0.000729852 | 0.519 | 0.106 |
| **981** | Tapbp | 2.97E-08 | 0.779418222 | 0.000738485 | 0.702 | 0.202 |

| | | | | | | |
|---|---|---|---|---|---|---|
| **982** | Stx5a | 2.97E-08 | 0.997442548 | 0.000738568 | 0.471 | 0.087 |
| **983** | Ppp2ca | 3.00E-08 | 0.7338315 | 0.000745687 | 0.712 | 0.212 |
| **984** | Ctbp1 | 3.01E-08 | 1.049045567 | 0.000749506 | 0.481 | 0.096 |
| **985** | Zfp652 | 3.04E-08 | 0.399254359 | 0.000755803 | 0.769 | 0.24 |
| **986** | Aplnr | 3.08E-08 | 5.422386406 | 0.000765391 | 0.26 | 0 |
| **987** | Cox4i2 | 3.08E-08 | 5.317755539 | 0.000765391 | 0.26 | 0 |
| **988** | Map2 | 3.08E-08 | 5.513872486 | 0.000765391 | 0.26 | 0 |
| **989** | Ndufa4l2 | 3.08E-08 | 5.642154606 | 0.000765391 | 0.26 | 0 |
| **990** | Rtl8a | 3.08E-08 | 5.424664584 | 0.000765391 | 0.26 | 0 |
| **991** | Smim11 | 3.08E-08 | 5.112486689 | 0.000765391 | 0.26 | 0 |
| **992** | Robo4 | 3.08E-08 | 0.760580687 | 0.000766093 | 0.673 | 0.183 |
| **993** | Cox7a2 | 3.10E-08 | 1.757655973 | 0.000770947 | 0.337 | 0.029 |
| **994** | Fam168b | 3.18E-08 | 0.475273453 | 0.00079075 | 0.548 | 0.125 |
| **995** | Cep295 | 3.22E-08 | 1.494713057 | 0.000801608 | 0.337 | 0.029 |
| **996** | Rtf1 | 3.23E-08 | 0.748399405 | 0.000804692 | 0.625 | 0.163 |
| **997** | Slc39a7 | 3.29E-08 | 1.156733846 | 0.000818287 | 0.423 | 0.067 |
| **998** | Ldha | 3.30E-08 | 1.628089644 | 0.000821507 | 0.356 | 0.038 |
| **999** | Armcx3 | 3.33E-08 | 3.34271826 | 0.000829088 | 0.308 | 0.019 |
| **1000** | Dnajc8 | 3.36E-08 | 1.168818146 | 0.000834929 | 0.452 | 0.077 |
| **1001** | Astn2 | 3.37E-08 | 1.737005149 | 0.000837377 | 0.413 | 0.067 |
| **1002** | Zranb2 | 3.44E-08 | 0.740884974 | 0.00085599 | 0.538 | 0.115 |
| **1003** | Sbds | 3.55E-08 | 1.502271872 | 0.000883524 | 0.337 | 0.029 |
| **1004** | Atp6v1e1 | 3.56E-08 | 1.657006349 | 0.000886255 | 0.375 | 0.048 |
| **1005** | Rbbp4 | 3.58E-08 | 0.683372773 | 0.00089078 | 0.596 | 0.144 |
| **1006** | Txnrd1 | 3.59E-08 | 1.604652258 | 0.000892491 | 0.423 | 0.067 |
| **1007** | Etfdh | 3.62E-08 | 1.570174395 | 0.000900855 | 0.337 | 0.029 |
| **1008** | Slit3 | 3.65E-08 | 0.516993928 | 0.000907946 | 0.99 | 0.663 |
| **1009** | Tcim | 3.65E-08 | 0.878917908 | 0.00090848 | 0.385 | 0.048 |
| **1010** | Mob1a | 3.66E-08 | 1.815371197 | 0.000909658 | 0.481 | 0.106 |
| **1011** | Ptpn13 | 3.66E-08 | 1.007260104 | 0.000909658 | 0.49 | 0.096 |
| **1012** | St13 | 3.68E-08 | 0.646706525 | 0.000914419 | 0.635 | 0.163 |
| **1013** | Acaa2 | 3.70E-08 | 0.746102378 | 0.000920688 | 0.404 | 0.058 |
| **1014** | Ptges3 | 3.81E-08 | 0.360710722 | 0.000947508 | 0.625 | 0.163 |
| **1015** | 9930111J21Ril | 3.82E-08 | 0.968968578 | 0.000949856 | 0.51 | 0.106 |
| **1016** | Ndrg3 | 3.83E-08 | 1.135094232 | 0.000953878 | 0.404 | 0.058 |
| **1017** | Itga7 | 3.84E-08 | 2.393510915 | 0.000955885 | 0.308 | 0.019 |
| **1018** | Myct1 | 3.86E-08 | 0.794079732 | 0.000959314 | 0.538 | 0.125 |
| **1019** | Ppp1r2 | 3.92E-08 | 0.805869892 | 0.000975102 | 0.904 | 0.337 |
| **1020** | Klhl24 | 3.93E-08 | 0.580822461 | 0.000977886 | 0.712 | 0.212 |
| **1021** | Tcf25 | 4.03E-08 | 0.410812961 | 0.001002668 | 0.798 | 0.25 |
| **1022** | Pcnp | 4.06E-08 | 0.594640353 | 0.00100962 | 0.567 | 0.135 |

| | | | | | | |
|---|---|---|---|---|---|---|
| **1023** | Dnaja2 | 4.10E-08 | 1.098799689 | 0.001020459 | 0.442 | 0.077 |
| **1024** | Ndufa6 | 4.17E-08 | 2.440804037 | 0.001036583 | 0.308 | 0.019 |
| **1025** | Rab14 | 4.18E-08 | 0.680719552 | 0.001039612 | 0.76 | 0.231 |
| **1026** | 2700081O15R | 4.19E-08 | 0.556948934 | 0.001042828 | 0.423 | 0.067 |
| **1027** | Atp6v0c | 4.20E-08 | 1.993508821 | 0.001044836 | 0.375 | 0.048 |
| **1028** | Rnf11 | 4.24E-08 | 1.375348727 | 0.001055549 | 0.442 | 0.077 |
| **1029** | Sirt1 | 4.26E-08 | 1.34034217 | 0.001060559 | 0.404 | 0.058 |
| **1030** | Smim10l1 | 4.26E-08 | 0.897335434 | 0.001060975 | 0.423 | 0.067 |
| **1031** | Nars | 4.30E-08 | 1.400073669 | 0.001068868 | 0.413 | 0.067 |
| **1032** | Setbp1 | 4.40E-08 | 0.717562754 | 0.001094817 | 0.692 | 0.212 |
| **1033** | Pdgfb | 4.40E-08 | 0.891596646 | 0.0010954 | 0.712 | 0.212 |
| **1034** | Swi5 | 4.42E-08 | 1.895779892 | 0.001100208 | 0.327 | 0.029 |
| **1035** | Neat1 | 4.45E-08 | -1.22048177 | 0.001108427 | 0.942 | 0.846 |
| **1036** | Med23 | 4.49E-08 | 0.64990995 | 0.001117252 | 0.423 | 0.067 |
| **1037** | Vdac2 | 4.50E-08 | 1.059886746 | 0.001119361 | 0.462 | 0.087 |
| **1038** | Bcas2 | 4.52E-08 | 1.78780909 | 0.001123869 | 0.308 | 0.019 |
| **1039** | Stard8 | 4.55E-08 | 0.38432983 | 0.001131521 | 0.692 | 0.202 |
| **1040** | Araf | 4.60E-08 | 0.84462224 | 0.001145495 | 0.529 | 0.115 |
| **1041** | Cdkl3 | 4.61E-08 | 1.917552862 | 0.001146781 | 0.308 | 0.019 |
| **1042** | 2900097C17Ri | 4.62E-08 | 1.201869522 | 0.001148469 | 0.442 | 0.077 |
| **1043** | Tmem204 | 4.64E-08 | 0.372203602 | 0.001154358 | 0.471 | 0.087 |
| **1044** | Zbtb11 | 4.67E-08 | 1.386326775 | 0.001161289 | 0.394 | 0.058 |
| **1045** | Nsa2 | 4.69E-08 | 2.443978583 | 0.001166471 | 0.327 | 0.029 |
| **1046** | Ddx46 | 4.70E-08 | 0.772664666 | 0.001168633 | 0.577 | 0.144 |
| **1047** | Lamtor2 | 4.75E-08 | 3.860163104 | 0.001181601 | 0.279 | 0.01 |
| **1048** | Anxa2 | 4.84E-08 | 0.623071003 | 0.001203776 | 0.644 | 0.173 |
| **1049** | Rap2c | 4.87E-08 | 2.08302393 | 0.001212778 | 0.327 | 0.029 |
| **1050** | Zup1 | 4.94E-08 | 0.725181277 | 0.001228175 | 0.519 | 0.115 |
| **1051** | Esyt1 | 4.95E-08 | 1.342031268 | 0.001230785 | 0.375 | 0.048 |
| **1052** | Zc3h15 | 4.95E-08 | 0.529500815 | 0.001232684 | 0.471 | 0.087 |
| **1053** | Brd3 | 5.05E-08 | 0.811245136 | 0.001256543 | 0.462 | 0.087 |
| **1054** | H2-Q7 | 5.05E-08 | 1.097284986 | 0.001256543 | 0.462 | 0.087 |
| **1055** | Rabl6 | 5.13E-08 | 0.833531934 | 0.001277425 | 0.462 | 0.087 |
| **1056** | Anp32b | 5.14E-08 | 0.719101897 | 0.001277737 | 0.49 | 0.096 |
| **1057** | Ccl11 | 5.17E-08 | 3.519219052 | 0.001286246 | 0.279 | 0.01 |
| **1058** | Map1lc3a | 5.17E-08 | 3.801949172 | 0.001286246 | 0.279 | 0.01 |
| **1059** | Ndufb7 | 5.17E-08 | 3.649825522 | 0.001286246 | 0.279 | 0.01 |
| **1060** | Wipi1 | 5.19E-08 | 1.742987168 | 0.001291432 | 0.394 | 0.058 |
| **1061** | Zfyve9 | 5.20E-08 | 0.535461994 | 0.001294525 | 0.663 | 0.183 |
| **1062** | Atad1 | 5.24E-08 | 0.727855849 | 0.001304091 | 0.423 | 0.067 |
| **1063** | Ggps1 | 5.24E-08 | 1.194881774 | 0.001304091 | 0.423 | 0.067 |

| | | | | | | |
|---|---|---|---|---|---|---|
| **1064** | Slc25a12 | 5.26E-08 | 0.77461689 | 0.001307541 | 0.471 | 0.096 |
| **1065** | Atp6v1g1 | 5.28E-08 | 2.211712792 | 0.001313528 | 0.346 | 0.038 |
| **1066** | Lrfn2 | 5.28E-08 | 3.225556546 | 0.00131378 | 0.279 | 0.01 |
| **1067** | Dusp7 | 5.34E-08 | 3.493648822 | 0.00132759 | 0.298 | 0.019 |
| **1068** | Trpv2 | 5.39E-08 | 2.598627853 | 0.001341884 | 0.279 | 0.01 |
| **1069** | Ttc39b | 5.43E-08 | 0.465473601 | 0.001351627 | 0.663 | 0.183 |
| **1070** | Pank3 | 5.46E-08 | 0.893932741 | 0.001359651 | 0.548 | 0.125 |
| **1071** | Serpine2 | 5.51E-08 | 1.441988031 | 0.00137057 | 0.279 | 0.01 |
| **1072** | Tmem134 | 5.51E-08 | 2.688106475 | 0.00137057 | 0.279 | 0.01 |
| **1073** | Ddb1 | 5.52E-08 | 0.823910821 | 0.00137284 | 0.423 | 0.067 |
| **1074** | Sf3b3 | 5.54E-08 | 0.816338051 | 0.001377829 | 0.529 | 0.125 |
| **1075** | Gm37494 | 5.59E-08 | 0.521780203 | 0.00138965 | 0.625 | 0.163 |
| **1076** | Rhoc | 5.67E-08 | 0.998183316 | 0.001410586 | 0.394 | 0.058 |
| **1077** | Tmx1 | 5.67E-08 | 1.210498396 | 0.001410586 | 0.394 | 0.058 |
| **1078** | Lysmd2 | 5.69E-08 | 2.027029806 | 0.001416479 | 0.327 | 0.029 |
| **1079** | Hnrnpdl | 5.72E-08 | 0.858130524 | 0.001422937 | 0.5 | 0.106 |
| **1080** | Ostf1 | 5.73E-08 | 0.24694368 | 0.001426002 | 0.673 | 0.183 |
| **1081** | Plaa | 5.76E-08 | 1.518318945 | 0.001434137 | 0.413 | 0.067 |
| **1082** | Lsm14a | 5.81E-08 | 0.367164734 | 0.001446235 | 0.798 | 0.231 |
| **1083** | Fstl1 | 5.84E-08 | 0.652522714 | 0.001452246 | 0.712 | 0.212 |
| **1084** | Csnk2a2 | 5.90E-08 | 0.677880722 | 0.001466786 | 0.606 | 0.154 |
| **1085** | Fkbp9 | 5.90E-08 | 0.914175002 | 0.00146804 | 0.548 | 0.125 |
| **1086** | B3galt1 | 5.91E-08 | 0.973187906 | 0.00147023 | 0.76 | 0.26 |
| **1087** | Efcab14 | 5.96E-08 | 0.915340409 | 0.00148197 | 0.519 | 0.115 |
| **1088** | Tmem260 | 5.96E-08 | 1.165082937 | 0.00148197 | 0.519 | 0.115 |
| **1089** | 4930502A04Ri | 5.96E-08 | 5.433875469 | 0.00148398 | 0.25 | 0 |
| **1090** | Atp5g1 | 5.96E-08 | 4.963061826 | 0.00148398 | 0.25 | 0 |
| **1091** | Cirbp | 5.96E-08 | 4.637745772 | 0.00148398 | 0.25 | 0 |
| **1092** | Fzd1 | 5.96E-08 | 5.21048126 | 0.00148398 | 0.25 | 0 |
| **1093** | Isg15 | 5.96E-08 | 5.49512065 | 0.00148398 | 0.25 | 0 |
| **1094** | Ndufaf8 | 5.96E-08 | 4.779455521 | 0.00148398 | 0.25 | 0 |
| **1095** | Ptger2 | 5.96E-08 | 5.442363679 | 0.00148398 | 0.25 | 0 |
| **1096** | S1pr3 | 5.96E-08 | 5.315971504 | 0.00148398 | 0.25 | 0 |
| **1097** | Tmem258 | 5.96E-08 | 5.030784954 | 0.00148398 | 0.25 | 0 |
| **1098** | Tubb4b | 5.96E-08 | 4.929654581 | 0.00148398 | 0.25 | 0 |
| **1099** | Ndufs7 | 6.03E-08 | 3.322709371 | 0.001499517 | 0.298 | 0.019 |
| **1100** | Timm23 | 6.04E-08 | 1.660034397 | 0.001501897 | 0.375 | 0.048 |
| **1101** | C4b | 6.10E-08 | 1.859871959 | 0.001517818 | 0.365 | 0.048 |
| **1102** | Pde5a | 6.14E-08 | 0.993310962 | 0.001527515 | 0.442 | 0.077 |
| **1103** | Dazap2 | 6.15E-08 | 2.1420161 | 0.001530395 | 0.327 | 0.029 |
| **1104** | Tecpr1 | 6.29E-08 | 0.740878939 | 0.001564333 | 0.529 | 0.115 |

| | | | | | | |
|---|---|---|---|---|---|---|
| **1105** | Unc45b | 6.29E-08 | 0.596826049 | 0.001564864 | 0.5 | 0.106 |
| **1106** | Snx27 | 6.35E-08 | 0.529764195 | 0.001579107 | 0.606 | 0.154 |
| **1107** | Ppp1r15b | 6.49E-08 | 0.861330721 | 0.001614163 | 0.375 | 0.048 |
| **1108** | Tmem219 | 6.49E-08 | 1.36248762 | 0.001614163 | 0.375 | 0.048 |
| **1109** | Thap12 | 6.49E-08 | 1.497939782 | 0.001614365 | 0.346 | 0.038 |
| **1110** | Vat1 | 6.53E-08 | 1.322654651 | 0.001623704 | 0.394 | 0.058 |
| **1111** | Sar1a | 6.55E-08 | 1.471110664 | 0.001629005 | 0.433 | 0.077 |
| **1112** | Ilk | 6.61E-08 | 1.706552361 | 0.001644809 | 0.346 | 0.038 |
| **1113** | Slc50a1 | 6.64E-08 | 1.816211645 | 0.001653165 | 0.327 | 0.029 |
| **1114** | Snapc3 | 6.65E-08 | 0.305786958 | 0.001655201 | 0.875 | 0.288 |
| **1115** | Fam43a | 6.66E-08 | 1.343643179 | 0.00165657 | 0.433 | 0.077 |
| **1116** | 4632427E13Ri | 6.72E-08 | 0.220391357 | 0.001673066 | 0.577 | 0.135 |
| **1117** | Paics | 6.74E-08 | 1.934513133 | 0.001675808 | 0.346 | 0.038 |
| **1118** | Mtmr6 | 6.75E-08 | 0.892800106 | 0.001678659 | 0.519 | 0.115 |
| **1119** | Usp14 | 6.76E-08 | 1.36621682 | 0.001681675 | 0.394 | 0.058 |
| **1120** | Ap2m1 | 6.77E-08 | 1.379762495 | 0.001685318 | 0.327 | 0.029 |
| **1121** | Selenot | 6.90E-08 | 0.333594492 | 0.001716279 | 0.481 | 0.096 |
| **1122** | Pdha1 | 6.91E-08 | 1.162471002 | 0.001718076 | 0.327 | 0.029 |
| **1123** | Eif3c | 7.07E-08 | 0.79433766 | 0.001760184 | 0.49 | 0.096 |
| **1124** | Rbm3 | 7.14E-08 | 0.644363012 | 0.001775586 | 0.5 | 0.106 |
| **1125** | Ripor1 | 7.15E-08 | 0.7525072 | 0.001779473 | 0.538 | 0.125 |
| **1126** | Mgst1 | 7.26E-08 | 0.735156933 | 0.001805785 | 0.577 | 0.144 |
| **1127** | Sfxn1 | 7.26E-08 | 1.604758195 | 0.001805838 | 0.385 | 0.058 |
| **1128** | Adgra3 | 7.30E-08 | 0.754645635 | 0.001816759 | 0.567 | 0.144 |
| **1129** | Tppp3 | 7.38E-08 | 2.577736692 | 0.001835078 | 0.298 | 0.019 |
| **1130** | Hp1bp3 | 7.41E-08 | 0.83120675 | 0.001844138 | 0.567 | 0.144 |
| **1131** | Mgat4b | 7.45E-08 | 1.634231636 | 0.001853043 | 0.365 | 0.048 |
| **1132** | Hoxc6 | 7.53E-08 | 1.820150444 | 0.001874038 | 0.346 | 0.038 |
| **1133** | Mllt6 | 7.62E-08 | 1.121599624 | 0.001896836 | 0.471 | 0.096 |
| **1134** | Krcc1 | 7.64E-08 | 0.504428027 | 0.001900367 | 0.587 | 0.144 |
| **1135** | Bod1l | 7.74E-08 | 0.720771122 | 0.001926876 | 0.615 | 0.163 |
| **1136** | Rassf1 | 7.84E-08 | 0.922625359 | 0.001951408 | 0.5 | 0.106 |
| **1137** | Clic1 | 8.15E-08 | 0.234775633 | 0.00202758 | 0.51 | 0.106 |
| **1138** | Uxs1 | 8.15E-08 | 1.621769035 | 0.002028152 | 0.365 | 0.048 |
| **1139** | Gpalpp1 | 8.16E-08 | 1.89393445 | 0.002029077 | 0.298 | 0.019 |
| **1140** | Thap2 | 8.32E-08 | 2.534170673 | 0.002070192 | 0.298 | 0.019 |
| **1141** | Vcp | 8.33E-08 | 0.682059602 | 0.00207279 | 0.596 | 0.154 |
| **1142** | B3glct | 8.34E-08 | 0.812541195 | 0.002073968 | 0.394 | 0.058 |
| **1143** | Tecpr2 | 8.35E-08 | 1.207118422 | 0.002078801 | 0.385 | 0.058 |
| **1144** | Hnrnph2 | 8.40E-08 | 1.105634472 | 0.002091117 | 0.462 | 0.096 |
| **1145** | Ube2i | 8.40E-08 | 0.223201853 | 0.002091138 | 0.452 | 0.087 |

| | | | | | | |
|---|---|---|---|---|---|---|
| **1146** | Fbxo32 | 8.49E-08 | 1.978855832 | 0.002112114 | 0.298 | 0.019 |
| **1147** | Ulk2 | 8.54E-08 | 0.382027848 | 0.002123811 | 0.721 | 0.212 |
| **1148** | Dync1h1 | 8.62E-08 | 0.680874292 | 0.002144969 | 0.856 | 0.317 |
| **1149** | Mif | 8.66E-08 | 2.021515155 | 0.002154857 | 0.298 | 0.019 |
| **1150** | Clec3b | 8.67E-08 | 1.610056068 | 0.002158054 | 0.442 | 0.087 |
| **1151** | Plgrkt | 8.67E-08 | 0.14125256 | 0.002158106 | 0.51 | 0.106 |
| **1152** | Tob2 | 8.76E-08 | 1.671159084 | 0.002179675 | 0.365 | 0.048 |
| **1153** | Bag1 | 8.84E-08 | 1.438766218 | 0.002198437 | 0.298 | 0.019 |
| **1154** | Nme2 | 8.84E-08 | 1.962255194 | 0.002198437 | 0.298 | 0.019 |
| **1155** | Ankrd40 | 8.84E-08 | 1.147950828 | 0.00220043 | 0.433 | 0.077 |
| **1156** | Trim12c | 8.97E-08 | 0.987780699 | 0.002232118 | 0.452 | 0.087 |
| **1157** | Prpf6 | 8.99E-08 | 0.440082317 | 0.002236654 | 0.538 | 0.125 |
| **1158** | mt-Nd1 | 9.22E-08 | 0.6686482 | 0.002294264 | 1 | 0.731 |
| **1159** | Hif1a | 9.27E-08 | 0.872567554 | 0.002305284 | 0.558 | 0.135 |
| **1160** | Yars | 9.41E-08 | 1.55365172 | 0.002340861 | 0.346 | 0.038 |
| **1161** | Thrsp | 9.47E-08 | 0.494048087 | 0.002355025 | 0.5 | 0.106 |
| **1162** | Itgav | 9.85E-08 | 0.736534621 | 0.002450895 | 0.471 | 0.096 |
| **1163** | Leng8 | 9.92E-08 | 0.528838102 | 0.002467344 | 0.692 | 0.192 |
| **1164** | Phc3 | 9.96E-08 | 0.373635565 | 0.002478355 | 0.798 | 0.26 |
| **1165** | Sh3glb1 | 1.01E-07 | 0.747183613 | 0.002520595 | 0.923 | 0.394 |
| **1166** | Pcnx4 | 1.01E-07 | 1.401754549 | 0.002521015 | 0.337 | 0.038 |
| **1167** | Nkap | 1.04E-07 | 3.343395325 | 0.00258379 | 0.269 | 0.01 |
| **1168** | Tmco1 | 1.05E-07 | 0.778626503 | 0.00260499 | 0.413 | 0.067 |
| **1169** | Tmem184b | 1.05E-07 | 0.34291577 | 0.00261 | 0.769 | 0.24 |
| **1170** | Rnf4 | 1.06E-07 | 0.120175178 | 0.002626623 | 0.519 | 0.115 |
| **1171** | CAAA0114733 | 1.06E-07 | 2.952680952 | 0.00263868 | 0.269 | 0.01 |
| **1172** | Fkbp8 | 1.06E-07 | 2.252792729 | 0.00263868 | 0.269 | 0.01 |
| **1173** | Rps27l | 1.06E-07 | 2.003961021 | 0.00263868 | 0.269 | 0.01 |
| **1174** | Sidt2 | 1.07E-07 | 1.246053263 | 0.002655488 | 0.385 | 0.058 |
| **1175** | Pura | 1.08E-07 | 0.361532023 | 0.002682448 | 0.692 | 0.192 |
| **1176** | Anxa4 | 1.09E-07 | 1.694087596 | 0.002702138 | 0.385 | 0.058 |
| **1177** | Gns | 1.09E-07 | 1.351614492 | 0.00272064 | 0.404 | 0.067 |
| **1178** | Stx12 | 1.10E-07 | 0.119073486 | 0.002748064 | 0.635 | 0.173 |
| **1179** | Pnrc2 | 1.11E-07 | 2.136141026 | 0.002766517 | 0.337 | 0.038 |
| **1180** | Ppm1a | 1.12E-07 | 0.509839619 | 0.002788114 | 0.529 | 0.125 |
| **1181** | Fip1l1 | 1.12E-07 | 0.80297251 | 0.002796665 | 0.683 | 0.202 |
| **1182** | Adh1 | 1.14E-07 | 0.788854658 | 0.002834251 | 0.413 | 0.067 |
| **1183** | Lrpap1 | 1.15E-07 | 1.372498506 | 0.00285166 | 0.365 | 0.048 |
| **1184** | Ffar2 | 1.15E-07 | 5.439366188 | 0.002861451 | 0.24 | 0 |
| **1185** | Mrps10 | 1.15E-07 | 4.817290928 | 0.002861451 | 0.24 | 0 |
| **1186** | Yipf3 | 1.15E-07 | 4.760109112 | 0.002861451 | 0.24 | 0 |

| | | | | | | |
|---|---|---|---|---|---|---|
| **1187** | Gm26917 | 1.15E-07 | -1.26887486 | 0.002867466 | 0.962 | 0.923 |
| **1188** | Plk2 | 1.15E-07 | 1.882465349 | 0.002871063 | 0.337 | 0.038 |
| **1189** | Kpna1 | 1.16E-07 | 0.705578525 | 0.00287729 | 0.587 | 0.154 |
| **1190** | Sik1 | 1.16E-07 | 0.888871067 | 0.002882338 | 0.462 | 0.096 |
| **1191** | Creb5 | 1.17E-07 | 0.201652028 | 0.002921264 | 0.625 | 0.173 |
| **1192** | Dr1 | 1.19E-07 | 1.871933614 | 0.00295954 | 0.317 | 0.029 |
| **1193** | Tmem88 | 1.19E-07 | 2.267021907 | 0.00295954 | 0.317 | 0.029 |
| **1194** | Smarca5 | 1.20E-07 | 0.476410233 | 0.002991711 | 0.625 | 0.163 |
| **1195** | Hspa12a | 1.21E-07 | 0.7569256 | 0.003008281 | 0.365 | 0.048 |
| **1196** | C1ra | 1.21E-07 | 2.040762173 | 0.003016673 | 0.317 | 0.029 |
| **1197** | Setd1b | 1.23E-07 | 1.281952188 | 0.00305163 | 0.385 | 0.058 |
| **1198** | Sugt1 | 1.23E-07 | 1.23474105 | 0.003063029 | 0.433 | 0.077 |
| **1199** | Clpx | 1.23E-07 | 0.642356564 | 0.003072016 | 0.538 | 0.125 |
| **1200** | Trpc1 | 1.24E-07 | 1.352881593 | 0.003074873 | 0.317 | 0.029 |
| **1201** | Gps2 | 1.24E-07 | 2.025604316 | 0.003091744 | 0.337 | 0.038 |
| **1202** | Plod1 | 1.25E-07 | 0.850363158 | 0.003110578 | 0.471 | 0.096 |
| **1203** | Sf3b1 | 1.25E-07 | 0.576009776 | 0.003120181 | 0.865 | 0.298 |
| **1204** | Matr3 | 1.26E-07 | 0.376520302 | 0.003122682 | 0.625 | 0.163 |
| **1205** | Prelp | 1.26E-07 | 1.448989973 | 0.003134158 | 0.317 | 0.029 |
| **1206** | Rassf8 | 1.26E-07 | 0.720504054 | 0.003145044 | 0.615 | 0.173 |
| **1207** | Dynll2 | 1.27E-07 | 1.118776626 | 0.003159267 | 0.385 | 0.058 |
| **1208** | Scand1 | 1.27E-07 | 3.150561116 | 0.003168693 | 0.288 | 0.019 |
| **1209** | Fuca1 | 1.28E-07 | 1.419331681 | 0.003194549 | 0.317 | 0.029 |
| **1210** | Tmem263 | 1.30E-07 | 2.357980607 | 0.003232806 | 0.288 | 0.019 |
| **1211** | Lbr | 1.30E-07 | 0.756823911 | 0.003238834 | 0.567 | 0.144 |
| **1212** | Aamp | 1.31E-07 | 1.760145974 | 0.003256064 | 0.317 | 0.029 |
| **1213** | Pcdh12 | 1.31E-07 | 1.336979513 | 0.003256064 | 0.317 | 0.029 |
| **1214** | Map3k5 | 1.33E-07 | 0.65478627 | 0.003305967 | 0.875 | 0.327 |
| **1215** | Brd1 | 1.34E-07 | 0.378605892 | 0.00332876 | 0.702 | 0.202 |
| **1216** | Ndufa13 | 1.34E-07 | 1.936872014 | 0.003328796 | 0.337 | 0.038 |
| **1217** | Etfb | 1.34E-07 | 0.889858744 | 0.003333933 | 0.404 | 0.067 |
| **1218** | Psmc6 | 1.36E-07 | 0.675385305 | 0.003393022 | 0.529 | 0.125 |
| **1219** | Shroom3 | 1.36E-07 | 0.680718774 | 0.003393022 | 0.529 | 0.125 |
| **1220** | Atf7 | 1.37E-07 | 0.513228626 | 0.003405746 | 0.894 | 0.327 |
| **1221** | Eprs | 1.37E-07 | 1.222082269 | 0.003405851 | 0.442 | 0.087 |
| **1222** | Eef1g | 1.38E-07 | 0.545194511 | 0.0034361 | 0.433 | 0.077 |
| **1223** | Fam53b | 1.39E-07 | 1.789992179 | 0.003453823 | 0.337 | 0.038 |
| **1224** | Ccn1 | 1.39E-07 | 1.31523001 | 0.003460842 | 0.308 | 0.029 |
| **1225** | Vti1b | 1.41E-07 | 1.100825285 | 0.00351195 | 0.433 | 0.087 |
| **1226** | Plscr2 | 1.43E-07 | 1.06777886 | 0.003566614 | 0.404 | 0.067 |
| **1227** | Smc3 | 1.45E-07 | 0.408879136 | 0.003600989 | 0.558 | 0.135 |

| | | | | | | |
|---|---|---|---|---|---|---|
| **1228** | Safb | 1.45E-07 | 0.981248227 | 0.003602283 | 0.606 | 0.163 |
| **1229** | Enpp4 | 1.47E-07 | 1.444483709 | 0.003665601 | 0.423 | 0.077 |
| **1230** | Rev1 | 1.48E-07 | 0.363863026 | 0.003677469 | 0.538 | 0.125 |
| **1231** | Tbc1d15 | 1.48E-07 | 0.565329227 | 0.00369231 | 0.5 | 0.106 |
| **1232** | Arhgef10l | 1.51E-07 | 1.160151215 | 0.003754952 | 0.385 | 0.058 |
| **1233** | Atp5c1 | 1.51E-07 | 0.766285513 | 0.003765457 | 0.51 | 0.115 |
| **1234** | Nelfa | 1.52E-07 | 1.550317453 | 0.003786587 | 0.337 | 0.038 |
| **1235** | 2-Mar | 1.52E-07 | 0.359237467 | 0.003789006 | 0.538 | 0.125 |
| **1236** | Ipo8 | 1.53E-07 | 0.449243765 | 0.003818665 | 0.471 | 0.096 |
| **1237** | Dnajc3 | 1.56E-07 | 0.307041973 | 0.003879884 | 0.625 | 0.173 |
| **1238** | Secisbp2l | 1.58E-07 | 0.237762089 | 0.003924653 | 0.606 | 0.163 |
| **1239** | Pcnt | 1.60E-07 | 1.520074028 | 0.003977421 | 0.394 | 0.067 |
| **1240** | Ankle2 | 1.62E-07 | 0.893616738 | 0.004022471 | 0.385 | 0.058 |
| **1241** | Trp53 | 1.62E-07 | 2.407966257 | 0.004026244 | 0.288 | 0.019 |
| **1242** | Nedd8 | 1.64E-07 | 1.483144107 | 0.004074922 | 0.337 | 0.038 |
| **1243** | Cct2 | 1.64E-07 | 0.694434945 | 0.004080004 | 0.404 | 0.067 |
| **1244** | Zfp467 | 1.66E-07 | 1.325091382 | 0.004130518 | 0.433 | 0.087 |
| **1245** | Spon1 | 1.67E-07 | 0.705863181 | 0.004147637 | 0.683 | 0.212 |
| **1246** | Cenpb | 1.68E-07 | 1.529873233 | 0.004189453 | 0.288 | 0.019 |
| **1247** | Phax | 1.68E-07 | 1.739526393 | 0.004189453 | 0.288 | 0.019 |
| **1248** | Trim28 | 1.68E-07 | 2.06444621 | 0.004189453 | 0.288 | 0.019 |
| **1249** | Ltbp1 | 1.68E-07 | 0.211459452 | 0.00419115 | 0.702 | 0.212 |
| **1250** | Camk2g | 1.69E-07 | 0.445964119 | 0.004202149 | 0.683 | 0.202 |
| **1251** | Sun1 | 1.69E-07 | 0.413340939 | 0.004213844 | 0.548 | 0.135 |
| **1252** | Mitd1 | 1.70E-07 | 0.636145278 | 0.00422694 | 0.337 | 0.038 |
| **1253** | Etv3 | 1.70E-07 | 0.937785574 | 0.004235111 | 0.385 | 0.058 |
| **1254** | Meis1 | 1.71E-07 | 0.83043253 | 0.004249447 | 0.567 | 0.163 |
| **1255** | Reep3 | 1.71E-07 | 0.227646806 | 0.004249668 | 0.779 | 0.25 |
| **1256** | Stx18 | 1.72E-07 | 0.148842382 | 0.004268773 | 0.538 | 0.125 |
| **1257** | Dhrs7 | 1.72E-07 | 1.337306902 | 0.004270948 | 0.356 | 0.048 |
| **1258** | Xbp1 | 1.73E-07 | 1.166427522 | 0.004293876 | 0.452 | 0.096 |
| **1259** | C3 | 1.75E-07 | 1.291186507 | 0.004360978 | 0.558 | 0.163 |
| **1260** | Pwwp3a | 1.77E-07 | 0.10063492 | 0.004392208 | 0.452 | 0.087 |
| **1261** | Mrps21 | 1.78E-07 | 2.586746254 | 0.004440399 | 0.308 | 0.029 |
| **1262** | Lmna | 1.79E-07 | 1.540151346 | 0.004465294 | 0.337 | 0.038 |
| **1263** | Eif2s3x | 1.80E-07 | 0.954792262 | 0.00447835 | 0.442 | 0.087 |
| **1264** | Adcy3 | 1.81E-07 | 1.867179619 | 0.004504547 | 0.356 | 0.048 |
| **1265** | Gigyf1 | 1.82E-07 | 0.92427841 | 0.004537211 | 0.423 | 0.077 |
| **1266** | Lmod1 | 1.83E-07 | 3.698390022 | 0.004549987 | 0.26 | 0.01 |
| **1267** | C130074G19R | 1.88E-07 | 0.484837914 | 0.004671255 | 0.548 | 0.135 |
| **1268** | Gbp9 | 1.89E-07 | 0.672326049 | 0.004698808 | 0.442 | 0.087 |

| | | | | | | |
|---|---|---|---|---|---|---|
| **1269** | Sdhb | 1.89E-07 | 1.619032586 | 0.004701908 | 0.308 | 0.029 |
| **1270** | Col4a1 | 1.90E-07 | 0.5409399 | 0.004722444 | 0.981 | 0.663 |
| **1271** | Ntrk2 | 1.91E-07 | 1.317154077 | 0.00474909 | 0.471 | 0.106 |
| **1272** | Atp6v1f | 1.95E-07 | 3.053247418 | 0.004844895 | 0.26 | 0.01 |
| **1273** | N4bp2l2 | 1.95E-07 | 0.620484901 | 0.004860015 | 0.76 | 0.25 |
| **1274** | Uhmk1 | 1.96E-07 | 1.238035753 | 0.004873131 | 0.452 | 0.096 |
| **1275** | Pbdc1 | 1.98E-07 | 0.519396253 | 0.004921585 | 0.356 | 0.048 |
| **1276** | Pot1a | 1.99E-07 | 1.530378653 | 0.004962732 | 0.375 | 0.058 |
| **1277** | Nln | 2.00E-07 | 1.56725784 | 0.004978275 | 0.308 | 0.029 |
| **1278** | Tma7 | 2.00E-07 | 1.834757052 | 0.004978275 | 0.308 | 0.029 |
| **1279** | Vamp8 | 2.00E-07 | 1.962688977 | 0.004978275 | 0.308 | 0.029 |
| **1280** | Rpn2 | 2.01E-07 | 1.190596652 | 0.005009125 | 0.442 | 0.087 |
| **1281** | Mob4 | 2.02E-07 | 0.26031224 | 0.005029347 | 0.452 | 0.096 |
| **1282** | Msn | 2.03E-07 | 0.562870591 | 0.00504069 | 0.885 | 0.337 |
| **1283** | Mplkip | 2.03E-07 | 3.106149863 | 0.005051675 | 0.26 | 0.01 |
| **1284** | Armcx1 | 2.03E-07 | 2.129212258 | 0.005059786 | 0.346 | 0.048 |
| **1285** | Eif4ebp2 | 2.08E-07 | 0.542979675 | 0.005180613 | 0.635 | 0.173 |
| **1286** | Nxf1 | 2.10E-07 | 0.272331625 | 0.005226061 | 0.49 | 0.106 |
| **1287** | Mtss2 | 2.11E-07 | 1.412209474 | 0.005242946 | 0.413 | 0.077 |
| **1288** | Mcl1 | 2.12E-07 | 0.798818894 | 0.005263833 | 0.587 | 0.163 |
| **1289** | Vbp1 | 2.12E-07 | 1.469370287 | 0.005270312 | 0.308 | 0.029 |
| **1290** | Metap2 | 2.12E-07 | 0.539457281 | 0.005280437 | 0.519 | 0.125 |
| **1291** | Hsd17b11 | 2.15E-07 | 0.532905396 | 0.005349661 | 0.587 | 0.154 |
| **1292** | Cerk | 2.15E-07 | 0.643431047 | 0.005359726 | 0.519 | 0.125 |
| **1293** | Oip5os1 | 2.18E-07 | 0.512780588 | 0.005416778 | 0.75 | 0.25 |
| **1294** | Ptpn9 | 2.18E-07 | 0.410308864 | 0.005422496 | 0.663 | 0.202 |
| **1295** | Xpr1 | 2.18E-07 | 0.291132653 | 0.005426636 | 0.721 | 0.231 |
| **1296** | Ndufb8 | 2.20E-07 | 2.028299951 | 0.005465294 | 0.327 | 0.038 |
| **1297** | Anapc2 | 2.20E-07 | 0.978685798 | 0.005471415 | 0.356 | 0.048 |
| **1298** | Ctsz | 2.20E-07 | 1.606426709 | 0.005474128 | 0.308 | 0.029 |
| **1299** | Ndufb11 | 2.20E-07 | 1.161423271 | 0.005474128 | 0.308 | 0.029 |
| **1300** | 2410006H16Ri | 2.21E-07 | 4.910459636 | 0.005487889 | 0.231 | 0 |
| **1301** | Eva1b | 2.21E-07 | 4.67872625 | 0.005487889 | 0.231 | 0 |
| **1302** | Mrpl57 | 2.21E-07 | 4.934681161 | 0.005487889 | 0.231 | 0 |
| **1303** | Nop10 | 2.21E-07 | 4.7662925 | 0.005487889 | 0.231 | 0 |
| **1304** | Scarf2 | 2.21E-07 | 4.617949729 | 0.005487889 | 0.231 | 0 |
| **1305** | Sspn | 2.21E-07 | 4.921898842 | 0.005487889 | 0.231 | 0 |
| **1306** | Timm10b | 2.21E-07 | 4.764429567 | 0.005487889 | 0.231 | 0 |
| **1307** | Vkorc1 | 2.21E-07 | 4.757873236 | 0.005487889 | 0.231 | 0 |
| **1308** | Cdk6 | 2.21E-07 | 0.006104639 | 0.005497483 | 0.625 | 0.163 |
| **1309** | Dpf3 | 2.24E-07 | 0.764152403 | 0.005568644 | 0.356 | 0.048 |

| | | | | | | |
|---|---|---|---|---|---|---|
| **1310** | Mrc2 | 2.26E-07 | 2.15601581 | 0.005630167 | 0.346 | 0.048 |
| **1311** | Smc1a | 2.28E-07 | 0.38850888 | 0.005665491 | 0.673 | 0.202 |
| **1312** | Nap1l1 | 2.30E-07 | 0.228938024 | 0.005712642 | 0.567 | 0.144 |
| **1313** | Kmt2d | 2.35E-07 | 0.759529242 | 0.00584413 | 0.558 | 0.144 |
| **1314** | Cpe | 2.37E-07 | 1.126674549 | 0.005891809 | 0.375 | 0.058 |
| **1315** | Tmem167 | 2.38E-07 | 0.568086253 | 0.005925172 | 0.51 | 0.115 |
| **1316** | Hcfc1 | 2.42E-07 | 1.707876186 | 0.006017547 | 0.308 | 0.029 |
| **1317** | Hdac9 | 2.42E-07 | 0.61528628 | 0.006019553 | 0.731 | 0.269 |
| **1318** | Clic5 | 2.43E-07 | 0.396830913 | 0.00604072 | 0.577 | 0.154 |
| **1319** | Gabpb2 | 2.43E-07 | 0.337098986 | 0.006045671 | 0.538 | 0.135 |
| **1320** | Uqcrfs1 | 2.46E-07 | 1.037854742 | 0.006132317 | 0.308 | 0.029 |
| **1321** | Jag1 | 2.49E-07 | 0.045281378 | 0.006195837 | 0.654 | 0.192 |
| **1322** | Man1a2 | 2.51E-07 | 0.626692006 | 0.006256257 | 0.702 | 0.221 |
| **1323** | Lman2 | 2.52E-07 | 0.843377968 | 0.006258596 | 0.394 | 0.067 |
| **1324** | Prune2 | 2.52E-07 | 1.293743296 | 0.006273414 | 0.442 | 0.096 |
| **1325** | Yy1 | 2.57E-07 | 0.07847032 | 0.006398654 | 0.76 | 0.231 |
| **1326** | Vps13a | 2.60E-07 | 0.249565408 | 0.006464519 | 0.798 | 0.25 |
| **1327** | Pgap2 | 2.60E-07 | 0.855809778 | 0.006470552 | 0.394 | 0.067 |
| **1328** | Ftx | 2.61E-07 | 0.742367326 | 0.006482933 | 0.413 | 0.077 |
| **1329** | Megf8 | 2.64E-07 | 1.6973436 | 0.006561458 | 0.327 | 0.038 |
| **1330** | Spaar | 2.67E-07 | 0.17809768 | 0.006647162 | 0.721 | 0.212 |
| **1331** | Sh3rf1 | 2.68E-07 | 0.13812501 | 0.006678879 | 0.933 | 0.375 |
| **1332** | Fasn | 2.71E-07 | 0.841211029 | 0.006753995 | 0.375 | 0.058 |
| **1333** | Nbr1 | 2.73E-07 | 0.198456362 | 0.006790752 | 0.625 | 0.183 |
| **1334** | Kdelr1 | 2.77E-07 | 1.844304795 | 0.006886604 | 0.365 | 0.058 |
| **1335** | Pdcd6 | 2.77E-07 | 1.907117424 | 0.006895723 | 0.298 | 0.029 |
| **1336** | Fundc2 | 2.78E-07 | 2.315061798 | 0.006912436 | 0.279 | 0.019 |
| **1337** | Flii | 2.78E-07 | 0.785744935 | 0.006918411 | 0.413 | 0.077 |
| **1338** | Grrp1 | 2.88E-07 | 2.389703244 | 0.007163167 | 0.298 | 0.029 |
| **1339** | Thbs1 | 2.91E-07 | 0.847572191 | 0.007229587 | 0.375 | 0.058 |
| **1340** | Tsn | 2.92E-07 | 1.133770602 | 0.007263422 | 0.413 | 0.077 |
| **1341** | Cap1 | 2.92E-07 | 0.56126842 | 0.00726896 | 0.5 | 0.115 |
| **1342** | Desi2 | 2.96E-07 | 0.652175573 | 0.007354772 | 0.596 | 0.163 |
| **1343** | 2610507B11Ri | 2.98E-07 | 0.519867557 | 0.007415358 | 0.587 | 0.154 |
| **1344** | Mecp2 | 3.00E-07 | 0.424180078 | 0.007456245 | 0.788 | 0.26 |
| **1345** | Socs4 | 3.01E-07 | 0.883627582 | 0.007479353 | 0.375 | 0.058 |
| **1346** | Pttg1 | 3.01E-07 | 2.35956325 | 0.007480928 | 0.279 | 0.019 |
| **1347** | Zdhhc20 | 3.05E-07 | 0.803001827 | 0.007592558 | 0.702 | 0.231 |
| **1348** | Ikbkb | 3.05E-07 | 0.605520078 | 0.007601139 | 0.481 | 0.106 |
| **1349** | Fam92a | 3.07E-07 | 2.007722438 | 0.007629956 | 0.279 | 0.019 |
| **1350** | Spcs3 | 3.07E-07 | 1.51481416 | 0.007629956 | 0.279 | 0.019 |

| | | | | | | |
|---|---|---|---|---|---|---|
| **1351** | Fam124a | 3.11E-07 | 1.207860636 | 0.00774937 | 0.413 | 0.077 |
| **1352** | BC005624 | 3.13E-07 | 1.304914474 | 0.007781848 | 0.279 | 0.019 |
| **1353** | Tmem71 | 3.13E-07 | 2.31683299 | 0.007781848 | 0.279 | 0.019 |
| **1354** | Ncbp1 | 3.15E-07 | 1.585016102 | 0.00783521 | 0.385 | 0.067 |
| **1355** | Srsf9 | 3.17E-07 | 2.218087311 | 0.007876332 | 0.298 | 0.029 |
| **1356** | Lsm7 | 3.19E-07 | 1.808443268 | 0.007936658 | 0.279 | 0.019 |
| **1357** | Zmym4 | 3.22E-07 | 0.207540148 | 0.008006193 | 0.808 | 0.26 |
| **1358** | Snhg20 | 3.22E-07 | 1.569202047 | 0.00801231 | 0.327 | 0.038 |
| **1359** | Zeb2 | 3.30E-07 | 0.502801194 | 0.00822275 | 0.971 | 0.692 |
| **1360** | Orc4 | 3.33E-07 | 0.08188926 | 0.008280419 | 0.606 | 0.163 |
| **1361** | Tob1 | 3.36E-07 | 1.315257839 | 0.008351235 | 0.423 | 0.087 |
| **1362** | Atp6v0a2 | 3.38E-07 | 0.252966048 | 0.00840107 | 0.413 | 0.077 |
| **1363** | Ccz1 | 3.38E-07 | 0.968390176 | 0.00840107 | 0.413 | 0.077 |
| **1364** | Mef2c | 3.39E-07 | 0.43208102 | 0.00844526 | 0.952 | 0.462 |
| **1365** | Wdr82 | 3.40E-07 | 0.583943988 | 0.008452181 | 0.481 | 0.106 |
| **1366** | Vps29 | 3.48E-07 | 1.321890363 | 0.008657816 | 0.298 | 0.029 |
| **1367** | Sptssa | 3.55E-07 | 1.124641591 | 0.008822853 | 0.298 | 0.029 |
| **1368** | Pdcd5 | 3.56E-07 | 3.505927279 | 0.008850942 | 0.25 | 0.01 |
| **1369** | Mkrn1 | 3.61E-07 | 1.592999215 | 0.008990925 | 0.298 | 0.029 |
| **1370** | Morf4l2 | 3.63E-07 | 0.892674849 | 0.009033623 | 0.49 | 0.115 |
| **1371** | Yme1l1 | 3.63E-07 | 0.306831774 | 0.009040238 | 0.635 | 0.183 |
| **1372** | Ehmt2 | 3.64E-07 | 0.990734952 | 0.009051983 | 0.365 | 0.058 |
| **1373** | Derl2 | 3.64E-07 | 1.595201035 | 0.009061705 | 0.346 | 0.048 |
| **1374** | Uba3 | 3.77E-07 | 0.840688671 | 0.009383956 | 0.346 | 0.048 |
| **1375** | Atp5d | 3.79E-07 | 3.135258063 | 0.00942048 | 0.25 | 0.01 |
| **1376** | Ccnd1 | 3.79E-07 | 3.064174903 | 0.00942048 | 0.25 | 0.01 |
| **1377** | Rab11b | 3.80E-07 | 0.664264838 | 0.009442767 | 0.625 | 0.183 |
| **1378** | Tns2 | 3.81E-07 | 0.612995568 | 0.009475284 | 0.51 | 0.125 |
| **1379** | Cdv3 | 3.87E-07 | 0.907672725 | 0.009619294 | 0.433 | 0.087 |
| **1380** | Glg1 | 3.88E-07 | 0.608311757 | 0.009651071 | 0.808 | 0.288 |
| **1381** | Mex3c | 3.91E-07 | 1.290297924 | 0.009722863 | 0.385 | 0.067 |
| **1382** | Snrpb | 3.96E-07 | 0.865267878 | 0.009854161 | 0.365 | 0.058 |
| **1383** | Plpp6 | 3.97E-07 | 2.13665575 | 0.00987871 | 0.298 | 0.029 |
| **1384** | 8-Sep | 4.02E-07 | 0.684871079 | 0.009995 | 0.596 | 0.163 |
| **1385** | B3galnt2 | 4.04E-07 | 0.931148444 | 0.010061959 | 0.346 | 0.048 |
| **1386** | Med17 | 4.04E-07 | 1.033685869 | 0.010061959 | 0.346 | 0.048 |
| **1387** | Ephx1 | 4.07E-07 | 0.710385146 | 0.010116482 | 0.5 | 0.125 |
| **1388** | mt-Nd2 | 4.07E-07 | 0.686700676 | 0.010119092 | 0.99 | 0.779 |
| **1389** | Psmd1 | 4.11E-07 | 0.263096187 | 0.010234445 | 0.606 | 0.173 |
| **1390** | Ptp4a3 | 4.11E-07 | 0.170458513 | 0.010235561 | 0.5 | 0.115 |
| **1391** | Cldn15 | 4.12E-07 | 0.949285784 | 0.010242909 | 0.433 | 0.087 |

| | | | | | | |
|---|---|---|---|---|---|---|
| **1392** | AU020206 | 4.12E-07 | 0.122723141 | 0.01024653 | 0.644 | 0.192 |
| **1393** | Abhd16a | 4.12E-07 | 1.696947572 | 0.010257013 | 0.298 | 0.029 |
| **1394** | Kdsr | 4.12E-07 | 1.769905891 | 0.010257013 | 0.298 | 0.029 |
| **1395** | Sesn3 | 4.15E-07 | 1.753098357 | 0.010332821 | 0.317 | 0.038 |
| **1396** | Sec24a | 4.16E-07 | 0.396450486 | 0.010338147 | 0.538 | 0.135 |
| **1397** | Ank | 4.18E-07 | 0.328565579 | 0.010388052 | 0.5 | 0.115 |
| **1398** | Aebp2 | 4.19E-07 | 0.430132803 | 0.010416319 | 0.567 | 0.154 |
| **1399** | Mrpl52 | 4.21E-07 | 5.081411819 | 0.010469712 | 0.221 | 0 |
| **1400** | Myom1 | 4.21E-07 | 5.298525885 | 0.010469712 | 0.221 | 0 |
| **1401** | Prdx5 | 4.21E-07 | 4.630173558 | 0.010469712 | 0.221 | 0 |
| **1402** | Snhg8 | 4.21E-07 | 4.85909723 | 0.010469712 | 0.221 | 0 |
| **1403** | Dlat | 4.24E-07 | 1.125225927 | 0.010544879 | 0.365 | 0.058 |
| **1404** | Arpp19 | 4.26E-07 | 1.091455142 | 0.010590178 | 0.462 | 0.106 |
| **1405** | Ndufs2 | 4.26E-07 | 1.077294386 | 0.010601234 | 0.346 | 0.048 |
| **1406** | Sec23a | 4.27E-07 | -0.1172274 | 0.010618261 | 0.452 | 0.096 |
| **1407** | Gnl3l | 4.28E-07 | 1.615303714 | 0.010649275 | 0.298 | 0.029 |
| **1408** | Polr2m | 4.28E-07 | 0.563923846 | 0.010649275 | 0.298 | 0.029 |
| **1409** | Lpcat1 | 4.30E-07 | 2.055281751 | 0.010704389 | 0.269 | 0.019 |
| **1410** | Pea15a | 4.36E-07 | 0.6794257 | 0.010851692 | 0.471 | 0.106 |
| **1411** | Hmg20b | 4.38E-07 | 0.388431243 | 0.01090992 | 0.317 | 0.038 |
| **1412** | Ctsa | 4.39E-07 | 3.007438958 | 0.01091793 | 0.269 | 0.019 |
| **1413** | Paip1 | 4.43E-07 | 0.576899928 | 0.011016599 | 0.471 | 0.106 |
| **1414** | Pcdh1 | 4.43E-07 | 0.400044922 | 0.011022687 | 0.673 | 0.212 |
| **1415** | Phc2 | 4.45E-07 | 0.691267698 | 0.011063267 | 0.548 | 0.144 |
| **1416** | Nmi | 4.47E-07 | 0.547474404 | 0.0111312 | 0.519 | 0.135 |
| **1417** | Ier2 | 4.48E-07 | 1.766284853 | 0.011135578 | 0.269 | 0.019 |
| **1418** | Zkscan1 | 4.53E-07 | 1.058273033 | 0.011264926 | 0.298 | 0.029 |
| **1419** | Mark1 | 4.53E-07 | 1.650536952 | 0.011282193 | 0.365 | 0.058 |
| **1420** | Fyttd1 | 4.54E-07 | 0.420328909 | 0.011294801 | 0.51 | 0.125 |
| **1421** | Dhx9 | 4.57E-07 | 0.70102356 | 0.011378266 | 0.5 | 0.125 |
| **1422** | Gsr | 4.58E-07 | 0.315964512 | 0.01138876 | 0.404 | 0.077 |
| **1423** | Tgs1 | 4.68E-07 | 0.820897158 | 0.011642208 | 0.452 | 0.096 |
| **1424** | Hdgf | 4.69E-07 | 1.146419652 | 0.011669257 | 0.365 | 0.058 |
| **1425** | Smarce1 | 4.70E-07 | 0.823009231 | 0.011699384 | 0.442 | 0.096 |
| **1426** | Ptbp1 | 4.75E-07 | 2.527182483 | 0.011813927 | 0.269 | 0.019 |
| **1427** | Edem1 | 4.78E-07 | 1.207905382 | 0.01190193 | 0.394 | 0.077 |
| **1428** | Pabpn1 | 4.80E-07 | 0.307434281 | 0.011937614 | 0.558 | 0.144 |
| **1429** | Tmem127 | 4.84E-07 | 0.915145237 | 0.012045763 | 0.385 | 0.067 |
| **1430** | Cul3 | 4.91E-07 | 0.392456183 | 0.012228451 | 0.875 | 0.327 |
| **1431** | Atn1 | 4.96E-07 | 0.716118581 | 0.012340353 | 0.49 | 0.115 |
| **1432** | Sec61b | 5.04E-07 | 2.288394187 | 0.012532046 | 0.269 | 0.019 |

| | | | | | | |
|---|---|---|---|---|---|---|
| **1433** | Ifi27 | 5.12E-07 | 1.732606648 | 0.012729981 | 0.356 | 0.058 |
| **1434** | Dpt | 5.13E-07 | 1.817825841 | 0.012773101 | 0.337 | 0.048 |
| **1435** | Pigp | 5.14E-07 | 2.7050104 | 0.01278064 | 0.269 | 0.019 |
| **1436** | Supt5 | 5.20E-07 | 0.869390251 | 0.01294758 | 0.433 | 0.087 |
| **1437** | Nufip2 | 5.22E-07 | 0.623661908 | 0.012976596 | 0.779 | 0.279 |
| **1438** | Tmem39a | 5.25E-07 | 1.68063037 | 0.013066845 | 0.317 | 0.038 |
| **1439** | Naa20 | 5.28E-07 | 1.034803433 | 0.013127419 | 0.365 | 0.058 |
| **1440** | Cd164 | 5.29E-07 | 1.35060577 | 0.013156465 | 0.404 | 0.077 |
| **1441** | Tollip | 5.32E-07 | 1.327108475 | 0.013226261 | 0.337 | 0.048 |
| **1442** | Rnf114 | 5.37E-07 | 0.611499346 | 0.013349562 | 0.365 | 0.058 |
| **1443** | Fbxw8 | 5.37E-07 | 0.261802296 | 0.013355767 | 0.692 | 0.212 |
| **1444** | Egln1 | 5.38E-07 | 0.344760746 | 0.013388461 | 0.635 | 0.183 |
| **1445** | P2rx4 | 5.44E-07 | 1.56946634 | 0.013545042 | 0.317 | 0.038 |
| **1446** | Rbp4 | 5.45E-07 | 1.606440099 | 0.013555284 | 0.269 | 0.019 |
| **1447** | Hexim1 | 5.46E-07 | 1.080575589 | 0.013575327 | 0.365 | 0.058 |
| **1448** | Lactb2 | 5.54E-07 | 1.683919876 | 0.013772735 | 0.308 | 0.038 |
| **1449** | Creb3 | 5.56E-07 | 2.4483281 | 0.013823415 | 0.269 | 0.019 |
| **1450** | Gm15581 | 5.56E-07 | 2.726020247 | 0.013823415 | 0.269 | 0.019 |
| **1451** | Pcolce | 5.56E-07 | 2.298902924 | 0.013823415 | 0.269 | 0.019 |
| **1452** | Trim30d | 5.57E-07 | 1.555488487 | 0.013854426 | 0.356 | 0.058 |
| **1453** | Tmbim1 | 5.59E-07 | 0.224831014 | 0.013896713 | 0.519 | 0.125 |
| **1454** | Jup | 5.64E-07 | 0.512644731 | 0.014025763 | 0.615 | 0.183 |
| **1455** | Trp53bp2 | 5.64E-07 | 0.296265562 | 0.014037951 | 0.365 | 0.058 |
| **1456** | Ipo9 | 5.66E-07 | -0.07922473 | 0.014090577 | 0.49 | 0.115 |
| **1457** | D16Ertd472e | 5.67E-07 | 1.748027736 | 0.014096655 | 0.269 | 0.019 |
| **1458** | Scaf11 | 5.69E-07 | 0.468829115 | 0.014154994 | 0.837 | 0.298 |
| **1459** | Igkc | 5.71E-07 | 4.16157948 | 0.014205612 | 0.24 | 0.01 |
| **1460** | Cul5 | 5.71E-07 | 0.198428827 | 0.014218886 | 0.471 | 0.106 |
| **1461** | Serp1 | 5.74E-07 | 1.113679822 | 0.014294109 | 0.317 | 0.038 |
| **1462** | Anapc7 | 5.78E-07 | 2.386262384 | 0.014375099 | 0.269 | 0.019 |
| **1463** | Armc8 | 5.81E-07 | 0.461812892 | 0.014453809 | 0.731 | 0.231 |
| **1464** | Zc2hc1a | 5.81E-07 | 0.195273164 | 0.014460624 | 0.529 | 0.144 |
| **1465** | Pnkd | 5.83E-07 | 0.590986106 | 0.014504751 | 0.442 | 0.096 |
| **1466** | Slc43a3 | 5.84E-07 | 0.267986749 | 0.014529532 | 0.74 | 0.24 |
| **1467** | Plekha3 | 5.85E-07 | 1.512807397 | 0.014552557 | 0.317 | 0.038 |
| **1468** | Psmd2 | 5.85E-07 | 1.414297036 | 0.014552557 | 0.317 | 0.038 |
| **1469** | Ostm1 | 5.89E-07 | 1.956004178 | 0.014658841 | 0.269 | 0.019 |
| **1470** | Psmc5 | 5.89E-07 | 1.820352678 | 0.014658841 | 0.269 | 0.019 |
| **1471** | Pde1a | 5.95E-07 | 0.901961925 | 0.014802731 | 0.596 | 0.183 |
| **1472** | H2-T22 | 5.95E-07 | 0.989257442 | 0.014815507 | 0.317 | 0.038 |
| **1473** | Ap2s1 | 6.01E-07 | 1.098640327 | 0.014947979 | 0.269 | 0.019 |

| | | | | | | |
|---|---|---|---|---|---|---|
| **1474** | Prelid1 | 6.01E-07 | 1.31551711 | 0.014947979 | 0.269 | 0.019 |
| **1475** | Ncbp3 | 6.01E-07 | 0.583094438 | 0.014952994 | 0.423 | 0.087 |
| **1476** | Scamp1 | 6.03E-07 | 0.481628147 | 0.015013249 | 0.587 | 0.163 |
| **1477** | Arl8a | 6.06E-07 | 1.343977191 | 0.015083033 | 0.317 | 0.038 |
| **1478** | Serinc5 | 6.10E-07 | 0.532671055 | 0.015187292 | 0.404 | 0.077 |
| **1479** | Cacnb2 | 6.10E-07 | 1.328340275 | 0.015187737 | 0.413 | 0.096 |
| **1480** | Grina | 6.22E-07 | 1.103593523 | 0.015464539 | 0.337 | 0.048 |
| **1481** | Arhgef6 | 6.23E-07 | 0.244302119 | 0.015508585 | 0.558 | 0.154 |
| **1482** | Xaf1 | 6.25E-07 | 0.486989871 | 0.01555202 | 0.452 | 0.096 |
| **1483** | Hspe1 | 6.37E-07 | 1.840341527 | 0.015850026 | 0.288 | 0.029 |
| **1484** | Mapre1 | 6.44E-07 | -0.05983841 | 0.016029168 | 0.558 | 0.144 |
| **1485** | Oat | 6.47E-07 | 3.486270975 | 0.016089567 | 0.24 | 0.01 |
| **1486** | Pds5b | 6.56E-07 | 0.315079389 | 0.016318883 | 0.846 | 0.308 |
| **1487** | Chchd1 | 6.60E-07 | 3.614141421 | 0.016426112 | 0.24 | 0.01 |
| **1488** | Ate1 | 6.74E-07 | 3.521555094 | 0.016769436 | 0.24 | 0.01 |
| **1489** | Mybbp1a | 6.74E-07 | 3.395663166 | 0.016769436 | 0.24 | 0.01 |
| **1490** | Smim15 | 6.74E-07 | 3.487314068 | 0.016769436 | 0.24 | 0.01 |
| **1491** | Esam | 6.75E-07 | 0.547920793 | 0.016805332 | 0.865 | 0.317 |
| **1492** | Mbd1 | 6.78E-07 | 1.477513676 | 0.016861421 | 0.337 | 0.048 |
| **1493** | Sh2d3c | 6.84E-07 | 0.360557757 | 0.017028759 | 0.596 | 0.173 |
| **1494** | Atp6v1a | 6.88E-07 | 0.810613719 | 0.017119862 | 0.519 | 0.135 |
| **1495** | Cdc37 | 6.91E-07 | 0.510018401 | 0.017197905 | 0.423 | 0.087 |
| **1496** | Ldlrap1 | 6.99E-07 | 2.148583184 | 0.017402363 | 0.288 | 0.029 |
| **1497** | Lin7c | 7.01E-07 | 0.935883067 | 0.017453562 | 0.337 | 0.048 |
| **1498** | Arhgef25 | 7.02E-07 | 3.188883273 | 0.017476946 | 0.24 | 0.01 |
| **1499** | Ide | 7.05E-07 | 0.447937802 | 0.017551884 | 0.49 | 0.115 |
| **1500** | Mia3 | 7.08E-07 | 0.727162603 | 0.017614141 | 0.519 | 0.135 |
| **1501** | Mta2 | 7.13E-07 | 2.220454425 | 0.017729942 | 0.288 | 0.029 |
| **1502** | Vcl | 7.13E-07 | 0.276293978 | 0.017745491 | 0.885 | 0.375 |
| **1503** | Cdc26 | 7.17E-07 | 1.292772603 | 0.017832122 | 0.356 | 0.058 |
| **1504** | Ythdf1 | 7.17E-07 | 1.164508218 | 0.017832122 | 0.356 | 0.058 |
| **1505** | Mrps33 | 7.17E-07 | 2.819085455 | 0.017841401 | 0.24 | 0.01 |
| **1506** | Fkbp1a | 7.19E-07 | 0.706504179 | 0.01789461 | 0.606 | 0.183 |
| **1507** | Mios | 7.26E-07 | 1.092381318 | 0.018065714 | 0.337 | 0.048 |
| **1508** | Casc4 | 7.32E-07 | 1.931172511 | 0.018213173 | 0.24 | 0.01 |
| **1509** | Cited2 | 7.32E-07 | 1.967178522 | 0.018213173 | 0.24 | 0.01 |
| **1510** | Cope | 7.32E-07 | 1.95984885 | 0.018213173 | 0.24 | 0.01 |
| **1511** | Fam217b | 7.32E-07 | 1.718149781 | 0.018213173 | 0.24 | 0.01 |
| **1512** | Fhl5 | 7.32E-07 | 2.353512574 | 0.018213173 | 0.24 | 0.01 |
| **1513** | Hgf | 7.32E-07 | 1.783910708 | 0.018213173 | 0.24 | 0.01 |
| **1514** | Pbx2 | 7.32E-07 | 2.963924061 | 0.018213173 | 0.24 | 0.01 |

| | | | | | | |
|---|---|---|---|---|---|---|
| **1515** | Rpl7l1 | 7.32E-07 | 3.120927086 | 0.018213173 | 0.24 | 0.01 |
| **1516** | Selenos | 7.32E-07 | 1.417334519 | 0.018213173 | 0.24 | 0.01 |
| **1517** | Ston1 | 7.34E-07 | 0.450599735 | 0.018267402 | 0.538 | 0.144 |
| **1518** | Dync1i2 | 7.34E-07 | 0.33378329 | 0.018270031 | 0.692 | 0.221 |
| **1519** | Id1 | 7.37E-07 | 1.298413302 | 0.018338644 | 0.413 | 0.087 |
| **1520** | Adcy6 | 7.38E-07 | 0.505444049 | 0.018364863 | 0.548 | 0.144 |
| **1521** | Ttc14 | 7.42E-07 | 0.330568892 | 0.018455308 | 0.712 | 0.221 |
| **1522** | Retreg3 | 7.47E-07 | 0.978678678 | 0.018591325 | 0.433 | 0.096 |
| **1523** | Coro6 | 7.52E-07 | 0.938100364 | 0.018698524 | 0.337 | 0.048 |
| **1524** | Kdelr2 | 7.52E-07 | 0.90422653 | 0.018698524 | 0.337 | 0.048 |
| **1525** | Zc3h13 | 7.65E-07 | 0.217883085 | 0.019027372 | 0.731 | 0.24 |
| **1526** | Filip1l | 7.71E-07 | 0.830863056 | 0.019176738 | 0.808 | 0.337 |
| **1527** | Rps6kb1 | 7.78E-07 | 0.2966302 | 0.019361959 | 0.712 | 0.231 |
| **1528** | Ogn | 7.80E-07 | 1.82385345 | 0.019402735 | 0.308 | 0.038 |
| **1529** | Rcsd1 | 7.80E-07 | 0.113240642 | 0.019408102 | 0.798 | 0.26 |
| **1530** | Sgta | 7.82E-07 | 1.741290576 | 0.019458986 | 0.288 | 0.029 |
| **1531** | Mtmr1 | 7.86E-07 | 0.848142428 | 0.019566591 | 0.404 | 0.077 |
| **1532** | Psmb9 | 7.92E-07 | 2.474582781 | 0.019711066 | 0.26 | 0.019 |
| **1533** | Rbp7 | 7.96E-07 | 1.482732535 | 0.019807744 | 0.404 | 0.087 |
| **1534** | Yipf4 | 7.97E-07 | 0.86563402 | 0.01982377 | 0.288 | 0.029 |
| **1535** | Tmx3 | 7.98E-07 | 0.168388579 | 0.019858753 | 0.577 | 0.163 |
| **1536** | Abca8a | 7.98E-07 | 0.890555782 | 0.019859473 | 0.375 | 0.067 |
| **1537** | Cyb5r4 | 7.98E-07 | 0.635885563 | 0.019859473 | 0.375 | 0.067 |
| **1538** | Bloc1s1 | 7.99E-07 | 5.005278692 | 0.019871317 | 0.212 | 0 |
| **1539** | Cdc34 | 7.99E-07 | 4.725370956 | 0.019871317 | 0.212 | 0 |
| **1540** | Cdk4 | 7.99E-07 | 4.596643534 | 0.019871317 | 0.212 | 0 |
| **1541** | Evi5l | 7.99E-07 | 4.867760226 | 0.019871317 | 0.212 | 0 |
| **1542** | Fam219b | 7.99E-07 | 4.482225546 | 0.019871317 | 0.212 | 0 |
| **1543** | Hoxb8 | 7.99E-07 | 4.73077739 | 0.019871317 | 0.212 | 0 |
| **1544** | Ptpn18 | 7.99E-07 | 4.567905269 | 0.019871317 | 0.212 | 0 |
| **1545** | Slc7a2 | 7.99E-07 | 5.126201395 | 0.019871317 | 0.212 | 0 |
| **1546** | Smpdl3a | 7.99E-07 | 4.747463203 | 0.019871317 | 0.212 | 0 |
| **1547** | Trmt9b | 7.99E-07 | 5.006826321 | 0.019871317 | 0.212 | 0 |
| **1548** | Larp1 | 8.05E-07 | 0.311696225 | 0.020027177 | 0.663 | 0.221 |
| **1549** | mt-Nd5 | 8.10E-07 | 0.754512557 | 0.020152471 | 1 | 0.644 |
| **1550** | Cox7b | 8.12E-07 | 2.016835958 | 0.020195136 | 0.288 | 0.029 |
| **1551** | Ddost | 8.12E-07 | 1.111935851 | 0.020195136 | 0.288 | 0.029 |
| **1552** | Trp53inp1 | 8.12E-07 | 1.619020165 | 0.020195136 | 0.288 | 0.029 |
| **1553** | Pik3ca | 8.13E-07 | 0.441336086 | 0.020233655 | 0.865 | 0.327 |
| **1554** | Flt4 | 8.19E-07 | -0.47372924 | 0.020375365 | 0.337 | 0.048 |
| **1555** | Mcrip1 | 8.19E-07 | 1.04079735 | 0.020375365 | 0.337 | 0.048 |

| | | | | | | |
|---|---|---|---|---|---|---|
| **1556** | Map1b | 8.22E-07 | 0.599718939 | 0.020453186 | 0.51 | 0.135 |
| **1557** | E130307A14R | 8.22E-07 | 0.968745765 | 0.020455117 | 0.462 | 0.115 |
| **1558** | Etf1 | 8.24E-07 | 1.11181378 | 0.0205134 | 0.394 | 0.077 |
| **1559** | Brap | 8.27E-07 | 1.80769187 | 0.020573198 | 0.288 | 0.029 |
| **1560** | Uba2 | 8.29E-07 | 1.130656493 | 0.020634053 | 0.452 | 0.106 |
| **1561** | Glul | 8.35E-07 | 0.833184386 | 0.020785359 | 0.49 | 0.125 |
| **1562** | U2af1 | 8.38E-07 | 0.501859357 | 0.020838437 | 0.404 | 0.077 |
| **1563** | Eif4b | 8.41E-07 | 0.760779795 | 0.020921913 | 0.5 | 0.125 |
| **1564** | Leprot | 8.48E-07 | 0.976975757 | 0.021090289 | 0.413 | 0.087 |
| **1565** | Manbal | 8.53E-07 | 0.995314382 | 0.021218973 | 0.308 | 0.038 |
| **1566** | Cant1 | 8.58E-07 | 1.079975251 | 0.021349875 | 0.288 | 0.029 |
| **1567** | Cdc42se1 | 8.58E-07 | 1.857108686 | 0.021349875 | 0.288 | 0.029 |
| **1568** | Irs2 | 8.58E-07 | 1.773795579 | 0.021349875 | 0.288 | 0.029 |
| **1569** | Pomp | 8.58E-07 | 0.72424184 | 0.021349875 | 0.288 | 0.029 |
| **1570** | Vamp3 | 8.58E-07 | 1.588494262 | 0.021349875 | 0.288 | 0.029 |
| **1571** | Arl8b | 8.73E-07 | 0.582708487 | 0.021711872 | 0.596 | 0.173 |
| **1572** | Gpr137b | 8.74E-07 | 2.896101633 | 0.021737903 | 0.26 | 0.019 |
| **1573** | Poldip3 | 8.76E-07 | 0.825703789 | 0.021785803 | 0.356 | 0.058 |
| **1574** | Scn7a | 8.78E-07 | 1.39324342 | 0.021854739 | 0.394 | 0.077 |
| **1575** | Esf1 | 8.84E-07 | 1.090140418 | 0.021990394 | 0.308 | 0.038 |
| **1576** | Ptar1 | 8.87E-07 | -0.1672941 | 0.022072315 | 0.462 | 0.106 |
| **1577** | Emg1 | 8.91E-07 | 2.810806918 | 0.022166683 | 0.26 | 0.019 |
| **1578** | Mapk12 | 8.92E-07 | 0.839832623 | 0.022203046 | 0.394 | 0.077 |
| **1579** | Rras | 9.04E-07 | 1.363655934 | 0.022486026 | 0.327 | 0.048 |
| **1580** | Acbd3 | 9.05E-07 | 0.503791896 | 0.02252558 | 0.558 | 0.154 |
| **1581** | Krit1 | 9.21E-07 | 0.022780035 | 0.022919741 | 0.615 | 0.173 |
| **1582** | Dlg2 | 9.25E-07 | 0.197093195 | 0.023010188 | 0.462 | 0.115 |
| **1583** | Itgb5 | 9.35E-07 | 0.388854756 | 0.023255834 | 0.548 | 0.144 |
| **1584** | Prr5l | 9.45E-07 | 0.956410422 | 0.023501182 | 0.413 | 0.087 |
| **1585** | Man2a2 | 9.52E-07 | 1.298876457 | 0.023679872 | 0.327 | 0.048 |
| **1586** | Golga1 | 9.59E-07 | 0.350925116 | 0.023866548 | 0.413 | 0.087 |
| **1587** | Zfp62 | 9.59E-07 | 0.820824402 | 0.023866548 | 0.413 | 0.087 |
| **1588** | Ndufc1 | 9.63E-07 | 2.826395325 | 0.023964677 | 0.26 | 0.019 |
| **1589** | Polr2b | 9.66E-07 | 0.347803507 | 0.024038921 | 0.308 | 0.038 |
| **1590** | Tcp1 | 9.67E-07 | 0.874538957 | 0.024066814 | 0.356 | 0.058 |
| **1591** | Nfya | 9.69E-07 | 0.388605347 | 0.02412008 | 0.375 | 0.067 |
| **1592** | Pfkp | 9.70E-07 | 0.192652703 | 0.024141726 | 0.538 | 0.144 |
| **1593** | Psma2 | 9.79E-07 | 1.208849821 | 0.024346102 | 0.385 | 0.077 |
| **1594** | Hcfc1r1 | 1.00E-06 | 2.379196647 | 0.024915532 | 0.26 | 0.019 |
| **1595** | Csf1 | 1.00E-06 | 1.120356628 | 0.024976759 | 0.346 | 0.058 |
| **1596** | Tbc1d23 | 1.01E-06 | 0.545999537 | 0.025160245 | 0.596 | 0.173 |

| | | | | | | |
|---|---|---|---|---|---|---|
| **1597** | Golga3 | 1.02E-06 | 0.488445414 | 0.02534388 | 0.471 | 0.115 |
| **1598** | Sntb1 | 1.02E-06 | 0.869342964 | 0.025346734 | 0.587 | 0.192 |
| **1599** | Zcchc24 | 1.02E-06 | 0.635437475 | 0.025378692 | 0.654 | 0.202 |
| **1600** | Arpc1b | 1.02E-06 | 0.43010472 | 0.025383452 | 0.413 | 0.087 |
| **1601** | Kdm1a | 1.03E-06 | 0.399939654 | 0.025601656 | 0.558 | 0.163 |
| **1602** | Glud1 | 1.03E-06 | 0.190126794 | 0.025657994 | 0.635 | 0.192 |
| **1603** | Psma3 | 1.03E-06 | 0.050176963 | 0.025657994 | 0.635 | 0.192 |
| **1604** | Sgpl1 | 1.03E-06 | 0.481101015 | 0.025658905 | 0.423 | 0.087 |
| **1605** | Amotl2 | 1.03E-06 | 0.601798151 | 0.025686349 | 0.596 | 0.183 |
| **1606** | Vapa | 1.03E-06 | 0.325035996 | 0.025710115 | 0.654 | 0.202 |
| **1607** | Ppp1r18 | 1.04E-06 | 1.404679596 | 0.025806414 | 0.327 | 0.048 |
| **1608** | Gpr89 | 1.04E-06 | 2.243790753 | 0.025902649 | 0.26 | 0.019 |
| **1609** | Lsm6 | 1.06E-06 | 1.658337509 | 0.026410218 | 0.26 | 0.019 |
| **1610** | Rab5a | 1.07E-06 | 0.584559359 | 0.026540363 | 0.596 | 0.173 |
| **1611** | Crppa | 1.07E-06 | 0.836355897 | 0.026638179 | 0.462 | 0.115 |
| **1612** | Ilf2 | 1.07E-06 | 1.324841051 | 0.026740169 | 0.308 | 0.038 |
| **1613** | Wbp2 | 1.08E-06 | 3.640627596 | 0.026800082 | 0.231 | 0.01 |
| **1614** | Wdr83os | 1.08E-06 | 2.159485517 | 0.026927351 | 0.26 | 0.019 |
| **1615** | Mpc2 | 1.09E-06 | 0.640024466 | 0.02699838 | 0.375 | 0.067 |
| **1616** | Sdf2 | 1.09E-06 | 1.000245738 | 0.02699838 | 0.375 | 0.067 |
| **1617** | Ifit2 | 1.09E-06 | 1.226687388 | 0.027147662 | 0.346 | 0.058 |
| **1618** | Ak2 | 1.09E-06 | 1.395645403 | 0.027217887 | 0.308 | 0.038 |
| **1619** | Spg11 | 1.10E-06 | 0.580236246 | 0.027435631 | 0.375 | 0.067 |
| **1620** | Wars | 1.10E-06 | 0.444455759 | 0.027435631 | 0.375 | 0.067 |
| **1621** | Lims1 | 1.11E-06 | 0.599671653 | 0.027566 | 0.962 | 0.452 |
| **1622** | Trim37 | 1.11E-06 | 0.504802206 | 0.027638842 | 0.327 | 0.048 |
| **1623** | Mfn1 | 1.12E-06 | 0.422528564 | 0.027879693 | 0.375 | 0.067 |
| **1624** | Ago3 | 1.12E-06 | 0.351748448 | 0.027970163 | 0.74 | 0.25 |
| **1625** | Psmb4 | 1.12E-06 | 1.776355822 | 0.027991007 | 0.26 | 0.019 |
| **1626** | Ctnnbip1 | 1.13E-06 | 0.04991638 | 0.028034406 | 0.558 | 0.154 |
| **1627** | Ngly1 | 1.13E-06 | 0.409952868 | 0.028070139 | 0.615 | 0.183 |
| **1628** | Cnrip1 | 1.13E-06 | 0.978525666 | 0.028198081 | 0.308 | 0.038 |
| **1629** | Ppm1g | 1.13E-06 | 1.139129508 | 0.028198081 | 0.308 | 0.038 |
| **1630** | Pon2 | 1.13E-06 | 0.572504429 | 0.028198268 | 0.442 | 0.096 |
| **1631** | Gba2 | 1.15E-06 | 0.878151002 | 0.028700829 | 0.308 | 0.038 |
| **1632** | Map1lc3b | 1.15E-06 | 1.311470686 | 0.028700829 | 0.308 | 0.038 |
| **1633** | Ube2v1 | 1.15E-06 | 0.552225303 | 0.028700829 | 0.308 | 0.038 |
| **1634** | Alkbh5 | 1.16E-06 | 0.985962525 | 0.028831713 | 0.433 | 0.096 |
| **1635** | Col4a2 | 1.17E-06 | 0.610639373 | 0.029047644 | 1 | 0.587 |
| **1636** | Cd200 | 1.18E-06 | 0.292381783 | 0.029371343 | 0.76 | 0.269 |
| **1637** | Clip2 | 1.18E-06 | 1.082820834 | 0.029435304 | 0.385 | 0.077 |

| | | | | | | |
|---|---|---|---|---|---|---|
| **1638** | Hnrnpa1 | 1.19E-06 | 0.924776759 | 0.02970967 | 0.433 | 0.096 |
| **1639** | Psmc1 | 1.21E-06 | 0.546174263 | 0.030105928 | 0.327 | 0.048 |
| **1640** | H2-Q6 | 1.21E-06 | 0.62887711 | 0.030161345 | 0.49 | 0.125 |
| **1641** | Nsmce4a | 1.22E-06 | 3.400011783 | 0.030330514 | 0.231 | 0.01 |
| **1642** | Sptbn1 | 1.22E-06 | 0.347023669 | 0.030364102 | 1 | 0.942 |
| **1643** | Max | 1.23E-06 | 0.01371933 | 0.030529193 | 0.538 | 0.144 |
| **1644** | Ranbp1 | 1.24E-06 | 3.529547462 | 0.030960831 | 0.231 | 0.01 |
| **1645** | Rexo2 | 1.25E-06 | 0.292256941 | 0.031192431 | 0.481 | 0.115 |
| **1646** | Abcb1b | 1.25E-06 | 0.603613333 | 0.031224339 | 0.529 | 0.144 |
| **1647** | Cr1l | 1.27E-06 | 0.17136838 | 0.031629937 | 0.769 | 0.26 |
| **1648** | Ttl | 1.27E-06 | 2.02353835 | 0.03164893 | 0.279 | 0.029 |
| **1649** | Plekhg3 | 1.29E-06 | 0.212667354 | 0.032024032 | 0.654 | 0.202 |
| **1650** | mt-Nd3 | 1.31E-06 | 0.529095125 | 0.032494296 | 0.913 | 0.423 |
| **1651** | Sipa1 | 1.31E-06 | 0.179945595 | 0.03256375 | 0.481 | 0.115 |
| **1652** | Slc25a4 | 1.32E-06 | 1.733750185 | 0.03283972 | 0.279 | 0.029 |
| **1653** | Aig1 | 1.34E-06 | -0.34298113 | 0.033227201 | 0.779 | 0.25 |
| **1654** | Rif1 | 1.34E-06 | 0.599539796 | 0.033311837 | 0.49 | 0.125 |
| **1655** | AI480526 | 1.34E-06 | 0.727981972 | 0.033346646 | 0.327 | 0.048 |
| **1656** | Stat2 | 1.34E-06 | 0.61636232 | 0.033364027 | 0.471 | 0.115 |
| **1657** | Zscan26 | 1.35E-06 | 0.080971141 | 0.033473748 | 0.606 | 0.173 |
| **1658** | Slc30a5 | 1.35E-06 | 0.46612477 | 0.033603578 | 0.452 | 0.106 |
| **1659** | Aifm1 | 1.35E-06 | 2.947257969 | 0.033610441 | 0.231 | 0.01 |
| **1660** | Cops7a | 1.35E-06 | 2.725344508 | 0.033610441 | 0.231 | 0.01 |
| **1661** | Ppp1ca | 1.35E-06 | 3.334197453 | 0.033610441 | 0.231 | 0.01 |
| **1662** | Snrpb2 | 1.35E-06 | 2.681989337 | 0.033610441 | 0.231 | 0.01 |
| **1663** | Copg1 | 1.36E-06 | 0.525019377 | 0.033907687 | 0.394 | 0.077 |
| **1664** | Pde4a | 1.36E-06 | 0.461084572 | 0.033907687 | 0.394 | 0.077 |
| **1665** | Wtap | 1.38E-06 | -0.01659617 | 0.034232892 | 0.587 | 0.163 |
| **1666** | Dtx3l | 1.38E-06 | 1.164839959 | 0.034235757 | 0.346 | 0.058 |
| **1667** | Ankfy1 | 1.38E-06 | 0.135836354 | 0.034303123 | 0.702 | 0.221 |
| **1668** | Hgs | 1.38E-06 | 2.834909325 | 0.034306164 | 0.231 | 0.01 |
| **1669** | Plscr4 | 1.40E-06 | 0.301071235 | 0.034869886 | 0.817 | 0.308 |
| **1670** | Dusp11 | 1.41E-06 | 0.16631869 | 0.035003994 | 0.663 | 0.202 |
| **1671** | Trappc6b | 1.42E-06 | 0.799835417 | 0.035251794 | 0.49 | 0.125 |
| **1672** | Syn3 | 1.42E-06 | 1.67925703 | 0.03535188 | 0.279 | 0.029 |
| **1673** | Peg3 | 1.43E-06 | 1.090847402 | 0.035540145 | 0.385 | 0.077 |
| **1674** | Snw1 | 1.43E-06 | 1.468111307 | 0.035690796 | 0.327 | 0.048 |
| **1675** | Ndufb5 | 1.45E-06 | 1.289113454 | 0.036008221 | 0.279 | 0.029 |
| **1676** | Ganab | 1.45E-06 | 0.883392778 | 0.036077042 | 0.413 | 0.087 |
| **1677** | Clstn1 | 1.47E-06 | 0.467622748 | 0.036482777 | 0.615 | 0.183 |
| **1678** | Efr3a | 1.47E-06 | 0.475553478 | 0.03651552 | 0.625 | 0.192 |

| | | | | | | |
|---|---|---|---|---|---|---|
| **1679** | Dyrk2 | 1.47E-06 | 1.422889662 | 0.036568064 | 0.346 | 0.058 |
| **1680** | Ldb1 | 1.47E-06 | 0.655680751 | 0.036628826 | 0.413 | 0.087 |
| **1681** | Glrx3 | 1.47E-06 | 0.907394933 | 0.036669399 | 0.385 | 0.077 |
| **1682** | Odf2l | 1.48E-06 | 0.538693239 | 0.036921511 | 0.327 | 0.048 |
| **1683** | Psmd12 | 1.48E-06 | 0.589983845 | 0.036921511 | 0.327 | 0.048 |
| **1684** | Immt | 1.49E-06 | 0.316127365 | 0.037070688 | 0.548 | 0.154 |
| **1685** | Itga6 | 1.50E-06 | 0.327239443 | 0.037351177 | 0.942 | 0.442 |
| **1686** | Ube2s | 1.50E-06 | 1.810057021 | 0.037356229 | 0.279 | 0.029 |
| **1687** | Orc6 | 1.51E-06 | 2.263099299 | 0.037490628 | 0.298 | 0.038 |
| **1688** | Aurkaip1 | 1.51E-06 | 4.585871186 | 0.03752639 | 0.202 | 0 |
| **1689** | Cetn2 | 1.51E-06 | 4.637911219 | 0.03752639 | 0.202 | 0 |
| **1690** | Emc4 | 1.51E-06 | 4.566308979 | 0.03752639 | 0.202 | 0 |
| **1691** | Higd1b | 1.51E-06 | 4.811999079 | 0.03752639 | 0.202 | 0 |
| **1692** | Msantd4 | 1.51E-06 | 4.497280436 | 0.03752639 | 0.202 | 0 |
| **1693** | Psenen | 1.51E-06 | 4.746790972 | 0.03752639 | 0.202 | 0 |
| **1694** | Rasl11a | 1.51E-06 | 5.074933247 | 0.03752639 | 0.202 | 0 |
| **1695** | Serpinf1 | 1.51E-06 | 4.631530757 | 0.03752639 | 0.202 | 0 |
| **1696** | Tmem173 | 1.51E-06 | 4.477062785 | 0.03752639 | 0.202 | 0 |
| **1697** | Adipor1 | 1.52E-06 | 0.457680852 | 0.037730023 | 0.462 | 0.115 |
| **1698** | Gatad1 | 1.52E-06 | 0.804192065 | 0.03783311 | 0.385 | 0.077 |
| **1699** | Atp1a1 | 1.53E-06 | 0.600829384 | 0.037947066 | 0.423 | 0.096 |
| **1700** | Dcbld2 | 1.53E-06 | 0.288329887 | 0.037966924 | 0.471 | 0.115 |
| **1701** | Orc3 | 1.53E-06 | 0.764752271 | 0.038048889 | 0.442 | 0.106 |
| **1702** | Tia1 | 1.53E-06 | 0.516470924 | 0.038168288 | 0.587 | 0.173 |
| **1703** | Cyb5b | 1.54E-06 | -0.1739819 | 0.038274802 | 0.558 | 0.154 |
| **1704** | Tmem87a | 1.54E-06 | 0.227324171 | 0.038275012 | 0.673 | 0.212 |
| **1705** | Gpr160 | 1.54E-06 | 0.037633175 | 0.038319808 | 0.654 | 0.202 |
| **1706** | Rasa4 | 1.55E-06 | 0.894600758 | 0.038485836 | 0.404 | 0.087 |
| **1707** | B230307C23R | 1.56E-06 | 1.415603632 | 0.038839852 | 0.298 | 0.038 |
| **1708** | Prkx | 1.56E-06 | 0.645603417 | 0.038856383 | 0.365 | 0.067 |
| **1709** | Tyw1 | 1.59E-06 | 0.352197486 | 0.039482213 | 0.365 | 0.067 |
| **1710** | Nr1d2 | 1.61E-06 | 0.4360371 | 0.04001865 | 0.654 | 0.212 |
| **1711** | Shisa5 | 1.61E-06 | 0.649389565 | 0.040027482 | 0.433 | 0.096 |
| **1712** | Sec61g | 1.61E-06 | 1.303340095 | 0.040029985 | 0.317 | 0.048 |
| **1713** | Cpsf7 | 1.62E-06 | 0.400466929 | 0.040200345 | 0.548 | 0.154 |
| **1714** | Sncg | 1.62E-06 | 0.689643216 | 0.040235691 | 0.298 | 0.038 |
| **1715** | Setd7 | 1.62E-06 | 0.559772161 | 0.040293006 | 0.519 | 0.144 |
| **1716** | Prpsap1 | 1.62E-06 | 0.843404182 | 0.040355176 | 0.346 | 0.058 |
| **1717** | Spred1 | 1.63E-06 | 0.430501891 | 0.040579492 | 0.779 | 0.298 |
| **1718** | Atf2 | 1.64E-06 | 0.414739311 | 0.040708106 | 0.817 | 0.288 |
| **1719** | H2-Aa | 1.64E-06 | 1.235250114 | 0.040718692 | 0.317 | 0.048 |

| | | | | | | |
|---|---|---|---|---|---|---|
| **1720** | Samd4b | 1.64E-06 | 0.456449986 | 0.040899802 | 0.385 | 0.077 |
| **1721** | Mpp5 | 1.66E-06 | 0.021158443 | 0.041214858 | 0.587 | 0.163 |
| **1722** | Ythdf2 | 1.66E-06 | -0.08275216 | 0.041230917 | 0.433 | 0.096 |
| **1723** | 5-Mar | 1.66E-06 | 0.52194589 | 0.041244277 | 0.51 | 0.135 |
| **1724** | Acly | 1.66E-06 | 0.679877542 | 0.041413704 | 0.519 | 0.144 |
| **1725** | Zfp68 | 1.66E-06 | 0.790587167 | 0.041418381 | 0.365 | 0.067 |
| **1726** | Sars | 1.66E-06 | 1.474410449 | 0.04141878 | 0.317 | 0.048 |
| **1727** | Hmgxb4 | 1.68E-06 | 0.702148854 | 0.041863061 | 0.337 | 0.058 |
| **1728** | Top3b | 1.70E-06 | 0.555697602 | 0.042387402 | 0.346 | 0.058 |
| **1729** | Tnfaip1 | 1.71E-06 | 0.292221779 | 0.042426249 | 0.548 | 0.154 |
| **1730** | Azi2 | 1.72E-06 | 0.42075418 | 0.04280284 | 0.529 | 0.144 |
| **1731** | Vps26a | 1.75E-06 | 0.606176161 | 0.043655901 | 0.49 | 0.135 |
| **1732** | Glt8d1 | 1.77E-06 | 0.936325455 | 0.04393944 | 0.298 | 0.038 |
| **1733** | Hmgn1 | 1.77E-06 | 1.02993361 | 0.04393944 | 0.298 | 0.038 |
| **1734** | Gli2 | 1.77E-06 | 0.694990378 | 0.044142331 | 0.365 | 0.067 |
| **1735** | Brip1os | 1.78E-06 | 0.782415477 | 0.044204684 | 0.385 | 0.077 |
| **1736** | Ndfip2 | 1.80E-06 | 0.560344403 | 0.044777831 | 0.49 | 0.125 |
| **1737** | Atr | 1.80E-06 | 0.624675592 | 0.044895774 | 0.385 | 0.077 |
| **1738** | Flywch1 | 1.82E-06 | 0.49514768 | 0.045250516 | 0.346 | 0.058 |
| **1739** | Mapk1ip1l | 1.82E-06 | 0.646750108 | 0.045375541 | 0.548 | 0.154 |
| **1740** | Foxp1 | 1.83E-06 | -0.99403588 | 0.045414796 | 0.99 | 0.913 |
| **1741** | Fam49a | 1.84E-06 | 0.368158045 | 0.045683493 | 0.683 | 0.221 |
| **1742** | Rad54l2 | 1.84E-06 | 0.179790741 | 0.045886085 | 0.587 | 0.173 |
| **1743** | Ppt1 | 1.85E-06 | 0.324918515 | 0.045994819 | 0.346 | 0.058 |
| **1744** | Zfp326 | 1.86E-06 | 0.228087397 | 0.046182168 | 0.519 | 0.135 |
| **1745** | Anapc11 | 1.86E-06 | 1.153450831 | 0.046217841 | 0.337 | 0.058 |
| **1746** | Cebpz | 1.86E-06 | 0.661946774 | 0.046297426 | 0.365 | 0.067 |
| **1747** | Blvra | 1.87E-06 | 2.417251609 | 0.046437739 | 0.25 | 0.019 |
| **1748** | Rcc2 | 1.87E-06 | 2.188024652 | 0.046437739 | 0.25 | 0.019 |
| **1749** | Lgals3bp | 1.88E-06 | 1.753961721 | 0.04665353 | 0.317 | 0.048 |
| **1750** | Gli3 | 1.90E-06 | 0.391261357 | 0.047238641 | 0.548 | 0.154 |
| **1751** | Arf1 | 1.90E-06 | 0.589095758 | 0.047309291 | 0.615 | 0.192 |
| **1752** | Pa2g4 | 1.90E-06 | 1.514936709 | 0.047340959 | 0.25 | 0.019 |
| **1753** | Dctn1 | 1.90E-06 | 0.687078027 | 0.047364458 | 0.442 | 0.106 |
| **1754** | Tcirg1 | 1.92E-06 | 1.592275191 | 0.047763813 | 0.337 | 0.058 |
| **1755** | Ssbp1 | 1.92E-06 | 0.751990404 | 0.047765389 | 0.385 | 0.077 |
| **1756** | Rcbtb2 | 1.92E-06 | 0.585941402 | 0.047789938 | 0.365 | 0.067 |
| **1757** | Rab21 | 1.94E-06 | 0.133842628 | 0.048240975 | 0.538 | 0.154 |
| **1758** | Ahr | 1.94E-06 | 0.079043398 | 0.048379687 | 0.471 | 0.115 |
| **1759** | Mast2 | 1.95E-06 | 0.206833449 | 0.048482935 | 0.625 | 0.192 |
| **1760** | Armc9 | 1.96E-06 | 1.313335247 | 0.048817253 | 0.298 | 0.038 |

| | | | | | | |
|---|---|---|---|---|---|---|
| **1761** | Mdh1 | 1.96E-06 | 0.61716643 | 0.048817253 | 0.298 | 0.038 |
| **1762** | Ipmk | 1.97E-06 | 1.340570699 | 0.049086572 | 0.317 | 0.048 |
| **1763** | Ajuba | 1.98E-06 | 1.772633398 | 0.049198296 | 0.25 | 0.019 |
| **1764** | Prkaca | 1.98E-06 | 1.465778865 | 0.049198296 | 0.25 | 0.019 |
| **1765** | Sf3a3 | 1.98E-06 | 2.204205431 | 0.049198296 | 0.25 | 0.019 |
| **1766** | Pink1 | 1.98E-06 | 1.114313474 | 0.049359403 | 0.337 | 0.058 |
| **1767** | Dcp1b | 1.98E-06 | 0.359489232 | 0.049380819 | 0.49 | 0.125 |

**Extended Table 3**

DEG from endothelial SnC (SenMayo strategy) versus old non-senescent

| | gene | mean_pval | mean_log2FC | p_val_adj | mean_pct_1 | mean_pct_2 |
|---|---|---|---|---|---|---|
| 1 | Dlc1 | 1.54E-13 | 1.966041809 | 3.83E-09 | 1 | 0.735 |
| 2 | Pla2r1 | 1.69E-11 | 2.48400063 | 4.21E-07 | 0.837 | 0.122 |
| 3 | Col6a3 | 1.85E-11 | 2.610711381 | 4.59E-07 | 0.939 | 0.204 |
| 4 | Mrvi1 | 4.40E-11 | 2.744893517 | 1.09E-06 | 0.816 | 0.102 |
| 5 | Cacna1c | 8.41E-11 | 2.725726783 | 2.09E-06 | 0.878 | 0.163 |
| 6 | Col5a3 | 1.13E-10 | 2.661541124 | 2.82E-06 | 0.857 | 0.184 |
| 7 | Ebf2 | 1.26E-10 | 2.401378282 | 3.13E-06 | 0.918 | 0.265 |
| 8 | Pde3a | 1.42E-10 | 2.079468968 | 3.54E-06 | 0.98 | 0.265 |
| 9 | Col6a2 | 1.60E-10 | 2.298717338 | 3.99E-06 | 0.878 | 0.163 |
| 10 | Hspa9 | 2.33E-10 | 3.374767319 | 5.80E-06 | 0.714 | 0.061 |
| 11 | Slit3 | 2.36E-10 | 1.779040672 | 5.86E-06 | 1 | 0.633 |
| 12 | Gucy1a2 | 2.95E-10 | 2.957471322 | 7.35E-06 | 0.796 | 0.163 |
| 13 | Adgrl3 | 3.16E-10 | 2.752773951 | 7.87E-06 | 0.776 | 0.082 |
| 14 | Abcc9 | 3.78E-10 | 3.250026878 | 9.40E-06 | 0.755 | 0.102 |
| 15 | Adamts12 | 4.27E-10 | 1.813606646 | 1.06E-05 | 0.939 | 0.224 |
| 16 | Agtr1a | 4.40E-10 | 3.086284236 | 1.09E-05 | 0.755 | 0.102 |
| 17 | Cacna2d1 | 4.71E-10 | 2.159455622 | 1.17E-05 | 0.857 | 0.163 |
| 18 | Notch3 | 6.64E-10 | 3.374572232 | 1.65E-05 | 0.755 | 0.143 |
| 19 | Dtx3 | 9.05E-10 | 2.840627515 | 2.25E-05 | 0.694 | 0.061 |
| 20 | C1s1 | 1.03E-09 | 2.021858603 | 2.55E-05 | 0.816 | 0.122 |
| 21 | Plcl1 | 1.08E-09 | 2.086876169 | 2.68E-05 | 0.939 | 0.245 |
| 22 | Arhgap6 | 1.11E-09 | 1.838844866 | 2.76E-05 | 1 | 0.327 |
| 23 | Dusp6 | 1.36E-09 | 4.468583087 | 3.39E-05 | 0.592 | 0.02 |
| 24 | Postn | 1.54E-09 | 3.114221008 | 3.83E-05 | 0.776 | 0.143 |
| 25 | Arhgap42 | 1.65E-09 | 2.004612898 | 4.10E-05 | 0.857 | 0.204 |
| 26 | Col6a1 | 1.69E-09 | 1.877689985 | 4.22E-05 | 0.796 | 0.122 |
| 27 | Tmem65 | 1.94E-09 | 1.747362632 | 4.82E-05 | 0.796 | 0.102 |
| 28 | Synpo2 | 2.15E-09 | 2.993460377 | 5.36E-05 | 0.694 | 0.082 |
| 29 | Pdia4 | 2.89E-09 | 4.089652304 | 7.18E-05 | 0.592 | 0.02 |
| 30 | Enox1 | 4.51E-09 | 2.089866317 | 0.000112218 | 0.816 | 0.143 |
| 31 | Pdgfrb | 4.99E-09 | 2.723245209 | 0.000124234 | 0.755 | 0.143 |
| 32 | Tmed9 | 5.49E-09 | 1.731815828 | 0.000136486 | 0.694 | 0.061 |
| 33 | Slc24a3 | 6.12E-09 | 2.071336451 | 0.000152274 | 0.918 | 0.306 |
| 34 | Plekhg3 | 6.16E-09 | 1.435065323 | 0.000153239 | 0.857 | 0.163 |
| 35 | Dock7 | 6.30E-09 | 1.836906165 | 0.00015674 | 0.837 | 0.184 |
| 36 | Ror1 | 7.85E-09 | 1.874707207 | 0.000195349 | 0.878 | 0.204 |
| 37 | Col5a1 | 7.95E-09 | 1.659532685 | 0.000197775 | 0.755 | 0.143 |
| 38 | Sox5 | 8.10E-09 | 1.595173611 | 0.000201501 | 1 | 0.408 |

| | | | | | | |
|---|---|---|---|---|---|---|
| **39** | Gucy1a1 | 8.73E-09 | 1.975069845 | 0.000217265 | 0.776 | 0.143 |
| **40** | Dag1 | 9.90E-09 | 1.388454346 | 0.000246439 | 0.837 | 0.204 |
| **41** | Adap2 | 9.97E-09 | 2.345734724 | 0.000248146 | 0.735 | 0.122 |
| **42** | Cd63 | 1.07E-08 | 2.148643314 | 0.000266717 | 0.776 | 0.122 |
| **43** | Arhgap32 | 1.22E-08 | 1.302346746 | 0.000303637 | 0.878 | 0.204 |
| **44** | Steap4 | 1.64E-08 | 1.628015061 | 0.000408578 | 0.796 | 0.184 |
| **45** | Robo1 | 1.64E-08 | 2.913811613 | 0.000408668 | 0.694 | 0.122 |
| **46** | Pakap | 2.04E-08 | 3.778894852 | 0.000507558 | 0.612 | 0.082 |
| **47** | Bzw2 | 2.09E-08 | 2.852887218 | 0.000519966 | 0.633 | 0.061 |
| **48** | Sema5a | 2.17E-08 | 1.92999406 | 0.000540376 | 0.837 | 0.224 |
| **49** | Tmf1 | 2.47E-08 | 1.332750854 | 0.000614017 | 0.755 | 0.122 |
| **50** | Gpc6 | 2.60E-08 | 1.83206869 | 0.000646317 | 0.878 | 0.265 |
| **51** | Anxa1 | 2.71E-08 | 1.861021096 | 0.000673871 | 0.755 | 0.143 |
| **52** | Aspn | 2.72E-08 | 3.664286448 | 0.000677443 | 0.612 | 0.082 |
| **53** | Ikbip | 3.18E-08 | 4.758487666 | 0.000790056 | 0.49 | 0 |
| **54** | Dennd2a | 3.21E-08 | 2.563325006 | 0.000798019 | 0.592 | 0.041 |
| **55** | Stac | 3.71E-08 | 1.636966962 | 0.00092317 | 0.673 | 0.102 |
| **56** | Rbms3 | 3.86E-08 | 1.510207308 | 0.000960825 | 0.98 | 0.408 |
| **57** | Ano1 | 4.26E-08 | 2.211815947 | 0.001059277 | 0.673 | 0.102 |
| **58** | Lamb1 | 4.34E-08 | 1.56997686 | 0.001080963 | 0.857 | 0.184 |
| **59** | Rtn4rl1 | 4.97E-08 | 2.26274338 | 0.001237806 | 0.592 | 0.041 |
| **60** | Fam129a | 5.03E-08 | 1.675081703 | 0.001252241 | 0.837 | 0.245 |
| **61** | Gsn | 5.05E-08 | 1.832080478 | 0.001255506 | 1 | 0.653 |
| **62** | Rbbp7 | 5.16E-08 | 2.542400774 | 0.00128497 | 0.612 | 0.061 |
| **63** | Zfhx3 | 5.31E-08 | 1.507026243 | 0.001321904 | 0.98 | 0.286 |
| **64** | Inpp4b | 6.04E-08 | 1.779599986 | 0.001502615 | 0.837 | 0.224 |
| **65** | Adam23 | 6.57E-08 | 2.89903353 | 0.001635116 | 0.571 | 0.041 |
| **66** | Rnase4 | 7.12E-08 | 2.857317587 | 0.001771981 | 0.592 | 0.061 |
| **67** | Igf2r | 8.17E-08 | 1.704106862 | 0.002033691 | 0.837 | 0.204 |
| **68** | Kirrel | 8.17E-08 | 1.705386357 | 0.002033691 | 0.816 | 0.224 |
| **69** | Nbr1 | 8.75E-08 | 1.093509628 | 0.002176338 | 0.776 | 0.143 |
| **70** | Sp3 | 8.88E-08 | 1.40078123 | 0.002210419 | 0.837 | 0.204 |
| **71** | Col1a2 | 9.65E-08 | 1.978899621 | 0.002401976 | 0.816 | 0.224 |
| **72** | Fap | 1.02E-07 | 1.334395842 | 0.002542722 | 0.714 | 0.102 |
| **73** | Tbc1d1 | 1.10E-07 | 1.527486387 | 0.002736171 | 0.837 | 0.184 |
| **74** | Tapbp | 1.10E-07 | 1.602449316 | 0.002737505 | 0.816 | 0.184 |
| **75** | Nckap5 | 1.16E-07 | 1.309334125 | 0.002897293 | 0.918 | 0.265 |
| **76** | Ecm2 | 1.27E-07 | 1.86489057 | 0.00317012 | 0.653 | 0.082 |
| **77** | Prrx1 | 1.55E-07 | 2.020529276 | 0.003862032 | 0.755 | 0.184 |
| **78** | Rps28 | 1.57E-07 | 1.541263749 | 0.003914047 | 0.898 | 0.286 |
| **79** | Zscan26 | 1.69E-07 | 1.161596007 | 0.004195461 | 0.755 | 0.122 |

| | | | | | | |
|---|---|---|---|---|---|---|
| **80** | Col1a1 | 1.80E-07 | 1.423897806 | 0.004480973 | 0.816 | 0.245 |
| **81** | Frg1 | 1.90E-07 | 1.436008728 | 0.004719823 | 0.531 | 0.041 |
| **82** | Nr1h3 | 1.90E-07 | 1.181201101 | 0.004723013 | 0.592 | 0.061 |
| **83** | Jade1 | 1.92E-07 | 2.25980818 | 0.004777605 | 0.633 | 0.082 |
| **84** | Tnxb | 1.96E-07 | 1.153758745 | 0.004873673 | 0.878 | 0.286 |
| **85** | Ugp2 | 2.09E-07 | 1.721408066 | 0.005201406 | 0.694 | 0.122 |
| **86** | Chn1 | 2.33E-07 | 2.817733049 | 0.005791135 | 0.633 | 0.122 |
| **87** | Samd4 | 2.48E-07 | 1.40078925 | 0.006168458 | 0.857 | 0.245 |
| **88** | Hnrnpul2 | 2.70E-07 | 1.400385421 | 0.006726215 | 0.796 | 0.224 |
| **89** | Aoc3 | 2.75E-07 | 1.589631185 | 0.006840535 | 0.673 | 0.102 |
| **90** | H2afz | 2.79E-07 | 2.473790965 | 0.006947654 | 0.551 | 0.061 |
| **91** | Itpripl2 | 2.83E-07 | 1.417524679 | 0.007036736 | 0.776 | 0.163 |
| **92** | Frmd6 | 2.83E-07 | 1.958558379 | 0.007051017 | 0.735 | 0.184 |
| **93** | Ccz1 | 2.90E-07 | 3.237078795 | 0.007221446 | 0.49 | 0.02 |
| **94** | Lhfp | 3.03E-07 | 2.2319763 | 0.007551259 | 0.673 | 0.163 |
| **95** | Lin7a | 3.20E-07 | 3.390275388 | 0.007973578 | 0.571 | 0.082 |
| **96** | Pgap2 | 3.34E-07 | 3.54661802 | 0.008314139 | 0.469 | 0.02 |
| **97** | Snrnp48 | 3.57E-07 | 1.08428916 | 0.008887987 | 0.571 | 0.061 |
| **98** | Angptl4 | 3.61E-07 | 0.771074258 | 0.008980188 | 0.878 | 0.224 |
| **99** | Gjc1 | 3.63E-07 | 1.644443087 | 0.009033788 | 0.694 | 0.122 |
| **100** | Dazap1 | 3.67E-07 | 1.977749972 | 0.009126806 | 0.571 | 0.082 |
| **101** | Bsg | 3.84E-07 | 1.672773331 | 0.009551709 | 0.755 | 0.184 |
| **102** | Abca8a | 3.96E-07 | 2.389700926 | 0.00984268 | 0.653 | 0.122 |
| **103** | Trim12c | 4.32E-07 | 3.076474544 | 0.010753408 | 0.51 | 0.041 |
| **104** | Ttc17 | 4.38E-07 | 1.495467627 | 0.010893337 | 0.776 | 0.184 |
| **105** | Map3k20 | 4.79E-07 | 1.424863703 | 0.01191582 | 0.878 | 0.245 |
| **106** | Nop53 | 4.87E-07 | 1.42968649 | 0.012114197 | 0.612 | 0.082 |
| **107** | Angptl2 | 5.14E-07 | 1.459003629 | 0.012784905 | 0.776 | 0.184 |
| **108** | Gli3 | 5.40E-07 | 1.74261304 | 0.013434112 | 0.673 | 0.143 |
| **109** | Egfr | 5.59E-07 | 1.503172146 | 0.013903681 | 0.837 | 0.224 |
| **110** | P3h2 | 5.79E-07 | 2.348876718 | 0.014399857 | 0.653 | 0.143 |
| **111** | Chd4 | 5.87E-07 | 1.25232835 | 0.014611448 | 0.857 | 0.245 |
| **112** | Enpep | 5.99E-07 | 2.095759324 | 0.014896305 | 0.633 | 0.122 |
| **113** | Hsp90ab1 | 6.07E-07 | 1.291709495 | 0.015110772 | 0.939 | 0.306 |
| **114** | Arhgap31 | 6.21E-07 | -1.33666973 | 0.015458242 | 0.939 | 0.98 |
| **115** | Calm2 | 6.29E-07 | 0.998438676 | 0.015639617 | 0.796 | 0.184 |
| **116** | Atn1 | 6.35E-07 | 2.347445462 | 0.015801767 | 0.551 | 0.061 |
| **117** | Cnot2 | 6.47E-07 | 1.103450126 | 0.016098886 | 0.918 | 0.286 |
| **118** | Prkca | 7.27E-07 | 0.758713213 | 0.018093684 | 0.837 | 0.184 |
| **119** | Hoxc6 | 7.46E-07 | 1.625479963 | 0.018555091 | 0.531 | 0.061 |
| **120** | Lrp1 | 7.53E-07 | 1.563034762 | 0.018726662 | 0.898 | 0.306 |

| | | | | | | |
|---|---|---|---|---|---|---|
| **121** | Yars | 7.61E-07 | 3.479059593 | 0.01893313 | 0.449 | 0.02 |
| **122** | Tpm1 | 7.80E-07 | 1.261029404 | 0.019411754 | 0.776 | 0.204 |
| **123** | Akirin2 | 7.80E-07 | 2.772562092 | 0.019413059 | 0.531 | 0.061 |
| **124** | Ccdc58 | 7.84E-07 | 4.272190411 | 0.019508642 | 0.408 | 0 |
| **125** | Pdcl | 7.84E-07 | 4.17566865 | 0.019508642 | 0.408 | 0 |
| **126** | Ypel3 | 8.36E-07 | 1.202494485 | 0.02079787 | 0.571 | 0.061 |
| **127** | Arhgef10l | 8.50E-07 | 2.847315597 | 0.021136757 | 0.531 | 0.082 |
| **128** | Rarres2 | 8.50E-07 | 2.047726558 | 0.021158406 | 0.571 | 0.082 |
| **129** | Slc38a6 | 8.52E-07 | 1.228209346 | 0.021197606 | 0.694 | 0.143 |
| **130** | Trpc3 | 8.54E-07 | 2.737303953 | 0.021245065 | 0.531 | 0.061 |
| **131** | Atp5a1 | 8.59E-07 | 1.685200336 | 0.021377826 | 0.755 | 0.204 |
| **132** | Larp4b | 8.65E-07 | 0.826951929 | 0.021511833 | 0.959 | 0.265 |
| **133** | Sae1 | 8.68E-07 | 3.052832963 | 0.021586361 | 0.51 | 0.061 |
| **134** | Atm | 8.93E-07 | 2.431315194 | 0.022222417 | 0.531 | 0.061 |
| **135** | Xpr1 | 9.02E-07 | 0.5920105 | 0.022447019 | 0.837 | 0.245 |
| **136** | Rmnd5a | 9.29E-07 | 1.007919606 | 0.023117565 | 0.776 | 0.184 |
| **137** | Arhgap10 | 9.43E-07 | 1.400056955 | 0.023457334 | 0.898 | 0.306 |
| **138** | Cygb | 9.49E-07 | 2.516902176 | 0.023613615 | 0.633 | 0.122 |
| **139** | H13 | 9.90E-07 | 0.944165152 | 0.024634494 | 0.735 | 0.163 |
| **140** | Fstl1 | 1.02E-06 | 1.505667196 | 0.025430943 | 0.837 | 0.265 |
| **141** | Kcnq5 | 1.02E-06 | 2.062916226 | 0.025493336 | 0.735 | 0.245 |
| **142** | Ssr1 | 1.03E-06 | 1.72219627 | 0.025645773 | 0.653 | 0.143 |
| **143** | Fam49a | 1.09E-06 | 1.622209456 | 0.027014894 | 0.755 | 0.204 |
| **144** | Hnrnpk | 1.11E-06 | 1.35216605 | 0.027597182 | 0.857 | 0.265 |
| **145** | Axin1 | 1.12E-06 | 1.501363059 | 0.027948528 | 0.653 | 0.122 |
| **146** | Ghitm | 1.13E-06 | 2.153550087 | 0.028196835 | 0.612 | 0.102 |
| **147** | Il34 | 1.19E-06 | 3.342768321 | 0.029632978 | 0.51 | 0.061 |
| **148** | Tmem273 | 1.21E-06 | 4.938446536 | 0.030175543 | 0.429 | 0.02 |
| **149** | Nectin3 | 1.22E-06 | 2.396980425 | 0.030381748 | 0.531 | 0.061 |
| **150** | Gfm1 | 1.24E-06 | 3.312651627 | 0.030799807 | 0.449 | 0.02 |
| **151** | Sod1 | 1.27E-06 | 1.936411252 | 0.031530419 | 0.633 | 0.122 |
| **152** | Rps24 | 1.30E-06 | 0.940486804 | 0.032387481 | 0.959 | 0.367 |
| **153** | Rnpep | 1.37E-06 | 2.100087649 | 0.034029755 | 0.571 | 0.082 |
| **154** | Adcy3 | 1.47E-06 | 3.98569971 | 0.036692788 | 0.429 | 0.02 |
| **155** | Vrk2 | 1.51E-06 | 2.142126808 | 0.037601162 | 0.592 | 0.102 |
| **156** | Col3a1 | 1.57E-06 | 1.091320108 | 0.038977794 | 1 | 0.429 |
| **157** | Ccpg1 | 1.61E-06 | 0.37802249 | 0.040016195 | 0.673 | 0.122 |
| **158** | Olfm2 | 1.65E-06 | 4.01206266 | 0.040938313 | 0.469 | 0.061 |
| **159** | Myh11 | 1.65E-06 | 2.511539186 | 0.041123161 | 0.633 | 0.143 |
| **160** | Ccl11 | 1.69E-06 | 4.616159101 | 0.041939368 | 0.388 | 0 |
| **161** | Gpx8 | 1.69E-06 | 4.463014113 | 0.041939368 | 0.388 | 0 |

| | | | | | | |
|---|---|---|---|---|---|---|
| **162** | Zfc3h1 | 1.75E-06 | 1.17109456 | 0.043615822 | 0.796 | 0.224 |
| **163** | Pard3 | 1.77E-06 | 1.134720544 | 0.04395536 | 0.918 | 0.286 |
| **164** | Ppp4r2 | 1.83E-06 | 0.585705376 | 0.045416683 | 0.857 | 0.245 |
| **165** | Fbh1 | 1.87E-06 | 1.452517152 | 0.046417111 | 0.612 | 0.102 |
| **166** | Fam110b | 1.90E-06 | 3.231274661 | 0.047193196 | 0.51 | 0.082 |
| **167** | Pik3r1 | 1.95E-06 | 1.403199231 | 0.048606192 | 0.959 | 0.408 |
| **168** | 4632427E13Ri | 1.99E-06 | 1.27682826 | 0.049541725 | 0.694 | 0.163 |

**Extended Table 4**

DEG from endothelial SnC (SEAC strategy) versus old non-senescent

| | gene | mean_pval | mean_log2FC | p_val_adj | mean_pct_1 | mean_pct_2 |
|---|---|---|---|---|---|---|
| **1** | Cacna1c | 2.37E-25 | 3.005438214 | 5.89E-21 | 0.904 | 0.135 |
| **2** | Pdgfrb | 1.36E-22 | 3.07880465 | 3.37E-18 | 0.837 | 0.125 |
| **3** | Notch3 | 1.37E-22 | 3.004247742 | 3.41E-18 | 0.827 | 0.125 |
| **4** | Adgrl3 | 4.08E-22 | 3.033482425 | 1.02E-17 | 0.817 | 0.096 |
| **5** | Abcc9 | 2.21E-21 | 2.823274111 | 5.50E-17 | 0.817 | 0.125 |
| **6** | Enox1 | 9.51E-21 | 3.188114868 | 2.37E-16 | 0.76 | 0.077 |
| **7** | Mrvi1 | 9.99E-21 | 2.255177999 | 2.49E-16 | 0.827 | 0.115 |
| **8** | Pde3a | 1.81E-20 | 1.889567565 | 4.51E-16 | 0.933 | 0.192 |
| **9** | Plcl1 | 2.13E-20 | 2.425127314 | 5.30E-16 | 0.933 | 0.269 |
| **10** | Dlc1 | 2.93E-20 | 1.459161835 | 7.28E-16 | 1 | 0.75 |
| **11** | Stac | 3.25E-20 | 2.466016741 | 8.09E-16 | 0.75 | 0.077 |
| **12** | Gucy1a1 | 1.81E-19 | 2.277494172 | 4.51E-15 | 0.808 | 0.135 |
| **13** | Ano1 | 2.89E-19 | 2.611646966 | 7.20E-15 | 0.76 | 0.106 |
| **14** | Pakap | 3.24E-19 | 2.934269712 | 8.06E-15 | 0.731 | 0.077 |
| **15** | Sox5 | 6.05E-19 | 1.87241618 | 1.50E-14 | 0.971 | 0.346 |
| **16** | Inpp4b | 6.63E-19 | 2.11030153 | 1.65E-14 | 0.894 | 0.221 |
| **17** | Lhfp | 1.14E-18 | 2.603952199 | 2.85E-14 | 0.731 | 0.077 |
| **18** | Pla2r1 | 1.51E-18 | 2.117333012 | 3.76E-14 | 0.808 | 0.144 |
| **19** | Lin7a | 4.17E-18 | 3.833149754 | 1.04E-13 | 0.654 | 0.067 |
| **20** | Sema5a | 7.53E-18 | 2.267877335 | 1.87E-13 | 0.827 | 0.154 |
| **21** | Agtr1a | 7.57E-18 | 3.061803653 | 1.88E-13 | 0.702 | 0.087 |
| **22** | Postn | 1.33E-17 | 2.008775309 | 3.32E-13 | 0.827 | 0.173 |
| **23** | Ebf2 | 1.43E-17 | 1.241134271 | 3.56E-13 | 0.885 | 0.231 |
| **24** | Adamts12 | 1.46E-17 | 1.772440072 | 3.62E-13 | 0.933 | 0.288 |
| **25** | Arhgef17 | 2.54E-17 | 3.279508302 | 6.33E-13 | 0.663 | 0.077 |
| **26** | Gjc1 | 3.89E-17 | 3.130743091 | 9.68E-13 | 0.673 | 0.087 |
| **27** | Arhgap42 | 4.15E-17 | 2.376673522 | 1.03E-12 | 0.76 | 0.125 |
| **28** | Cdh2 | 8.53E-17 | 2.501058609 | 2.12E-12 | 0.635 | 0.058 |
| **29** | Rgs5 | 2.62E-16 | 2.008120383 | 6.52E-12 | 0.885 | 0.279 |
| **30** | Cacna2d1 | 2.67E-16 | 1.815201738 | 6.64E-12 | 0.808 | 0.154 |
| **31** | Chn1 | 3.91E-16 | 2.506256404 | 9.74E-12 | 0.673 | 0.096 |
| **32** | Aspn | 4.37E-16 | 2.618398233 | 1.09E-11 | 0.663 | 0.087 |
| **33** | Tmem273 | 1.01E-15 | 2.90004292 | 2.52E-11 | 0.606 | 0.058 |
| **34** | Dmd | 1.33E-15 | 1.658442114 | 3.32E-11 | 0.904 | 0.317 |
| **35** | Pten | 1.44E-15 | 1.557275327 | 3.59E-11 | 0.99 | 0.462 |
| **36** | Kcnq5 | 1.77E-15 | 1.993716154 | 4.41E-11 | 0.769 | 0.192 |
| **37** | Synpo2 | 2.30E-15 | 2.139203024 | 5.73E-11 | 0.663 | 0.087 |
| **38** | Daam2 | 7.32E-15 | 2.535017997 | 1.82E-10 | 0.635 | 0.087 |

| | | | | | | |
|---|---|---|---|---|---|---|
| **39** | Steap4 | 1.05E-14 | 1.692909113 | 2.61E-10 | 0.779 | 0.173 |
| **40** | Ednra | 1.10E-14 | 2.322328221 | 2.73E-10 | 0.606 | 0.067 |
| **41** | Adap2 | 1.73E-14 | 1.815736109 | 4.29E-10 | 0.692 | 0.106 |
| **42** | Ror1 | 2.19E-14 | 1.503538024 | 5.44E-10 | 0.808 | 0.192 |
| **43** | Arhgap6 | 2.40E-14 | 1.351046098 | 5.96E-10 | 0.942 | 0.298 |
| **44** | Cygb | 2.50E-14 | 2.976343207 | 6.22E-10 | 0.587 | 0.067 |
| **45** | Sgip1 | 5.88E-14 | 1.53126854 | 1.46E-09 | 0.913 | 0.365 |
| **46** | Ppp2r3a | 6.42E-14 | 1.625532788 | 1.60E-09 | 0.769 | 0.192 |
| **47** | Nckap5 | 7.43E-14 | 2.022297335 | 1.85E-09 | 0.702 | 0.135 |
| **48** | Angptl2 | 7.50E-14 | 2.330841679 | 1.87E-09 | 0.644 | 0.115 |
| **49** | Ddr2 | 1.03E-13 | 1.187598038 | 2.57E-09 | 0.817 | 0.192 |
| **50** | Spry4 | 1.08E-13 | 2.365945384 | 2.69E-09 | 0.519 | 0.038 |
| **51** | Col5a3 | 1.09E-13 | 1.343121688 | 2.72E-09 | 0.817 | 0.221 |
| **52** | Slc24a3 | 1.20E-13 | 1.157092264 | 2.99E-09 | 0.837 | 0.221 |
| **53** | Gucy1a2 | 1.63E-13 | 1.641346053 | 4.05E-09 | 0.75 | 0.173 |
| **54** | Slit3 | 1.89E-13 | 1.082207489 | 4.69E-09 | 0.99 | 0.654 |
| **55** | Col6a3 | 2.07E-13 | 1.394703572 | 5.15E-09 | 0.885 | 0.269 |
| **56** | Pcdh19 | 2.22E-13 | 1.770679465 | 5.52E-09 | 0.808 | 0.231 |
| **57** | Enpep | 3.55E-13 | 2.08810387 | 8.82E-09 | 0.615 | 0.096 |
| **58** | Bmpr1a | 3.58E-13 | 1.422233251 | 8.90E-09 | 0.692 | 0.125 |
| **59** | Dock10 | 4.59E-13 | 1.473562147 | 1.14E-08 | 0.837 | 0.231 |
| **60** | Kcnq4 | 6.73E-13 | 2.756445175 | 1.67E-08 | 0.433 | 0.01 |
| **61** | Carmn | 7.23E-13 | 1.822781792 | 1.80E-08 | 0.606 | 0.096 |
| **62** | Rnf152 | 9.69E-13 | 1.873772926 | 2.41E-08 | 0.644 | 0.125 |
| **63** | Gcnt2 | 9.94E-13 | 1.446229754 | 2.47E-08 | 0.798 | 0.221 |
| **64** | Arhgap32 | 1.46E-12 | 1.193913089 | 3.63E-08 | 0.76 | 0.192 |
| **65** | Map3k20 | 1.51E-12 | 1.302087674 | 3.76E-08 | 0.837 | 0.26 |
| **66** | Trpc3 | 1.60E-12 | 1.86726904 | 3.99E-08 | 0.567 | 0.077 |
| **67** | Anxa1 | 1.73E-12 | 1.553982636 | 4.30E-08 | 0.702 | 0.154 |
| **68** | Daam1 | 1.81E-12 | 1.283360292 | 4.50E-08 | 0.837 | 0.24 |
| **69** | Bgn | 2.04E-12 | 2.42863346 | 5.07E-08 | 0.615 | 0.125 |
| **70** | Rasl12 | 2.76E-12 | 2.235827423 | 6.86E-08 | 0.51 | 0.058 |
| **71** | Rbms3 | 3.16E-12 | 1.125852491 | 7.87E-08 | 0.933 | 0.346 |
| **72** | Kirrel | 3.99E-12 | 1.744040744 | 9.94E-08 | 0.692 | 0.163 |
| **73** | Pard3 | 6.39E-12 | 1.091928997 | 1.59E-07 | 0.846 | 0.279 |
| **74** | Gpc6 | 6.80E-12 | 1.653625341 | 1.69E-07 | 0.817 | 0.317 |
| **75** | Tuba1b | 6.98E-12 | 1.967547466 | 1.74E-07 | 0.644 | 0.144 |
| **76** | Rbbp4 | 1.01E-11 | 1.484513257 | 2.52E-07 | 0.596 | 0.106 |
| **77** | Cobll1 | 1.03E-11 | 1.256891185 | 2.57E-07 | 0.846 | 0.269 |
| **78** | Arhgap10 | 1.29E-11 | 1.221428433 | 3.20E-07 | 0.894 | 0.308 |
| **79** | Dock7 | 1.36E-11 | 1.549602669 | 3.38E-07 | 0.712 | 0.183 |

| | | | | | | |
|---|---|---|---|---|---|---|
| **80** | Nbea | 1.43E-11 | 1.006365965 | 3.56E-07 | 1 | 0.413 |
| **81** | Myh11 | 1.93E-11 | 1.936726425 | 4.80E-07 | 0.615 | 0.135 |
| **82** | Col6a2 | 2.10E-11 | 1.415196248 | 5.22E-07 | 0.721 | 0.173 |
| **83** | Arap2 | 2.20E-11 | 1.143873399 | 5.47E-07 | 0.779 | 0.221 |
| **84** | Il34 | 2.32E-11 | 2.26787989 | 5.77E-07 | 0.519 | 0.077 |
| **85** | Cyp4b1 | 2.45E-11 | 1.518534556 | 6.10E-07 | 0.846 | 0.365 |
| **86** | Serping1 | 2.64E-11 | 2.245860155 | 6.56E-07 | 0.51 | 0.067 |
| **87** | Col6a1 | 2.91E-11 | 1.082393767 | 7.24E-07 | 0.692 | 0.163 |
| **88** | B3galt1 | 2.99E-11 | 0.753516995 | 7.43E-07 | 0.76 | 0.212 |
| **89** | Prrx1 | 3.00E-11 | 1.485705167 | 7.47E-07 | 0.644 | 0.144 |
| **90** | Iqsec2 | 3.11E-11 | 1.275203207 | 7.75E-07 | 0.654 | 0.144 |
| **91** | Rgs4 | 3.56E-11 | 3.102427046 | 8.85E-07 | 0.423 | 0.029 |
| **92** | Fap | 3.57E-11 | 1.698009944 | 8.89E-07 | 0.596 | 0.106 |
| **93** | Dync1i2 | 4.44E-11 | 0.947769443 | 1.10E-06 | 0.692 | 0.154 |
| **94** | Prkg1 | 4.62E-11 | 0.9244632 | 1.15E-06 | 0.952 | 0.519 |
| **95** | Mfge8 | 5.96E-11 | 0.97546017 | 1.48E-06 | 0.712 | 0.183 |
| **96** | Vstm4 | 6.09E-11 | 3.479629992 | 1.51E-06 | 0.442 | 0.048 |
| **97** | Rtn4rl1 | 6.16E-11 | 1.675917046 | 1.53E-06 | 0.481 | 0.058 |
| **98** | Pawr | 7.04E-11 | 0.777426353 | 1.75E-06 | 0.894 | 0.298 |
| **99** | Aff3 | 7.04E-11 | 2.181509674 | 1.75E-06 | 0.596 | 0.135 |
| **100** | Pik3r1 | 7.25E-11 | 1.096979329 | 1.80E-06 | 0.952 | 0.394 |
| **101** | Gli3 | 7.51E-11 | 1.973628537 | 1.87E-06 | 0.548 | 0.096 |
| **102** | Vasp | 8.42E-11 | 1.728342037 | 2.09E-06 | 0.558 | 0.096 |
| **103** | Eif4a1 | 9.15E-11 | 1.459530549 | 2.28E-06 | 0.587 | 0.115 |
| **104** | Ehbp1 | 1.17E-10 | 1.232028525 | 2.91E-06 | 0.856 | 0.327 |
| **105** | Hnrnpul2 | 1.21E-10 | 1.414581841 | 3.02E-06 | 0.673 | 0.173 |
| **106** | Dok6 | 1.26E-10 | 2.048603204 | 3.13E-06 | 0.548 | 0.096 |
| **107** | Zeb2 | 1.31E-10 | 0.905947172 | 3.25E-06 | 0.971 | 0.567 |
| **108** | Ifitm3 | 1.57E-10 | 1.054567803 | 3.91E-06 | 0.731 | 0.212 |
| **109** | Peli2 | 1.78E-10 | 0.882665035 | 4.43E-06 | 0.885 | 0.279 |
| **110** | Snx18 | 1.87E-10 | 1.371208645 | 4.66E-06 | 0.606 | 0.125 |
| **111** | Cpne1 | 2.08E-10 | 1.213753836 | 5.17E-06 | 0.721 | 0.202 |
| **112** | Adgra3 | 2.11E-10 | 1.375940961 | 5.26E-06 | 0.567 | 0.106 |
| **113** | Itih5 | 2.26E-10 | 1.426769426 | 5.62E-06 | 0.779 | 0.279 |
| **114** | Robo1 | 2.39E-10 | 2.100013252 | 5.96E-06 | 0.548 | 0.125 |
| **115** | Pip5k1b | 2.68E-10 | 1.507853153 | 6.66E-06 | 0.529 | 0.087 |
| **116** | Kctd12 | 2.86E-10 | 0.942049563 | 7.12E-06 | 0.683 | 0.173 |
| **117** | Trarg1 | 3.23E-10 | 1.84757733 | 8.03E-06 | 0.462 | 0.058 |
| **118** | Dag1 | 3.27E-10 | 0.662726891 | 8.14E-06 | 0.76 | 0.212 |
| **119** | Edem3 | 3.33E-10 | 1.489618758 | 8.27E-06 | 0.596 | 0.125 |
| **120** | Extl3 | 3.39E-10 | 0.655754624 | 8.44E-06 | 0.731 | 0.183 |

| | | | | | | |
|---|---|---|---|---|---|---|
| **121** | Mnd1 | 3.72E-10 | 2.127046361 | 9.26E-06 | 0.433 | 0.048 |
| **122** | Gucy1b1 | 4.03E-10 | 1.238870285 | 1.00E-05 | 0.596 | 0.135 |
| **123** | Morrbid | 4.20E-10 | 1.294398064 | 1.05E-05 | 0.76 | 0.24 |
| **124** | Stxbp3 | 4.38E-10 | 1.067128563 | 1.09E-05 | 0.596 | 0.125 |
| **125** | Tnxb | 4.58E-10 | 0.711096387 | 1.14E-05 | 0.769 | 0.231 |
| **126** | Gas1 | 5.47E-10 | 2.235940491 | 1.36E-05 | 0.394 | 0.029 |
| **127** | C1qtnf1 | 5.69E-10 | 1.899350507 | 1.42E-05 | 0.462 | 0.067 |
| **128** | Leng8 | 5.78E-10 | 0.886981682 | 1.44E-05 | 0.692 | 0.173 |
| **129** | Txnip | 6.22E-10 | 1.084228029 | 1.55E-05 | 0.971 | 0.615 |
| **130** | Mylk | 6.48E-10 | 0.930450646 | 1.61E-05 | 0.769 | 0.24 |
| **131** | Myo1b | 6.58E-10 | 1.073976304 | 1.64E-05 | 0.933 | 0.462 |
| **132** | Polr2a | 6.59E-10 | 0.614842653 | 1.64E-05 | 0.596 | 0.125 |
| **133** | Rasgrp2 | 6.99E-10 | 0.872193932 | 1.74E-05 | 0.75 | 0.221 |
| **134** | Limd1 | 7.67E-10 | 1.150010258 | 1.91E-05 | 0.625 | 0.154 |
| **135** | Olfm2 | 7.94E-10 | 2.985732517 | 1.98E-05 | 0.413 | 0.048 |
| **136** | Rpl7 | 8.47E-10 | 1.484151218 | 2.11E-05 | 0.625 | 0.154 |
| **137** | Stk38l | 9.28E-10 | 1.591752007 | 2.31E-05 | 0.538 | 0.106 |
| **138** | Fblim1 | 9.90E-10 | 2.838841161 | 2.46E-05 | 0.404 | 0.038 |
| **139** | 4632427E13Ri | 1.04E-09 | 1.193185743 | 2.58E-05 | 0.577 | 0.115 |
| **140** | Sirt1 | 1.07E-09 | 2.557875441 | 2.67E-05 | 0.404 | 0.038 |
| **141** | Ppfia2 | 1.13E-09 | 2.037452024 | 2.82E-05 | 0.49 | 0.087 |
| **142** | Gipr | 1.18E-09 | 3.292196082 | 2.94E-05 | 0.394 | 0.038 |
| **143** | Plekhg3 | 1.21E-09 | 0.957944936 | 3.00E-05 | 0.654 | 0.154 |
| **144** | Slc30a5 | 1.25E-09 | 1.10382306 | 3.10E-05 | 0.452 | 0.058 |
| **145** | Timp3 | 1.34E-09 | 1.039879061 | 3.34E-05 | 1 | 0.673 |
| **146** | Kifc3 | 1.36E-09 | 0.889258483 | 3.38E-05 | 0.538 | 0.106 |
| **147** | Nap1l1 | 1.44E-09 | 1.375768446 | 3.60E-05 | 0.567 | 0.125 |
| **148** | Frmd4a | 1.49E-09 | 0.817410404 | 3.71E-05 | 0.856 | 0.298 |
| **149** | Eps8 | 1.53E-09 | 0.831356041 | 3.80E-05 | 0.894 | 0.356 |
| **150** | Irf2bp2 | 1.56E-09 | 0.668982439 | 3.89E-05 | 0.663 | 0.173 |
| **151** | Arl8b | 1.58E-09 | 1.098976673 | 3.92E-05 | 0.596 | 0.135 |
| **152** | Dazap1 | 1.60E-09 | 1.401147468 | 3.97E-05 | 0.481 | 0.077 |
| **153** | Myl9 | 1.62E-09 | 3.190094136 | 4.02E-05 | 0.385 | 0.038 |
| **154** | Fam110b | 1.73E-09 | 1.120753725 | 4.30E-05 | 0.529 | 0.106 |
| **155** | Marcks | 1.79E-09 | 1.265315443 | 4.46E-05 | 0.837 | 0.337 |
| **156** | Zdhhc14 | 1.81E-09 | 0.767543982 | 4.50E-05 | 0.769 | 0.25 |
| **157** | Ctdsp2 | 1.82E-09 | 0.952532397 | 4.53E-05 | 0.721 | 0.192 |
| **158** | Pdzrn3 | 1.83E-09 | 1.016083511 | 4.56E-05 | 0.654 | 0.183 |
| **159** | Mark1 | 1.91E-09 | 2.77861856 | 4.74E-05 | 0.365 | 0.029 |
| **160** | Slc4a4 | 2.20E-09 | 0.69694785 | 5.47E-05 | 0.644 | 0.173 |
| **161** | Astn2 | 2.51E-09 | 2.78984893 | 6.24E-05 | 0.413 | 0.058 |

| | | | | | | |
|---|---|---|---|---|---|---|
| **162** | Prkar1a | 2.66E-09 | 0.832009675 | 6.62E-05 | 0.837 | 0.288 |
| **163** | Slc12a2 | 2.99E-09 | 1.360686936 | 7.44E-05 | 0.481 | 0.087 |
| **164** | Ppp1r12b | 3.29E-09 | 0.823824092 | 8.18E-05 | 0.827 | 0.279 |
| **165** | Mamdc2 | 3.43E-09 | 0.901577609 | 8.54E-05 | 0.5 | 0.096 |
| **166** | Rpl35a | 3.53E-09 | 0.711463958 | 8.77E-05 | 0.856 | 0.288 |
| **167** | Cspg4 | 3.70E-09 | 2.319332926 | 9.22E-05 | 0.385 | 0.038 |
| **168** | Rarres2 | 3.71E-09 | 1.789592515 | 9.24E-05 | 0.51 | 0.106 |
| **169** | Irs1 | 3.77E-09 | 1.424427518 | 9.37E-05 | 0.654 | 0.192 |
| **170** | Rps12 | 3.81E-09 | 0.914013752 | 9.48E-05 | 0.798 | 0.288 |
| **171** | Psma3 | 3.96E-09 | 0.863870487 | 9.85E-05 | 0.635 | 0.163 |
| **172** | Trim28 | 4.08E-09 | 5.294968345 | 0.000101506 | 0.288 | 0 |
| **173** | Klf9 | 4.26E-09 | 0.923553408 | 0.000106079 | 0.894 | 0.346 |
| **174** | Nbr1 | 4.54E-09 | 0.809431475 | 0.00011295 | 0.625 | 0.154 |
| **175** | Mpp5 | 4.88E-09 | 0.523233704 | 0.000121461 | 0.587 | 0.135 |
| **176** | Tpm1 | 5.00E-09 | 1.008987273 | 0.000124433 | 0.731 | 0.231 |
| **177** | Bcl9l | 5.13E-09 | 0.759518175 | 0.000127628 | 0.596 | 0.144 |
| **178** | Dtx3 | 5.38E-09 | 1.384854475 | 0.000133889 | 0.587 | 0.154 |
| **179** | Aldh2 | 5.46E-09 | 1.074374818 | 0.000135896 | 0.538 | 0.115 |
| **180** | Atf2 | 5.52E-09 | 0.651831045 | 0.000137464 | 0.817 | 0.26 |
| **181** | Emc7 | 5.55E-09 | 1.591336795 | 0.000138121 | 0.5 | 0.096 |
| **182** | Npepps | 5.80E-09 | 0.506602431 | 0.00014423 | 0.712 | 0.212 |
| **183** | Kank1 | 5.84E-09 | 1.286248336 | 0.000145374 | 0.5 | 0.096 |
| **184** | Atp1b2 | 6.14E-09 | 2.282192924 | 0.00015265 | 0.375 | 0.038 |
| **185** | Zdhhc5 | 6.18E-09 | 2.809764979 | 0.00015379 | 0.356 | 0.029 |
| **186** | Cfl1 | 6.60E-09 | 1.002544132 | 0.000164132 | 0.683 | 0.202 |
| **187** | Crppa | 6.69E-09 | 1.890074363 | 0.000166512 | 0.462 | 0.087 |
| **188** | Lamb1 | 7.08E-09 | 0.587638167 | 0.000176038 | 0.712 | 0.183 |
| **189** | P3h2 | 7.13E-09 | 0.853608315 | 0.000177354 | 0.606 | 0.163 |
| **190** | St3gal2 | 7.17E-09 | 0.795865783 | 0.000178482 | 0.587 | 0.135 |
| **191** | Rplp2 | 7.25E-09 | 0.955623554 | 0.000180411 | 0.817 | 0.327 |
| **192** | Lbh | 7.46E-09 | 1.471135274 | 0.000185715 | 0.452 | 0.077 |
| **193** | Prdm10 | 7.84E-09 | 1.096326469 | 0.000195116 | 0.423 | 0.058 |
| **194** | Gli2 | 8.33E-09 | 2.936740606 | 0.000207292 | 0.365 | 0.038 |
| **195** | Rps24 | 8.39E-09 | 0.770511951 | 0.000208812 | 0.875 | 0.356 |
| **196** | Ngf | 8.82E-09 | 1.277804356 | 0.000219475 | 0.529 | 0.115 |
| **197** | Pear1 | 8.88E-09 | 1.120323805 | 0.000221025 | 0.644 | 0.183 |
| **198** | Antxr1 | 8.88E-09 | 0.928469054 | 0.000221053 | 0.529 | 0.106 |
| **199** | Jag1 | 9.51E-09 | 0.620395721 | 0.000236715 | 0.654 | 0.173 |
| **200** | Ssb | 9.54E-09 | 1.078585776 | 0.00023725 | 0.606 | 0.154 |
| **201** | Rhob | 9.60E-09 | 0.702878772 | 0.000238774 | 0.788 | 0.269 |
| **202** | Rplp1 | 1.03E-08 | 0.997967377 | 0.000256711 | 0.875 | 0.346 |

| | | | | | | |
|---|---|---|---|---|---|---|
| **203** | Fzd4 | 1.04E-08 | 1.078831751 | 0.000257612 | 0.846 | 0.356 |
| **204** | Zfp704 | 1.09E-08 | 0.996561098 | 0.000270641 | 0.673 | 0.202 |
| **205** | Ndrg2 | 1.12E-08 | 2.478177166 | 0.000278655 | 0.423 | 0.067 |
| **206** | Zbtb38 | 1.15E-08 | 1.108511982 | 0.000286755 | 0.587 | 0.144 |
| **207** | Col5a1 | 1.20E-08 | 1.043117735 | 0.00029949 | 0.644 | 0.183 |
| **208** | Ccdc58 | 1.21E-08 | 2.645720426 | 0.000300213 | 0.327 | 0.019 |
| **209** | Cyp20a1 | 1.23E-08 | 1.398811066 | 0.000305604 | 0.49 | 0.096 |
| **210** | Vta1 | 1.25E-08 | 1.107834241 | 0.000310784 | 0.548 | 0.125 |
| **211** | Adcy6 | 1.31E-08 | 1.142567002 | 0.000326098 | 0.548 | 0.125 |
| **212** | Ghitm | 1.34E-08 | 1.36373157 | 0.000334606 | 0.471 | 0.087 |
| **213** | Elavl1 | 1.37E-08 | 0.993170101 | 0.0003406 | 0.587 | 0.144 |
| **214** | Brd2 | 1.38E-08 | 1.290595839 | 0.000342927 | 0.596 | 0.173 |
| **215** | Spon1 | 1.38E-08 | 0.872169405 | 0.000344276 | 0.683 | 0.212 |
| **216** | Adamts2 | 1.39E-08 | 1.108965859 | 0.00034602 | 0.558 | 0.125 |
| **217** | Shroom3 | 1.42E-08 | 1.123164354 | 0.000352312 | 0.529 | 0.115 |
| **218** | Tmem131l | 1.43E-08 | 1.551581562 | 0.000356437 | 0.462 | 0.087 |
| **219** | Ift43 | 1.46E-08 | 1.333043141 | 0.000362704 | 0.433 | 0.067 |
| **220** | Rpl37a | 1.47E-08 | 0.959419995 | 0.000365239 | 0.837 | 0.346 |
| **221** | Nop53 | 1.55E-08 | 1.055770634 | 0.000386826 | 0.452 | 0.077 |
| **222** | Rpl17 | 1.56E-08 | 0.77035969 | 0.000387366 | 0.673 | 0.221 |
| **223** | Bzw2 | 1.62E-08 | 0.867808557 | 0.000402303 | 0.596 | 0.154 |
| **224** | Nab1 | 1.65E-08 | 0.471932211 | 0.000409479 | 0.615 | 0.163 |
| **225** | Tnks | 1.66E-08 | 1.024317673 | 0.000411912 | 0.827 | 0.317 |
| **226** | Pde1a | 1.66E-08 | 1.11306537 | 0.000412191 | 0.596 | 0.173 |
| **227** | Nr1d1 | 1.71E-08 | 1.297338788 | 0.000424413 | 0.471 | 0.087 |
| **228** | Thbs2 | 1.78E-08 | 1.581878231 | 0.000443993 | 0.385 | 0.048 |
| **229** | Atp9a | 1.92E-08 | 0.812695681 | 0.000478168 | 0.75 | 0.24 |
| **230** | Tsc1 | 2.01E-08 | 1.830827594 | 0.000501337 | 0.404 | 0.058 |
| **231** | Kcnj8 | 2.03E-08 | 3.46125586 | 0.000505957 | 0.346 | 0.038 |
| **232** | Actb | 2.08E-08 | 0.718105502 | 0.000518587 | 0.962 | 0.49 |
| **233** | Rem1 | 2.10E-08 | 2.162608238 | 0.00052147 | 0.365 | 0.038 |
| **234** | Smo | 2.11E-08 | 2.851259789 | 0.000526204 | 0.317 | 0.019 |
| **235** | Pura | 2.12E-08 | 0.792931076 | 0.00052738 | 0.692 | 0.192 |
| **236** | Stk39 | 2.13E-08 | 0.46541485 | 0.000530059 | 0.74 | 0.231 |
| **237** | S100a6 | 2.18E-08 | 0.623704075 | 0.000543057 | 0.779 | 0.269 |
| **238** | Ptms | 2.27E-08 | 0.896399079 | 0.000564516 | 0.75 | 0.288 |
| **239** | Cpe | 2.33E-08 | 1.889266999 | 0.000579755 | 0.375 | 0.048 |
| **240** | Metap2 | 2.33E-08 | 1.358658011 | 0.000580246 | 0.519 | 0.125 |
| **241** | Fam124a | 2.37E-08 | 1.919926916 | 0.000589022 | 0.413 | 0.067 |
| **242** | Zfhx3 | 2.42E-08 | 0.402650251 | 0.000602277 | 0.885 | 0.346 |
| **243** | Kdelr2 | 2.45E-08 | 1.904662001 | 0.000609519 | 0.337 | 0.029 |

| | | | | | | |
|---|---|---|---|---|---|---|
| **244** | Rps15 | 2.46E-08 | 0.603652444 | 0.000612408 | 0.721 | 0.221 |
| **245** | Mapre1 | 2.49E-08 | 0.854143364 | 0.000619381 | 0.558 | 0.135 |
| **246** | 8-Sep | 2.50E-08 | 1.343061105 | 0.000621337 | 0.596 | 0.173 |
| **247** | Hnrnpa0 | 2.53E-08 | 1.469187571 | 0.000629111 | 0.529 | 0.125 |
| **248** | Igfbp7 | 2.58E-08 | 0.771239527 | 0.000641627 | 0.981 | 0.577 |
| **249** | Scp2 | 2.80E-08 | 0.984133443 | 0.000696398 | 0.577 | 0.144 |
| **250** | Sqstm1 | 2.81E-08 | 0.923995125 | 0.000700045 | 0.625 | 0.173 |
| **251** | Sort1 | 2.82E-08 | 1.346681932 | 0.000702875 | 0.529 | 0.125 |
| **252** | Jph1 | 2.89E-08 | 2.143752689 | 0.000719555 | 0.356 | 0.038 |
| **253** | Tbc1d1 | 2.90E-08 | 0.801759457 | 0.000721529 | 0.779 | 0.26 |
| **254** | Cnbp | 3.00E-08 | 0.645079781 | 0.000745295 | 0.779 | 0.279 |
| **255** | Tspan12 | 3.05E-08 | 0.936607901 | 0.000757664 | 0.769 | 0.269 |
| **256** | Eef1b2 | 3.06E-08 | 1.223060323 | 0.00076052 | 0.519 | 0.125 |
| **257** | Matr3 | 3.12E-08 | 0.986653574 | 0.000776882 | 0.625 | 0.173 |
| **258** | Sp1 | 3.14E-08 | 0.749973376 | 0.00078111 | 0.673 | 0.202 |
| **259** | Hnrnpk | 3.16E-08 | 0.705902844 | 0.000786823 | 0.76 | 0.269 |
| **260** | Mxra7 | 3.36E-08 | 1.244717729 | 0.000836035 | 0.529 | 0.125 |
| **261** | Akap7 | 3.38E-08 | 0.997917811 | 0.000841401 | 0.721 | 0.24 |
| **262** | Impad1 | 3.50E-08 | 1.035352954 | 0.000869773 | 0.452 | 0.087 |
| **263** | Lims1 | 3.52E-08 | 0.720729369 | 0.000875757 | 0.962 | 0.471 |
| **264** | Sec23a | 3.55E-08 | 1.307628316 | 0.000884484 | 0.452 | 0.087 |
| **265** | Cog3 | 3.57E-08 | 1.085396034 | 0.000887278 | 0.538 | 0.125 |
| **266** | Lhfpl3 | 3.57E-08 | 1.634199856 | 0.000889107 | 0.394 | 0.058 |
| **267** | Jund | 3.74E-08 | 0.97855805 | 0.000930518 | 0.692 | 0.221 |
| **268** | Myof | 3.77E-08 | 0.596492511 | 0.00093725 | 0.615 | 0.163 |
| **269** | Plxdc2 | 3.83E-08 | 1.076927668 | 0.000951908 | 0.798 | 0.317 |
| **270** | Rps25 | 3.85E-08 | 0.860183877 | 0.000957386 | 0.51 | 0.115 |
| **271** | Cmss1 | 3.85E-08 | -1.45620448 | 0.000959153 | 1 | 0.981 |
| **272** | Taok2 | 3.86E-08 | 0.952439586 | 0.000959691 | 0.577 | 0.144 |
| **273** | Tmem87a | 3.96E-08 | 0.819247293 | 0.000984202 | 0.673 | 0.202 |
| **274** | Peg3 | 3.96E-08 | 2.252221694 | 0.000985889 | 0.385 | 0.058 |
| **275** | Pgap2 | 3.98E-08 | 1.993046016 | 0.000989603 | 0.394 | 0.058 |
| **276** | Cald1 | 4.00E-08 | 0.832128315 | 0.000994045 | 0.952 | 0.452 |
| **277** | Trpv2 | 4.01E-08 | 4.440846724 | 0.000996497 | 0.279 | 0.01 |
| **278** | Ptpn9 | 4.13E-08 | 0.68864007 | 0.001026832 | 0.663 | 0.202 |
| **279** | Efl1 | 4.17E-08 | 1.151531349 | 0.001036738 | 0.49 | 0.106 |
| **280** | Rpl13 | 4.19E-08 | 0.916861933 | 0.001041483 | 0.798 | 0.288 |
| **281** | Bcap31 | 4.21E-08 | 1.417989784 | 0.001047347 | 0.558 | 0.154 |
| **282** | Ubb | 4.28E-08 | 1.057799011 | 0.001064037 | 0.663 | 0.221 |
| **283** | Spred1 | 4.32E-08 | 0.602825291 | 0.00107524 | 0.779 | 0.279 |
| **284** | Cdkn1b | 4.37E-08 | 1.580308979 | 0.001087591 | 0.413 | 0.067 |

| | | | | | | |
|---|---|---|---|---|---|---|
| **285** | Rpl21 | 4.37E-08 | 0.965774868 | 0.001088158 | 0.712 | 0.269 |
| **286** | Slc39a7 | 4.81E-08 | 1.294047128 | 0.001196857 | 0.423 | 0.067 |
| **287** | Ehd1 | 4.89E-08 | 2.051008858 | 0.001217596 | 0.413 | 0.077 |
| **288** | Hrct1 | 4.93E-08 | 1.780320024 | 0.001227852 | 0.413 | 0.067 |
| **289** | Tbx18 | 4.99E-08 | 2.295385365 | 0.001241688 | 0.365 | 0.048 |
| **290** | Prpsap1 | 5.08E-08 | 2.368111409 | 0.001265009 | 0.346 | 0.038 |
| **291** | Tgs1 | 5.22E-08 | 1.358184891 | 0.001297784 | 0.452 | 0.087 |
| **292** | Usp49 | 5.28E-08 | 1.694563972 | 0.001313528 | 0.346 | 0.038 |
| **293** | Ppp2r2a | 5.56E-08 | 0.508602224 | 0.001383281 | 0.817 | 0.298 |
| **294** | Mbd2 | 5.57E-08 | 0.686796694 | 0.001386678 | 0.856 | 0.346 |
| **295** | Angptl4 | 5.83E-08 | 0.817800048 | 0.001450822 | 0.74 | 0.25 |
| **296** | Dnajc6 | 5.96E-08 | 6.280845997 | 0.00148398 | 0.25 | 0 |
| **297** | Ggps1 | 6.08E-08 | 1.672643934 | 0.001512433 | 0.423 | 0.077 |
| **298** | Serf2 | 6.11E-08 | 0.884305011 | 0.001519163 | 0.663 | 0.221 |
| **299** | Col5a2 | 6.23E-08 | 0.452257569 | 0.001550739 | 0.731 | 0.25 |
| **300** | Rcsd1 | 6.27E-08 | 0.667566674 | 0.001559794 | 0.798 | 0.298 |
| **301** | Capn15 | 6.33E-08 | 0.550919741 | 0.001574046 | 0.365 | 0.048 |
| **302** | Tug1 | 6.43E-08 | 1.510750122 | 0.001599204 | 0.404 | 0.067 |
| **303** | Pde5a | 6.66E-08 | 1.041264394 | 0.001657744 | 0.442 | 0.087 |
| **304** | Mmp14 | 6.73E-08 | 1.331179286 | 0.001675464 | 0.471 | 0.106 |
| **305** | Anxa3 | 6.81E-08 | 0.547523791 | 0.001694123 | 0.692 | 0.231 |
| **306** | Rps8 | 6.83E-08 | 0.763762372 | 0.001699941 | 0.837 | 0.327 |
| **307** | Dlst | 6.85E-08 | 0.981311769 | 0.001704956 | 0.519 | 0.115 |
| **308** | Nipa2 | 6.92E-08 | 0.811904263 | 0.001722108 | 0.471 | 0.096 |
| **309** | Zcchc24 | 7.00E-08 | 0.912602282 | 0.001741304 | 0.654 | 0.202 |
| **310** | Vti1b | 7.00E-08 | 0.815591365 | 0.001742001 | 0.433 | 0.077 |
| **311** | Aoc3 | 7.24E-08 | 1.304567991 | 0.001800174 | 0.51 | 0.125 |
| **312** | Angpt1 | 7.24E-08 | 1.253135273 | 0.00180131 | 0.433 | 0.077 |
| **313** | Cul1 | 7.28E-08 | 0.705722725 | 0.001811649 | 0.798 | 0.279 |
| **314** | Hspa5 | 7.51E-08 | 0.919919651 | 0.001869509 | 0.654 | 0.202 |
| **315** | Prkca | 7.51E-08 | 0.624278887 | 0.001869509 | 0.663 | 0.192 |
| **316** | Hif1a | 7.67E-08 | 0.519681279 | 0.00190763 | 0.558 | 0.144 |
| **317** | Tmem65 | 7.74E-08 | 0.470349592 | 0.001926914 | 0.635 | 0.192 |
| **318** | Ptpn13 | 7.76E-08 | 1.469199284 | 0.001931683 | 0.49 | 0.106 |
| **319** | Bnip3l | 7.83E-08 | 1.218295689 | 0.001948844 | 0.519 | 0.135 |
| **320** | Rplp0 | 7.83E-08 | 0.926446266 | 0.001949423 | 0.596 | 0.173 |
| **321** | Uqcrq | 8.12E-08 | 1.296247266 | 0.002019695 | 0.452 | 0.096 |
| **322** | Flna | 8.32E-08 | 1.135795946 | 0.002070927 | 0.615 | 0.183 |
| **323** | Anxa2 | 8.34E-08 | 0.865652129 | 0.002074673 | 0.644 | 0.202 |
| **324** | Ccdc3 | 8.39E-08 | 0.90598708 | 0.002088316 | 0.442 | 0.087 |
| **325** | Csnk2a2 | 8.61E-08 | 0.841434485 | 0.002143051 | 0.606 | 0.163 |

| | | | | | | |
|---|---|---|---|---|---|---|
| **326** | Khdrbs1 | 8.68E-08 | 0.512371837 | 0.002159094 | 0.75 | 0.26 |
| **327** | Rhoj | 8.74E-08 | 0.812433575 | 0.002175173 | 0.971 | 0.481 |
| **328** | Rsl1d1 | 8.84E-08 | 1.320413764 | 0.002198437 | 0.298 | 0.019 |
| **329** | Eef1a1 | 8.91E-08 | 0.840996605 | 0.002217046 | 0.952 | 0.471 |
| **330** | Cox7a2l | 9.10E-08 | 1.413280523 | 0.002265282 | 0.423 | 0.077 |
| **331** | Ell | 9.22E-08 | 1.340204714 | 0.002294324 | 0.404 | 0.067 |
| **332** | Tnpo1 | 9.58E-08 | 0.55490698 | 0.00238373 | 0.856 | 0.317 |
| **333** | Cog7 | 9.61E-08 | 3.317124667 | 0.002390512 | 0.288 | 0.019 |
| **334** | Map3k2 | 9.61E-08 | 0.614280982 | 0.002391953 | 0.76 | 0.269 |
| **335** | Setbp1 | 9.67E-08 | 0.752132227 | 0.002404902 | 0.692 | 0.24 |
| **336** | Hspa4 | 9.75E-08 | 0.902164253 | 0.002425755 | 0.615 | 0.183 |
| **337** | Arcn1 | 9.80E-08 | 0.853756305 | 0.002438274 | 0.635 | 0.183 |
| **338** | Sar1a | 9.85E-08 | 1.460336786 | 0.002450303 | 0.433 | 0.087 |
| **339** | Piezo2 | 9.85E-08 | -0.21445779 | 0.002450524 | 0.519 | 0.135 |
| **340** | Sfr1 | 1.01E-07 | 1.283281775 | 0.002520165 | 0.385 | 0.058 |
| **341** | Rps28 | 1.04E-07 | 0.86099993 | 0.002580729 | 0.846 | 0.356 |
| **342** | Adcy3 | 1.04E-07 | 1.76859739 | 0.002586425 | 0.356 | 0.048 |
| **343** | Rsrc2 | 1.06E-07 | 0.538316271 | 0.002632976 | 0.721 | 0.231 |
| **344** | Ino80d | 1.07E-07 | 0.641234509 | 0.002654651 | 0.788 | 0.288 |
| **345** | Anks3 | 1.11E-07 | 0.754853407 | 0.002766517 | 0.337 | 0.038 |
| **346** | Clic1 | 1.11E-07 | 0.708780068 | 0.002772721 | 0.51 | 0.115 |
| **347** | Ahcyl1 | 1.14E-07 | 0.789183809 | 0.002828833 | 0.635 | 0.192 |
| **348** | Sh3kbp1 | 1.14E-07 | 0.75286443 | 0.002846295 | 0.663 | 0.212 |
| **349** | Rhoq | 1.16E-07 | 1.307643525 | 0.002889377 | 0.413 | 0.077 |
| **350** | Tpm4 | 1.18E-07 | 0.588418516 | 0.002933182 | 0.798 | 0.279 |
| **351** | Pde8b | 1.18E-07 | 1.880948321 | 0.002948168 | 0.375 | 0.058 |
| **352** | Uqcrh | 1.21E-07 | 0.615476848 | 0.003014939 | 0.596 | 0.183 |
| **353** | Eif4ebp2 | 1.23E-07 | 0.929179547 | 0.003049168 | 0.635 | 0.202 |
| **354** | Hnrnph3 | 1.24E-07 | 0.459233137 | 0.003086147 | 0.692 | 0.212 |
| **355** | Ammecr1l | 1.27E-07 | 0.590279856 | 0.003159543 | 0.423 | 0.077 |
| **356** | Cd63 | 1.28E-07 | 0.798759406 | 0.003188709 | 0.654 | 0.212 |
| **357** | Col3a1 | 1.34E-07 | 0.602808497 | 0.003329135 | 0.827 | 0.346 |
| **358** | Adamts15 | 1.34E-07 | 1.877096878 | 0.003331746 | 0.375 | 0.058 |
| **359** | Gpx4 | 1.35E-07 | 0.720104416 | 0.003350119 | 0.635 | 0.192 |
| **360** | Mlf2 | 1.37E-07 | 1.140734916 | 0.003413462 | 0.413 | 0.077 |
| **361** | Specc1l | 1.37E-07 | 0.579636897 | 0.003416928 | 0.779 | 0.26 |
| **362** | Rps29 | 1.37E-07 | 0.730107458 | 0.003418749 | 0.817 | 0.346 |
| **363** | Smg7 | 1.38E-07 | 0.791995697 | 0.003438601 | 0.692 | 0.221 |
| **364** | Tmem176b | 1.39E-07 | 1.472089884 | 0.00345527 | 0.433 | 0.087 |
| **365** | Rad54l2 | 1.42E-07 | 0.462877561 | 0.003524601 | 0.587 | 0.163 |
| **366** | Suz12 | 1.44E-07 | 0.66387336 | 0.003575782 | 0.587 | 0.163 |

| | | | | | | |
|---|---|---|---|---|---|---|
| **367** | Hoxc6 | 1.45E-07 | 2.138686835 | 0.003598594 | 0.346 | 0.048 |
| **368** | Gm29683 | 1.49E-07 | 2.429934817 | 0.00371807 | 0.288 | 0.019 |
| **369** | Tmed10 | 1.49E-07 | 0.808614651 | 0.003719565 | 0.692 | 0.231 |
| **370** | Tshz3 | 1.55E-07 | 0.930728963 | 0.003848003 | 0.548 | 0.144 |
| **371** | Prkar1b | 1.56E-07 | 1.973164996 | 0.003869191 | 0.288 | 0.019 |
| **372** | Bcl2l1 | 1.56E-07 | 0.19730942 | 0.003870537 | 0.683 | 0.221 |
| **373** | Creb3l2 | 1.56E-07 | 0.379086832 | 0.003875551 | 0.769 | 0.279 |
| **374** | Rps20 | 1.61E-07 | 0.409351826 | 0.004007867 | 0.817 | 0.317 |
| **375** | Naalad2 | 1.61E-07 | 1.199108555 | 0.004012122 | 0.404 | 0.067 |
| **376** | Myh9 | 1.64E-07 | 0.672336437 | 0.004087757 | 1 | 0.673 |
| **377** | Psmd1 | 1.64E-07 | 0.994933138 | 0.004090519 | 0.606 | 0.183 |
| **378** | Cdc5l | 1.65E-07 | 1.017490813 | 0.004111367 | 0.538 | 0.154 |
| **379** | Srpr | 1.66E-07 | 1.274091045 | 0.004121879 | 0.548 | 0.154 |
| **380** | Tril | 1.67E-07 | 2.095489927 | 0.00414582 | 0.327 | 0.038 |
| **381** | Etfdh | 1.67E-07 | 1.446824636 | 0.004150258 | 0.337 | 0.038 |
| **382** | Pcif1 | 1.68E-07 | 2.192729105 | 0.004189453 | 0.288 | 0.019 |
| **383** | Rpl41 | 1.74E-07 | 0.726552691 | 0.004323413 | 0.894 | 0.452 |
| **384** | Calu | 1.76E-07 | 0.677708094 | 0.004384801 | 0.644 | 0.192 |
| **385** | Hnrnpul1 | 1.78E-07 | 0.612120596 | 0.004421008 | 0.596 | 0.163 |
| **386** | Max | 1.78E-07 | -0.19241056 | 0.004438956 | 0.538 | 0.135 |
| **387** | Eefsec | 1.79E-07 | 0.9287522 | 0.004445197 | 0.481 | 0.106 |
| **388** | Ikbip | 1.79E-07 | 1.390083466 | 0.004464786 | 0.346 | 0.048 |
| **389** | Usp14 | 1.83E-07 | 1.603012062 | 0.004552467 | 0.394 | 0.067 |
| **390** | Mindy2 | 1.86E-07 | 0.536804651 | 0.004616038 | 0.519 | 0.125 |
| **391** | Rpl23a | 1.92E-07 | 0.790204103 | 0.00478419 | 0.625 | 0.202 |
| **392** | Wls | 1.94E-07 | 0.710402707 | 0.004821573 | 0.606 | 0.173 |
| **393** | Itpripl2 | 1.97E-07 | 0.804391863 | 0.004896264 | 0.625 | 0.192 |
| **394** | Akap8 | 2.01E-07 | 0.788470537 | 0.004991203 | 0.413 | 0.077 |
| **395** | Klhl29 | 2.01E-07 | 1.303784723 | 0.00501251 | 0.433 | 0.087 |
| **396** | Tnrc6a | 2.02E-07 | 0.584137425 | 0.005024552 | 0.904 | 0.385 |
| **397** | Mms19 | 2.11E-07 | 1.421391551 | 0.005242946 | 0.413 | 0.077 |
| **398** | Plekhh2 | 2.11E-07 | 1.187799595 | 0.00524598 | 0.423 | 0.087 |
| **399** | Ahdc1 | 2.17E-07 | 0.567690606 | 0.005387748 | 0.577 | 0.154 |
| **400** | Adcy9 | 2.19E-07 | 0.515266191 | 0.005438944 | 0.702 | 0.231 |
| **401** | Cdc42ep4 | 2.22E-07 | 0.628394137 | 0.005529285 | 0.442 | 0.096 |
| **402** | Lix1l | 2.25E-07 | 1.763306678 | 0.005600351 | 0.365 | 0.058 |
| **403** | Ncoa6 | 2.34E-07 | 0.450768824 | 0.005824609 | 0.788 | 0.26 |
| **404** | Tmem189 | 2.42E-07 | 2.402917423 | 0.006033532 | 0.298 | 0.029 |
| **405** | Parp11 | 2.47E-07 | 0.908416326 | 0.006134115 | 0.481 | 0.115 |
| **406** | Tspan15 | 2.48E-07 | 0.983814293 | 0.006161421 | 0.606 | 0.192 |
| **407** | Gpx8 | 2.49E-07 | 2.730869257 | 0.006190805 | 0.317 | 0.038 |

| | | | | | | |
|---|---|---|---|---|---|---|
| **408** | Tmem260 | 2.49E-07 | 1.068081986 | 0.006205401 | 0.519 | 0.135 |
| **409** | Scai | 2.50E-07 | 0.414229007 | 0.006213111 | 0.596 | 0.163 |
| **410** | Pdap1 | 2.53E-07 | 2.109854789 | 0.006297781 | 0.356 | 0.058 |
| **411** | Rpl22 | 2.56E-07 | 0.645639288 | 0.006381014 | 0.615 | 0.192 |
| **412** | Oip5os1 | 2.56E-07 | 0.582970048 | 0.006381476 | 0.75 | 0.269 |
| **413** | Myo7a | 2.61E-07 | 1.25837696 | 0.00648805 | 0.346 | 0.048 |
| **414** | Lrpap1 | 2.63E-07 | 1.556268181 | 0.006540588 | 0.365 | 0.058 |
| **415** | Rap2c | 2.73E-07 | 2.076260383 | 0.006804852 | 0.327 | 0.038 |
| **416** | Prrg1 | 2.75E-07 | 0.960789125 | 0.006834803 | 0.644 | 0.221 |
| **417** | Fzd1 | 2.77E-07 | 4.222941381 | 0.006887573 | 0.25 | 0.01 |
| **418** | Src | 2.77E-07 | 3.955478915 | 0.006887573 | 0.25 | 0.01 |
| **419** | Ccz1 | 2.78E-07 | 0.939365655 | 0.006918411 | 0.413 | 0.077 |
| **420** | Mob1b | 2.81E-07 | 0.980764 | 0.006999978 | 0.51 | 0.135 |
| **421** | 1110059E24Ri | 2.83E-07 | 0.740989828 | 0.007053436 | 0.51 | 0.125 |
| **422** | Elf4 | 2.86E-07 | 0.510543599 | 0.007104078 | 0.51 | 0.135 |
| **423** | Arpc4 | 2.92E-07 | 0.963323048 | 0.007263422 | 0.413 | 0.077 |
| **424** | Thra | 2.93E-07 | 0.677131537 | 0.00729609 | 0.692 | 0.24 |
| **425** | Cycs | 2.96E-07 | 1.883219381 | 0.007360735 | 0.356 | 0.058 |
| **426** | Myadm | 3.05E-07 | 0.598016189 | 0.007589719 | 0.837 | 0.327 |
| **427** | 4930523C07Ri | 3.07E-07 | 0.857339809 | 0.007639062 | 0.721 | 0.269 |
| **428** | Cul2 | 3.09E-07 | 1.006630144 | 0.007693266 | 0.442 | 0.096 |
| **429** | Smc3 | 3.12E-07 | 0.129478906 | 0.007775276 | 0.558 | 0.154 |
| **430** | Qrich1 | 3.14E-07 | 0.407448396 | 0.007813269 | 0.635 | 0.202 |
| **431** | Egflam | 3.21E-07 | 0.774812367 | 0.007980728 | 0.596 | 0.183 |
| **432** | Tasor2 | 3.23E-07 | 0.686464741 | 0.008048918 | 0.625 | 0.192 |
| **433** | Calm2 | 3.27E-07 | 0.72492897 | 0.008135024 | 0.683 | 0.221 |
| **434** | Atn1 | 3.27E-07 | 0.703279085 | 0.008135345 | 0.49 | 0.115 |
| **435** | Nr2f2 | 3.30E-07 | 0.759820796 | 0.008211234 | 0.673 | 0.24 |
| **436** | Glyr1 | 3.33E-07 | 0.560898342 | 0.008291125 | 0.625 | 0.183 |
| **437** | Dip2a | 3.37E-07 | 0.798485442 | 0.008384363 | 0.51 | 0.144 |
| **438** | Mon2 | 3.39E-07 | 0.571868663 | 0.008425506 | 0.731 | 0.26 |
| **439** | Pnrc1 | 3.39E-07 | 0.852490722 | 0.008428971 | 0.558 | 0.163 |
| **440** | Ghr | 3.40E-07 | -0.89035625 | 0.00845342 | 0.99 | 0.971 |
| **441** | Myl12a | 3.44E-07 | 0.744043743 | 0.008553473 | 0.596 | 0.183 |
| **442** | Prune2 | 3.55E-07 | 1.473685078 | 0.008824682 | 0.442 | 0.106 |
| **443** | Ankrd10 | 3.56E-07 | 0.425993153 | 0.008862782 | 0.558 | 0.144 |
| **444** | Pdzd8 | 3.58E-07 | 0.390216816 | 0.00890403 | 0.577 | 0.163 |
| **445** | Phf12 | 3.63E-07 | 0.232375621 | 0.009027526 | 0.673 | 0.212 |
| **446** | Arf1 | 3.63E-07 | 0.676330239 | 0.009040238 | 0.615 | 0.202 |
| **447** | Lasp1 | 3.70E-07 | 0.643234284 | 0.009210177 | 0.663 | 0.221 |
| **448** | Sdc2 | 3.72E-07 | 1.243339156 | 0.009254224 | 0.471 | 0.115 |

| | | | | | | |
|---|---|---|---|---|---|---|
| **449** | Ythdf2 | 3.81E-07 | 0.631234205 | 0.00946921 | 0.433 | 0.087 |
| **450** | Crtc3 | 3.85E-07 | 0.711429817 | 0.009569777 | 0.635 | 0.212 |
| **451** | Esco1 | 3.85E-07 | 0.642765642 | 0.009577343 | 0.635 | 0.202 |
| **452** | Gabpb2 | 3.87E-07 | 0.378265318 | 0.009620745 | 0.538 | 0.135 |
| **453** | Fam162a | 3.87E-07 | 1.35931941 | 0.009638068 | 0.394 | 0.077 |
| **454** | Sem1 | 3.94E-07 | 0.798699502 | 0.009804912 | 0.702 | 0.24 |
| **455** | Ubl5 | 4.09E-07 | 1.847327467 | 0.010172365 | 0.337 | 0.048 |
| **456** | Clip2 | 4.11E-07 | 1.026066946 | 0.010217094 | 0.385 | 0.067 |
| **457** | Rpl18 | 4.13E-07 | 0.625885778 | 0.010284881 | 0.692 | 0.221 |
| **458** | Naca | 4.24E-07 | 0.55575184 | 0.010545273 | 0.615 | 0.183 |
| **459** | Ubc | 4.25E-07 | 0.821163261 | 0.010570791 | 0.731 | 0.279 |
| **460** | Samd4 | 4.37E-07 | 0.792454416 | 0.010880718 | 0.798 | 0.337 |
| **461** | Bicc1 | 4.40E-07 | 0.847067556 | 0.010944772 | 0.644 | 0.221 |
| **462** | Araf | 4.44E-07 | 0.437978992 | 0.01105202 | 0.529 | 0.135 |
| **463** | Pcbp1 | 4.47E-07 | 0.962186381 | 0.011118052 | 0.452 | 0.106 |
| **464** | Tmem50a | 4.49E-07 | 0.574104353 | 0.011174029 | 0.567 | 0.154 |
| **465** | Rps19 | 4.52E-07 | 1.074600988 | 0.01123735 | 0.692 | 0.279 |
| **466** | Kansl2 | 4.53E-07 | 0.532568297 | 0.011267976 | 0.413 | 0.087 |
| **467** | Ubr2 | 4.55E-07 | 0.44710967 | 0.011325504 | 0.827 | 0.288 |
| **468** | Syde1 | 4.56E-07 | 0.721520424 | 0.011348797 | 0.51 | 0.135 |
| **469** | Oaz2 | 4.61E-07 | 0.650062209 | 0.011474337 | 0.577 | 0.173 |
| **470** | Igfbp5 | 4.62E-07 | 0.983706326 | 0.011491767 | 0.548 | 0.173 |
| **471** | Zfp236 | 4.63E-07 | 0.561312101 | 0.011511871 | 0.529 | 0.144 |
| **472** | Ube3c | 4.71E-07 | 0.534388419 | 0.011723485 | 0.779 | 0.279 |
| **473** | 1700025G04R | 4.72E-07 | 0.187569079 | 0.011747042 | 0.788 | 0.279 |
| **474** | Clk2 | 4.78E-07 | 1.345384954 | 0.01188104 | 0.442 | 0.096 |
| **475** | Nhsl2 | 4.79E-07 | 0.466843163 | 0.011910698 | 0.788 | 0.327 |
| **476** | Tmbim1 | 4.81E-07 | 0.658336147 | 0.011964562 | 0.519 | 0.135 |
| **477** | Trim35 | 4.82E-07 | 0.291866643 | 0.011997771 | 0.452 | 0.106 |
| **478** | Foxo3 | 4.83E-07 | 0.442951338 | 0.012016688 | 0.769 | 0.269 |
| **479** | Heyl | 4.83E-07 | 4.337505276 | 0.012021788 | 0.24 | 0.01 |
| **480** | Ndufb1-ps | 4.88E-07 | 1.79645257 | 0.012141672 | 0.375 | 0.067 |
| **481** | Zfp618 | 4.90E-07 | 0.575490883 | 0.012181597 | 0.452 | 0.106 |
| **482** | Pcnx | 4.90E-07 | 0.418020472 | 0.012186032 | 0.76 | 0.269 |
| **483** | Enpp2 | 4.91E-07 | 0.683957105 | 0.012221699 | 0.519 | 0.144 |
| **484** | Mrc2 | 5.02E-07 | 2.331246532 | 0.012486837 | 0.346 | 0.058 |
| **485** | Rps23 | 5.05E-07 | 0.575334229 | 0.012576131 | 0.808 | 0.337 |
| **486** | Ccdc85b | 5.06E-07 | 0.496009095 | 0.01258632 | 0.654 | 0.212 |
| **487** | Sfmbt2 | 5.08E-07 | 2.426123078 | 0.012649691 | 0.288 | 0.029 |
| **488** | Ndufa4l2 | 5.13E-07 | 3.881888772 | 0.012760027 | 0.26 | 0.019 |
| **489** | Arl13b | 5.16E-07 | 0.493507923 | 0.01283089 | 0.442 | 0.096 |

| | | | | | | |
|---|---|---|---|---|---|---|
| **490** | Rnpc3 | 5.29E-07 | 0.392178214 | 0.013170032 | 0.5 | 0.125 |
| **491** | Ago2 | 5.30E-07 | 0.668267233 | 0.013196791 | 0.567 | 0.163 |
| **492** | Chst1 | 5.34E-07 | 0.707800803 | 0.013293197 | 0.462 | 0.115 |
| **493** | Irak3 | 5.38E-07 | 0.85204354 | 0.013394692 | 0.548 | 0.163 |
| **494** | Lrp1 | 5.42E-07 | 0.722181148 | 0.013488843 | 0.731 | 0.279 |
| **495** | Ston1 | 5.46E-07 | 0.707706904 | 0.013583729 | 0.538 | 0.154 |
| **496** | Mtmr2 | 5.53E-07 | 0.402297997 | 0.013766121 | 0.644 | 0.192 |
| **497** | Etv1 | 5.59E-07 | 4.036072599 | 0.013913047 | 0.24 | 0.01 |
| **498** | Per1 | 5.65E-07 | 1.04264988 | 0.014054995 | 0.413 | 0.087 |
| **499** | Camkmt | 5.68E-07 | 0.427338149 | 0.014128756 | 0.721 | 0.269 |
| **500** | Ahcyl2 | 5.72E-07 | 1.076691595 | 0.014222617 | 0.654 | 0.24 |
| **501** | Tmem30a | 5.76E-07 | 0.589816252 | 0.014320051 | 0.76 | 0.288 |
| **502** | Csnk1e | 5.78E-07 | 1.182005872 | 0.014372578 | 0.49 | 0.135 |
| **503** | Kat14 | 5.80E-07 | 1.213192861 | 0.014428008 | 0.337 | 0.048 |
| **504** | Ptma | 5.85E-07 | 0.573691936 | 0.014555455 | 0.894 | 0.413 |
| **505** | Mapkbp1 | 5.95E-07 | 1.2553841 | 0.014816185 | 0.385 | 0.077 |
| **506** | Pwwp2a | 5.97E-07 | 0.145682688 | 0.014852576 | 0.673 | 0.202 |
| **507** | B130024G19R | 6.01E-07 | 0.321992085 | 0.014955022 | 0.442 | 0.096 |
| **508** | Wdr82 | 6.06E-07 | 0.633154847 | 0.015070175 | 0.481 | 0.115 |
| **509** | Cab39l | 6.06E-07 | 0.643552555 | 0.015080109 | 0.635 | 0.212 |
| **510** | Afap1 | 6.10E-07 | 0.470741648 | 0.015184973 | 0.683 | 0.212 |
| **511** | Ndufb9 | 6.15E-07 | 1.511995997 | 0.015292324 | 0.375 | 0.067 |
| **512** | Washc4 | 6.27E-07 | 0.453305615 | 0.015606966 | 0.692 | 0.24 |
| **513** | Rassf8 | 6.41E-07 | 0.76810499 | 0.015937372 | 0.615 | 0.202 |
| **514** | Arfgef2 | 6.44E-07 | 0.640891157 | 0.016018409 | 0.567 | 0.163 |
| **515** | Fam118b | 6.46E-07 | 0.9749868 | 0.016063347 | 0.375 | 0.067 |
| **516** | Nono | 6.49E-07 | 0.580633041 | 0.016141608 | 0.5 | 0.125 |
| **517** | Ecm2 | 6.53E-07 | 0.783511297 | 0.01625929 | 0.519 | 0.144 |
| **518** | Sufu | 6.67E-07 | 1.709457346 | 0.016587651 | 0.385 | 0.077 |
| **519** | Terf1 | 6.69E-07 | 0.246912893 | 0.016641031 | 0.462 | 0.106 |
| **520** | Vps13a | 6.72E-07 | 0.171812729 | 0.016729015 | 0.798 | 0.279 |
| **521** | Lars | 6.79E-07 | 0.435429039 | 0.016897016 | 0.529 | 0.144 |
| **522** | Ntrk2 | 6.86E-07 | 1.404823278 | 0.017077621 | 0.471 | 0.125 |
| **523** | Foxn2 | 6.88E-07 | 0.829534367 | 0.017128001 | 0.462 | 0.115 |
| **524** | Taok3 | 7.02E-07 | 0.607205146 | 0.017461056 | 0.615 | 0.192 |
| **525** | Emg1 | 7.04E-07 | 3.271550216 | 0.017518556 | 0.26 | 0.019 |
| **526** | Npc2 | 7.13E-07 | 0.692733476 | 0.017731955 | 0.615 | 0.202 |
| **527** | Cox4i2 | 7.18E-07 | 3.051577023 | 0.017866891 | 0.26 | 0.019 |
| **528** | Jcad | 7.18E-07 | 0.098752147 | 0.017875633 | 0.817 | 0.298 |
| **529** | Dtx4 | 7.32E-07 | 3.263474937 | 0.018221893 | 0.26 | 0.019 |
| **530** | Cyld | 7.39E-07 | 0.562311575 | 0.018381874 | 0.606 | 0.183 |

| | | | | | | |
|---|---|---|---|---|---|---|
| **531** | Sf1 | 7.49E-07 | 0.517751036 | 0.018645266 | 0.673 | 0.212 |
| **532** | Akap12 | 7.52E-07 | 0.649766168 | 0.018703502 | 0.769 | 0.317 |
| **533** | Ralgds | 7.68E-07 | 1.515568597 | 0.019096186 | 0.346 | 0.058 |
| **534** | AI480526 | 7.73E-07 | 1.360142999 | 0.019242137 | 0.327 | 0.048 |
| **535** | Sh3rf1 | 7.84E-07 | 0.532510462 | 0.019509083 | 0.933 | 0.413 |
| **536** | Cdc40 | 7.84E-07 | 1.298396147 | 0.019515979 | 0.413 | 0.087 |
| **537** | Dhx29 | 7.99E-07 | 4.726045547 | 0.019871317 | 0.212 | 0 |
| **538** | Egfr | 8.15E-07 | 0.749770875 | 0.020288813 | 0.663 | 0.221 |
| **539** | Uqcr11 | 8.22E-07 | 0.949992881 | 0.020443135 | 0.51 | 0.144 |
| **540** | Mapre3 | 8.23E-07 | 2.247993419 | 0.020473646 | 0.308 | 0.038 |
| **541** | Tsc22d1 | 8.33E-07 | 0.774693325 | 0.020714126 | 0.875 | 0.413 |
| **542** | Xrcc4 | 8.34E-07 | 0.18693783 | 0.020750671 | 0.654 | 0.212 |
| **543** | Gosr1 | 8.51E-07 | 0.719728941 | 0.021173843 | 0.394 | 0.077 |
| **544** | Laptm4b | 8.51E-07 | 0.737464699 | 0.021173843 | 0.394 | 0.077 |
| **545** | Btbd10 | 8.55E-07 | 0.465858914 | 0.021272522 | 0.471 | 0.115 |
| **546** | Tmem263 | 8.58E-07 | 1.296690842 | 0.021349875 | 0.288 | 0.029 |
| **547** | Ccl11 | 8.60E-07 | 2.742971921 | 0.021407725 | 0.279 | 0.029 |
| **548** | Eif3h | 8.68E-07 | 0.727602161 | 0.021593978 | 0.587 | 0.192 |
| **549** | Sycp2 | 8.74E-07 | 2.733310155 | 0.021737903 | 0.26 | 0.019 |
| **550** | Mlec | 8.76E-07 | 0.996243145 | 0.021785803 | 0.356 | 0.058 |
| **551** | Stx5a | 8.80E-07 | 0.720726107 | 0.021904286 | 0.471 | 0.115 |
| **552** | Atp2a2 | 8.88E-07 | 0.549531126 | 0.022083242 | 0.808 | 0.317 |
| **553** | Rps3a1 | 8.93E-07 | 0.510477431 | 0.022221626 | 0.817 | 0.327 |
| **554** | Ulk2 | 8.94E-07 | 0.569743345 | 0.022249793 | 0.721 | 0.26 |
| **555** | Ssr1 | 9.11E-07 | 0.789010542 | 0.022673569 | 0.538 | 0.163 |
| **556** | Mapk1 | 9.13E-07 | 0.609937356 | 0.022716219 | 0.846 | 0.346 |
| **557** | Eps15l1 | 9.22E-07 | 0.610152925 | 0.022943244 | 0.702 | 0.231 |
| **558** | Rpl23 | 9.23E-07 | 0.554962122 | 0.022973702 | 0.769 | 0.308 |
| **559** | Lamc3 | 9.31E-07 | 2.237006219 | 0.023165584 | 0.365 | 0.077 |
| **560** | Il15ra | 9.35E-07 | 1.052279641 | 0.023255834 | 0.529 | 0.163 |
| **561** | Tmem68 | 9.51E-07 | 0.486986236 | 0.023650483 | 0.394 | 0.077 |
| **562** | Unc45b | 9.56E-07 | 0.479378413 | 0.023797296 | 0.5 | 0.125 |
| **563** | Nfic | 9.60E-07 | 0.828155981 | 0.023874524 | 0.51 | 0.144 |
| **564** | Cct2 | 9.60E-07 | 0.765646516 | 0.023881625 | 0.404 | 0.087 |
| **565** | Sec22a | 9.70E-07 | 1.201408602 | 0.024136408 | 0.413 | 0.096 |
| **566** | Tra2b | 9.71E-07 | 0.527091357 | 0.02417115 | 0.75 | 0.288 |
| **567** | Zfp91 | 9.75E-07 | 0.557162755 | 0.024253957 | 0.654 | 0.212 |
| **568** | Fth1 | 9.80E-07 | 0.62885089 | 0.024383242 | 0.74 | 0.288 |
| **569** | Ppp4r2 | 9.98E-07 | 0.219920242 | 0.024822511 | 0.731 | 0.24 |
| **570** | Morf4l2 | 1.01E-06 | 0.632259042 | 0.025054502 | 0.49 | 0.125 |
| **571** | Cdk11b | 1.01E-06 | 0.436438788 | 0.02513512 | 0.558 | 0.154 |

| | | | | | | |
|---|---|---|---|---|---|---|
| **572** | S1pr3 | 1.02E-06 | 3.863539026 | 0.025374051 | 0.25 | 0.019 |
| **573** | Abr | 1.02E-06 | -0.18034818 | 0.025432581 | 0.519 | 0.135 |
| **574** | Mgat4b | 1.03E-06 | 1.167690737 | 0.025558733 | 0.365 | 0.067 |
| **575** | Lgals8 | 1.03E-06 | 0.673543973 | 0.025652164 | 0.423 | 0.096 |
| **576** | Bcl7b | 1.04E-06 | 0.980893132 | 0.025806728 | 0.279 | 0.029 |
| **577** | Abcd1 | 1.04E-06 | 2.294178334 | 0.025902649 | 0.26 | 0.019 |
| **578** | Fbxo3 | 1.05E-06 | 0.67349035 | 0.026091593 | 0.471 | 0.115 |
| **579** | Pla1a | 1.06E-06 | 1.092137276 | 0.026258353 | 0.346 | 0.058 |
| **580** | Rufy3 | 1.07E-06 | 0.753623591 | 0.026738154 | 0.538 | 0.163 |
| **581** | Mast4 | 1.08E-06 | 0.577662033 | 0.026805428 | 1 | 0.683 |
| **582** | Carhsp1 | 1.10E-06 | 3.633406736 | 0.027359669 | 0.231 | 0.01 |
| **583** | Dcp1a | 1.10E-06 | 0.508856677 | 0.027427084 | 0.462 | 0.115 |
| **584** | Ccar1 | 1.11E-06 | 0.320248008 | 0.027570662 | 0.625 | 0.202 |
| **585** | Rpl24 | 1.12E-06 | 0.822457996 | 0.027845304 | 0.558 | 0.173 |
| **586** | Zfp36l2 | 1.13E-06 | 0.167062139 | 0.0281233 | 0.577 | 0.183 |
| **587** | Stim1 | 1.13E-06 | 0.382968951 | 0.028211682 | 0.846 | 0.337 |
| **588** | Hs2st1 | 1.15E-06 | 0.080106123 | 0.028554471 | 0.702 | 0.25 |
| **589** | Pgrmc2 | 1.15E-06 | 0.554895795 | 0.028646676 | 0.51 | 0.144 |
| **590** | Setd1a | 1.18E-06 | 1.692891479 | 0.02938979 | 0.317 | 0.048 |
| **591** | Ppt1 | 1.19E-06 | 1.060120467 | 0.029499561 | 0.346 | 0.058 |
| **592** | Dhx40 | 1.20E-06 | 0.548149822 | 0.029939911 | 0.51 | 0.135 |
| **593** | Ost4 | 1.22E-06 | 1.04039438 | 0.030376526 | 0.481 | 0.135 |
| **594** | Rbm27 | 1.23E-06 | 0.738885209 | 0.0305647 | 0.577 | 0.173 |
| **595** | Arhgap35 | 1.23E-06 | 0.387338365 | 0.03062394 | 0.798 | 0.288 |
| **596** | Kpnb1 | 1.24E-06 | 0.319812534 | 0.030954152 | 0.558 | 0.163 |
| **597** | Tor1aip2 | 1.25E-06 | 0.529565763 | 0.031042877 | 0.548 | 0.163 |
| **598** | Hoxb3 | 1.26E-06 | 0.354360723 | 0.031366539 | 0.394 | 0.077 |
| **599** | Hdgf | 1.27E-06 | 0.988105214 | 0.031540127 | 0.365 | 0.067 |
| **600** | Ankrd40 | 1.27E-06 | 0.625971763 | 0.03154338 | 0.433 | 0.096 |
| **601** | Plgrkt | 1.27E-06 | 0.557295304 | 0.031583743 | 0.51 | 0.144 |
| **602** | Plekhg2 | 1.29E-06 | 1.251356915 | 0.032027673 | 0.404 | 0.087 |
| **603** | Chmp2a | 1.29E-06 | 0.787893814 | 0.032052236 | 0.365 | 0.067 |
| **604** | Pitpnm2 | 1.30E-06 | 0.062822957 | 0.032302495 | 0.692 | 0.24 |
| **605** | Epb41l1 | 1.31E-06 | 1.066424941 | 0.032664233 | 0.423 | 0.096 |
| **606** | Crispld2 | 1.32E-06 | 3.051192489 | 0.032928298 | 0.231 | 0.01 |
| **607** | Vkorc1 | 1.32E-06 | 3.386425005 | 0.032928298 | 0.231 | 0.01 |
| **608** | Crlf3 | 1.34E-06 | 0.778075966 | 0.033270974 | 0.471 | 0.125 |
| **609** | Serpinh1 | 1.36E-06 | 0.804110912 | 0.0337623 | 0.654 | 0.24 |
| **610** | Syf2 | 1.36E-06 | 1.940119914 | 0.03389367 | 0.356 | 0.067 |
| **611** | Ubap2l | 1.36E-06 | 0.541212501 | 0.033948755 | 0.712 | 0.26 |
| **612** | Sparc | 1.38E-06 | 0.714657812 | 0.034249522 | 0.99 | 0.606 |

| | | | | | | |
|---|---|---|---|---|---|---|
| **613** | Ubxn8 | 1.38E-06 | 2.790933961 | 0.034306164 | 0.231 | 0.01 |
| **614** | Thap3 | 1.38E-06 | 0.957828119 | 0.034402591 | 0.49 | 0.135 |
| **615** | Sptlc1 | 1.38E-06 | 0.904222897 | 0.034444372 | 0.385 | 0.077 |
| **616** | Rpl27a | 1.39E-06 | 0.827980812 | 0.034556007 | 0.712 | 0.279 |
| **617** | Eif4b | 1.39E-06 | 0.968614034 | 0.034697876 | 0.5 | 0.135 |
| **618** | Timm17a | 1.42E-06 | 1.277467159 | 0.035383436 | 0.346 | 0.058 |
| **619** | Mpc2 | 1.42E-06 | 1.315736124 | 0.035416615 | 0.375 | 0.077 |
| **620** | Foxk2 | 1.43E-06 | 0.481823328 | 0.035501233 | 0.433 | 0.106 |
| **621** | Atp5a1 | 1.44E-06 | 0.63371308 | 0.035938709 | 0.567 | 0.173 |
| **622** | Rbpj | 1.45E-06 | 0.376857278 | 0.036155437 | 0.808 | 0.317 |
| **623** | Btf3 | 1.49E-06 | 0.563760521 | 0.037190139 | 0.462 | 0.115 |
| **624** | Pkig | 1.50E-06 | 0.656062828 | 0.037220483 | 0.654 | 0.221 |
| **625** | Lama2 | 1.50E-06 | 1.075471532 | 0.037257898 | 0.538 | 0.173 |
| **626** | Tet2 | 1.50E-06 | 0.674733132 | 0.037352601 | 0.635 | 0.221 |
| **627** | Cetn2 | 1.51E-06 | 4.637911219 | 0.03752639 | 0.202 | 0 |
| **628** | Agps | 1.52E-06 | 0.473005204 | 0.037702438 | 0.731 | 0.279 |
| **629** | Rpl10a | 1.53E-06 | 1.032607741 | 0.037997975 | 0.49 | 0.144 |
| **630** | Syn3 | 1.53E-06 | 1.825907294 | 0.038048299 | 0.279 | 0.029 |
| **631** | Cyth3 | 1.54E-06 | 0.478813771 | 0.038275012 | 0.654 | 0.231 |
| **632** | Arf4 | 1.56E-06 | 0.549450861 | 0.038745283 | 0.615 | 0.221 |
| **633** | Ndufc2 | 1.56E-06 | 0.200498962 | 0.038745283 | 0.635 | 0.202 |
| **634** | Tmem87b | 1.59E-06 | 0.360905918 | 0.03956539 | 0.51 | 0.135 |
| **635** | 4931406C07Ri | 1.59E-06 | 1.049834021 | 0.039677307 | 0.404 | 0.087 |
| **636** | Fgfr1op2 | 1.60E-06 | 0.539438902 | 0.039764949 | 0.644 | 0.221 |
| **637** | Atad1 | 1.61E-06 | 1.059889507 | 0.040027482 | 0.423 | 0.106 |
| **638** | Zfp36l1 | 1.61E-06 | 0.758400981 | 0.040057363 | 0.875 | 0.452 |
| **639** | Rpl19 | 1.61E-06 | 0.376685036 | 0.040090121 | 0.721 | 0.288 |
| **640** | Tomm7 | 1.62E-06 | 1.12546734 | 0.040293006 | 0.51 | 0.154 |
| **641** | Top1 | 1.63E-06 | 0.492670407 | 0.040466775 | 0.817 | 0.327 |
| **642** | Zfp386 | 1.63E-06 | 0.949972564 | 0.040501198 | 0.337 | 0.058 |
| **643** | Hprt | 1.64E-06 | -0.11805457 | 0.040682378 | 0.538 | 0.144 |
| **644** | Sfxn1 | 1.64E-06 | 1.282472103 | 0.040839809 | 0.385 | 0.087 |
| **645** | Adam23 | 1.64E-06 | 0.88340822 | 0.040917898 | 0.442 | 0.115 |
| **646** | Ppp2r1a | 1.65E-06 | 0.510048432 | 0.040943832 | 0.481 | 0.125 |
| **647** | Wtap | 1.66E-06 | 0.49242258 | 0.041334869 | 0.587 | 0.183 |
| **648** | Oxct1 | 1.66E-06 | 0.674343618 | 0.041377997 | 0.452 | 0.115 |
| **649** | Map2 | 1.67E-06 | 3.573459679 | 0.041659265 | 0.26 | 0.029 |
| **650** | Srsf3 | 1.69E-06 | 0.387785133 | 0.04193685 | 0.75 | 0.279 |
| **651** | Hspa9 | 1.71E-06 | 0.861502701 | 0.042633823 | 0.481 | 0.135 |
| **652** | Eif4h | 1.72E-06 | 0.614729158 | 0.042725676 | 0.548 | 0.173 |
| **653** | Lgals1 | 1.73E-06 | 1.303803793 | 0.042934595 | 0.404 | 0.096 |

| | | | | | | |
|---|---|---|---|---|---|---|
| **654** | Rcan2 | 1.74E-06 | 1.315560189 | 0.043268879 | 0.337 | 0.058 |
| **655** | 4930502A04Ri | 1.76E-06 | 2.774659885 | 0.043826341 | 0.25 | 0.019 |
| **656** | Uba5 | 1.78E-06 | 1.264199533 | 0.044201408 | 0.394 | 0.087 |
| **657** | Cox7c | 1.78E-06 | 0.55852134 | 0.044275774 | 0.663 | 0.24 |
| **658** | Atf1 | 1.80E-06 | 0.807816209 | 0.044705606 | 0.423 | 0.096 |
| **659** | Cox16 | 1.80E-06 | 0.222384033 | 0.044737518 | 0.894 | 0.413 |
| **660** | Cd81 | 1.81E-06 | 0.484276429 | 0.044961851 | 0.817 | 0.337 |
| **661** | Tagln | 1.81E-06 | 2.151631683 | 0.045062112 | 0.269 | 0.029 |
| **662** | Gnb4 | 1.84E-06 | 0.475285623 | 0.045708453 | 0.663 | 0.231 |
| **663** | Heca | 1.85E-06 | 0.52395452 | 0.046058094 | 0.615 | 0.202 |
| **664** | Golph3 | 1.86E-06 | 0.892012481 | 0.046161267 | 0.49 | 0.144 |
| **665** | Dmtf1 | 1.86E-06 | 0.467517571 | 0.046202983 | 0.519 | 0.144 |
| **666** | Ucp2 | 1.86E-06 | 0.919091532 | 0.046269683 | 0.433 | 0.106 |
| **667** | Hsp90aa1 | 1.87E-06 | 1.206884012 | 0.046457479 | 0.452 | 0.125 |
| **668** | Ube2q2 | 1.87E-06 | 0.786222681 | 0.04649991 | 0.51 | 0.144 |
| **669** | Zbtb11 | 1.88E-06 | 1.002668054 | 0.046891064 | 0.394 | 0.096 |
| **670** | Ptpn11 | 1.89E-06 | 0.279475091 | 0.047112301 | 0.548 | 0.163 |
| **671** | Ksr1 | 1.90E-06 | 0.476965819 | 0.047340818 | 0.644 | 0.231 |
| **672** | Atp5e | 1.91E-06 | 0.651913017 | 0.047544928 | 0.558 | 0.173 |
| **673** | Ap2s1 | 1.91E-06 | 2.453734882 | 0.047627332 | 0.269 | 0.029 |
| **674** | Flot2 | 1.92E-06 | 1.073854587 | 0.047712992 | 0.375 | 0.077 |
| **675** | Afap1l2 | 1.92E-06 | 0.01586905 | 0.047867592 | 0.529 | 0.154 |
| **676** | Mosmo | 1.93E-06 | 0.198142036 | 0.047940909 | 0.654 | 0.221 |
| **677** | Echdc1 | 1.93E-06 | 0.903198335 | 0.047969659 | 0.298 | 0.038 |
| **678** | Tram2 | 1.95E-06 | 1.38167536 | 0.048555312 | 0.337 | 0.058 |
| **679** | Arhgdia | 1.96E-06 | 0.717068313 | 0.048787445 | 0.49 | 0.144 |
| **680** | Btbd3 | 1.98E-06 | 0.303241073 | 0.049189037 | 0.712 | 0.26 |
| **681** | Dnaja2 | 1.99E-06 | 1.120580098 | 0.049392415 | 0.442 | 0.115 |
| **682** | Ppp6r2 | 1.99E-06 | 0.77566453 | 0.049460902 | 0.548 | 0.173 |
| **683** | Ptpn2 | 1.99E-06 | 0.826568808 | 0.049474045 | 0.442 | 0.106 |
| **684** | Morc2a | 1.99E-06 | 0.846987186 | 0.049526932 | 0.471 | 0.125 |
| **685** | Nhs | 2.00E-06 | 0.838813401 | 0.049795034 | 0.51 | 0.144 |

**Extended Table 5**

DEG from SnC (SenMayo strategy) predicted to encode secreted proteins

| Protein Name | Gene Name | Localizations | Signals | Cytoplasm | Nucleus | Extracellular |
|---|---|---|---|---|---|---|
| Disintegrin and metalloproteinase domain-containing protein 23 | Adam23 | Extracellular | Signal peptide | 0.2228 | 0.1962 | 0.872900009 |
| Ig-like domain-containing protein | Adgra3 | Extracellular | Signal peptide | 0.1882 | 0.1737 | 0.782299995 |
| Adiponectin | Adipoq | Extracellular | Signal peptide | 0.232 | 0.1185 | 0.899100006 |
| AF4/FMR2 family, member 3 | Aff3 | Nucleus\|Extracellular | | 0.4613 | 0.5905 | 0.671599984 |
| fructose-bisphosphate aldolase | Aldoa | Extracellular | Signal peptide | 0.3498 | 0.2837 | 0.744000018 |
| Angiopoietin 2 | Angpt2 | Extracellular | Signal peptide | 0.1681 | 0.0628 | 0.881099999 |
| Angiopoietin-like 2 | Angptl2 | Extracellular | Signal peptide | 0.1372 | 0.0949 | 0.862600029 |
| Angiopoietin-like 4 | Angptl4 | Extracellular | Signal peptide | 0.1324 | 0.1287 | 0.831799984 |
| Anthrax toxin receptor 1 | Antxr1 | Extracellular | | 0.4415 | 0.292 | 0.66109997 |
| Annexin | Anxa1 | Cytoplasm\|Nucleus\|Extracellul | | 0.4776 | 0.5319 | 0.836300015 |
| Annexin A4 | Anxa4 | Cytoplasm\|Extracellular\|Lysos | | 0.7347 | 0.4551 | 0.654900014 |
| Annexin A5 | Anxa5 | Cytoplasm\|Extracellular\|Lysos | | 0.7837 | 0.4363 | 0.669399977 |
| Apolipoprotein E | Apoe | Extracellular | Signal peptide | 0.1566 | 0.0715 | 0.878199995 |
| Aspn protein | Aspn | Extracellular | Signal peptide | 0.1825 | 0.0713 | 0.901899993 |
| LRRNT domain-containing protein | Aspn | Extracellular | Signal peptide | 0.1046 | 0.0522 | 0.92019999 |
| Beta-2-microglobulin | B2m | Extracellular | Signal peptide | 0.1346 | 0.0455 | 0.771399975 |
| Beta-1,3-glucosyltransferase | B3glct | Extracellular\|E | Signal peptide | 0.179 | 0.0775 | 0.665799975 |
| Basal cell adhesion molecule | Bcam | Extracellular | Signal peptide | 0.1469 | 0.1064 | 0.808799982 |
| Biglycan | Bgn | Extracellular | Signal peptide | 0.1482 | 0.1458 | 0.878400028 |
| Biglycan | Bgn | Extracellular | Signal peptide | 0.1891 | 0.0925 | 0.862600029 |
| 4632419I22Rik protein | Brip1os | Extracellular | Signal peptide | 0.1933 | 0.1183 | 0.838100016 |
| Basigin | Bsg | Extracellular | Signal peptide | 0.299 | 0.2612 | 0.8046 |
| Adiponectin p | C1qtnf1 | Extracellular | Signal peptide | 0.2566 | 0.1366 | 0.886200011 |
| Complement C1q and tumor necrosis factor-related protein 9 | C1qtnf9 | Extracellular | Signal peptide | 0.1764 | 0.1079 | 0.907700002 |
| Complement C1r-A subcomponent | C1ra | Extracellular | Signal peptide | 0.1842 | 0.1227 | 0.879599988 |
| Complement C1s subcomponent | C1s1 | Extracellular | Signal peptide | 0.1625 | 0.0814 | 0.754800022 |
| Complement C3 | C3 | Extracellular | Signal peptide | 0.3199 | 0.1107 | 0.900699973 |

| | | | | | | |
|---|---|---|---|---|---|---|
| Complement C4-B | C4b | Extracellular | Signal peptide | 0.2813 | 0.1946 | 0.929400027 |
| Calcium channel, voltage-dependent, alpha2/delta subunit 1 | Cacna2d1 | Extracellular | Signal peptide | 0.2416 | 0.1121 | 0.696500003 |
| Calmodulin 2 | Calm2 | Extracellular | Signal peptide | 0.4661 | 0.2898 | 0.811699986 |
| Calreticulin | Calr | Extracellular\|Er | Signal peptide | 0.214 | 0.0771 | 0.693300009 |
| Calumenin | Calu | Extracellular | Signal peptide | 0.1517 | 0.0529 | 0.74000001 |
| Eotaxin | Ccl11 | Extracellular | Signal peptide | 0.1038 | 0.0946 | 0.935500026 |
| CD44 antigen | Cd44 | Extracellular | Signal peptide | 0.1219 | 0.1284 | 0.896799982 |
| CD93 antigen | Cd93 | Extracellular | Signal peptide | 0.4033 | 0.1758 | 0.959100008 |
| CEA cell adhesion molecule 1 | Ceacam1 | Extracellular | Signal peptide | 0.2918 | 0.1163 | 0.746699989 |
| C-type lectin domain family 3, member b | Clec3b | Extracellular | Signal peptide | 0.1849 | 0.0552 | 0.930299997 |
| Alpha-1 type I procollagen | Col1a1 | Extracellular | Signal peptide | 0.4143 | 0.1948 | 0.860499978 |
| Col1a2 protein | Col1a2 | Extracellular | Signal peptide | 0.3974 | 0.2616 | 0.935400009 |
| Col3a1 protein | Col3a1 | Extracellular | Signal peptide | 0.4002 | 0.3474 | 0.943099976 |
| Collagen alpha-1 | Col5a1 | Extracellular | Signal peptide | 0.2762 | 0.1446 | 0.892499983 |
| Adipocyte-specific protein 6 | Col5a3 | Extracellular | Signal peptide | 0.2859 | 0.1799 | 0.892799973 |
| Col6a1 protein | Col6a1 | Extracellular | Signal peptide | 0.3478 | 0.2244 | 0.970399976 |
| Collagen alpha-2 | Col6a2 | Extracellular | Signal peptide | 0.1707 | 0.1226 | 0.917699993 |
| Col6a3 protein | Col6a3 | Extracellular | Signal peptide | 0.3275 | 0.1932 | 0.943499982 |
| Cytochrome c oxidase, subunit 6B1 | Cox6b1 | Extracellular | | 0.3002 | 0.0735 | 0.393700004 |
| Ceruloplasmin | Cp | Extracellular | Signal peptide | 0.1841 | 0.115 | 0.657899976 |
| Cellular repressor of E1A-stimulated genes 1 | Creg1 | Extracellular\|Er | Signal peptide | 0.1439 | 0.0472 | 0.853500009 |
| Chondroitin sulfate proteoglycan 4 | Cspg4 | Extracellular | Signal peptide | 0.3729 | 0.2632 | 0.848999977 |
| Cystatin C | Cst3 | Extracellular | Signal peptide | 0.1363 | 0.0815 | 0.845200002 |
| Protein CTLA-2-alpha | Ctla2a | Extracellular\|Ly | Signal peptide | 0.1549 | 0.0773 | 0.691100001 |
| Cathepsin B | Ctsb | Extracellular\|Ly | Signal peptide | 0.4092 | 0.1553 | 0.850399971 |
| Cathepsin L | Ctsl | Extracellular | Signal peptide | 0.4234 | 0.1265 | 0.705799997 |
| Stromal cell-derived factor 1 | Cxcl12 | Extracellular | Signal peptide | 0.1116 | 0.0948 | 0.944500029 |
| DAN domain family member 5 | Dand5 | Extracellular | Signal peptide | 0.1474 | 0.1175 | 0.922100008 |
| Discoidin domain receptor family, member 2 | Ddr2 | Extracellular | Signal peptide | 0.19 | 0.1353 | 0.794200003 |
| Dermatopontin | Dpt | Extracellular | Signal peptide | 0.1315 | 0.0613 | 0.824299991 |

| | | | | | | |
|---|---|---|---|---|---|---|
| Dermatan sulfate epimerase | Dse | Extracellular | Signal peptide | 0.2618 | 0.0749 | 0.839299977 |
| Extracellular matrix protein 2 | Ecm2 | Extracellular | Signal peptide | 0.1854 | 0.1387 | 0.854300022 |
| Eukaryotic translation elongation factor 1 beta 2 | Eef1b2 | Extracellular | Signal peptide | 0.4394 | 0.5141 | 0.534099996 |
| 4732418C07Rik protein | Efcab14 | Extracellular | Signal peptide | 0.4453 | 0.1264 | 0.83130002 |
| Pikachurin | Egflam | Extracellular | Signal peptide | 0.2831 | 0.1213 | 0.943099976 |
| Epidermal growth factor receptor | Egfr | Extracellular | Signal peptide | 0.4594 | 0.3088 | 0.867299974 |
| Initiation factor eIF-4A long form | Eif4a1 | Extracellular | Signal peptide | 0.3147 | 0.2949 | 0.623000026 |
| Eukaryotic translation initiation factor 5 | Eif5 | Extracellular | Signal peptide | 0.3057 | 0.4877 | 0.675599992 |
| Aminopeptidase | Erap1 | Cytoplasm\|Extı | Signal peptide | 0.516 | 0.2657 | 0.714699984 |
| Fibroblast activation protein | Fap | Extracellular | Signal peptide | 0.3932 | 0.2834 | 0.580299973 |
| EGF-like domain-containing protein | Fbln1 | Extracellular | | 0.3813 | 0.4965 | 0.712300003 |
| Fc fragment of IgG receptor and transporter | Fcgrt | Extracellular | Signal peptide | 0.1882 | 0.1323 | 0.644900024 |
| Growth hormone receptor | Ghr | Extracellular | Signal peptide | 0.156 | 0.0881 | 0.878600001 |
| Gastric inhibitory polypeptide receptor | Gipr | Extracellular | Signal peptide | 0.2779 | 0.2737 | 0.652999997 |
| Golgi apparatus protein 1 | Glg1 | Extracellular | Signal peptide | 0.1259 | 0.1535 | 0.878400028 |
| Uncharacterized protein | Gm10252 | Cytoplasm\|Extı | Signal peptide | 0.5496 | 0.5039 | 0.765200019 |
| Uncharacterized protein | Gm10575 | Cytoplasm\|Extı | Signal peptide | 0.6281 | 0.3581 | 0.826600015 |
| Predicted gene 21970 | Gm21970 | Extracellular\|Cell membrane | | 0.1966 | 0.1117 | 0.726599991 |
| Glucosamine | Gns | Extracellular | Signal peptide | 0.1568 | 0.2253 | 0.785600007 |
| Glypican-6 | Gpc6 | Extracellular | Signal peptide | 0.144 | 0.0975 | 0.777499974 |
| GPI-anchored HDL-binding protein 1 | Gpihbp1 | Extracellular | Signal peptide | 0.1323 | 0.1123 | 0.906400025 |
| Glutathione peroxidase | Gpx3 | Extracellular | Signal peptide | 0.2062 | 0.0776 | 0.820200026 |
| Granulin | Grn | Extracellular | Signal peptide | 0.2534 | 0.1 | 0.891099989 |
| Granzyme N | Gzmn | Extracellular | Signal peptide | 0.2504 | 0.0797 | 0.930400014 |
| MHC class I antigen H-2D | H2-D1 | Cytoplasm\|Extı | Signal peptide | 0.5788 | 0.1239 | 0.75880003 |
| Histocompatibility 2, K1, K region | H2-K1 | Extracellular | Signal peptide | 0.201 | 0.1437 | 0.703499973 |
| H2-T23 protein | H2-T23 | Extracellular | Signal peptide | 0.2869 | 0.1846 | 0.717499971 |
| Hepatocyte growth factor | Hgf | Extracellular | Signal peptide | 0.1688 | 0.0956 | 0.93900001 |
| Heat shock protein 90, alpha | Hsp90aa1 | Extracellular | Signal peptide | 0.248 | 0.5058 | 0.771300018 |
| Heat shock protein 90 alpha | Hsp90ab1 | Extracellular | Signal peptide | 0.3674 | 0.3834 | 0.4991 |

| | | | | | | |
|---|---|---|---|---|---|---|
| Heat shock protein 90, beta | Hsp90b1 | Extracellular | Signal peptide | 0.29 | 0.3716 | 0.520799994 |
| 78 kDa glucose-regulated protein | Hspa5 | Extracellular | Signal peptide | 0.166 | 0.0999 | 0.772800028 |
| Heat shock cognate 71 kDa protein | Hspa8 | Cytoplasm\|Ext | Signal peptide | 0.6409 | 0.5154 | 0.719099998 |
| Iduronate sulfatase | Ids | Extracellular | Signal peptide | 0.1431 | 0.1368 | 0.599799991 |
| Ifnar2 protein | Ifnar2 | Extracellular\|C | Signal peptide | 0.193 | 0.1072 | 0.694700003 |
| Interferon gamma receptor 1 | Ifngr1 | Extracellular | Signal peptide | 0.1368 | 0.1173 | 0.787599981 |
| Insulin-like growth factor binding protein 4 | Igfbp4 | Extracellular | Signal peptide | 0.2528 | 0.1425 | 0.799799979 |
| Igfbp7 protein | Igfbp7 | Extracellular | Signal peptide | 0.238 | 0.1076 | 0.901899993 |
| VH186.2-D-J-C mu protein | Ighm | Extracellular | Signal peptide | 0.3237 | 0.3386 | 0.564100027 |
| Interleukin 10 receptor, beta | Il10rb | Extracellular | Signal peptide | 0.1214 | 0.058 | 0.825399995 |
| Interleukin-34 | Il34 | Extracellular | Signal peptide | 0.2884 | 0.151 | 0.726300001 |
| Integrin beta 1 | Itgb1 | Extracellular | Signal peptide | 0.1162 | 0.0597 | 0.948800027 |
| Inter-alpha-trypsin inhibitor heavy chain H5 | Itih5 | Extracellular | Signal peptide | 0.2524 | 0.0968 | 0.886600018 |
| Lama2 protein | Lama2 | Extracellular | Signal peptide | 0.2818 | 0.1252 | 0.982599974 |
| Lamb1-1 protein | Lamb1 | Cytoplasm\|Ext | Signal peptide | 0.5757 | 0.1145 | 0.86650002 |
| Galectin-1 | Lgals1 | Cytoplasm\|Extracellular | | 0.7546 | 0.2794 | 0.730300009 |
| Galectin | Lgals8 | Cytoplasm\|Extracellular | | 0.6086 | 0.3365 | 0.742500007 |
| EGF-like domain-containing protein | Lrp1 | Cytoplasm\|Ext | Signal peptide | 0.5361 | 0.182 | 0.930800021 |
| Latent transforming growth factor beta binding protein 1 | Ltbp1 | Extracellular | Signal peptide | 0.1787 | 0.1839 | 0.926900029 |
| Lymphocyte antigen 6 family member A | Ly6a | Extracellular | Signal peptide | 0.0848 | 0.0907 | 0.860300004 |
| Lymphocyte antigen 6 family member C1 | Ly6c1 | Extracellular | Signal peptide | 0.1608 | 0.1082 | 0.918200016 |
| Ly6e protein | Ly6e | Extracellular | Signal peptide | 0.1829 | 0.1389 | 0.923500001 |
| lysozyme | Lyz2 | Extracellular | Signal peptide | 0.0782 | 0.0639 | 0.905099988 |
| MAM domain containing 2 | Mamdc2 | Extracellular | Signal peptide | 0.211 | 0.1068 | 0.93809998 |
| LRP chaperone MESD | Mesd | Extracellular | Signal peptide | 0.147 | 0.1291 | 0.769299984 |
| Lactadherin | Mfge8 | Extracellular | Signal peptide | 0.2162 | 0.1072 | 0.881900012 |
| Matrix metallopeptidase 14 | Mmp14 | Extracellular | Signal peptide | 0.1463 | 0.1294 | 0.686399996 |
| Multimerin 2 | Mmrn2 | Extracellular | Signal peptide | 0.4561 | 0.0935 | 0.782100022 |
| C-type mannose receptor 2 | Mrc2 | Extracellular | Signal peptide | 0.2128 | 0.1342 | 0.858699977 |
| 5-methyltetrahydrofolate-homocysteine methyltransferase | Mtr | Extracellular | Signal peptide | 0.2725 | 0.4246 | 0.656000018 |

| | | | | | | |
|---|---|---|---|---|---|---|
| Beta-nerve growth factor | Ngf | Extracellular | Signal peptide | 0.103 | 0.1454 | 0.955900013 |
| Neurogenic locus notch homolog protein 3 | Notch3 | Extracellular | Signal peptide | 0.3297 | 0.4178 | 0.459600002 |
| NPC intracellular cholesterol transporter 2 | Npc2 | Extracellular | Signal peptide | 0.1159 | 0.0374 | 0.920799971 |
| Neurotrophic tyrosine kinase, receptor, type 2 | Ntrk2 | Extracellular | | 0.2172 | 0.4714 | 0.604900002 |
| Mimecan | Ogn | Extracellular | Signal peptide | 0.1247 | 0.0744 | 0.869599998 |
| Protein OS-9 | Os9 | Extracellular | Signal peptide | 0.2504 | 0.132 | 0.692900002 |
| Pappalysin 2 | Pappa2 | Extracellular | Signal peptide | 0.3518 | 0.2209 | 0.79369998 |
| EF-hand domain-containing protein | Pdcd6 | Extracellular | Signal peptide | 0.3843 | 0.4266 | 0.735700011 |
| Protein disulfide-isomerase A6 | Pdia3 | Extracellular | Signal peptide | 0.3737 | 0.3741 | 0.795799971 |
| Protein disulfide-isomerase A6 | Pdia6 | Extracellular\|E | Signal peptide | 0.169 | 0.1022 | 0.718699992 |
| Phosphatidylinositol-4-phosphate 5-kinase, type 1 gamma | Pip5k1c | Cytoplasm\|Extracellular | | 0.587 | 0.3761 | 0.717800021 |
| Secretory phospholipase A2 receptor | Pla2r1 | Extracellular | Signal peptide | 0.2276 | 0.111 | 0.851000011 |
| Plasminogen activator, urokinase | Plau | Extracellular | Signal peptide | 0.3219 | 0.1959 | 0.744899988 |
| Lysyl hydroxylase 2 | Plod2 | Extracellular | Signal peptide | 0.3253 | 0.2111 | 0.717000008 |
| Phospholipid transfer protein | Pltp | Extracellular | Signal peptide | 0.2041 | 0.0908 | 0.850799978 |
| Plexin A1 | Plxna1 | Extracellular | Signal peptide | 0.2775 | 0.3671 | 0.715300024 |
| Periostin | Postn | Extracellular | Signal peptide | 0.1903 | 0.0592 | 0.879100025 |
| Peptidyl-prolyl cis-trans isomerase | Ppia | Extracellular | Signal peptide | 0.337 | 0.3937 | 0.672699988 |
| Peptidyl-prolyl cis-trans isomerase | Ppib | Extracellular\|E | Signal peptide | 0.1825 | 0.0837 | 0.69599998 |
| Laminin G domain-containing protein | Pros1 | Extracellular | Signal peptide | 0.3197 | 0.1075 | 0.99000001 |
| Prosaposin | Psap | Extracellular | Signal peptide | 0.1282 | 0.0517 | 0.965900004 |
| Uncharacterized protein | Pten | Extracellular | Signal peptide | 0.4157 | 0.423 | 0.514900029 |
| Protein tyrosine phosphatase domain containing 1 | Ptpdc1 | Extracellular | | 0.3456 | 0.2442 | 0.465700001 |
| Pituitary tumor-transforming 1 interacting protein | Pttg1ip | Extracellular | Signal peptide | 0.1526 | 0.105 | 0.685199976 |
| Receptor | Ramp2 | Extracellular | Signal peptide | 0.1882 | 0.1831 | 0.656000018 |
| Retinoic acid receptor responder protein 2 | Rarres2 | Extracellular | Signal peptide | 0.1112 | 0.1401 | 0.923600018 |
| Uncharacterized protein | Rbx1 | Extracellular | Signal peptide | 0.3923 | 0.5089 | 0.809499979 |
| Resistin | Retn | Extracellular | Signal peptide | 0.1803 | 0.0726 | 0.907000005 |
| Ribonuclease 4 | Rnase4 | Extracellular | Signal peptide | 0.1157 | 0.1135 | 0.897599995 |
| Ribonuclease, RNase K | Rnasek | Extracellular\|C | Signal peptide\| | 0.2822 | 0.1568 | 0.696799994 |

| | | | | | | |
|---|---|---|---|---|---|---|
| Ig-like domain-containing protein | Robo1 | Extracellular | Signal peptide | 0.22 | 0.16 | 0.554700017 |
| Ror1 protein | Ror1 | Extracellular | Signal peptide | 0.1076 | 0.1247 | 0.88499999 |
| Ribosomal protein L27A | Rpl27a | Extracellular\|Ly | Signal peptide | 0.3612 | 0.2126 | 0.649500012 |
| Uncharacterized protein | Rsrc2 | Nucleus\|Extrac | Signal peptide | 0.3914 | 0.6945 | 0.714699984 |
| Receptor-like tyrosine kinase | Ryk | Extracellular | Signal peptide | 0.1569 | 0.1734 | 0.644800007 |
| S100 calcium binding protein A1 | S100a1 | Nucleus\|Extracellular | | 0.3159 | 0.6031 | 0.671700001 |
| 45 kDa calcium-binding protein | Sdf4 | Extracellular | Signal peptide | 0.2324 | 0.1388 | 0.702600002 |
| Selenoprotein F | Selenof | Extracellular | Signal peptide | 0.1693 | 0.1118 | 0.773800015 |
| Selenoprotein S | Selenos | Cytoplasm\|Ext | Signal peptide | 0.582 | 0.312 | 0.722299993 |
| Sema domain-containing protein | Sema5a | Extracellular | Signal peptide | 0.141 | 0.059 | 0.886399984 |
| Glia-derived nexin | Serpine2 | Extracellular | Signal peptide | 0.1694 | 0.0483 | 0.845099986 |
| Pigment epithelium-derived factor | Serpinf1 | Extracellular | Signal peptide | 0.1864 | 0.0612 | 0.932099998 |
| Plasma protease C1 inhibitor | Serping1 | Extracellular | Signal peptide | 0.1847 | 0.0832 | 0.897400022 |
| Serine | Serpinh1 | Extracellular | Signal peptide | 0.1654 | 0.0388 | 0.946099997 |
| Set protein | Set | Nucleus\|Extrac | Signal peptide | 0.1815 | 0.6549 | 0.737699986 |
| Glycosyl hydrolase family 13 catalytic domain-containing protein | Slc3a2 | Cytoplasm\|Ext | Signal peptide | 0.5444 | 0.1589 | 0.818000019 |
| SPARC | Sparc | Extracellular | Signal peptide | 0.1795 | 0.0699 | 0.91109997 |
| SPARC-like 1 | Sparcl1 | Extracellular | Signal peptide | 0.1101 | 0.0772 | 0.935000002 |
| Serglycin | Srgn | Extracellular | Signal peptide | 0.0926 | 0.0647 | 0.910099983 |
| Translocon-associated protein subunit alpha | Ssr1 | Extracellular | Signal peptide | 0.3708 | 0.1268 | 0.701900005 |
| Synemin | Synm | Extracellular | Signal peptide | 0.3041 | 0.1714 | 0.814100027 |
| Teneurin transmembrane protein 4 | Tenm4 | Extracellular | Signal peptide | 0.3505 | 0.3082 | 0.572000027 |
| Serotransferrin | Tf | Extracellular | Signal peptide | 0.2249 | 0.0516 | 0.88319999 |
| Metalloproteinase inhibitor 3 | Timp3 | Extracellular | Signal peptide | 0.1205 | 0.1096 | 0.938399971 |
| TM2 domain containing 1 | Tm2d1 | Extracellular | Signal peptide | 0.1769 | 0.1398 | 0.621800005 |
| Transmembrane 9 superfamily member 2 | Tm9sf2 | Extracellular | Signal peptide | 0.1081 | 0.1198 | 0.774999976 |
| GOLD domain-containing protein | Tmed5 | Extracellular | Signal peptide | 0.1526 | 0.0881 | 0.840799987 |
| Transmembrane protein 109 | Tmem109 | Extracellular | Signal peptide | 0.1492 | 0.1035 | 0.681200027 |
| Transmembrane protein 184b | Tmem184b | Extracellular | Signal peptide | 0.469 | 0.3431 | 0.645099998 |
| Insulin-like growth factor-binding protein 3 receptor | Tmem219 | Extracellular | Signal peptide | 0.3842 | 0.3055 | 0.580299973 |

| | | | | | | |
|---|---|---|---|---|---|---|
| Tumor necrosis factor receptor superfamily, member 1a | Tnfrsf1a | Cytoplasm\|Ext | Signal peptide | 0.5528 | 0.0838 | 0.654200017 |
| Clcn1 protein | Tnxb | Extracellular | Signal peptide | 0.3667 | 0.2912 | 0.781799972 |
| Transferrin | Trf | Extracellular | Signal peptide | 0.175 | 0.0739 | 0.845399976 |
| UDP-glucuronate decarboxylase 1 | Uxs1 | Extracellular | Signal peptide\| | 0.1882 | 0.207 | 0.590200007 |

**Extended Table 6**

DEG from SnC (SEAC strategy) predicted to encode secreted proteins

| Protein Name | Gene Name | Localizations | Signals | Cytoplasm | Nucleus | Extracellular |
|---|---|---|---|---|---|---|
| Bridge-like lipid transfer protein family member 2 | Bltp2 | Cytoplasm\|Ext | Signal peptide | 0.4902 | 0.1827 | 0.64749998 |
| Disintegrin and metalloproteinase domain-containing protein 23 | Adam23 | Extracellular | Signal peptide | 0.2228 | 0.1962 | 0.87290001 |
| A disintegrin and metalloproteinase with thrombospondin motifs 12 | Adamts12 | Extracellular | Signal peptide | 0.1274 | 0.1202 | 0.94370002 |
| A disintegrin and metalloproteinase with thrombospondin motifs 15 | Adamts15 | Extracellular | Signal peptide | 0.1921 | 0.1229 | 0.95850003 |
| Adhesion G protein-coupled receptor L3 | Adgrl3 | Extracellular | Signal peptide | 0.127 | 0.0981 | 0.9501 |
| Adiponectin | Adipoq | Extracellular | Signal peptide | 0.232 | 0.1185 | 0.89910001 |
| Angiopoietin-2 | Angpt2 | Extracellular | Signal peptide | 0.1681 | 0.0628 | 0.8811 |
| Angiopoietin-related protein 2 | Angptl2 | Extracellular | Signal peptide | 0.1578 | 0.1352 | 0.90619999 |
| Angiopoietin-related protein 4 | Angptl4 | Extracellular | Signal peptide | 0.1697 | 0.1125 | 0.87819999 |
| Annexin A4 | Anxa4 | Cytoplasm\|Extracellular\|Lysos | | 0.7347 | 0.4551 | 0.65490001 |
| Annexin A5 | Anxa5 | Cytoplasm\|Extracellular\|Lysos | | 0.7837 | 0.4363 | 0.66939998 |
| Apolipoprotein E | Apoe | Extracellular | Signal peptide | 0.1882 | 0.0727 | 0.90719998 |
| Asporin | Aspn | Extracellular | Signal peptide | 0.1825 | 0.0713 | 0.90189999 |
| Beta-2-microglobulin | B2m | Extracellular | Signal peptide | 0.1266 | 0.0439 | 0.78969997 |
| Beta-1,3-glucosyltransferase | B3glct | Extracellular\|Er | Signal peptide | 0.179 | 0.0775 | 0.66579998 |
| Biglycan | Bgn | Extracellular | Signal peptide | 0.1888 | 0.0938 | 0.86119998 |
| 4632419I22Rik protein | Brip1os | Extracellular | Signal peptide | 0.1933 | 0.1183 | 0.83810002 |
| Complement C1q tumor necrosis factor-related protein 1 | C1qtnf1 | Extracellular | Signal peptide | 0.2566 | 0.1366 | 0.88620001 |
| Complement C1q and tumor necrosis factor-related protein 9 | C1qtnf9 | Extracellular | Signal peptide | 0.3635 | 0.1862 | 0.75779998 |
| Complement C1r-A subcomponent | C1ra | Extracellular | Signal peptide | 0.1842 | 0.1227 | 0.87959999 |
| Complement C1s-1 subcomponent | C1s1 | Extracellular | Signal peptide | 0.1533 | 0.0821 | 0.72070003 |
| Complement C3 | C3 | Extracellular | Signal peptide | 0.3199 | 0.1107 | 0.90069997 |
| Complement C4-B | C4b | Extracellular | Signal peptide | 0.2813 | 0.1946 | 0.92940003 |
| Calreticulin | Calr | Extracellular\|Er | Signal peptide | 0.214 | 0.0771 | 0.69330001 |
| Eotaxin | Ccl11 | Extracellular | Signal peptide | 0.1038 | 0.0946 | 0.93550003 |
| CCN family member 1 | Ccn1 | Extracellular | Signal peptide | 0.2041 | 0.0944 | 0.97289997 |
| CMRF35-like molecule 9 | Cd300lg | Extracellular | Signal peptide | 0.1806 | 0.2244 | 0.77920002 |
| Membrane cofactor protein | Cd46 | Extracellular | Signal peptide | 0.195 | 0.1854 | 0.84189999 |
| Complement factor D | Cfd | Extracellular | Signal peptide | 0.1552 | 0.0523 | 0.9346 |
| Tetranectin | Clec3b | Extracellular | Signal peptide | 0.1888 | 0.0566 | 0.92159998 |

| | | | | | | |
|---|---|---|---|---|---|---|
| Collagen alpha-1 | Col1a1 | Extracellular | Signal peptide | 0.1657 | 0.0969 | 0.92650002 |
| Collagen alpha-2 | Col1a2 | Extracellular | Signal peptide | 0.1486 | 0.1225 | 0.92799997 |
| Collagen alpha-1 | Col3a1 | Extracellular | Signal peptide | 0.2368 | 0.1559 | 0.9188 |
| Collagen alpha-1 | Col4a1 | Extracellular | Signal peptide | 0.182 | 0.1111 | 0.89819998 |
| Collagen alpha-2 | Col4a2 | Extracellular | Signal peptide | 0.2199 | 0.1306 | 0.87510002 |
| Collagen alpha-1 | Col5a1 | Extracellular | Signal peptide | 0.2762 | 0.1446 | 0.89249998 |
| Collagen type V alpha 3 chain | Col5a3 | Extracellular | Signal peptide | 0.3116 | 0.1522 | 0.88959998 |
| Collagen alpha-1 | Col6a1 | Extracellular | Signal peptide | 0.2188 | 0.1551 | 0.92540002 |
| Collagen alpha-2 | Col6a2 | Extracellular | Signal peptide | 0.1707 | 0.1226 | 0.91769999 |
| Collagen, type VI, alpha 3 | Col6a3 | Extracellular | Signal peptide | 0.148 | 0.1238 | 0.89200002 |
| Carboxypeptidase E | Cpe | Extracellular | Signal peptide | 0.1612 | 0.1451 | 0.83200002 |
| Protein CREG1 | Creg1 | Extracellular | Signal peptide | 0.1508 | 0.0533 | 0.83289999 |
| Chondroitin sulfate proteoglycan 4 | Cspg4 | Extracellular | Signal peptide | 0.3729 | 0.2632 | 0.84899998 |
| Cystatin-C | Cst3 | Extracellular | Signal peptide | 0.1423 | 0.0881 | 0.85970002 |
| Protein CTLA-2-alpha | Ctla2a | Extracellular\|Ly | Signal peptide | 0.1549 | 0.0773 | 0.6911 |
| Lysosomal protective protein | Ctsa | Extracellular\|Ly | Signal peptide | 0.1249 | 0.0744 | 0.8039 |
| Procathepsin L | Ctsl | Extracellular\|Ly | Signal peptide | 0.157 | 0.0571 | 0.77469999 |
| Cathepsin Z | Ctsz | Extracellular | Signal peptide | 0.1348 | 0.0713 | 0.7385 |
| Stromal cell-derived factor 1 | Cxcl12 | Extracellular | Signal peptide | 0.0998 | 0.085 | 0.92799997 |
| C-X-C motif chemokine 9 | Cxcl9 | Extracellular | Signal peptide | 0.1276 | 0.1569 | 0.89029998 |
| Dermatopontin | Dpt | Extracellular | Signal peptide | 0.1315 | 0.0613 | 0.82429999 |
| Extracellular matrix protein 1 | Ecm1 | Extracellular | Signal peptide | 0.2492 | 0.1252 | 0.87819999 |
| Extracellular matrix protein 1 | Ecm1 | Extracellular | Signal peptide | 0.2315 | 0.0989 | 0.89789999 |
| Extracellular matrix protein 2 | Ecm2 | Extracellular | Signal peptide | 0.1854 | 0.1387 | 0.85430002 |
| SPARC-like protein 1 | Sparcl1 | Extracellular | Signal peptide | 0.1721 | 0.0853 | 0.88819999 |
| Pikachurin | Egflam | Extracellular | Signal peptide | 0.2894 | 0.124 | 0.9386 |
| Autotaxin | Enpp2 | Extracellular | Signal peptide | 0.1594 | 0.1028 | 0.8581 |
| Follistatin-related protein 1 | Fstl1 | Extracellular | Signal peptide | 0.1458 | 0.0901 | 0.93449998 |
| Tissue alpha-L-fucosidase | Fuca1 | Extracellular\|Ly | Signal peptide | 0.2145 | 0.083 | 0.79189998 |
| Neutral alpha-glucosidase AB | Ganab | Extracellular | Signal peptide | 0.3044 | 0.1069 | 0.68800002 |
| Growth hormone receptor | Ghr | Extracellular | Signal peptide | 0.1605 | 0.1138 | 0.86229998 |
| Progranulin | Grn | Extracellular | Signal peptide | 0.2341 | 0.083 | 0.90259999 |
| MHC class I-like antigen recognition-like domain-containing protein | H2-T22 | Extracellular | Signal peptide | 0.178 | 0.1338 | 0.80019999 |

| | | | | | | |
|---|---|---|---|---|---|---|
| Hepatocyte growth factor | Hgf | Extracellular | Signal peptide | 0.1688 | 0.0956 | 0.93900001 |
| Insulin-degrading enzyme | Ide | Extracellular | Peroxisomal ta | 0.3195 | 0.1324 | 0.59850001 |
| Interferon alpha/beta receptor 2 | Ifnar2 | Extracellular\|C | Signal peptide | 0.193 | 0.1072 | 0.6947 |
| Insulin-like growth factor-binding protein 5 | Igfbp5 | Extracellular | Signal peptide | 0.1519 | 0.1134 | 0.95230001 |
| Insulin-like growth factor-binding protein 7 | Igfbp7 | Extracellular | Signal peptide | 0.2192 | 0.1201 | 0.88050002 |
| Immunoglobulin kappa constant | Igkc | Extracellular | Signal peptide | 0.421 | 0.0811 | 0.94590002 |
| Interleukin-31 receptor subunit alpha | Il31ra | Extracellular\|C | Signal peptide | 0.2918 | 0.18 | 0.7001 |
| Interleukin-34 | Il34 | Extracellular | Signal peptide | 0.2906 | 0.1523 | 0.73100001 |
| Inter-alpha-trypsin inhibitor heavy chain H5 | Itih5 | Extracellular | Signal peptide | 0.2524 | 0.0968 | 0.88660002 |
| Vascular endothelial growth factor receptor 2 | Kdr | Extracellular | Signal peptide | 0.1962 | 0.139 | 0.84359998 |
| Laminin subunit beta-1 | Lamb1 | Extracellular | Signal peptide | 0.3128 | 0.0989 | 0.89279997 |
| Laminin subunit beta-2 | Lamb2 | Extracellular | Signal peptide | 0.3406 | 0.1769 | 0.85369998 |
| Laminin subunit gamma-3 | Lamc3 | Extracellular | Signal peptide | 0.3204 | 0.1443 | 0.84560001 |
| Galectin-1 | Lgals1 | Cytoplasm\|Extracellular | | 0.7546 | 0.2794 | 0.73030001 |
| Galectin-3-binding protein | Lgals3bp | Extracellular | Signal peptide | 0.3061 | 0.1768 | 0.65359998 |
| Alpha-2-macroglobulin receptor-associated protein | Lrpap1 | Extracellular\|E | Signal peptide | 0.1621 | 0.1504 | 0.81870002 |
| Latent-transforming growth factor beta-binding protein 1 | Ltbp1 | Extracellular | Signal peptide | 0.2477 | 0.2357 | 0.94220001 |
| MAM domain-containing protein 2 | Mamdc2 | Extracellular | Signal peptide | 0.2953 | 0.1646 | 0.92119998 |
| Lactadherin | Mfge8 | Extracellular | Signal peptide | 0.2149 | 0.1066 | 0.88239998 |
| Multimerin-2 | Mmrn2 | Extracellular | Signal peptide | 0.264 | 0.1541 | 0.86119998 |
| C-type mannose receptor 2 | Mrc2 | Extracellular | Signal peptide | 0.2128 | 0.1342 | 0.85869998 |
| Beta-nerve growth factor | Ngf | Extracellular | Signal peptide | 0.0715 | 0.1111 | 0.97039998 |
| Nidogen-1 | Nid1 | Extracellular | Signal peptide | 0.3254 | 0.1131 | 0.90429997 |
| Neurogenic locus notch homolog protein 3 | Notch3 | Extracellular | Signal peptide | 0.3297 | 0.4178 | 0.4596 |
| NPC intracellular cholesterol transporter 2 | Npc2 | Extracellular | Signal peptide | 0.1348 | 0.042 | 0.91109997 |
| Mimecan | Ogn | Extracellular | Signal peptide | 0.1247 | 0.0744 | 0.8696 |
| Noelin-2 | Olfm2 | Extracellular | Signal peptide | 0.2945 | 0.0827 | 0.86369997 |
| Protein OS-9 | Os9 | Extracellular | Signal peptide | 0.2504 | 0.132 | 0.6929 |
| Protocadherin 1 | Pcdh1 | Extracellular | Signal peptide | 0.2051 | 0.292 | 0.56040001 |
| Procollagen C-endopeptidase enhancer 1 | Pcolce | Extracellular | Signal peptide | 0.1542 | 0.0915 | 0.91170001 |
| Platelet-derived growth factor subunit B | Pdgfb | Extracellular | Signal peptide | 0.1804 | 0.1458 | 0.91710001 |
| Platelet endothelial aggregation receptor 1 | Pear1 | Extracellular | Signal peptide | 0.2551 | 0.1137 | 0.84909999 |
| Phospholipase A1 member A | Pla1a | Extracellular | Signal peptide | 0.2154 | 0.1227 | 0.8725 |

| | | | | | | |
|---|---|---|---|---|---|---|
| Secretory phospholipase A2 receptor | Pla2r1 | Extracellular | Signal peptide | 0.2276 | 0.111 | 0.85100001 |
| Urokinase-type plasminogen activator | Plau | Extracellular | Signal peptide | 0.1601 | 0.1011 | 0.96200001 |
| Procollagen-lysine,2-oxoglutarate 5-dioxygenase 1 | Plod1 | Extracellular\|Er | Signal peptide | 0.1733 | 0.0632 | 0.72549999 |
| Phospholipid transfer protein | Pltp | Extracellular | Signal peptide | 0.2218 | 0.1087 | 0.82370001 |
| Periostin | Postn | Extracellular | Signal peptide | 0.1863 | 0.057 | 0.8793 |
| Peptidyl-prolyl cis-trans isomerase B | Ppib | Extracellular\|Er | Signal peptide | 0.1825 | 0.0837 | 0.69599998 |
| Palmitoyl-protein thioesterase 1 | Ppt1 | Extracellular | Signal peptide | 0.1234 | 0.1124 | 0.79589999 |
| Prolargin | Prelp | Extracellular | Signal peptide | 0.193 | 0.1255 | 0.80690002 |
| Vitamin K-dependent protein S | Pros1 | Extracellular | Signal peptide | 0.2513 | 0.0839 | 0.96329999 |
| Prosaposin | Psap | Extracellular | Signal peptide | 0.1309 | 0.0511 | 0.96619999 |
| Retinoic acid receptor responder protein 2 | Rarres2 | Extracellular | Signal peptide | 0.1141 | 0.1423 | 0.92159998 |
| Retinol-binding protein 4 | Rbp4 | Extracellular | Signal peptide | 0.1754 | 0.1108 | 0.9156 |
| Resistin | Retn | Extracellular | Signal peptide | 0.1796 | 0.0702 | 0.9073 |
| Ribonuclease 4 | Rnase4 | Extracellular | Signal peptide | 0.1178 | 0.1133 | 0.89429998 |
| Stromal cell-derived factor 2 | Sdf2 | Extracellular\|Er | Signal peptide | 0.1378 | 0.108 | 0.69919997 |
| 45 kDa calcium-binding protein | Sdf4 | Extracellular | Signal peptide | 0.2324 | 0.1388 | 0.7026 |
| Selenoprotein P | Selenop | Extracellular | Signal peptide | 0.2319 | 0.1043 | 0.90219998 |
| Glia-derived nexin | Serpine2 | Extracellular | Signal peptide | 0.1694 | 0.0483 | 0.84509999 |
| Pigment epithelium-derived factor | Serpinf1 | Extracellular | Signal peptide | 0.1864 | 0.0612 | 0.9321 |
| Plasma protease C1 inhibitor | Serping1 | Extracellular | Signal peptide | 0.1847 | 0.0832 | 0.89740002 |
| Serpin H1 | Serpinh1 | Extracellular | Signal peptide | 0.1662 | 0.0792 | 0.9188 |
| Slit homolog 3 protein | Slit3 | Extracellular | Signal peptide | 0.2025 | 0.1543 | 0.93059999 |
| Cyclic GMP-AMP phosphodiesterase SMPDL3A | Smpdl3a | Extracellular | Signal peptide | 0.2267 | 0.0567 | 0.92259997 |
| DAN domain family member 5 | Dand5 | Extracellular | Signal peptide | 0.1474 | 0.1175 | 0.92210001 |
| SPARC | Sparc | Extracellular | Signal peptide | 0.1777 | 0.0689 | 0.91350001 |
| Spondin-1 | Spon1 | Extracellular | Signal peptide | 0.1757 | 0.1096 | 0.921 |
| Serglycin | Srgn | Extracellular | Signal peptide | 0.0926 | 0.0647 | 0.91009998 |
| Tapasin | Tapbp | Extracellular | Signal peptide | 0.3189 | 0.1371 | 0.61290002 |
| ADAMTS-like protein 2 | Adamtsl2 | Extracellular | Signal peptide | 0.1843 | 0.1276 | 0.95069999 |
| Thrombospondin-1 | Thbs1 | Extracellular | Signal peptide | 0.1868 | 0.0774 | 0.92210001 |
| Thrombospondin-2 | Thbs2 | Extracellular | Signal peptide | 0.2039 | 0.0825 | 0.93300003 |
| Metalloproteinase inhibitor 3 | Timp3 | Extracellular | Signal peptide | 0.1205 | 0.1096 | 0.93839997 |
| Metalloproteinase inhibitor 4 | Timp4 | Extracellular | Signal peptide | 0.1904 | 0.1331 | 0.90859997 |

| | | | | | | |
|---|---|---|---|---|---|---|
| Tubulointerstitial nephritis antigen-like | Tinagl1 | Extracellular | Signal peptide | 0.1889 | 0.1318 | 0.8926 |
| Insulin-like growth factor-binding protein 3 receptor | Tmem219 | Extracellular | Signal peptide | 0.3842 | 0.3055 | 0.58029997 |
| Transmembrane protein 273 | Tmem273 | Extracellular | Signal peptide | 0.1686 | 0.1582 | 0.78240001 |
| Tenascin X | Tnxb | Extracellular | Signal peptide | 0.2504 | 0.122 | 0.92940003 |
| Serotransferrin | Tf | Extracellular | Signal peptide | 0.2249 | 0.0516 | 0.88319999 |
| Vitronectin | Vtn | Extracellular | Signal peptide | 0.183 | 0.1548 | 0.90869999 |

**Extended Table 7**

| DEG in endothelial senotype compared to young endothelial nuclei within the monoatomic ion channel activity gene ontology term GO:0005216 | | | | | |
|---|---|---|---|---|---|
| **Gene** | **Mean_log2FC** | **p_val_adj** | **Mean_pct_1** | **Mean_pct_2** | |
| Abcc9 | 9.412 | 7.12E-26 | 0.817 | 0 | Regulatory subunit of ATP-sensitive potassium channel (subunit 9) |
| Cacna1c | 4.261 | 1.80E-25 | 0.904 | 0.077 | Voltage-dependent L-type calcium channel (subunit a1C) |
| Kcnq5 | 4.521 | 3.26E-19 | 0.769 | 0.067 | Voltage-gated potassium channel |
| Ano1 | 2.705 | 7.01E-16 | 0.76 | 0.077 | Calcium-activated chloride channel |
| Gjc1 | 3.010 | 7.53E-15 | 0.673 | 0.038 | Gap junction protein |
| Cacna2d1 | 1.997 | 1.61E-12 | 0.808 | 0.154 | Voltage-dependent calcium channel subunit alpha-2/delta-1 |
| Trpc3 | 2.789 | 6.89E-11 | 0.567 | 0.038 | Transient receptor potential cation channel (subunit C3) |
| Tmbim6 | 1.325 | 8.03E-10 | 0.846 | 0.183 | regulates calcium; anti-apoptotic; BAX inhibitor |
| Tmbim1 | 1.931 | 8.56E-09 | 0.519 | 0.038 | Bax inhibitor |
| Slc24a3 | 1.100 | 4.62E-08 | 0.837 | 0.231 | Potassium-dependent sodium/calcium exchanger |
| Kcnq4 | 3.945 | 8.59E-08 | 0.433 | 0.019 | Potassium voltage-gated channel (subunit Q4) |
| Ghitm | 2.552 | 1.95E-07 | 0.471 | 0.038 | Bax inhibitor |
| Kcnj8 | 6.451 | 1.52E-06 | 0.346 | 0 | ATP-sensitive potassium channel (Kir6.1) |
| Tmco1 | 2.483 | 1.88E-06 | 0.413 | 0.029 | Transmembrane and coiled-coil domains 1 |
| Clic1 | 0.936 | 1.75E-05 | 0.51 | 0.077 | Chloride intracellular channel 1 |
| Tmem109 | 1.850 | 3.21E-05 | 0.433 | 0.048 | Voltage-gated monoatomic cation channel permeable to K+ and Ca+2 |
| Trpv2 | 5.501 | 0.00020 | 0.279 | 0 | Transient receptor potential cation channel (subunit V2) |
| Grina | 1.994 | 0.00076 | 0.337 | 0.029 | Glutamate ionotropic receptor NMDA type associated protein 1 |
| Vdac2 | 1.182 | 0.00079 | 0.462 | 0.087 | Voltage-dependent anion channel 2 |
| Kcna5 | 4.874 | 0.00148 | 0.25 | 0 | Potassium voltage-gated channel (subunit A5) |
| Aqp1 | 1.879 | 0.00178 | 0.644 | 0.279 | Aquaporin1 |
| P2rx4 | 2.013 | 0.00279 | 0.317 | 0.029 | Purinoceptor 4; ATP-gated ion channel |
| Trpc1 | 2.017 | 0.00302 | 0.317 | 0.029 | Transient receptor potential cation channel (subunit C1) |
| Vdac1 | 0.934 | 0.00579 | 0.423 | 0.077 | Transient receptor potential melastatin 7 calcium channel |
| Trpm7 | 0.671 | 0.01042 | 0.846 | 0.298 | Transient receptor potential cation channel (subfamily M7) |
| Cacnb2 | 2.129 | 0.01076 | 0.413 | 0.096 | Voltage-dependent calcium channel (subunit b2) |
| Tmem175 | 0.651 | 0.04698 | 0.337 | 0.058 | Transmembrane protein 175 |
